# Genotype-Informed Interaction Landscape of the Dengue Envelope Glycoprotein with Host Cellular Proteins: Structural Dynamics, Thermodynamic Cooperativity, and Binding Specificity

**DOI:** 10.64898/2026.05.31.728572

**Authors:** Daniel Ferreira de Lima Neto, Juan Philippe Teixeira, Marcus Alexandre Finzi Corat, Miklos Maximiliano Bajay, Gustavo Henrique Goulart Trossini, Daniel Santos Mansur, Elcio de Souza Leal, Luiz Mario Ramos Janini, Caio Cesar de Melo Freire

## Abstract

Dengue virus (DENV) entry has historically been viewed as a receptor-centric process in which viral envelope proteins engage isolated host factors to initiate infection. However, mounting evidence reveals that viral attachment, internalization, and intracellular trafficking occur within complex molecular environments where multiple interacting host proteins shape the infection landscape. Here we introduce vectorial host interaction fields—a framework that represents intermolecular contacts as directional vectors embedded within host protein interaction networks, providing structural and systems-level insight into viral entry.

Docking analyses were performed across sixteen dengue envelope genotype variants, whereas molecular dynamics and MM/GBSA analyses were conducted on representative complexes selected from this genotype-informed screening. We characterized dengue envelope interactions with thirteen host proteins involved in membrane attachment, receptor signaling, cytoskeletal transport, vesicular trafficking, proteostasis, and immune regulation. Our multiscale approach integrated protein–protein docking, 200 ns molecular dynamics simulations of representative complexes, MM/GBSA binding free energy analysis, and vectorial hydrogen-bond formalism encoding orientation, persistence, and angular entropy. We identified four recurrent viral interface architectures: multivalent anchoring (BiP/GRP78, Betaglycan), electrostatic sliding (Glypican-1), focal regulation (Claudin-1, SUMO), and conserved cytoskeletal modules (Actin, Rab5). Ternary docking analyses revealed that viral stability is strongly conditioned by local network context, with conditional ΔΔ*G* values ranging from approximately −3.4 to +6.9 kcal mol^−1^ depending on neighboring proteins and assembly sequence.

These findings support a systems-level model of dengue entry driven by layered host interaction fields rather than a single dominant receptor, but they should be interpreted as computational structural hypotheses requiring biochemical, biophysical, and cellular validation. Detailed structural datasets, molecular dynamics trajectories, vectorial analyses, and per-residue MM/GBSA decomposition are provided in the companion Supplementary Material, which includes extended Results and Discussion, comprehensive Limitations assessment, and VectorPROT pipeline documentation supporting these findings.

## 2 Introduction

Dengue virus (DENV), a mosquito-borne flavivirus, causes approximately 390 million infections annually, with nearly 100 million symptomatic cases worldwide, making it one of the most significant global public health challenges among arboviruses [1]. Belonging to the family *Flaviviridae* alongside other medically important pathogens such as Zika, West Nile, and Yellow fever viruses, flaviviruses share a defining structural feature: a lipid envelope bearing the envelope glycoprotein (E). This multifunctional protein mediates viral attachment, receptor recognition, and membrane fusion during cell entry [2,3].

On the virion surface, DENV envelope proteins form antiparallel dimers that undergo major conformational rearrangements during infection. Upon receptor engagement and endocytosis, acidification in early endosomes triggers a structural transition where dimers reorganize into fusogenic trimers, enabling the viral membrane to fuse with the host endosomal membrane and release the viral genome into the cytoplasm [3]. Given its central role in viral entry, tropism, and immunogenicity, the E protein has become the primary focus of structural, immunological, and antiviral investigations [3].

Dengue entry is a complex, multifactorial process that exploits diverse host proteins rather than relying on a single receptor. Heparan sulfate proteoglycans are well-supported dengue attachment platforms [4]; in this study, Glypican-1 was modeled as a representative heparan sulfate proteoglycan platform potentially contributing to initial electrostatic attachment rather than as a dengue receptor established independently of its sulfated glycosaminoglycan context. GRP78/BiP, an endoplasmic reticulum chaperone capable of surface translocation under stress, directly interacts with the envelope protein to promote viral entry [5,6]. Cytoskeletal components, particularly actin, drive membrane dynamics, endocytosis, and vesicular transport—experimental disruption of actin dynamics and Rac1 signaling significantly impairs dengue entry and productive infection [7,8]. Following internalization, the GTPase Rab5 directs transport through early endosomes, regulating vesicle fusion and endosomal maturation [1].

Additional host factors relevant to dengue biology include Neuropilin-1 (NRP1), a multifunctional receptor involved in angiogenesis and vascular permeability whose direct role in dengue entry remains a computational hypothesis in this work rather than an established DENV mechanism [9]; the TGF-*β* receptor system (Betaglycan/TGFBR3 and TGFBR1), whose elevated signaling has been observed in dengue patients; and tight junction proteins such as Claudin-1, for which dengue prM/M-dependent entry has been experimentally reported [10]. Immune regulatory components including the NLRP3 inflammasome, the SUMOylation machinery, and the NK activating receptor NKp44 connect structural perturbations by dengue envelope to downstream innate immune responses and are supported by dengue-specific evidence for EIII-driven NLRP3 activation, DENV E–Ubc9 interaction, and NKp44 recognition of flaviviral envelope glycoproteins [11–13].

The diversity of host factors involved—spanning receptors, cytoskeletal scaffolds, chaperones, trafficking regulators, and immune sensors—indicates that simple receptor-binding models cannot fully explain dengue entry. Instead, viral attachment occurs within local molecular environments shaped by protein interactions, membrane organization, and cellular signaling. A systems-level understanding therefore requires integrating structural analysis with network-level context.

This study presents a vectorial interaction framework that encodes intermolecular contacts as directional vectors characterized by orientation, persistence, and angular entropy. The VectorPROT pipeline (detailed in the Supplementary Material, Sections 3.7–3.8) integrates protein–protein docking, molecular dynamics simulations, MM/GBSA energetics, and vectorial hydrogen-bond analysis across sixteen envelope genotypes and thirteen host proteins. This approach interprets structural interactions within their native cellular networks using a conditional energetic framework (AB/BC/ABC) that reveals how local protein interactions modulate viral stability. Under this framework, dengue entry emerges as a systems property of host cellular organization rather than a consequence of isolated receptor binding.

## 3 Materials and Methods

### 3.1 Sequence Retrieval and Structural Dataset Construction

Amino acid sequences for all thirteen host proteins — Actin, Betaglycan (TGFBR3), BiP/GRP78, Claudin-1, Clusterin, Glypican-1, Neuropilin-1, NKp44, NLRP3, Rab5, SUMO, TGFBR1, and VEGF — were retrieved from the UniProt Knowledgebase (UniProtKB), selecting the reviewed Swiss-Prot entry corresponding to *Homo sapiens* for each target. Full details of the sequence retrieval procedure, including UniProt accession numbers for each target, are provided in the Supplementary Material (Section 3.1, line 256).

Three-dimensional structures were obtained through a hierarchical pipeline: (i) experimentally resolved structures from the Protein Data Bank (PDB), prioritized by crystallographic resolution, structural completeness, and absence of extensive disordered regions; (ii) deep-learning models from AlphaFold [14], evaluated by pLDDT confidence score; (iii) template-based models generated with I-TASSER 5.2 [15], evaluated by C-score; and (iv) comparative refinement with MODELLER [16], selecting models with the lowest DOPE energy and most favorable Ramachandran statistics. The complete structural acquisition pipeline, AlphaFold model evaluation criteria (Section 3.3, line 284), I-TASSER threading parameters (Section 3.4, line 295), MODELLER refinement settings (Section 3.5, line 319), and Ramachandran validation results (Section 3.2, line 306) are detailed in the Supplementary Material.

All structural models underwent stereochemical validation using the PDBsum Generate pipeline and Ramachandran plot analysis, with high-quality models exhibiting >90% of residues in favored regions. Models requiring further optimization were subjected to molecular dynamics relaxation in GROMACS [17] using an established biomolecular force field, with explicit solvent, energy minimization, and sequential NVT/NPT equilibration prior to short production runs. MD relaxation protocols, including force field selection, explicit solvent parameters, and NVT/NPT equilibration settings, are described in the Supplementary Material (Section 3.6, line 330).

Sixteen structural variants representing distinct dengue envelope (ENV) genotypes were selected to capture the genetic diversity of circulating DENV1–4 populations, with detailed genotype dataset construction described in the Supplementary Material (Section 3.7, line 344). including intraserotype genotypic variants, using the same modeling and refinement pipeline. The genotype selection rationale and the full genotype dataset are described in Section 3.8 of the Supplementary Material.

### 3.2 Protein–Protein Docking

Protein–protein docking simulations were performed using the HDOCK platform [18], a hybrid algorithm integrating template-based and *ab initio* docking strategies. Each of the sixteen ENV genotypes was docked against each of the thirteen host proteins, generating a comprehensive docking matrix of all viral–host combinations. The highest-ranked conformations were retained for subsequent interaction analysis. Full docking parameters, ranking criteria, and the rationale for using HDOCK in the context of large-scale host– pathogen interaction screening are detailed in Section 3.9 of the Supplementary Material.

### 3.3 Vectorial Characterization of Hydrogen Bond Networks (VectorPROT)

To quantify the spatial organization of intermolecular contacts, hydrogen bond interactions were represented using a vector-based formalism encoding direction, magnitude, persistence, and angular variability. Complete mathematical formalization, parameter definitions, and implementation details are provided in the Supplementary Material (Sections 3.10–3.11, lines 460–680).

For each interacting residue pair (*i, j*), the hydrogen bond interaction vector was defined as:

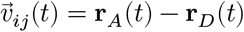

where **r**_*A*_ and **r**_*D*_ are the spatial coordinates of acceptor and donor atoms, respectively. Normalized interaction vectors preserve only the directional component. Interaction persistence was quantified through an occupancy parameter, and directional variability through angular entropy. Each viral–host interface was represented as a vector interaction network where residues are nodes and hydrogen bond interactions are edges characterized by mean vector orientation, occupancy, and angular entropy (Supplementary Material, Section 3.11, line 564).

#### 3.3.1 Circular Statistics for Directional Coherence Classification

To classify interaction regimes from the directional distribution of hydrogen bond vectors, circular statistics were computed for each interface independently for the envelope and host protein chains. Complete methodology including interpretation of circular metrics is provided in the Supplementary Material (Section 3.12, lines 564–680).

**Mean Resultant Length (R):** computed as the magnitude of the vector mean of all normalized interaction vectors over the trajectory:

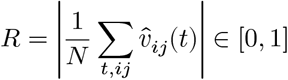

where *R* → 1 indicates perfect directional coherence (all vectors aligned) and *R* → 0 indicates isotropic angular dispersion. Anchoring interfaces such as BiP and Betaglycan show high directional coherence (*R* ≈ 0.72– 0.81), reflecting geometrically locked interaction vectors with low inter-frame variability. Electrostatic sliding interfaces such as GPC1 display markedly lower values (*R* ≈ 0.31–0.42), consistent with quasi-isotropic sampling of contact vectors as the viral particle explores the proteoglycan surface. Focal regulatory interfaces (NRP1, SUMO) occupy an intermediate regime (*R* ≈ 0.55–0.68), reflecting directionally constrained but conformationally adaptive recognition. Multi-macrostate exploration interfaces (TGFBR1) present the lowest coherence in the dataset, consistent with continuous angular resampling throughout the trajectory.

**Normalized Circular Entropy (***H*_**norm**_): defined as:

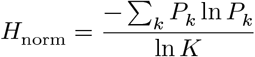

ln *K* where *K* is the number of orientation bins, scaling entropy to [0, 1] independently of bin resolution. *H*_norm_ → 0 denotes a near-deterministic directional distribution; *H*_norm_ → 1 denotes maximal angular randomness. Anchoring interfaces exhibit *H*_norm_ ≈ 0.18–0.27; sliding interfaces exhibit *H*_norm_ ≈ 0.61–0.74; and multi-macrostate systems such as TGFBR1 exhibit the highest values in the dataset (*H*_norm_ ≈ 0.78–0.85), reflecting continuous sampling of new vector orientations as the envelope explores the kinase surface without convergence. Together, *R* and *H*_norm_ provide an objective, quantitative basis for classifying interfaces into the four functional categories described in Section 4.8.

#### 3.3.2 Quadratic Surface Fitting as a Macroscopic Interface Descriptor

To characterize the global geometry of each vector field independently of local residue-level fluctuations, quadratic surfaces were fitted to the spatial coordinates of hydrogen bond vector tips (**r**_*A*_) across the trajectory ensemble for each genotype variant:

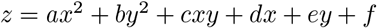

The fitted quadratic surface provides a macroscopic descriptor of the interaction field geometry through three interpretable parameters: (i) the mean curvature (*a* + *b*)/2, which distinguishes concave (cup-shaped, anchoring) from convex (dome-shaped, sliding) vector fields; (ii) the principal curvature anisotropy |*a* − *b*|, which quantifies directional bias in interface geometry; and (iii) the surface normal orientation, which provides a genotype-resolved descriptor of the dominant contact axis. Serotype-resolved comparison of fitted surface parameters reveals that curvature descriptors are broadly concordant across genotypic clusters within each serotype, confirming that dominant interaction geometry is encoded at the serotype level. Modest but detectable inter-serotype differences in principal curvature anisotropy — particularly at the DIII surface for GPC1 — represent testable structural hypotheses for serotype-specific differences in attachment efficiency.

### 3.4 Ternary Complex Analysis and Thermodynamic Cooperativity

To approximate cellular interaction environments, first-neighbor proteins for each host factor were identified through cross-referencing UniProt annotations with STRING, BioGRID, and IntAct, retaining only experimentally supported interactions. Three interaction scenarios were defined for ternary docking analysis:

| Scenario | Chain Pair | Biological Description |
| --- | --- | --- |
| AB | Virus (A) ↔ Host protein (B) | Direct viral–host protein contact |
| BC | Host protein (B) ↔ First neighbor (C) | Host protein engaged with its PPI-network neighbor |
| ABC | AB and BC simultaneously | Cooperative recruitment of the first neighbor |

Binding cooperativity was evaluated through ΔΔ*G* thermodynamic analysis:

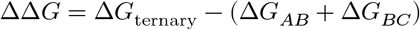

Negative ΔΔ*G* values indicate cooperative stabilization; positive values indicate competitive destabilization by the network neighbor. The complete rationale for the triple-docking design, the neighbor identification procedure, the data preprocessing and thermodynamic standardization pipeline, and the statistical analysis framework are detailed in Sections 3.10–3.14 of the Supplementary Material. The important limitations of this pairwise-decomposability approximation — including the inability to capture higher-order cooperative effects in simultaneous three-body contacts and the absence of post-docking MD relaxation for ternary assemblies — are discussed in the Limitations section of the Supplementary Material (Section 6.6).

## 4 Results

### 4.1 Functional Architecture of the Host Protein Network

#### 4.1.1 The Selected Host Proteins Form an Interconnected Signaling and Homeostasis Network

The thirteen host proteins selected for this study constitute an integrated cellular network in which signaling inputs, structural responses, and homeostatic mechanisms are functionally coupled across multiple layers of cellular organization (Supplementary Figure 2, *Functional interaction network of host proteins selected for envelope docking analysis*). The network can be organized into four functionally distinct but heavily interconnected clusters: (i) growth factor signaling and co-receptor complexes, (ii) intracellular trafficking and membrane dynamics, (iii) cell structure and adhesion, and (iv) protein homeostasis and inflammatory surveillance. Complete network topology analysis and functional characterization are provided in the Supplementary Material (Sections 3.1.1–3.1.2; Supplementary Figure 2, *Functional interaction network of host proteins selected for envelope docking analysis*).

The growth factor signaling cluster is anchored by two extracellular ligand systems — VEGF and TGF-*β* — whose activities are modulated by dedicated co-receptors before reaching canonical signaling receptors. Neuropilin-1 acts as a co-receptor that presents VEGF to VEGFR, potentiating pro-angiogenic and vascular permeability responses directly relevant to dengue-associated plasma leakage. Betaglycan (TGFBR3) presents TGF-*β* ligands to TGFBR1, enabling assembly of the canonical signaling receptor complex. Glypican-1 operates at the intersection of both ligand systems as a heparan sulfate proteoglycan capable of sequestering or presenting VEGF and FGF2, thereby fine-tuning the magnitude and spatial distribution of growth factor signaling at the cell surface.

The intracellular trafficking cluster is organized around Rab5 as the master regulator of early endosome identity and function. Rab5 GTPase activity controls vesicle docking, tethering, and fusion at the early endosomal compartment, directing the intracellular fate of internalized signaling receptors and viral particles following endocytosis. Actin provides the structural scaffold for this trafficking machinery through its roles in membrane deformation, vesicle budding, and intracellular cargo transport.

The homeostasis and inflammatory surveillance cluster monitors cellular stress and coordinates survival or death responses. BiP/GRP78 is the central sensor of the unfolded protein response (UPR), detecting misfolded protein accumulation induced by the high demand of viral protein production. Clusterin functions as an extracellular chaperone that clears aggregated proteins and directly interacts with BiP to suppress stress-induced apoptosis. NLRP3 operates downstream of UPR signals and perturbed actin dynamics as an intracellular danger sensor whose inflammasome activation drives the inflammatory pathology associated with dengue. SUMO post-translational modification regulates transcription factors within TGF-*β* and NF-*κ*B pathways, as well as the stability of NLRP3 itself.

#### 4.1.2 Network Topology Predicts Pleiotropic Consequences of Envelope Engagement

The network topology has direct implications for interpreting the biological significance of individual envelope–host protein interactions. Three network properties are particularly relevant. First, Actin, Rab5, and TGFBR1 function as connectivity hubs with disproportionate influence on network behavior, meaning that envelope engagement with any of these proteins could simultaneously perturb trafficking, structural, and signaling processes. Second, the network contains feedback loops whose disruption may produce non-linear inflammatory consequences: the BiP–Clusterin axis and the NLRP3–SUMO regulatory relationship are both feedback-stabilized circuits, and envelope engagement with either BiP or NLRP3 may contribute to inflammatory amplification consistent with the cytokine storm phenotype observed in severe dengue. Third, the network converges at Actin as a structural integration point receiving inputs from both signaling clusters and feeding outputs into trafficking, adhesion, and homeostasis.

### 4.2 Envelope–Actin Interface: Structural Stability and Cytoskeletal Tethering

Molecular dynamics simulations of the envelope–actin complex demonstrate rapid convergence to a stable conformational regime, with backbone RMSD values stabilizing near 0.4–0.6 nm without measurable drift or reduction in the radius of gyration of either partner. The continuous maintenance of atomic-scale interchain contacts — reflected in both minimum distance and contact number metrics — excludes the possibility that the observed interface represents a transient docking artifact. Complete RMSD/RMSF time-series data, PCA conformational ensembles, MM/GBSA decomposition plots, and VectorPROT vectorial field visualizations for the actin complex are provided in the Supplementary Material (Supplementary Figure 4, *Principal component analysis and RMSD-based hierarchical clustering of the dengue Envelope–actin molecular dynamics trajectory*; Supplementary Figure 5, *MM/GBSA energetic decomposition of the dengue virus Envelope–actin complex during molecular dynamics simulation*; Supplementary Figure 6, *Structural, interaction-field, and predicted energetic consequences of dengue Envelope–actin-centered ternary complexes with actin-associated proteins*).

Per-residue RMSF analysis reveals a characteristic asymmetric flexibility partitioning: the envelope protein functions as the comparatively rigid partner, maintaining a restricted and spatially consistent surface, while actin assumes the role of the conformationally adaptive host factor, with localized flexibility concentrated in interface-proximal subdomains. This partitioning — in which a viral protein presents a defined binding determinant to a plastically responsive host scaffold — is a recurrent organizational principle in virus–host cytoskeletal interactions.

MM/GBSA energetic analysis revealed an interface predominantly stabilized by electrostatic interactions, with gas-phase EEL contributions substantially offset by polar desolvation penalties, supplemented by van der Waals stabilization. Per-residue decomposition revealed a distributed energetic architecture spanning multiple spatially separated residue clusters on both partners rather than a single critical hotspot. For a viral envelope protein interacting with a highly conserved cellular scaffold such as actin, this energetic redundancy provides a structural basis for maintaining host factor engagement across sequence variants.

Ternary docking analyses indicate that the actin environment surrounding the envelope– actin interface is highly context dependent. Classical actin regulators such as *α*-actinin (ACTN2) and profilin (PFN1) produced destabilizing ΔΔ*G* effects consistent with competitive occupation of shared actin determinants, whereas dynactin-associated components (ACTR1A, ACTR10) showed patterns consistent with cooperative stabilization of the viral interface. ACTL6A (BAF53A), an actin-related component of the SWI/SNF chromatin-remodeling complex, emerged as the most prominent cooperative factor, generating a strongly negative ΔΔ*G* in the Viral–Cell Triple_Inverse configuration. These findings highlight that the ENV–actin complex is embedded within a dynamic network of cytoskeletal regulators that can either compete with or stabilize viral attachment depending on assembly order and local structural context.

### 4.3 Envelope–BiP/GRP78 Interface: Discrete Electrostatic Anchor and Stress-Chaperone Docking

The envelope–BiP complex exhibited global structural stability throughout the 200 ns trajectory, with neither partner undergoing measurable conformational drift. Full trajectory metrics, time-resolved contact analysis, and ternary network partner results (DNAJB11, HSPA5, FICD, DERL1, HYOU1, HSP90B1, PDIA3) are presented in the Supplementary Material (Supplementary Figure 10, *Principal component analysis and RMSD-based hierarchical clustering of the dengue Envelope–BiP molecular dynamics trajectory*; Supplementary Figure 11, *MM/GBSA energetic decomposition of the dengue virus Envelope–BiP complex during molecular dynamics simulation*; Supplementary Figure 12, *Structural, interaction-field, and predicted energetic consequences of dengue Envelope–BiP-centered ternary complexes with ER-associated BiP partners*). Per-residue RMSF analysis identified a specific electrostatic anchor involving A:ARG410 and B:GLU498, generating per-residue MM/GBSA contributions of approximately −3.1 to −3.4 kcal mol^−1^. This discrete anchor is supplemented by hydrophobic packing and distributed secondary contacts, yielding a total Δ*G* of approximately −70.5 kcal mol^−1^ (semi-quantitative MM/GBSA estimate).

The interfacial organization is characterized by electrostatic frustration at LYS435/LYS516/LYS447, where adjacent lysine residues in the nucleotide-binding domain generate competing favorable and unfavorable decomposed energies that may contribute to dynamic on-rate optimization. The interaction mode — stable yet dynamically permissive — is consistent with a co-receptor role in which cell-surface BiP increases viral residence time without irreversibly sequestering the particle. The identity of ARG410 as the primary anchor constitutes a specific, experimentally testable structural hypothesis.

### 4.4 Envelope–Glypican-1 Interface: Electrostatic Saturation and Dimensionality-Reduction Attachment

Glypican-1 generated the largest absolute electrostatic stabilization across all thirteen host proteins, with Δ*G*_total_ of approximately −90 kcal mol^−1^ and favorable per-residue contributions concentrated in the DIII electropositive patch — consistent with the established role of domain III as the primary heparan sulfate-binding site in flaviviruses. Full trajectory data, PCA ensemble analysis, per-residue decomposition identifying the DIII basic-residue hotspot cluster and peripheral frustration loci, and ternary results for HBEGF, FGF1, and FGF2 competitors are reported in the Supplementary Material (Supplementary Figure 17, *Principal component analysis and RMSD-based hierarchical clustering of the dengue Envelope–glypican-1 molecular dynamics trajectory*; Supplementary Figure 16, *MM/GBSA energetic decomposition of the dengue virus Envelope–glypican-1 complex during molecular dynamics simulation*; Supplementary Figure 18, *Structural, interaction-field, and predicted energetic consequences of dengue Envelope–glypican-1-centered ternary complexes with growth-factor-associated partners*). Both partners maintained stable global conformations throughout the trajectory with no inter-domain rearrangements, and the DIII region of the envelope displayed markedly reduced per-residue RMSF values relative to unliganded states, consistent with spatial constraint imposed by persistent electrostatic contacts.

The contact number rose rapidly during equilibration and thereafter stabilized at an elevated plateau without systematic decay, distinguishing the GPC1 interface from the betaglycan complex (which shows progressive contact growth) and suggesting that GPC1 achieves rapid electrostatic saturation and subsequently maintains it.

Ternary docking analysis incorporating GPC1 network neighbors revealed strong competitive displacement by HBEGF, consistent with a surf-riding model in which viral particles exploit the GPC1 heparan sulfate surface as a dimensionality-reduction platform for diffusion-assisted receptor search. The core protein component characterized here provides a baseline interaction energy that would be augmented by the full heparan sulfate-decorated proteoglycan in physiological conditions.

### 4.5 Envelope–Neuropilin-1 Interface: SLiM-Based Molecular Mimicry

The ENV–NRP1 complex displayed bilateral conformational restriction and energy concentration in a terminal basic residue–acidic cage arrangement qualitatively resembling characterized CendR–NRP1 complexes, supporting the hypothesis that the dengue envelope may engage NRP1 through SLiM-based molecular mimicry — a strategy documented across multiple viral families. The candidate CendR-like loop residue cluster was structurally present across all sixteen envelope variants examined, suggesting conservation of this interaction geometry across circulating DENV serotypes. A structural comparison with the SARS-CoV-2 NRP1 engagement geometry — revealing convergent b1-pocket geometry across hemorrhagic pathogens — and detailed analysis of the plexin-dominated ternary landscape (PLXNA1, PLXNA4, SEMA4A) are discussed in the Supplementary Material (Supplementary Material Sections 3.8 and 4.7; Supplementary Figure 20, *Principal component analysis and RMSD-based hierarchical clustering of the dengue Envelope– neuropilin-1 molecular dynamics trajectory*; Supplementary Figure 19, *MM/GBSA energetic decomposition of the dengue virus Envelope–neuropilin-1 complex during molecular dynamics simulation*; Supplementary Figure 21, *Structural, interaction-field, and predicted energetic consequences of dengue Envelope–neuropilin-1-centered ternary complexes with neuropilin-1-associated proteins*).

This hypothesis requires direct experimental confirmation through competition assays and mutagenesis of the candidate CendR-like loop. Cross-flavivirus NRP1 engagement profiling (Zika, West Nile, Yellow fever) is warranted to test whether b1-pocket engagement represents a shared flavivirus strategy.

### 4.6 Envelope–Betaglycan (TGFBR3) Interface: Interfacial Maturation and Avidity-Driven Tethering

The envelope–betaglycan complex exhibited monotonic contact number increase and concomitant reduction in mean interchain distance over the 200 ns trajectory — a structural signature of interfacial maturation in which local conformational exploration leads to progressive improvement of complementarity. Complete contact density evolution data, per-residue decomposition of the ARG107/LYS236 hotspot cluster, MM/GBSA profiles revealing the electrostatically driven binding with persistent polar frustration, and ternary results for the TGFBR1, TGFBR2, and FKBP1A network partners are provided in the Supplementary Material (Supplementary Figure 7, *Principal component analysis and RMSD-based hierarchical clustering of the dengue Envelope–betaglycan molecular dynamics trajectory*; Supplementary Figure 8, *MM/GBSA energetic decomposition of the dengue virus Envelope–betaglycan complex during molecular dynamics simulation*; Supplementary Figure 9, *Structural, interaction-field, and predicted energetic consequences of dengue Envelope–betaglycan-centered ternary complexes with betaglycan-associated partners*). The convergence of these metrics toward a tighter distributional regime in the final trajectory window indicates that the system accesses a conformational basin of greater stability than the initial docked configuration.

Per-residue decomposition identified an ARG107/LYS236 hotspot cluster on the Betaglycan surface, with corresponding energetic penalties at unfavorable charged residues reflecting electrostatic frustration at the interface periphery. The total Δ*G* is among the most favorable in the dataset, consistent with a high-avidity tethering function congruent with betaglycan’s established role as a co-receptor scaffold.

### 4.7 Envelope–Clusterin Interface: Hydrophobic Tail-Mediated Recognition

The envelope–clusterin complex presented an energetically distinct signature relative to betaglycan and BiP systems. Full PCA ensemble analysis confirming the asymmetric conformational landscape, per-residue decomposition identifying hydrophobic tail residues as primary contributors, and ternary docking results for BiP, HSPA8, and stress-context partners are reported in the Supplementary Material (Supplementary Figure 14, *Principal component analysis and RMSD-based hierarchical clustering of the dengue Envelope– clusterin molecular dynamics trajectory*; Supplementary Figure 13, *MM/GBSA energetic decomposition of the dengue virus Envelope–clusterin complex during molecular dynamics simulation*; Supplementary Figure 15, *Structural, interaction-field, and predicted energetic consequences of dengue Envelope–clusterin-centered ternary complexes with clusterin-associated proteins*). The gas-phase van der Waals term (VDWAALS) represented a proportionally larger share of Δ*G*_gas_ than in predominantly electrostatic profiles of other complexes, consistent with the hydrophobic character of the clusterin tail-mediated interface. The polar solvation penalty was comparatively attenuated, because partial — rather than full — burial of ionizable groups reduces the desolvation cost.

PCA of C*α* coordinates confirmed an asymmetric conformational landscape: the envelope protein underwent a single major conformational transition during the trajectory, while clusterin exhibited continuous multidirectional sampling reflecting the intrinsic multi-state dynamics of its disordered tail regions. Per-residue decomposition identified hydrophobic and amphipathic residues in the disordered tail as the primary contributors, with secondary favorable terms from the coiled-coil bundle and minimal contributions from the disulfide-stabilized domain. Crucially, the clusterin and BiP interaction epitopes on the envelope surface were non-overlapping, supporting the possibility of sequential or simultaneous chaperone engagement during viral entry.

### 4.8 Envelope–Rab5 Interface: Two-Registry Mechanism and Endosomal Trafficking Recruitment

The envelope–Rab5 complex presents the most mechanistically distinctive trajectory in the entire dataset. The global center-of-mass distance between chains exhibits a characteristic biphasic behavior — an initial plateau near approximately 2.81 nm during the first 20 ns, followed by a gradual transition spanning approximately 20–120 ns that leads to a new stable late-time plateau near approximately 3.11 nm — while the minimum interchain distance remains essentially constant at approximately 0.172 nm throughout the full 200 ns trajectory. This dissociation between global and local distance metrics is the defining structural signature of the Rab5 complex: the two proteins undergo a *rigid-body global reorientation* rather than partial dissociation, rotating relative to one another while maintaining a continuously anchored local contact set.

This behavior defines the *Two-Registry Model* for Rab5 engagement. In the early registry (0–20 ns), the complex initiates contact through a geometrically compact arrangement in which the molecular centroids are in closer proximity. In the late registry (120–200 ns), the complex pivots globally — the envelope reorients by approximately 15–20° relative to the Rab5 GTPase core — while anchor residues at the local interface remain engaged throughout. The molecular pivot identified in this complex is mechanistically consistent with the known architecture of Rab–effector interactions, which frequently tolerate global reorientations while maintaining focal contacts at switch I/switch II–proximal surfaces. Per-residue energetic decomposition reveals that the primary anchor on the Rab5 side involves the surface segment flanking the switch II domain, a region known to exhibit variable engagement geometry across different Rab5–effector complexes. This continuous anchoring behavior distinguishes the Rab5 complex from all other host factor systems in this study and suggests that the viral envelope can exploit the conformational plasticity of the Rab GTPase surface to achieve orientation-optimized productive endosomal recruitment.

Ternary docking analyses incorporating Rab5 regulatory network partners — GDI2, PIK3C3, RAB5B, RAB5C, and RABGEF1 — reveal a strikingly uniform pattern: all five partners destabilize the Rab5– envelope reference state, with no partner generating cooperative AB reinforcement. This uniformity is itself mechanistically informative, indicating that the Rab5–envelope interaction is systematically incompatible with the native Rab5 regulatory machinery and is therefore more likely to represent a transient trafficking-associated contact than a constitutive entry receptor engagement. These findings are consistent with the functional classification of Rab5 as a trafficking scaffold rather than a primary attachment factor. The mechanistic interpretation of this uniform antagonism — including the “endosomal window kinetics” and allosteric GTPase displacement model, and a discussion of Rab5 as a biological timer for viral endosomal routing — is developed in detail in the Supplementary Material (Supplementary Figure 23, *Principal component analysis and RMSD-based hierarchical clustering of the dengue Envelope–Rab5 molecular dynamics trajectory*; Supplementary Figure 22, *MM/GBSA energetic decomposition of the dengue virus Envelope–Rab5 complex during molecular dynamics simulation*; Supplementary Figure 24, *Structural, interaction-field, and predicted energetic consequences of dengue Envelope–Rab5-centered ternary complexes with Rab5-associated proteins*).

### 4.9 Envelope–SUMO/UBC9 Interface: Cooperative Induced-Fit and Packing-Dominant Recognition

The envelope–SUMO9-UBC1 complex exhibits the most extreme asymmetry in conformational behavior observed across all thirteen host protein systems. The SUMO9-UBC1 ligand displays exceptional structural rigidity, with backbone RMSD values stabilizing at approximately 0.22–0.24 nm within the first 20 ns and remaining within this narrow band throughout the full 200 ns trajectory — a behavior consistent with the conserved-grasp fold of SUMO proteins, whose hydrophobic core resists thermal fluctuation even in the absence of binding partners. In stark contrast, the dengue envelope protein undergoes a large and temporally structured conformational rearrangement: RMSD values increase from approximately 0.50 nm to a late-time plateau near approximately 0.73 nm, with the variance of RMSD values decreasing substantially during the final 100 ns. This transition from a high-variance early regime to a low-variance late regime represents a genuine conformational selection event in which the envelope enters a well-defined basin distinct from the initial docked geometry.

The defining feature of the SUMO complex is the mechanism of *Cooperative Induced Fit*. Unlike the Rab5 pivot (where global reorientation accompanies stable local anchoring) or the TGFBR1 system (where the envelope explores multiple macrostates), the SUMO interface convergence proceeds through two simultaneous and mutually reinforcing processes: (i) envelope-side compaction, in which the three-domain ectodomain architecture closes inter-domain degrees of freedom, reducing the radius of gyration by approximately 0.14 nm — the largest envelope Rg reduction observed across all complexes — and (ii) substate selection, in which the SUMO9-UBC1 ligand selects among accessible peripheral loop conformations to present a surface optimally complementary to the compacting envelope. Interface development proceeds through three kinetically identifiable phases: an initial heterogeneous exploration phase (0–20 ns), an intermediate reorganization plateau (20–100 ns), and a late-time densification phase (100–200 ns) in which contact number rises to approximately 4,622 with the lowest variance of the entire trajectory. Throughout all phases, the minimum interchain distance remains essentially constant at approximately 0.16–0.17 nm, confirming continuous interface engagement.

Per-residue RMSF and contact-gain analyses identify the envelope C-terminal stem-proximal region (residues ~464–471) as the primary adaptive interface element — the region simultaneously exhibiting the highest RMSF and the highest contact-gain ratio during the trajectory. This “flexible arm” model positions the envelope stem-proximal segment as a previously uncharacterized host-factor binding surface with potential functional relevance for SUMOylation axis co-option during infection, consistent with experimental evidence that the DENV-2 E protein co-immunoprecipitates with Ubc9 and that specific E protein lysines (K51, K241) are functional determinants of this interaction.

Energetically, the SUMO complex is characterized by a van der Waals-dominant interface architecture — a packing-dominant signature distinct from the electrostatic-dominant profiles of BiP, Betaglycan, and GPC1 — supplemented by localized electrostatic hotspots at the SIM-proximal groove. This energetic regime is consistent with a regulatory recognition mode in which the envelope engages the SUMO/Ubc9 machinery without disrupting its active-site geometry, a property that would permit virally induced modulation of SUMOylation targets while the enzymatic cascade remains functional. The hypothesis that the envelope acts as a non-competitive Ubc9 inhibitor through allosteric modulation of the catalytic cleft geometry — and the analysis of ternary partners spanning the full SUMOylation pathway (SAE1, UBA2, RANGAP1, IPO13, and SUMO paralogs) — is developed in the Supplementary Material (Supplementary Figure 26, *Principal component analysis and RMSD-based hierarchical clustering of the dengue Envelope–SUMO molecular dynamics trajectory*; Supplementary Figure 25, *MM/GBSA energetic decomposition of the dengue virus Envelope–SUMO complex during molecular dynamics simulation*; Supplementary Figure 27, *Structural, interaction-field, and predicted energetic consequences of dengue Envelope–SUMO-centered ternary complexes with SUMO-associated proteins*).

### 4.10 Envelope–TGFBR1 Interface: Multi-Macrostate Exploration Against a Structurally Stable Kinase Surface

The envelope–TGFBR1 complex presents the most extreme conformational behavior observed in this study. TGFBR1 maintains a structurally stable configuration throughout the 200 ns trajectory, with backbone RMSD rising only modestly from approximately 0.180 nm to approximately 0.224 nm — consistent with thermal relaxation of surface loops rather than global refolding of the kinase domain. The dengue envelope protein, by contrast, displays a sustained and monotonic RMSD increase across all four temporal windows (0.381, 0.490, 0.533, and 0.582 nm), without reaching a stable plateau anywhere within the simulation timescale.

This continuous drift without convergence is the defining feature of the *Multi-Macrostate Exploration* mechanism. Unlike the SUMO complex (where the envelope converges to a single well-defined basin after approximately 100 ns), the Rab5 complex (where a discrete pivot yields a stable late-time registry), or the NRP1 complex (where both partners are conformationally restricted), the TGFBR1 envelope explores a broad, heterogeneous conformational landscape against a structurally invariant kinase surface. Later-occupied conformational basins achieve higher interfacial contact densities — the TGFBR1 complex generates the highest absolute contact count observed across all host protein systems — but at the cost of sub-optimal electrostatic complementarity in transient late-stage geometries. The envelope simultaneously becomes more compact (radius of gyration decreasing from ~3.113 nm to ~3.049 nm) and more divergent from its initial structure, consistent with inter-domain rearrangements that redistribute internal contacts without expanding the molecular volume.

On the TGFBR1 side, a lysine/aspartate hotspot cluster on the extracellular face of the kinase domain provides an electrostatic scaffold that the continuously exploring viral ectodomain approaches with progressively improving geometric complementarity — a kinetic rather than thermodynamic optimization of the interface. This behavior is consistent with the structural polymorphism of flavivirus envelope proteins, whose multidomain architecture supports multi-stage receptor engagement and whose conformational landscape includes metastable intermediates accessible during normal virion dynamics. PCA confirms the absence of late-time convergence, with trajectory frames distributed broadly across conformational space throughout, contrasting with the compact, low-dispersion landscape of TGFBR1 itself.

The TGFBR1 system represents a distinct functional class of envelope–host factor engagement: one in which the viral protein acts as a continuous conformational explorer exploiting the stable geometric surface of a signaling receptor without committing to a single productive geometry — a mode of interaction that may contribute to TGF-pathway perturbation during dengue infection through transient, probabilistic rather than stoichiometric contacts. The energetic paradox arising from this system — in which accumulation of higher contact density yields progressively less favorable binding energy due to electrostatic incompatibility in late-stage geometries — is resolved and discussed in Section 5.9.2 of the Supplementary Material. The TGF-pathway modulation hypothesis, connecting envelope-mediated TGFBR1 perturbation to plasma leakage and endothelial barrier dysregulation in severe dengue, is developed in Section 5.9.3. Complete PCA trajectories, MM/GBSA profiles, and ternary partner data (TGFBR2, TGFB1, FKBP1A, SMAD2, SMAD3) are provided in the Supplementary Material (Supplementary Figure 29, *Principal component analysis and RMSD-based hierarchical clustering of the dengue Envelope–TGFBR1 molecular dynamics trajectory*; Supplementary Figure 28, *MM/GBSA energetic decomposition of the dengue virus Envelope–TGFBR1 complex during molecular dynamics simulation*; Supplementary Figure 30, *Structural, interaction-field, and predicted energetic consequences of dengue Envelope–TGFBR1-centered ternary complexes with TGFBR1-associated proteins*).

### 4.11 Comparative Ligand Dynamics: Distinct Regimes of Envelope Modulation

Across all thirteen host proteins, the dengue envelope does not behave as a passive binding scaffold. Instead, each ligand imposes a characteristic dynamic regime on the receptor, modulating backbone polar surface exposure, radius of gyration, RMSD drift, and residue-level RMSF in ligand-specific ways. The Supplementary Material presents a full comparative analysis of these descriptors — receptor-side and ligand-side — for all complexes simultaneously (Supplementary Figure 3, *Comparative molecular dynamics descriptors of dengue envelope–host ligand complexes*), revealing at least three dynamic classes: (i) ligands that stabilize the envelope near a compact, low-deviation state (consistent with structurally compatible anchoring), (ii) ligands that induce stronger receptor remodeling and expansion (consistent with induced-fit or conformational selection entry modes), and (iii) ligands with high intrinsic mobility whose global metrics overestimate instability because flexible distal regions dominate the signal while a stable interface core is maintained. The present vectorial analysis builds on this classification by identifying the directional coherence (*R*) and normalized circular entropy (*H*_norm_) signatures that distinguish each class at the vector-field level.

### 4.12 PRE versus POST Interaction States: Interface Capture

Comparison between basal host interaction states (PRE) and viral docking states (POST) revealed a recurring pattern across systems: dengue virus frequently exploits geometric configurations already accessible within host molecular networks rather than generating entirely new binding configurations. Viral docking consistently stabilizes interaction surfaces that are transiently exposed during normal cellular dynamics — configurations that already exist as metastable substates in the host protein’s native conformational ensemble.

This pattern — termed *interface capture* — is best interpreted as a computational mechanistic hypothesis: the dengue envelope may act as a *structural opportunist*, preferentially populating molecular geometries compatible with binding without requiring host protein refolding. Crucially, this strategy does not impose a thermodynamic refolding cost on the host protein, which means that the energetic barrier to initial contact formation is lower than it would be for a classical induced-fit mechanism. By targeting conformations already sampled in the host’s free energy landscape, the virus achieves rapid, productive engagement across a chemically diverse set of cellular proteins — GPC1, BiP, Betaglycan, Rab5, SUMO — without requiring structural specialization for each individual target.

The functional consequence of interface capture is that the DENV envelope is not recognized as a foreign structural element by the host interaction network; it is instead accommodated as a geometric mimic of transiently accessible host surface configurations. This deepens the interpretation of viral tropism: cellular permissiveness to dengue infection may reflect not only receptor expression levels but also the fraction of time that host proteins spend in capture-compatible substates — a property that would vary with cell type, activation state, and local molecular environment. The recurrence of interface-capture-like geometries across all four interaction regimes (anchoring, sliding, regulatory, trafficking) provides a unifying computational hypothesis to prioritize for experimental testing.

### 4.13 Hierarchical Architecture of Dengue Host Engagement

Integrating all pairwise and ternary interaction data, dengue host engagement proceeds through a four-stage hierarchical architecture:

1. *Electrostatic Capture* (GPC1): heparan sulfate-mediated dimensionality reduction at the cell surface, exploiting the DIII electropositive patch of the envelope to concentrate viral particles prior to productive receptor engagement.
2. *Anchoring Stabilization* (BiP, Betaglycan): high-avidity electrostatically dominated interfaces with discrete anchor residues (ARG410 in BiP; ARG107/LYS236 in Betaglycan) that extend viral residence time at the plasma membrane without irreversible capture.
3. *Regulatory Checkpoints* (NRP1, SUMO/UBC9): focal interfaces gating intracellular access and connecting viral docking to immune signaling cascades; NRP1 is consistent with a CendR-like pocket-engagement hypothesis requiring direct validation, while the SUMO/UBC9 axis suggests a possible link between envelope recognition and post-translational modification machinery through envelope compaction-driven induced fit.
4. *Cytoskeletal Trafficking* (Actin, Rab5): conserved trafficking modules directing internalized particles through productive endosomal pathways; the Rab5 complex is uniquely characterized by a Two-Registry rigid-body pivot mechanism, while actin engagement is regulated by the dynactin network context.

This integrated mechanistic hierarchy is summarized in Figures 1 and 2. The first figure places the envelope– host interaction landscape within the hierarchical modeling pipeline and genotype-informed mechanistic classes, whereas the second condenses the individual molecular dynamics signatures into a unified model of dengue envelope engagement across trafficking, regulatory-checkpoint, and capture-stabilization layers. The host protein interaction network that provides biological context for this hierarchy — including the four functional cluster organization (growth factor signaling, intracellular trafficking, cell structure and adhesion, homeostasis and inflammatory surveillance) and the network topology analysis identifying Actin, Rab5, and TGFBR1 as connectivity hubs — is described in the Supplementary Material (Sections 3.1.1– 3.1.2; Supplementary Figure 2, *Functional interaction network of host proteins selected for envelope docking analysis*).

**Figure 1:**
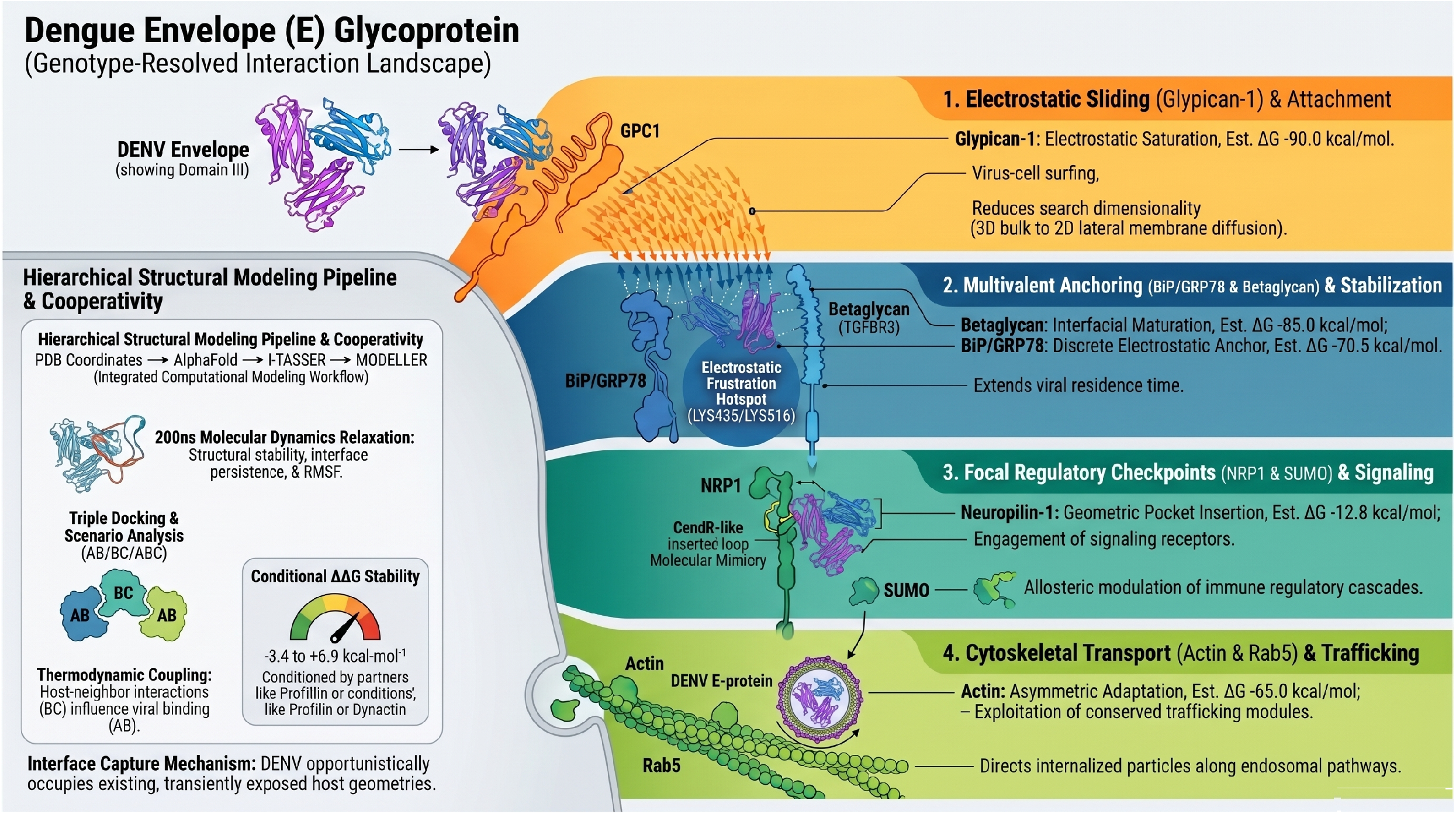
Genotype-informed interaction landscape of the dengue envelope glycoprotein. Integrative schematic of the hierarchical structural modeling and molecular dynamics framework used to classify dengue envelope engagement with host proteins across functionally distinct interaction regimes. The left panel summarizes the computational workflow, in which experimentally derived or predicted coordinates were refined through hierarchical structural modeling, AlphaFold- and I-TASSER-assisted reconstruction, molecular modeling, and 200 ns molecular dynamics relaxation, followed by triple-docking and scenario analyses of basal host complexes, viral docking states, and ternary host–virus assemblies. This workflow supports a cooperative interpretation of dengue entry, in which host–neighbor interactions can condition the stability of envelope binding, shifting conditional interaction energetics across favorable or unfavorable ranges depending on the local molecular context. The right panel organizes the resulting envelope–host landscape into four mechanistic layers. Glypican-1 is modeled as a representative heparan sulfate proteoglycan platform, in which the envelope may exploit sulfated proteoglycan surfaces to reduce the dimensionality of viral search from three-dimensional diffusion to two-dimensional lateral membrane exploration. Betaglycan/TGFBR3 and BiP/GRP78 define a multivalent anchoring and stabilization layer, extending viral residence time through interfacial maturation and discrete electrostatic anchoring, including charged hotspots that can also generate local electrostatic frustration. NRP1 and SUMO/Ubc9 define candidate focal regulatory checkpoints, coupling predicted geometric pocket insertion, CendR-like pocket-engagement resemblance, and possible allosteric modulation of immune-regulatory cascades to envelope recognition. Actin and Rab5 define a cytoskeletal transport and trafficking layer, linking asymmetric envelope adaptation to conserved intracellular transport modules and endosomal routing. Together, the figure frames dengue E not as a single-receptor ligand but as a genotype-informed, structurally opportunistic interaction hub that captures pre-existing host geometries, exploits electrostatic and steric complementarity, and converts local interface chemistry into higher-order trafficking, stabilization, and signaling outcomes.

**Figure 2:**
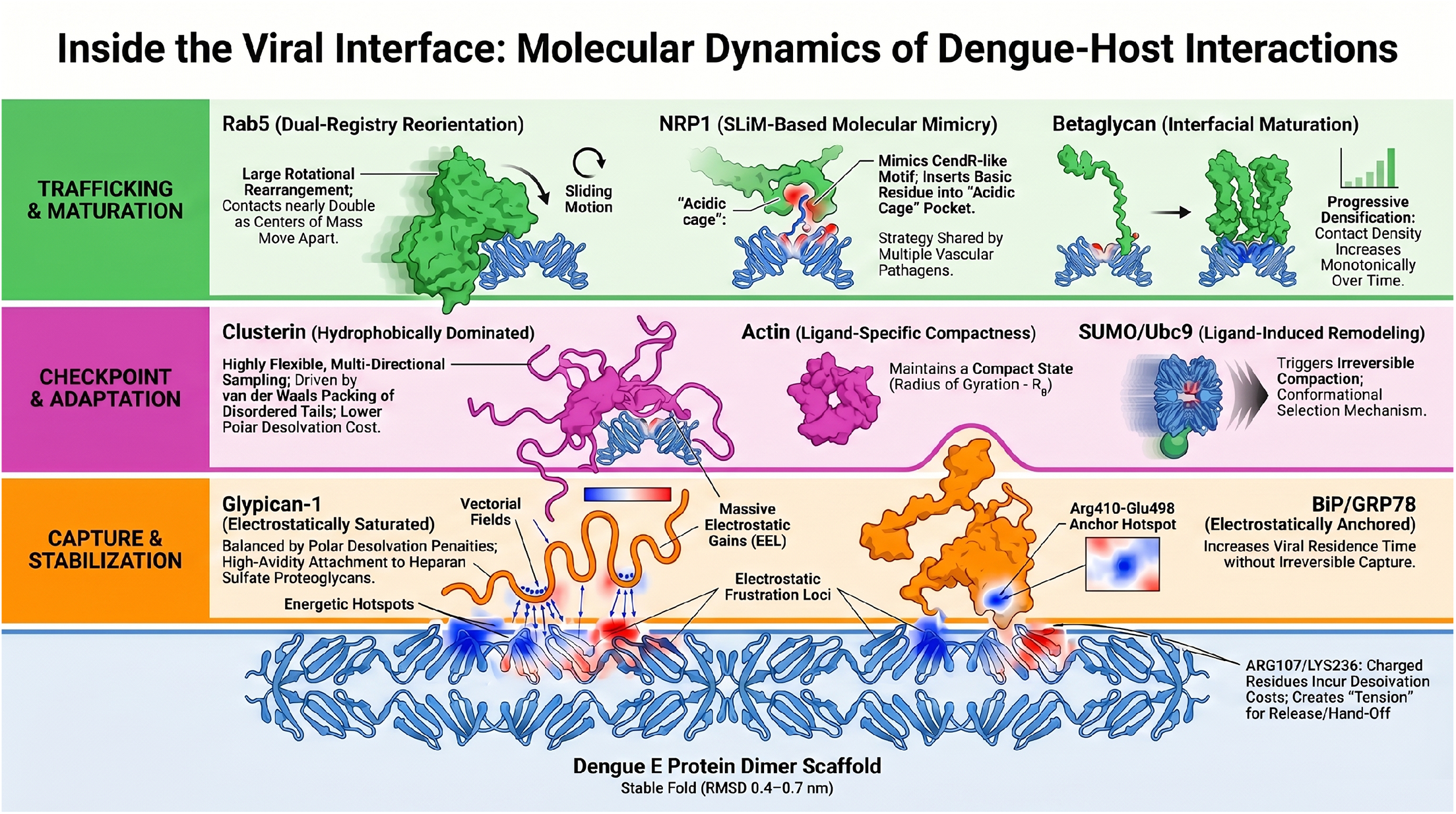
Integrated molecular dynamics model of dengue envelope–host protein interfaces. Schematic synthesis of the principal molecular dynamics regimes identified for dengue envelope engagement with host proteins. The dengue E protein dimer scaffold, shown as the conserved structural platform at the base of the model, remains globally stable across simulations while recruiting chemically and functionally distinct host factors through interface-specific mechanisms. Trafficking and maturation modules include Rab5, which undergoes a dual-registry reorientation with sliding motion and increased interfacial separation, and betaglycan/TGFBR3, which shows progressive contact densification consistent with interfacial maturation. Regulatory checkpoint and adaptive interfaces include NRP1, where an envelope basic element is positioned into an acidic pocket in a CendR-like molecular mimicry mode; clusterin, whose flexible hydrophobic tails support multi-directional sampling with comparatively lower polar desolvation cost; actin, which maintains a ligand-specific compact state; and SUMO/Ubc9, which promotes ligand-induced envelope remodeling and irreversible compaction. Capture and stabilization modules include glypican-1, dominated by electrostatic saturation and distributed vectorial fields over the DIII electropositive surface, and BiP/GRP78, characterized by an electrostatic anchor centered on the ARG410–GLU498 hotspot that increases viral residence time without requiring irreversible sequestration. Blue and red electrostatic fields denote favorable and unfavorable charge environments, respectively, highlighting how local energetic hotspots, desolvation penalties, and electrostatic frustration jointly define whether each interface behaves as a capture site, stabilizing anchor, regulatory checkpoint, or trafficking adaptor. Together, the figure presents dengue envelope recognition as a hierarchical, network-embedded process rather than as a single-receptor binding event. of interacting host proteins whose spatial organization determines which binding geometries become accessible. The concept of vectorial host interaction fields provides a quantitative framework for describing these environments, enabling integration of structural interaction data with systems-level molecular organization.

## 5 Discussion

### 5.1 Viral Entry as a Network-Embedded Structural Process

The principal conclusion of this study is that dengue virus entry cannot be adequately explained through single receptor interactions. Instead, viral docking occurs within structured molecular environments composed

### 5.2 Asymmetric Conformational Adaptation as a Cross-Interface Conserved Feature

Across all characterized complexes, molecular dynamics trajectories reveal a recurring pattern of *asymmetric conformational adaptation* in which the viral envelope protein functions as a structurally rigid scaffold — presenting defined binding determinants to the cellular environment — while host proteins absorb the thermodynamic cost of interface formation through localized flexibility rather than global refolding. Backbone RMSD values for the envelope protein typically converge to 0.4–0.7 nm over 200 ns, while host proteins exhibit greater conformational plasticity concentrated in interface-proximal subdomains. This partitioning is consistent with the established structural robustness of the flavivirus E protein, whose *β*-rich architecture confers conformational stability necessary for the pH-triggered fusion transition [2].

The asymmetry is not uniform but takes characteristic forms in each interaction class. In anchoring interfaces (BiP, Betaglycan), both partners converge to stable late-time geometries, with host protein flexibility localized to electrostatic accommodation loops. In the sliding interface (GPC1), the envelope maintains a rigid DIII electropositive patch while the proteoglycan surface permits lateral diffusion-like vector sampling. In the SUMO complex, the envelope undergoes active compaction — absorbing the induced-fit cost — while SUMO maintains its rigid *β*-grasp fold. In the Rab5 complex, the envelope drives a global pivot while maintaining local anchoring, with Rab5 acting as a geometrically tolerant tethering surface. In the TGFBR1 complex, the envelope assumes the role of continuous explorer against a kinase surface that provides structural invariance as the reference geometry.

This envelope-as-scaffold principle has a direct biological implication: the dengue particle does not require host protein refolding to engage a broad repertoire of cellular factors. Instead, it presents a conformationally pre-organized interaction surface that can be accommodated by the natural plasticity already encoded in host protein dynamics. This reduces the energetic barrier to initial contact formation and allows the virus to engage transiently with multiple host factors in rapid succession — consistent with the kinetic requirements of productive endocytic entry.

### 5.3 Electrostatically Dominated Interfaces and Distributed Energetic Architecture

MM/GBSA analysis revealed that envelope–host interfaces are predominantly stabilized by electrostatic interactions, with gas-phase EEL contributions ranging from approximately −180 to −750 kcal mol^−1^ across complexes, substantially offset by polar solvation penalties. Three energetic regimes were identified: electrostatically saturated interfaces (ENV–GPC1, Δ*G*_total_ ≈ −90 kcal mol^−1^), electrostatically anchored interfaces (ENV–BiP, ≈ −70.5 kcal mol^−1^), and hydrophobically dominated interfaces (ENV–Clusterin). These values should be interpreted as semi-quantitative rankings given known MM/GBSA limitations for highly charged interfaces. A detailed quantitative discussion of these limitations — including the published *R*^2^ benchmarks for MM/GBSA against experimental affinities, the overestimation at electrostatically saturated interfaces (such as ENV–GPC1), and recommendations for block-averaging and FEP follow-up at the highest-priority hotspot pairs — is provided in Section 6.5 of the Supplementary Material.

Per-residue decomposition consistently revealed a distributed energetic architecture spanning multiple spatially separated residue clusters on both partners, rather than single dominant hotspots. For interactions with highly conserved cellular scaffolds such as actin and BiP, this energetic redundancy may contribute to robustness of host factor engagement across the sixteen genotype variants examined. The distributed architecture also implies that the interactions are expected to be tolerant of individual point mutations in either partner — a property consistent with the functional conservation of actin-mediated and chaperone-mediated flavivirus processes across DENV serotypes.

### 5.4 Geometric Regimes Define Functional Interaction Classes

The four geometric regimes identified in this study represent distinct functional classes of host engagement that provide a unified structural interpretation for the heterogeneous set of host factors implicated in dengue infection:

- *Anchoring interfaces* provide multivalent stabilization surfaces that increase viral residence time at the cell surface without irreversible capture.
- *Sliding interfaces* facilitate electrostatic capture and surface exploration, enabling dimensionality reduction of the receptor search.
- *Focal regulatory interfaces* correspond to checkpoints whose engagement connects viral docking to immune signaling cascades.
- *Cytoskeletal transport interfaces* govern intracellular trafficking and viral progression after endocytosis.

### 5.5 Network Context Determines Viral Compatibility

Ternary docking analyses demonstrate that the surrounding host network can strongly modulate viral docking stability, with conditional ΔΔ*G* values ranging from approximately −3.4 to +6.9 kcal mol^−1^ depending on the neighboring protein and assembly sequence. Classical actin regulators (ACTN2, PFN1) produced destabilizing effects consistent with competitive occupation of shared actin determinants, whereas dynactin-associated components (ACTR1A, ACTR10) showed patterns consistent with cooperative stabilization. These observations suggest that the cellular cytoskeletal environment at the time of viral encounter may influence the thermodynamic favorability of actin engagement — a prediction directly testable through targeted knock-down experiments and infection assays in cells with perturbed cytoskeletal composition.

The finding that viral engagement with trafficking modules (Actin, Rab5) is temporally gated by the surrounding host protein network carries significant implications for understanding viral tropism. Viral entry efficiency may depend not only on host protein expression levels but also on which molecular network states are structurally populated within specific cell types and activation states. The complete per-protein ternary interaction tables — including all ΔΔ*G* values, partner identity, assembly order, and cooperative/competitive classification — are provided in the Supplementary Material (Section 4, one ternary subsection per host protein). A critical methodological note is that ternary assemblies in the present pipeline were generated by sequential rigid-body docking without post-assembly MD relaxation; the systematic implications of this approximation, including potential overestimation of competitive ΔΔ*G* from unrelaxed steric clashes, and the three priority ternary assemblies recommended for validation by short MD (NRP1–PLXNA4, SUMO1– IPO13, GPC1–HBEGF), are addressed in Section 6.6 of the Supplementary Material.

### 5.6 Genotype-Informed Framework: Conservation Across the Circulating DENV Panel

The basic residue clusters mediating electrostatic interactions — particularly in the DIII electropositive patch — were structurally present across all sixteen envelope variants examined, as was the structural position corresponding to ARG410 in the BiP complex and the candidate CendR-like loop residue cluster in the NRP1 complex. This conservation across the genotype panel suggests that the principal interaction architectures characterized here are not idiosyncratic properties of a single representative structure but are broadly encoded across the circulating sequence space of DENV1–4.

Serotype-resolved vectorial surface fitting confirmed that curvature and orientation of fitted quadratic surface descriptors are broadly concordant across genotypic clusters, while revealing modest but detectable inter-genotype differences in hydrogen-bond vector directionality, particularly at the DIII surface. These vectorial differences represent testable structural hypotheses for serotype-specific differences in attachment efficiency.

An important caveat accompanies the genotype-resolved designation: the molecular dynamics trajectories and MM/GBSA analyses reported here were conducted on single representative envelope structures per host protein complex, not across all sixteen genotype variants. Full quantitative comparison of Δ*G* values and hotspot contributions across the complete DENV1–4 panel — requiring approximately 208 additional independent 200 ns simulations — remains a priority deferred objective. This limitation, its implications for interpreting the term “genotype-informed” in a quantitative sense, and the recommended experimental approaches for closing this gap (SPR/BLI with purified genotype ectodomains) are discussed in Section 5.10 of the Supplementary Material.

### 5.7 Pharmacological Implications

The structural insights generated by this study identify several priority pharmacological opportunities, with the electrostatic frustration architecture of envelope–host interfaces providing a particularly actionable framework for drug design.

The frustration clusters identified at *LYS435/LYS516/LYS447* in BiP and at *ARG107/LYS236* in Betaglycan constitute priority targets for a *molecular glue* strategy. In this approach, small molecules would not compete with viral binding but would instead stabilize the envelope–host complex in a non-productive geometry — locking the interaction in a conformation that prevents the subsequent conformational progression required for membrane fusion. The electrostatic frustration topology at these clusters is particularly well-suited for molecular glue design: the adjacent charged residues generate competing favorable and unfavorable energetic contributions that produce a dynamic equilibrium between binding-permissive and binding-restrictive geometries. A molecular glue inserting at this frustration node could shift this equilibrium toward a non-productive dead-end complex, effectively trapping the viral particle at the cell surface while blocking endosomal progression. This strategy has precedent in the antiviral literature — lenacapavir’s stabilization of a non-productive HIV capsid state represents a well-characterized example — and the structural specificity of the dengue frustration sites (LYS435 in the BiP nucleotide-binding domain, ARG107 in the Betaglycan core-binding region) provides chemically tractable anchor points distinct from any current dengue therapeutic target.

The *NRP1 b1-pocket*, engaged by the dengue envelope through a CendR-like molecular mimicry mechanism, represents a second high-priority pharmacological target. This pocket is well-characterized structurally from peptide–NRP1 co-crystals, and small molecules or stapled peptides designed to occupy the b1 binding groove would directly compete with the CendR-like loop of the viral envelope, disrupting the focal regulatory interaction without requiring targeting of the viral protein itself — an approach that is inherently more resistant to viral escape through point mutation.

The *distributed energetic architecture* of the ENV–actin interface adds a third pharmacological opportunity. Because the interface lacks a single dominant hotspot, blocking it through competitive inhibition requires either large surface-mimetic compounds or allosteric approaches. However, the cooperative destabilization produced by dynactin-regulatory partners (ACTR1A, ACTR10) in ternary docking analyses identifies a network-level vulnerability: compounds that stabilize these cooperative partners at the actin surface could indirectly outcompete viral engagement through thermodynamic displacement rather than direct competition.

Collectively, the frustration-guided molecular glue targets (BiP, Betaglycan), the b1-pocket mimicry disruptors (NRP1), and the cytoskeletal network cooperativity targets (Actin/dynactin axis) suggest a preliminary three-tier druggability landscape corresponding to the first three stages of the hierarchical entry model (electrostatic capture → anchoring stabilization → regulatory checkpoint). Each tier offers mechanistically distinct intervention opportunities with complementary resistance profiles.

## 6 Conclusions

This study presents a multiscale computational characterization of the structural interaction landscape between sixteen dengue virus envelope genotype variants and thirteen host proteins implicated in viral attachment, endocytic trafficking, immune signaling, and vascular regulation. By integrating protein–protein docking, molecular dynamics simulations, MM/GBSA energetic analysis, and vectorial characterization of hydrogen bond networks — including circular directional statistics (*R, H*_norm_) and genotype-resolved quadratic surface fitting — the framework reveals consistent structural patterns that provide mechanistic context for understanding dengue virus–host protein engagement.

Four central conclusions emerge. *First*, dengue host engagement is organized as a four-stage hierarchical architecture proceeding from surface capture to intracellular delivery:

1. *Electrostatic Capture* — GPC1 was modeled as a representative heparan sulfate proteoglycan platform that may support initial dimensionality reduction through surface diffusion along sulfated glycosaminoglycan chains, exploiting the DIII electropositive patch of the envelope;
2. *Anchoring Stabilization* — BiP/GRP78 and Betaglycan extend viral residence time at the cell surface through high-avidity electrostatic interfaces with discrete anchor residues (ARG410 in BiP; ARG107/LYS236 in Betaglycan);
3. *Regulatory Checkpoints* — NRP1 (via a CendR-like pocket-engagement hypothesis) and the SUMO/UBC9 axis may connect viral docking to immune signaling cascades, representing candidate regulatory transitions between surface capture and productive endocytosis;
4. *Cytoskeletal Trafficking* — Actin and Rab5 direct internalized particles through productive endosomal pathways via context-dependent interfaces regulated by the surrounding network.

*Second*, viral docking consistently exploits molecular geometries already accessible within host interaction networks — a mechanism termed *interface capture* — in which the dengue envelope acts as a structural opportunist that selects pre-existing host protein conformations without requiring refolding. *Third*, the viral envelope functions as a conformationally rigid scaffold throughout this process, delegating the thermodynamic cost of interface formation to host proteins through asymmetric conformational adaptation. *Fourth*, host network context is a primary determinant of viral engagement efficacy: ternary interactions can either stabilize (DNAJB11, FICD for BiP; PLXNA1, SEMA4 for NRP1) or disrupt (the entire Rab5 regulatory panel; HYOU1/DERL1 for BiP) envelope docking depending on assembly order and partner identity, indicating that cellular microenvironment — not simply receptor expression level — governs site-specific infection susceptibility.

Priority experimental directions arising from this study include: (1) site-directed mutagenesis and SPR-based affinity measurement at the ARG410 hotspot in the ENV–BiP interface; (2) competition assays and cryo-EM structural determination for the ENV–NRP1 complex to test the CendR-like mimicry model; (3) CRISPR-based knockdown of ACTR1A and ACTR10 combined with dengue infectivity assays to assess the functional relevance of the predicted dynactin-cooperative interface; (4) live-cell single-virion tracking experiments in GPC1 knockout versus overexpressing cell lines to test the surf-riding dimensionality-reduction model; (5) Ubc9 catalytic activity assays in the presence of recombinant dengue envelope ectodomain using STAT1/STAT2 SUMO-acceptor peptide substrates; (6) cross-flavivirus NRP1 engagement profiling for Zika, West Nile, and Yellow fever envelope proteins to test the convergent geometric mimicry hypothesis; and (7) experimental molecular-glue screening campaigns prioritizing the LYS435/LYS516/LYS447 frustration cluster in BiP and the ARG107/LYS236 cluster in Betaglycan, using the non-productive complex stabilization strategy articulated in the Pharmacological Implications section.

Although developed here for dengue virus, the concept of vectorial host interaction fields — combining directional hydrogen bond formalism, circular coherence metrics, and quadratic surface descriptors — may apply broadly to viruses that exploit heterogeneous host factor repertoires. Understanding these interaction landscapes may therefore provide new opportunities for antiviral strategies targeting network-dependent stages of infection.

An extended Outlook section — elaborating the full set of eight priority experimental directions, discussing the broader applicability of vectorial interaction fields to other flaviviruses, and framing the pharmacological implications of electrostatic frustration-guided molecular glue targets within the wider antiviral landscape — is provided in Section 5.10 of the Supplementary Material. That section also contains the important qualification that full quantitative genotype-resolved affinity differentiation across the DENV1–4 panel remains a recommended next step, and outlines the specific experimental platforms (SPR, BLI, ITC, FEP) by which the computational predictions generated here can be most directly confronted with experimental evidence.

## 7 Limitations

Our computational framework provides valuable structural and energetic insights, yet several methodological constraints warrant consideration when interpreting results and assessing biological implications. Complete discussion with quantitative benchmarks, published error estimates, and experimental recommendations is provided in the Supplementary Material (Section 6, lines 4707–4744). Key constraints include:

### Single-conformation docking (Section 6.1, line 4711)

Rigid-body docking simulations used single representative structures, omitting conformational selection and induced-fit mechanisms. Ensemble docking with conformational diversity from MD pre-sampling would improve priority interface refinement.

### Virion context and membrane environment (Section 6.2, line 4715)

The dengue envelope protein exists as an icosahedral lattice on the virion surface, anchored to lipid membranes. Our simulations used soluble ectodomains, excluding orientational constraints from membrane anchoring, steric effects from dimer contacts, and contributions of stem residues to interface energetics.

### Unmodeled N-linked glycans (Section 6.3, line 4719)

Conserved glycosylation at Asn67 and Asn153 was omitted from structures and simulations, potentially affecting local electrostatics and contact geometry— particularly important for Glypican-1 and Neuropilin-1 interfaces where glycan-mediated steric masking could substantially alter predicted binding modes.

### MD timescale limitations (Section 6.4, line 4723)

The 200 ns trajectories capture local conformational adjustments but not dissociation kinetics on biologically relevant timescales (microseconds to milliseconds). The ENV–TGFBR1 complex notably fails to achieve PCA convergence, indicating MM/GBSA estimates represent a non-equilibrated ensemble rather than a true bound state.

### MM/GBSA semi-quantitative nature (Section 6.5, line 4727)

Energetics should be interpreted as rankings rather than absolute free energies. The Supplementary Material discusses published benchmarks (*R*^2^ = 0.3–0.6 against experimental values), systematic overestimation at electrostatically saturated interfaces, and recommends block-averaging and FEP refinement for priority complexes.

### Ternary framework assumptions (Section 6.6, line 4733)

The AB/BC/ABC framework assumes ternary energetics can be decomposed into pairwise contributions, potentially underestimating higher-order cooperativity. Post-assembly MD relaxation of ternary complexes would validate three priority assemblies: NRP1–PLXNA4, SUMO1–IPO13, and GPC1–HBEGF.

### Absence of macromolecular crowding (Section 6.7, line 4739)

All simulations used dilute solvent, omitting excluded-volume and viscosity effects of the cytoplasm, which generally favor compact states and may alter association kinetics relative to dilute predictions.

Additionally, molecular dynamics and MM/GBSA analyses used single representative envelope structures per complex; comprehensive cross-genotype analysis remains an important future objective, detailed in the Supplementary Material (Section 5.10, line 4666).

## 8 Data Availability

All structural models, docking result files, molecular dynamics trajectories, VectorPROT analysis outputs, and complete per-residue MM/GBSA decomposition data — including the 27 individual host-protein figures (PCA, MM/GBSA, and vectorial visualization panels) referenced throughout this manuscript — are provided in the companion Supplementary Material and its Supplementary Information. Complete data availability statement and information on accessing supplementary datasets is provided in the Main Manuscript (Section 10, line 4788). R and Python analysis scripts, input structures, docking outputs, GROMACS input files, MM/GBSA parameters, complete energetic tables, hotspot tables, software versions, and model-selection criteria should be deposited in a public repository such as GitHub, Zenodo, OSF, or an institutional repository with a persistent DOI before submission. Until deposition is complete, these files should be supplied as supplementary material or shared directly during review to preserve reproducibility. For computational resource credit and HPC infrastructure information, refer to the Supplementary Material (Section 11, line 4813).

## 9 Author Contributions

Daniel Ferreira de Lima Neto and Caio Cesar de Melo Freire contributed equally to this work and share corresponding authorship. Daniel Ferreira de Lima Neto conceived and designed the study, developed the complete computational framework, implemented the analytical methodologies, performed molecular docking experiments, molecular dynamics simulations, and energetic analyses, curated and interpreted the datasets, prepared the figures, and wrote the original manuscript draft, also coordinating the overall research workflow. Caio Cesar de Melo Freire co-conceived and co-designed the study, participated in the development and implementation of the computational framework, contributed to all stages of the molecular dynamics simulations and energetic analyses, performed data curation and interpretation, co-prepared the figures, and substantially contributed to writing and revising the manuscript. Juan Philippe Teixeira assisted with data organization, computational analysis support, and visualization. Marcus Alexandre Finzi Corat contributed with institutional support and critical revision of the manuscript. Miklos Maximiliano Bajay contributed to discussions on bioinformatic analyses and interpretation of computational results. Gustavo Henrique Goulart Trossini contributed to scientific discussion and provided expertise in molecular modeling and structural analysis. Daniel Santos Mansur contributed to conceptual discussions on virus–host interaction mechanisms and critical revision of the manuscript. Elcio de Souza Leal contributed to the interpretation of virological data and critical revision of the manuscript. Luiz Mario Ramos Janini contributed to scientific discussion and provided expertise in dengue virology and manuscript revision. All authors reviewed and approved the final version of the manuscript.

Correspondence: Daniel Ferreira de Lima Neto and Caio Cesar de Melo Freire.

## 12 Acknowledgements

The authors acknowledge the computational infrastructure and technical support provided by the high-performance computing (HPC) facilities coordinated by Dr. Edroaldo Lummertz da Rocha (Federal University of Santa Catarina), whose support was essential for enabling the large-scale molecular simulations and computational analyses performed in this study. We would like to extend our sincere thanks to Prof. Dr. Carlos Francisco Sampaio Bonafé (*in memoriam*). His profound teachings on protein thermodynamics were fundamental to shaping our understanding of the concepts explored in this work. We hope this paper honors his memory.

## 14 Code Availability and Reproducibility

Computational workflows, analysis pipelines, and scripts for molecular docking, molecular dynamics simulations, structural analysis, and trajectory processing should be deposited with the public reproducibility package. All computational procedures employed established molecular modeling frameworks (GROMACS, AutoDock, and others) with standardized protocols detailed in the Materials and Methods section, enabling independent replication and verification of results.

## 15 Data Availability

All data generated during this study are included in the published article and supplementary information files. Additional datasets from molecular docking and molecular dynamics simulations, complete vectorial analyses, and per-residue MM/GBSA decompositions should be deposited in the same public reproducibility package before submission.

