## Supplementary Material for "Genotype-Informed Interaction Landscape of the Dengue Envelope Glycoprotein with Host Cellular Proteins: Structural Dynamics, Thermodynamic Cooperativity, and Binding Specificity"

May 31, 2026

Daniel Ferreira de Lima Neto[1,*], Juan Philippe Teixeira[4], Marcus Alexandre Finzi Corat[5], Miklos Maximiliano Bajay[4], Gustavo Henrique Goulart Trossini[2,3], Daniel Santos Mansur[1], Elcio de Souza Leal[6], Luiz Mario Ramos Janini[7], Caio Cesar de Melo Freire[8,*]

*[1]Laboratory of Immunobiology, Department of Microbiology, Immunology and Parasitology, Center of Biological Sciences, Federal University of Santa Catarina (UFSC), Av. Prof. Henrique da Silva Fontes 2754, Florianópolis, SC 88040-900, Brazil*

*[2]Laboratory for the Integration of Experimental and Computational Techniques (LITEC), Department of Pharmacy, School of Pharmaceutical Sciences, University of São Paulo (USP), Av. Prof. Lineu Prestes 580, São Paulo, SP 05508-000, Brazil*

*[3]Institute of Advanced Studies of the University of São Paulo, University of São Paulo, Praça do Relógio 109, Ground floor, Cidade Universitária, São Paulo, SP 05508-050, Brazil*

*[4]Graduate Program in Animal Science and Department of Fishery Engineering and Biological Sciences, Santa Catarina State University (UDESC), Rua Beloni Trombeta Zanini 680, Chapecó, SC 89815-630, Brazil*

*[5]Multidisciplinary Center for Biological Research on Laboratory Animal Science (CEMIB), University of Campinas (UNICAMP), Rua 5 de Junho 250, Campinas, SP 13083-970, Brazil*

*[6]Laboratory of Virology, Institute of Biological Sciences, Federal University of Pará (UFPA), Belém, PA, Brazil*

*[7]Laboratory of Retrovirology, Department of Microbiology, Immunology and Parasitology, Escola Paulista de Medicina, Federal University of São Paulo (UNIFESP), São Paulo, SP, Brazil*

*[8]Department of Genetics and Evolution, Center of Biological and Health Sciences, Federal University of São Carlos (UFSCar), Rod. Washington Luís, km 235, São Carlos, SP 13565-905, Brazil*

*[*]These authors contributed equally to this work. Correspondence:* `danielvirogmail.com` *(D.F.L.N.);*

caio.freireufscar.br (C.C.M.F.)


#### Abstract

Dengue virus (DENV) is a mosquito-borne flavivirus responsible for approximately 390 million infections annually, representing one of the most significant global public health challenges among arboviruses. The dengue envelope (E) glycoprotein is the principal mediator of viral attachment, receptor recognition, and membrane fusion during host cell entry. This study characterizes a genotype-informed structural interaction landscape between sixteen dengue envelope genotype variants and a curated panel of host proteins involved in viral attachment, endocytic trafficking, immune signaling, and vascular regulation. Three-dimensional structural models were obtained through a hierarchical pipeline integrating experimentally determined structures, AlphaFold predictions, I-TASSER template-based modeling, and MODELLER refinement, followed by molecular dynamics relaxation. Protein–protein docking simulations were performed using HDOCK across the sixteen-variant envelope panel, whereas molecular dynamics and MM/GBSA analyses were conducted on representative complexes selected from this genotype-informed screening. Interaction interfaces were quantitatively characterized through a vector-based formalism encoding hydrogen bond directionality, occupancy, and angular entropy. To approximate cellular interaction environments, triple docking simulations incorporating first-neighbor host proteins were implemented under both pre-assembly and post-assembly scenarios, and binding cooperativity was evaluated through $\Delta\Delta G$ thermodynamic analysis. This framework provides a multiscale representation of viral–host interaction architecture and identifies structural determinants potentially associated with viral entry, host immune modulation, and vascular pathogenesis. These results should be interpreted as computational structural hypotheses that require biochemical, biophysical, and cellular validation.

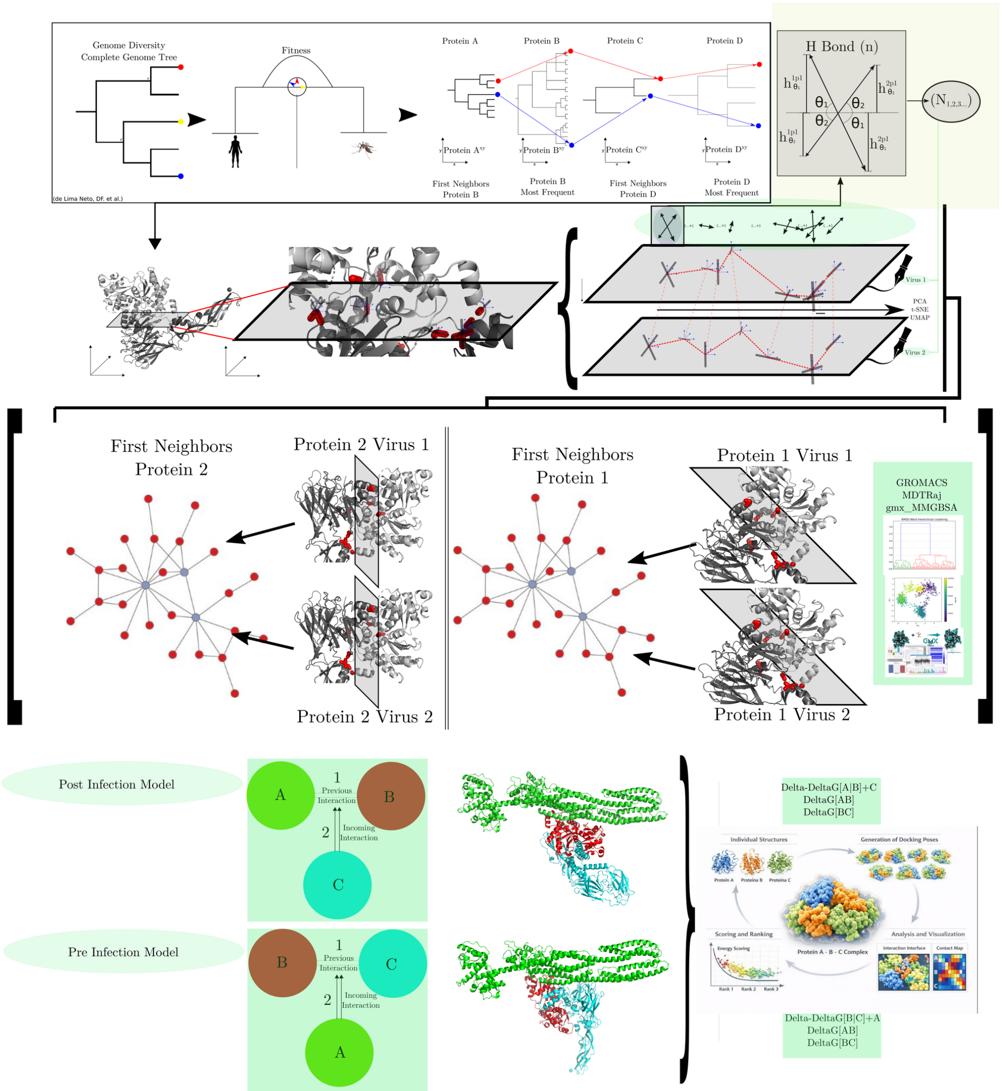


**Figure 1:** Graphical abstract

### 1 Introduction

Dengue virus (DENV) is a mosquito-borne flavivirus responsible for approximately 390 million infections annually, with nearly 100 million symptomatic cases worldwide, representing one of the most significant global public health challenges among arboviruses [1]. The virus belongs to the genus *Flavivirus* within the family *Flaviviridae*, which also includes medically important pathogens such as Zika virus, West Nile virus, and Yellow fever virus. A defining feature of flaviviruses is the presence of a lipid envelope containing the envelope glycoprotein (E), a multifunctional protein that mediates viral attachment to host cells, receptor recognition, and membrane fusion during entry [2,3].

The DENV envelope protein is arranged as antiparallel dimers on the virion surface and undergoes major conformational rearrangements during infection. Following receptor engagement and endocytosis, acidification within early endosomes induces a structural transition in which envelope dimers reorganize into fusogenic trimers, enabling the viral membrane to fuse with the host endosomal membrane and release the viral genome into the cytoplasm [4,5]. Because of its central role in viral entry and tropism, the E protein has been extensively studied as the primary mediator of host–virus interactions and represents a critical target for structural, immunological, and antiviral investigations [6].

Dengue virus entry into host cells is a complex and multifactorial process involving a wide range of host surface molecules and intracellular trafficking pathways. Rather than relying on a single receptor, flaviviruses typically exploit a combination of attachment factors, co-receptors, and signaling molecules that collectively facilitate viral adsorption, internalization, and intracellular transport [7,8]. Several classes of host proteins have been implicated in these processes, including glycosaminoglycan-binding proteins, lectins, heat-shock proteins, cytoskeletal components, and regulators of endosomal trafficking.

#### 1.1 Heparan Sulfate Proteoglycans as Attachment Factors

One of the best-characterized attachment mechanisms for dengue virus involves interactions with heparan sulfate proteoglycans, which function as initial attachment factors that increase viral concentration at the cell surface prior to engagement with more specific receptors [9–11]. Glypican-1 was modeled here as a representative heparan sulfate proteoglycan platform rather than as an independently established dengue receptor. Proteoglycans such as Glypican-1, which contain sulfated glycosaminoglycan chains anchored to the plasma membrane via glycosylphosphatidylinositol (GPI), represent plausible candidates for envelope interaction studies due to their ability to mediate electrostatic interactions with viral surface proteins. These molecules have been shown to facilitate the attachment of multiple viruses, including flaviviruses and alphaviruses, highlighting their importance in viral tropism and host specificity.

#### 1.2 Stress Chaperones and Endosomal Regulators

Another host factor strongly associated with dengue virus entry is the endoplasmic reticulum chaperone BiP/GRP78 (glucose-regulated protein 78). Although primarily located in the endoplasmic reticulum, GRP78 can translocate to the cell surface under stress conditions and act as a viral attachment factor. Experimental studies have demonstrated that GRP78 interacts with the dengue envelope protein and promotes viral entry in mammalian cells [12,13]. Similar roles for GRP78 have also been observed in infections caused by other RNA viruses, suggesting that stress-induced chaperones may act as conserved viral docking platforms.

Cytoskeletal components also play a fundamental role in viral entry and intracellular trafficking. The actin cytoskeleton is essential for membrane dynamics, endocytosis, and vesicular transport, processes widely exploited by viruses during infection. Several studies have shown that actin remodeling is required for efficient dengue virus internalization and that disruption of actin dynamics significantly reduces dengue entry and productive infection [14–16]. Once internalized, dengue virus particles are transported through the endosomal pathway, where early endosomes marked by the small GTPase Rab5 play a crucial role. Rab5 regulates vesicle docking, membrane fusion, and early endosome maturation, and its activity is required for the trafficking of flaviviruses to acidified compartments where membrane fusion occurs [17,18].

#### 1.3 Vascular Signaling and Immune Modulators

In addition to proteins directly involved in viral entry, several membrane receptors associated with signaling pathways relevant to dengue pathogenesis were included in this study. Neuropilin-1 (NRP1) is a multifunctional receptor involved in angiogenesis and vascular permeability through its interactions with vascular endothelial growth factor (VEGF). Neuropilin-1 has been identified as an entry factor for SARS-CoV-2 and participates in endothelial signaling networks that regulate vascular integrity [19]. Because vascular leakage and endothelial dysfunction are hallmark features of severe dengue disease, NRP1 was included here as a biologically plausible vascular-signaling receptor; however, ENV–NRP1 engagement is treated as a computational hypothesis for dengue rather than as an established DENV entry mechanism.

Similarly, the TGF-$\beta$ receptor system, including Betaglycan (TGFBR3) and TGFBR1, regulates immune modulation, extracellular matrix remodeling, and inflammatory responses. Elevated TGF-$\beta$ signaling has been observed in dengue patients and may contribute to vascular permeability and immune dysregulation during infection [20,21]. The VEGF signaling pathway itself is strongly implicated in dengue pathophysiology;

elevated levels of VEGF and its receptors have been detected in patients with severe dengue and are associated with plasma leakage and vascular permeability [22].

Tight junction proteins such as Claudin-1 were also included because of their role in maintaining epithelial and endothelial barrier integrity. Claudin-1 is a known entry factor for hepatitis C virus and has also been experimentally implicated in dengue entry through interaction between Claudin-1 and viral prM/M [23,24]. In the context of host immune responses, the activating natural killer cell receptor NKp44 (NCR2) was included because NKp44 has been shown to recognize flaviviral envelope glycoproteins from dengue and West Nile viruses [25]. The NLRP3 inflammasome, a multiprotein complex responsible for sensing cellular stress and viral infection, has been shown to be activated during dengue virus infection, leading to the production of pro-inflammatory cytokines such as IL-1$\beta$ [26].

Additional host factors included in this analysis represent regulatory components involved in protein stability and immune modulation. The SUMOylation system regulates transcription, protein localization, and antiviral responses, and several RNA viruses manipulate SUMOylation pathways to enhance replication or suppress host immune responses [27]. Finally, Clusterin (Apolipoprotein J) was included because of its role as an extracellular chaperone involved in lipid transport, complement regulation, and cellular stress responses [28].

Taken together, the proteins selected in this study represent multiple layers of host biology relevant to dengue infection, including viral attachment factors, membrane receptors, cytoskeletal components, endocytic regulators, immune sensors, and signaling molecules associated with vascular regulation and inflammation. Structural modeling and protein–protein docking approaches enable the characterization of interaction interfaces at atomic resolution, allowing identification of hydrogen bond networks, electrostatic contacts, and hydrophobic interaction surfaces that stabilize protein complexes [29]. By integrating structural interaction modeling with host protein selection grounded in biological evidence, this study aims to characterize potential interaction networks between the dengue envelope glycoprotein and host factors involved in membrane dynamics, immune regulation, and intracellular trafficking.

### 2 Materials and Methods

#### 2.1 Sequence Retrieval and Dataset Construction

The amino acid sequences corresponding to the host factors investigated in this study — Actin, Betaglycan (TGFBR3), BiP/GRP78, Claudin-1, Clusterin, Glypican-1, Neuropilin-1, NKp44, NLRP3, Rab5, SUMO, TGFBR1, and VEGF — were retrieved from the UniProt Knowledgebase (UniProtKB). UniProt integrates protein sequence information with functional annotations and cross-references to multiple biological databases, including structural repositories such as the Protein Data Bank (PDB) and the AlphaFold Protein Structure Database [30]. For each protein included in the study, the reviewed Swiss-Prot entry corresponding to *Homo sapiens* was selected to ensure the highest level of annotation quality and experimental validation. In cases where multiple isoforms were available, the canonical isoform annotated in UniProt was preferentially selected unless previous experimental studies suggested functional relevance of a specific isoform in viral infection pathways. The selected sequences were downloaded in FASTA format and organized into a standardized dataset serving as input for structural modeling and protein–protein interaction analyses.

#### 2.2 Structural Model Acquisition

Three-dimensional structures for the host proteins were obtained through a hierarchical strategy integrating experimentally determined structures and computational modeling approaches. Whenever experimentally resolved structures were available, atomic coordinates were retrieved from the Protein Data Bank, the primary international repository for experimentally determined macromolecular structures obtained through X-ray crystallography, nuclear magnetic resonance spectroscopy, and cryo-electron microscopy [31]. When multiple structural entries were available for a given protein, candidate structures were evaluated according to crystallographic resolution, structural completeness, domain coverage, and the absence of extensive unresolved regions.

#### 2.3 Structure Prediction Using AlphaFold

For proteins lacking experimentally determined structures or presenting incomplete domain coverage, structural models were generated using AlphaFold [32]. AlphaFold is a deep learning–based structure prediction system capable of inferring highly accurate three-dimensional protein structures directly from amino acid sequences, integrating evolutionary information derived from multiple sequence alignments with neural network architectures trained on experimentally determined protein structures. Model reliability was evaluated using the predicted Local Distance Difference Test (pLDDT) confidence score. Models exhibiting globally high confidence scores and coherent domain organization were retained for further structural validation.

#### 2.4 Template-Based Structural Modeling with I-TASSER

In cases where AlphaFold models were unavailable or insufficient, additional structural predictions were generated using the I-TASSER 5.2 standalone software [33]. I-TASSER performs hierarchical protein structure prediction through template threading, fragment assembly simulations, and iterative atomic-level refinement. The resulting structural models were evaluated using the C-score metric, which reflects the significance of template alignments and the convergence of structural assembly simulations. To accelerate modeling of multiple proteins, the I-TASSER standalone implementation was executed using GNU Parallel, enabling simultaneous structural predictions across multiple computational cores.

#### 2.5 Structural Validation and Quality Assessment

All predicted and experimentally obtained structures were subjected to structural validation using the PDBsum Generate pipeline, which provides automated structural summaries including secondary structure topology diagrams, hydrogen bond interaction maps, domain organization information, and stereochemical validation reports derived from the PROCHECK algorithm [34]. Stereochemical quality was further evaluated using Ramachandran plot analysis, which describes the distribution of backbone dihedral angles $\phi$ and $\psi$ for amino acid residues and provides a widely accepted metric for assessing structural quality [35]. High-quality protein structures typically present more than 90% of residues within favored conformational regions and minimal residues in disallowed regions [36]. Structures exhibiting unfavorable stereochemical geometry were subjected to additional structural refinement.

#### 2.6 Structural Refinement Using MODELLER

Structural models requiring geometric optimization were refined using MODELLER, a comparative protein modeling program that improves protein structures through spatial restraints derived from homologous templates [37]. During the refinement procedure, predicted structures were aligned with homologous structural templates when available, and multiple refined structural models were generated through iterative optimization of spatial restraints. The resulting models were evaluated using the DOPE (Discrete Optimized Protein Energy) scoring function. Models presenting the lowest DOPE energy values and the most favorable stereochemical properties were selected for subsequent analyses.

#### 2.7 Molecular Dynamics Structural Relaxation

To further improve structural stability and eliminate potential steric clashes, selected protein models were subjected to molecular dynamics simulations using GROMACS [38]. Protein structures were converted into molecular simulation systems by generating topology files using an appropriate biomolecular force field. The systems were solvated in explicit water molecules within a simulation box and neutralized by the addition of counterions. Energy minimization was performed using the steepest descent algorithm. Following minimization, equilibration simulations were conducted in two stages: first under the NVT ensemble for temperature stabilization, and subsequently under the NPT ensemble for pressure and density equilibration. Short production simulations were performed to allow relaxation of backbone and side-chain conformations, and structural stability was evaluated through trajectory analysis to ensure convergence of the simulated systems.

#### 2.8 Construction of the Dengue Envelope Genotype Dataset

Sixteen structural variants representing distinct dengue virus envelope (ENV) genotypes were selected to represent the genetic diversity of circulating dengue virus populations. Structural models for each genotype were generated using the same structural modeling and refinement pipeline applied to host proteins, producing a comprehensive dataset enabling systematic investigation of genotype-specific differences in viral interaction interfaces.

#### 2.9 Protein–Protein Docking Simulations

Protein–protein docking simulations were performed using the HDOCK docking platform, a hybrid docking algorithm that integrates template-based modeling and *ab initio* docking strategies to predict protein–protein complexes [39]. The HDOCK scoring function accounts for geometric complementarity, electrostatic interactions, and desolvation energy. Each ENV genotype was docked against each host protein included in the dataset, resulting in a comprehensive docking matrix representing all possible viral–host interaction combinations. For each docking simulation, multiple candidate complexes were generated and ranked according to the HDOCK scoring function, and the highest-ranked conformations were selected for subsequent interaction analysis.

All docking calculations used single representative structures for each protein partner, without incorporation of conformational diversity through ensemble docking approaches. This rigid-body approximation may not fully capture interactions governed by conformational selection or induced-fit mechanisms. Ensemble docking approaches that incorporate conformational diversity sampled by molecular dynamics could refine the interaction landscape identified here and are recommended for follow-up studies of the highest-priority interfaces [40].

#### 2.10 Vectorial Representation of Hydrogen Bond Interactions

Protein–protein interfaces are stabilized by networks of non-covalent interactions including hydrogen bonds, electrostatic interactions, van der Waals forces, and hydrophobic contacts [29,41]. To quantitatively characterize the spatial organization of these interactions across viral–host interfaces, hydrogen bond interactions were represented using a vector-based formalism encoding both the direction and magnitude of intermolecular contacts.

##### 2.10.1 Mathematical Formalization of Interaction Vectors

For each interacting residue pair $(i)$ and $(j)$, a hydrogen bond interaction was represented as a three-dimensional vector connecting the donor atom $(D)$ and the acceptor atom $(A)$, defined as:

$$\vec{v}_{ij}(t) = \mathbf{r}_A(t) - \mathbf{r}_D(t) \tag{1}$$

where $\mathbf{r}_A(t)$ represents the spatial coordinate of the acceptor atom and $\mathbf{r}_D(t)$ corresponds to the coordinate of the donor atom at structural configuration $t$. To isolate the directional component of the interaction independently of bond length, normalized interaction vectors were calculated as:

$$\hat{v}_{ij}(t) = \frac{\vec{v}_{ij}(t)}{|\vec{v}_{ij}(t)|} \tag{2}$$

which preserves only the orientation of the interaction in three-dimensional space.

##### 2.10.2 Temporal Occupancy of Interactions

The persistence of each hydrogen bond interaction across structural configurations was quantified through an occupancy parameter defined as:

$$p_{ij} = \langle w_{ij}(t) \rangle \tag{3}$$

where $w_{ij}(t)$ represents a binary variable indicating the presence or absence of the interaction at time $t$. The occupancy parameter therefore represents the fraction of structural frames in which the hydrogen bond is observed.

###### 2.10.3 Angular Entropy of Interaction Vectors

Directional variability of hydrogen bond vectors was quantified through angular entropy analysis. Angular entropy describes the dispersion of vector orientations across structural configurations and is defined as:

$$S_{\mathrm{ang},ij} = -\sum_k P_k \ln P_k \tag{4}$$

where $P_k$ represents the probability that the interaction vector occupies orientation bin $k$ within the angular distribution. Low angular entropy values correspond to geometrically stable interactions with consistent directional orientation, whereas high angular entropy values indicate flexible or transient interactions.

###### 2.10.4 Construction of Vector Interaction Networks

Using the parameters described above, each viral–host interface was represented as a vector interaction network in which residues correspond to nodes and hydrogen bond interactions correspond to edges. Each edge was characterized by the average orientation of the interaction vector ($\hat{v}_{ij}$), the occupancy ($p_{ij}$), and the angular entropy ($S_{\mathrm{ang},ij}$). This representation enables quantitative comparison of interaction architectures across viral genotypes and host proteins and allows identification of conserved structural motifs associated with receptor recognition.

##### 2.11 Identification of First-Neighbor Interaction Partners

After establishing the structural models for each host protein and generating the viral docking complexes, an additional layer of structural context was incorporated through the identification of first-neighbor proteins within the host cellular interaction network. In cellular systems, proteins interact through cooperative assemblies, transient complexes, and signaling scaffolds that can modulate binding affinity, steric accessibility, and conformational dynamics of receptor interfaces [42,43].

First neighbors were defined as proteins that directly interact with each host factor according to curated protein–protein interaction databases and experimentally validated interaction datasets. Interaction partners were identified through cross-referencing UniProt annotations with interaction resources such as STRING, BioGRID, and IntAct. Only interactions supported by experimental evidence or high-confidence interaction scores were considered. In network theory terms, these proteins correspond to nodes connected through edges representing experimentally validated physical interactions [44]. From the resulting interaction network, first neighbors were selected based on biological relevance, structural availability, and functional association with membrane signaling, cytoskeletal dynamics, or immune pathways. Structural models for the selected first neighbors were obtained using the same modeling pipeline previously described.

#### 2.12 Vectorial Analysis of Molecular Interactions at the Protein–Protein Interface (VectorPROT)

##### 2.12.1 Overview and Experimental Logic

To characterize the directional organization of intermolecular contacts at the envelope–host protein interface, a pipeline for the vectorial analysis of molecular interactions was developed and applied to structural models representing three experimental conditions: (i) the basal reference state of host protein–first neighbor contacts (`3D_neighbors`); (ii) the pre-infection configuration, in which the host–first neighbor complex is assembled prior to viral engagement (`PRE`); and (iii) the post-infection configuration, in which the viral envelope protein is present in the ternary complex (`POS`). Three interaction scenarios were defined to reflect the progression of viral engagement across the host protein network (Table 1).

**Table 1:** Interaction scenarios used in the vectorial analysis pipeline.

| Scenario | Chain Pair | Biological Description |
|---|---|---|
| AB | Virus (A) ↔ Host protein (B) | Direct viral–host protein contact. |
| BC | Host protein (B) ↔ First neighbor (C) | Host protein engaged with its first PPI-network neighbor. |
| ABC | AB and BC simultaneously | Two sub-experiments per PDB: *exp01* (A–B); *exp02* (B–C). Models cooperative recruitment of the first neighbor. |

Experimental conditions are encoded in the directory hierarchy and automatically recognized by the pipeline during batch processing. This design ensures full traceability of each interaction vector to its source structure, chain pair, and condition.

##### 2.12.2 Structural Extraction and Local Interface Reference Frame

Each PDB file was parsed using the `Bio.PDB.PDBParser` module from Biopython [45], and chain detection was performed automatically on model 0 of each assembly. Interface atoms were selected using a proximity criterion: an atom of chain A was defined as belonging to the interface if at least one atom of chain B was located within a cutoff radius of 6.0 Å (adjustable via `--interface-cutoff`). The geometric centroids of the interface atom sets for each chain were computed as:

$$\mathbf{c}_A = \langle \mathbf{r}_i \mid a_i \in \text{interface}(A) \rangle \,, \quad \mathbf{c}_B = \left\langle \mathbf{r}_j \mid a_j \in \text{interface}(B) \right\rangle \tag{7}$$

A local orthonormal reference frame was constructed such that the $\hat{z}$ axis is normal to the mean interface plane and oriented from chain A toward chain B:

$$\hat{z} = \frac{\mathbf{c}_B - \mathbf{c}_A}{\|\mathbf{c}_B - \mathbf{c}_A\|} \tag{8}$$

When centroids coincide (norm $< 10^{-6}$ Å), $\hat{z} = [0, 0, 1]$ is used as a fallback. The complete orthogonal basis $\{\hat{x}, \hat{y}, \hat{z}\}$ was constructed by Gram–Schmidt orthogonalization using $[1, 0, 0]$ as the initial preference vector (substituted by $[0, 1, 0]$ if too parallel to $\hat{z}$). The frame origin is defined as the midpoint of the two centroids:

$$\mathbf{o} = \tfrac{1}{2}\left(\mathbf{c}_A + \mathbf{c}_B\right) \tag{9}$$

All subsequent coordinates and vectors are expressed in this local frame, rendering analyses invariant to the absolute position and orientation of the structure in Cartesian space.

###### 2.12.3 Detection and Vectorization of Intermolecular Contacts

Two classes of intermolecular interactions were detected and parametrized:

**Hydrogen bonds** were identified when (a) the interatomic distance between atoms from the two chains was $\leq 3.5$ Å (`--hbond-cutoff`), and (b) both atoms were nitrogen (N) or oxygen (O), qualifying them as hydrogen bond donors or acceptors. This criterion was applied to both backbone and side-chain atoms.

**Hydrophobic contacts** were registered when (a) the interatomic distance was $\leq 5.0$ Å (`--hydrophobic-cutoff`), (b) both atoms were side-chain carbons (element C, excluding backbone atoms N, C$\alpha$, C, O, and OXT), and (c) both residues belonged to the canonical hydrophobic set: Ala, Val, Leu, Ile, Met, Phe, Tyr, Trp, Pro, and Cys.

For each detected interaction, the parameters listed in Table 2 were computed and stored. Each interaction was thus represented as a (midpoint, vector) pair in the local interface frame, encoding both the spatial position and the directional orientation of the contact relative to the interface plane.

**Table 2:** Vectorial parameters computed for each detected interaction.

| Field | Formula | Description |
|---|---|---|
| `dist_Ang` | $\|\mathbf{r}_B - \mathbf{r}_A\|$ | Interatomic distance (Å). |
| `mid_local` (x, y, z) | $T_{\text{local}}\big(\frac{1}{2}(\mathbf{r}_A + \mathbf{r}_B),\, \mathbf{o},\, \mathcal{B}\big)$ | Midpoint in local frame. |
| `vec_local` (x, y, z) | $T_{\text{local}}(\mathbf{r}_B - \mathbf{r}_A,\, \mathcal{B})$ | Interaction vector in local frame. |
| `angle_deg_to_plane` | $\arctan 2(\lvert v_z\rvert,\, \sqrt{v_x^2 + v_y^2})$ | Elevation angle from the interface plane (°). |

###### 2.12.4 Circular Statistics and Normal/Tangential Decomposition

From the local-frame vector components ($v_x$, $v_y$, $v_z$), the angular descriptors and decomposition metrics defined in Table 3 were derived for each interaction.

**Table 3:** Angular quantities and decomposition metrics derived from local-frame interaction vectors.

| Quantity | Formula | Interpretation |
|---|---|---|
| $\varphi_{xy}$ (rad/°) | $\arctan 2(v_y, v_x)$ | Azimuthal angle in the interface plane. |
| $\theta_{xz}$ (rad/°) | $\arctan 2(v_z, v_x)$ | Elevation angle in the $xz$ plane. |
| $v_\parallel$ | $\sqrt{v_x^2 + v_y^2}$ | Tangential component magnitude. |
| $Q$ | $\lvert v_z \rvert/(v_\parallel + \varepsilon)$ | Normal/tangential ratio; $Q \gg 1$ indicates perpendicular contact. |

Interactions were grouped by {protein, first_neighbor, condition, chain pair, interaction type} and the following circular statistics were computed over $\varphi_{xy}$ and $\theta_{xz}$ distributions for each group [46]:

- **Mean resultant length ($R$):** $R = \sqrt{\bar{c}^2 + \bar{s}^2}$, where $\bar{c}$ and $\bar{s}$ are the means of $\cos\theta$ and $\sin\theta$. $R \in [0, 1]$; values approaching 1 indicate directional coherence; values approaching 0 indicate isotropic dispersal.
- **Normalized circular entropy ($H_{\text{norm}}$):** Computed over an 18-bin histogram on $[-\pi, \pi]$ as $H_{\text{norm}} = -\sum_k p_k \ln p_k / \ln N_{\text{bins}}$. Values near 1 indicate uniform angular distribution; values near 0 indicate high directional concentration.
- **Circular mean ($\varphi_{\text{mean}}$):** $\arctan 2(\bar{s}, \bar{c})$ in degrees; preferred interaction direction in the interface plane.
- **Circular RMS deviation ($\varphi_{\text{rms}}$):** Root mean square of angular deviations around the circular mean (wrapped to $[-\pi, \pi]$), in degrees.
- **$Q_{\text{mean}}$ and $Q_{\text{median}}$:** Mean and median of the normal/tangential ratio across the group.

These metrics were consolidated into a single output file (`all_vectors_metrics_by_protein_neighbor_condition.csv`), with one row per unique combination of grouping variables. This file constitutes the primary quantitative output of the pipeline and is directly used for statistical comparisons between experimental conditions.

###### 2.12.5 Visualization and Serotype-Resolved Analysis

For each unique combination of protein, first neighbor, chain pair, and interaction type, a panel of six subplots was generated — one per experimental condition — containing: (i) a polar rose diagram (18 bins) of the azimuthal angle distribution $\varphi_{xy}$, representing the directional preference of interaction vectors within the interface plane; and (ii) a histogram of the elevation angle $\theta_{xz}$ in degrees, revealing the normal penetration character of the contacts. Three-dimensional quiver visualizations of the local interaction vectors were generated for each individual experiment, with bidirectional arrows scaled by vector length and the interface plane (z = 0 in the local frame) rendered with transparency to aid interpretation of normal versus tangential contact geometry.

For serotype-resolved analysis, interaction vectors from all experiments sharing a given dengue genotype (RQH index) were aggregated and a quadratic surface $z = a + bx + cy + dx^2 + exy + fy^2$ was fitted by least-squares to the vector tip coordinates ($\mathbf{r}_{\mathrm{tip}} = \mathbf{r}_{\mathrm{mid}} + \hat{u}L$, where $\hat{u}$ is the unit vector and $L$ the interaction length). When fewer than six tips were available, a plane was fitted by singular value decomposition (SVD) as fallback. The curvature and orientation of the fitted surface provides a macroscopic descriptor of the vectorial interaction field geometry for each genotype.

###### 2.12.6 Configurable Parameters and Batch Execution

The full pipeline is executed in five sequential stages (Table 5), each driven by a dedicated script and designed for batch processing over arbitrarily large sets of PDB files. Key parameters and their defaults are listed in Table 4.

**Table 4:** Configurable parameters of the VectorPROT pipeline.

| Parameter | Default | Description |
|---|---|---|
| `--hbond-cutoff` (Å) | 3.5 | Maximum interatomic distance for H-bond detection (N/O atoms). |
| `--hydrophobic-cutoff` (Å) | 5.0 | Maximum distance for hydrophobic contacts (side-chain C atoms). |
| `--interface-cutoff` (Å) | 6.0 | Radius for interface atom selection and local frame calculation. |
| `--bins` | 18 | Number of bins for rose diagrams and circular entropy calculation. |
| `--fit {plane\|surface}` | surface | Vector-tip fitting: SVD plane or least-squares quadratic surface. |
| `--surface-res` | 40 | Grid resolution ($N \times N$) for quadratic surface rendering. |

**Table 5:** Sequential execution stages of the VectorPROT pipeline.

| Stage | Script | Input | Primary Output |
|---|---|---|---|
| 1 | `vectorPROT_v_3_AB_BC_ABC.py` | Multi-chain PDB file | Per-experiment interaction CSVs and 3D qu[illegible] |
| 2 | `batch_vector.sh` | Root PDB directory | Batch execution over all PDB files |
| 3 | `concat_All_struc_V3.py` | Structured CSV tree | `all_vectors_structured.csv` + audit inde[illegible] |
| 4 | `Analise_combo_[...].py` | `all_vectors_structured.csv` | Enriched CSV, circular metrics, visualizatio[illegible] |
| 5 | `batch_serotype_planes.py` | Root directory + serotype map | Per-serotype 3D figures and concatenated ti[illegible] |

###### 2.12.7 Interpretation of Circular Metrics

The circular statistics produced by the pipeline are interpreted as follows in the context of the dengue envelope–host interaction analysis. $R$ values near 1 for $\varphi_{xy}$ indicate that the interaction vectors of a given group point coherently in a preferred azimuthal direction within the interface plane — a signature of geometrically oriented molecular recognition — whereas low $R$ ($< 0.3$) reflects isotropic dispersal consistent with non-specific or conformationally heterogeneous contacts. A significant reduction in $H_{\mathrm{norm}}$ between the PRE and POS conditions indicates that the presence of the viral envelope protein imposes a more restricted directional pattern on interfacial contacts, suggesting viral-induced conformational reorganization of the host interface. The $Q$ ratio differentiates compression-mode from sliding-mode interactions: $Q > 1$ identifies contacts with a dominant normal component (perpendicular to the interface plane), characteristic of high-affinity embedding; $Q < 1$ identifies predominantly tangential contacts, consistent with lateral surface scanning. Asymmetries in $Q$ between chain pairs AB and BC reveal differential recognition modes at the viral–host versus host–neighbor interfaces, providing mechanistic resolution beyond what is accessible from global binding free energies alone.

#### 2.13 Rationale for Triple Docking Simulations

Traditional protein–protein docking approaches evaluate interactions between two isolated proteins. However, biological protein interfaces frequently exist within multiprotein assemblies where binding of one protein can modify the accessibility, geometry, or energetic landscape of additional interaction interfaces [47]. To account for these cooperative effects, triple docking simulations were implemented, in which the dengue virus envelope protein was docked to a host protein already associated with one of its first neighbors, thereby forming a ternary complex.

Two complementary docking scenarios were investigated. In the **pre-assembly scenario**, docking simulations were first performed between the host protein and its first neighbor, producing a host–host complex representing the cellular interaction state prior to viral binding. The viral envelope protein was subsequently docked onto this complex. In the **post-assembly scenario**, the viral envelope protein was first docked onto the host protein, and a first-neighbor host protein was then docked onto the resulting complex to simulate the recruitment of additional host proteins following viral binding. Comparing these two scenarios provides insight into the directionality of interaction cooperativity within the interface network.

#### 2.14 Thermodynamic Analysis of Binding Energies

To quantify the energetic consequences of viral binding and host complex formation, binding free energies for the predicted complexes were estimated using post-docking energetic analysis [48]. The MM/GBSA method provides a computationally tractable approximation of binding free energy that is useful for ranking interactions and identifying mechanistic trends, but it systematically underestimates entropic contributions and overestimates absolute affinities, particularly for electrostatically dominated interfaces [49,50]. All $\Delta G$ values reported in this study should therefore be interpreted comparatively across complexes within the same structural class rather than as quantitative binding affinities, and $\Delta G$ magnitudes exceeding $-100$ $\mathrm{kcal \cdot mol^{-1}}$ are characteristic MM/GBSA overestimates for highly charged interfaces rather than physically meaningful absolute values.

The binding free energy of a binary complex is expressed as:

$$\Delta G_{\text{bind}} = G_{\text{complex}} - \left(G_{\text{protein}_1} + G_{\text{protein}_2}\right) \tag{5}$$

where $G_{\text{complex}}$ represents the free energy of the assembled complex and $G_{\text{protein}_1}$ and $G_{\text{protein}_2}$ represent the free energies of the individual proteins in the unbound state. To evaluate how the presence of neighboring proteins influences viral binding, the change in binding free energy between docking scenarios was quantified through $\Delta\Delta G$ calculations:

$$\Delta\Delta G = \Delta G_{\text{condition}_2} - \Delta G_{\text{condition}_1} \tag{6}$$

Negative $\Delta\Delta G$ values indicate that the second condition stabilizes the complex relative to the first, suggesting cooperative interactions that may enhance viral binding. Conversely, positive $\Delta\Delta G$ values indicate destabilization, consistent with the additional protein interaction introducing steric or energetic constraints that reduce binding affinity. Mapping $\Delta\Delta G$ values onto vector-based interaction networks allows identification of cooperative structural motifs: regions exhibiting both persistent hydrogen bond vectors and energetically favorable $\Delta\Delta G$ values represent candidate hotspots of structural cooperativity within the interaction network.

The AB/BC/ABC framework assumes pairwise decomposability of interaction energetics — that is, that the total free energy of the ternary complex can be approximated as the sum of binary contributions — and therefore approximates higher-order cooperativity rather than explicitly modeling three-body interaction terms [41,51]. This assumption is most valid when the three binding interfaces are spatially separated and conformationally independent. When all three proteins are in simultaneous contact, higher-order cooperative effects not captured by pairwise sums may be present. All $\Delta\Delta G$ values derived from this framework should be interpreted as directional indicators of assembly-order effects rather than as quantitative measurements of three-body cooperativity.

#### 2.15 Integrated Analytical Pipeline for Ternary Docking Data

##### 2.15.1 AB/BC/ABC Interaction Framework

Docking results were analyzed through a structured AB/BC/ABC framework that models sequential stages of ternary complex assembly. Four interaction contexts were defined: (i) the AB context, representing the binary viral–host interaction that establishes the baseline binding affinity; (ii) the BC context, representing the native host–neighbor interaction prior to viral engagement; (iii) the ABC_post_BC_A scenario (Triple), in which the viral protein docks onto a pre-formed BC complex; and (iv) the ABC_post_AB_C scenario (Triple_Inverse), in which the first-neighbor host protein is recruited to a pre-formed AB complex. This bidirectional design enables discrimination between cooperative binding, competitive displacement, and independent interaction modes [43,47], and accommodates replicate docking experiments for reproducibility assessment.

##### 2.15.2 Data Preprocessing and Thermodynamic Standardization

Primary docking outputs were provided as structured tabular data with mandatory fields for interaction pair identity, context label, and binding free energy ($\Delta G$, kcal · mol$^{-1}$). When the dissociation constant ($K_d$) was available, bidirectional conversion between $\Delta G$ and $K_d$ was performed using the standard thermodynamic relation $\Delta G = RT \ln K_d$, with $R = 1.9872 \times 10^{-3}$ kcal · mol$^{-1}$ · K$^{-1}$ and $T = 310.15$ K [48]. Interaction pair identifiers were parsed to handle multiple separator conventions, and context labels were harmonized to the canonical AB/BC/ABC notation to ensure compatibility across input datasets.

##### 2.15.3 Differential Binding Energetics and Thermodynamic Coupling

Differential binding free energies ($\Delta\Delta G$) were computed for two primary transitions: $\Delta\Delta G_{\mathrm{AB} \to \mathrm{ABC}}$ (change in viral binding energy upon encountering a pre-formed BC complex) and $\Delta\Delta G_{\mathrm{BC} \to \mathrm{ABC}}$ (change in host–neighbor binding energy upon viral pre-engagement). The thermodynamic coupling parameter was calculated as $\alpha = \exp(\Delta\Delta G/RT)$, providing a dimensionless index of binding cooperativity. Values of $\alpha < 1$ indicate mutual stabilization; $\alpha > 1$ indicates competitive destabilization; $\alpha \approx 1$ indicates independent binding events [52]. An order-asymmetry parameter was defined as the difference between the two $\Delta\Delta G$ values, quantifying the degree to which the thermodynamic outcome depends on the temporal order of complex assembly. Perturbations with $|\Delta\Delta G| > 1.0$ kcal · mol$^{-1}$ were classified as energetically significant, consistent with thresholds established for computational cooperativity analyses [43].

###### 2.15.4 Integration with Protein Interaction Networks and Functional Annotation

Docking-derived interaction pairs were annotated using the STRING protein interaction database (v12.0) [53], which provides pairwise confidence scores aggregating co-expression, genomic co-occurrence, text mining, and experimental evidence. Each edge was tagged with the STRING combined confidence score (range 0–1000) and the experimentally determined interaction score, enabling downstream stratification of predictions by network support level. Protein functional annotations were derived from Gene Ontology (GO) enrichment data, with an FDR threshold of $\leq 0.01$ applied to retain only statistically robust biological process terms [54]. An edge-level GO score was computed as the arithmetic mean of the GO enrichment strengths of the two interacting proteins, providing a functional relevance metric for each predicted interaction.

###### 2.15.5 Network Topology Analysis

Protein interaction networks were constructed as undirected weighted graphs using the igraph package [55], with edge weights corresponding to absolute $\Delta G$ values. Three standard centrality metrics were computed for each node: betweenness centrality (proportion of shortest paths traversing each node), closeness centrality (inverse mean shortest path distance), and degree centrality (number of direct connections), collectively identifying hub proteins and structural bridges within the interaction architecture [56]. Community structure was resolved using the Louvain modularity optimization algorithm, which detects densely interconnected subgraphs without requiring *a priori* specification of cluster number [57].

###### 2.15.6 Statistical Analysis

Thermodynamic differences between interaction contexts were assessed using paired Wilcoxon signed-rank tests, a non-parametric approach appropriate for binding free energy data, which frequently deviate from normality [58]. Tests were conducted both within individual protein systems and globally across all cases, requiring a minimum of three paired observations. Where contact-type data (intermolecular contacts, ICs; non-interacting surface residues, NIS) were available, supervised machine learning models were trained to identify structural predictors of thermodynamic outcomes. LASSO regression with five-fold cross-validation [59] provided sparse linear models identifying the most predictive contact categories. Random Forest regression [60] with 500 trees and permutation-based variable importance provided complementary non-linear models robust to feature colinearity. A minimum of ten observations was required for model training.

##### 2.15.7 Integrative Scoring and Target Prioritization

An integrative composite score was computed for each interaction by normalizing and linearly combining three orthogonal metrics: binding free energy ($z$-scored), STRING combined confidence ($z$-scored), and GO enrichment strength ($z$-scored). Default weights were $w_{\Delta G} = 0.6$, $w_{\text{STRING}} = 0.3$, and $w_{\text{GO}} = 0.1$, reflecting the primary importance of thermodynamic predictions in docking-based studies while incorporating network and functional context. A separate target ranking score combined the absolute $|\Delta\Delta G|$ magnitudes for both binding transitions (weighted equally at 45% each) with a reproducibility robustness metric across replicates (10%), producing a prioritized candidate list for experimental follow-up.

##### 2.15.8 Reproducibility and Computational Implementation

All analyses were implemented in R (v4.3+) within a reproducible R Markdown workflow [61]. Data manipulation and visualization utilized the tidyverse ecosystem [62]. Network analysis employed igraph [55]; regularized regression used glmnet [63]; ensemble modeling used ranger [64]; graph visualization used ggraph [65]. Fixed random seeds (default: 1) ensured reproducibility of stochastic components including cross-validation fold assignments and Random Forest ensemble construction. Quality control flags were applied to extreme values ($|\Delta G| > 40$ kcal $\cdot$ mol$^{-1}$; $K_d < 10^{-15}$ M or $K_d > 10^{-3}$ M) to document potential outliers without automatic exclusion. Complete source code is provided in the Supplementary Information.

### 3 Results

#### 3.1 Functional Architecture of the Host Protein Network

##### 3.1.1 The Selected Host Proteins Form an Interconnected Signaling and Homeostasis Network

The thirteen host proteins selected for this study constitute an integrated cellular network in which signaling inputs, structural responses, and homeostatic mechanisms are functionally coupled across multiple layers of cellular organization (Figure 2). Understanding these functional interconnections is essential for interpreting the biological significance of individual envelope–host protein interactions, since viral perturbation of one node in the network will propagate through existing molecular relationships to affect connected partners. The network can be organized into four functionally distinct but heavily interconnected clusters: growth factor signaling and co-receptor complexes, intracellular trafficking and membrane dynamics, cell structure and adhesion, and protein homeostasis with inflammatory surveillance.

The growth factor signaling cluster is anchored by two extracellular ligand systems — VEGF and TGF-$\beta$ — whose activities are modulated by dedicated co-receptors before reaching canonical signaling receptors. Neuropilin-1 acts as a co-receptor that presents VEGF to VEGFR, potentiating pro-angiogenic and vascular permeability responses directly relevant to dengue-associated plasma leakage [19,22]. In parallel, Betaglycan (TGFBR3) presents TGF-$\beta$ ligands — particularly the TGF-$ $2 isoform — to TGFBR1, enabling assembly of the canonical signaling receptor complex [66,67]. Glypican-1 operates at the intersection of both ligand systems as a heparan sulfate proteoglycan capable of sequestering or presenting VEGF and FGF2, thereby fine-tuning the magnitude and spatial distribution of growth factor signaling at the cell surface [11]. Together, these co-receptors constitute an integrated surface-sensing layer whose collective activity regulates endothelial permeability, immune modulation, and extracellular matrix remodeling — all processes dysregulated during severe dengue disease.

The intracellular trafficking cluster is organized around Rab5 as the master regulator of early endosome identity and function. Rab5 GTPase activity controls vesicle docking, tethering, and fusion at the early endosomal compartment, directing the intracellular fate of internalized signaling receptors, including TGFBR1 complexes, as well as viral particles following endocytosis [17,18]. Glypican-1 and Betaglycan are themselves substrates of endocytic recycling regulated by Rab5-dependent vesicular transport, creating bidirectional coupling between the surface co-receptor layer and the endosomal trafficking network. Actin provides the structural scaffold for this trafficking machinery through its roles in membrane deformation, vesicle budding, and intracellular cargo transport [15].

The cell structure and adhesion cluster connects intracellular signaling outputs to macroscopic cellular organization. Actin filament dynamics are a principal downstream effector of growth factor receptor activation, translating chemical signals into morphological changes including junction remodeling and endocytic membrane deformation. Claudin-1 has dengue-specific experimental support through prM/M-dependent entry assays [24] and functions as a structural component of tight junctions physically anchored to the actin cytoskeleton through scaffold proteins, creating a mechanical link between signal-responsive actin dynamics and epithelial/endothelial barrier integrity [23].

The homeostasis and inflammatory surveillance cluster monitors cellular stress and coordinates survival or death responses. BiP/GRP78 is the central sensor of the unfolded protein response (UPR) in the ER, detecting misfolded protein accumulation induced by the high demand of viral protein production [68,69]. Clusterin functions as an extracellular chaperone that clears aggregated proteins; its intracellular isoform directly interacts with BiP to suppress stress-induced apoptosis, forming a functional axis linking extracellular proteostasis to intracellular ER stress responses [28]. NLRP3 operates downstream of UPR signals and perturbed actin dynamics as an intracellular danger sensor whose inflammasome activation drives the inflammatory pathology associated with dengue [26]. SUMO post-translational modification regulates transcription factors within TGF-$\beta$ and NF-$\kappa$B pathways, as well as the stability of NLRP3 itself, acting as a molecular rheostat that tunes pathway amplitude [27]. NKp44 closes the circuit from cellular stress to immune recognition, and its reported interaction with dengue and West Nile envelope glycoproteins provides a direct rationale for inclusion in this host-factor panel [25,70].

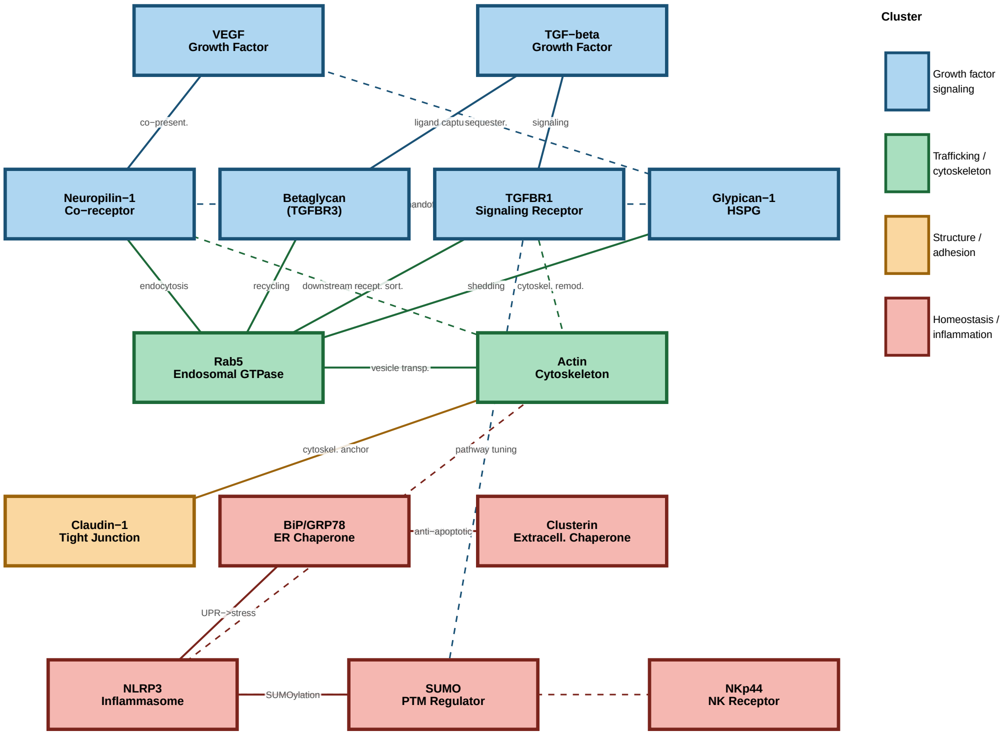


**Figure 2:** Functional interaction network of host proteins selected for envelope docking analysis. Nodes are color-coded by functional cluster: blue, growth factor signaling and co-receptors; green, intracellular trafficking and membrane dynamics; orange, cell structure and adhesion; red, protein homeostasis and inflammatory surveillance. Solid arrows indicate direct, experimentally supported functional relationships; dashed arrows indicate indirect or context-dependent connections. Envelope interaction overlays will be added in subsequent analyses.

##### 3.1.2 Network Topology Predicts Pleiotropic Consequences of Envelope Engagement

The network topology described above has direct implications for interpreting the results of envelope–host protein docking and molecular dynamics analyses. Because the selected proteins occupy distinct positions within a shared interaction network, envelope binding to any individual node is unlikely to produce isolated effects. Instead, disruption or stabilization of a single interaction can propagate through established functional edges, producing cascading consequences for connected partners. Three network properties are particularly relevant in this regard.

First, Actin, Rab5, and TGFBR1 function as connectivity hubs — nodes with high functional connectivity that are disproportionately influential on network behavior. Each participates simultaneously in multiple functional clusters, meaning that envelope engagement with any of these proteins could perturb trafficking, structural, and signaling processes in parallel. The characterization of binding affinity and interface architecture for these hub proteins is therefore of heightened biological significance relative to peripheral network participants.

Second, the network contains feedback loops whose disruption may produce non-linear inflammatory consequences. The BiP–Clusterin axis and the NLRP3–SUMO regulatory relationship are both feedback-stabilized circuits; envelope engagement with either BiP or NLRP3 may contribute to inflammatory amplification, consistent with the cytokine storm phenotype observed in severe dengue, though the magnitude of any such effect would be modulated by the macromolecular crowding of the cellular environment, which can substantially alter effective association kinetics and stabilize interaction states not captured in dilute-solution simulations [71,72].

Third, the network converges at Actin as a structural integration point receiving inputs from both signaling clusters and feeding outputs into trafficking, adhesion, and homeostasis. This centrality means that the properties of the envelope–actin interface have functional relevance extending well beyond simple cytoskeletal anchorage, motivating the detailed characterization presented in the following section.

An important interpretive framework for the subsequent analysis is the functional classification of the host proteins examined. Not all thirteen proteins interact with the viral envelope at the same stage of infection or through the same biological mechanism [73,74]. Three functional classes are distinguished throughout this discussion: (i) *attachment and surface-concentration factors* — GPC1, NRP1, and BiP — whose primary predicted role is to concentrate or stabilize viral particles at the plasma membrane before productive internalization; (ii) *trafficking and intracellular scaffold proteins* — actin and Rab5 — which are implicated in endocytic and post-endocytic transport rather than primary attachment; and (iii) *signaling and pathogenesis-associated proteins* — TGFBR1, NLRP3, and the SUMO/UBC9 axis — which do not participate directly in viral entry but are implicated in the immunopathological consequences of dengue infection. Conclusions about the biological relevance of interactions in each class must be calibrated to the appropriate biological context: an interaction that is thermodynamically favorable in silico does not imply an entry receptor function unless the protein is accessible at the plasma membrane during the relevant stage of infection. Subsequent sections systematically characterize each envelope–host protein interaction within this network framework, proceeding from cytoskeletal and chaperone interactions to co-receptor complexes, trafficking regulators, and immune sentinels.

#### 3.2 Ligand-specific modulation of dengue envelope dynamics reveals distinct regimes of structural stabilization and conformational remodeling

To investigate how the dengue virus envelope protein responds to different host-associated ligands, we compared backbone polar surface exposure, average backbone polar surface area, radius of gyration, RMSD, and RMSF for both the envelope protein, treated here as the receptor, and each interacting ligand. The comparative analysis revealed that the dengue envelope protein retains a globally coherent structural behavior across all complexes, but its local flexibility, compactness, and conformational drift are clearly modulated by the identity of the ligand. Conversely, the ligands displayed markedly heterogeneous dynamic responses, indicating that each host factor imposes a distinct interaction regime on the viral envelope surface.

Across all systems, the receptor-side trajectories showed that the dengue envelope protein does not behave as a passive binding scaffold. Instead, the envelope undergoes ligand-dependent conformational adjustments that differ in magnitude, persistence, and spatial distribution. Although none of the complexes showed evidence of complete structural disruption of the envelope, several ligands induced measurable shifts in receptor compactness, backbone exposure, and residue-level flexibility. These observations support a model in which host-protein engagement by the dengue envelope is governed not simply by static complementarity at the docking interface, but by dynamic coupling between ligand binding and envelope conformational plasticity.

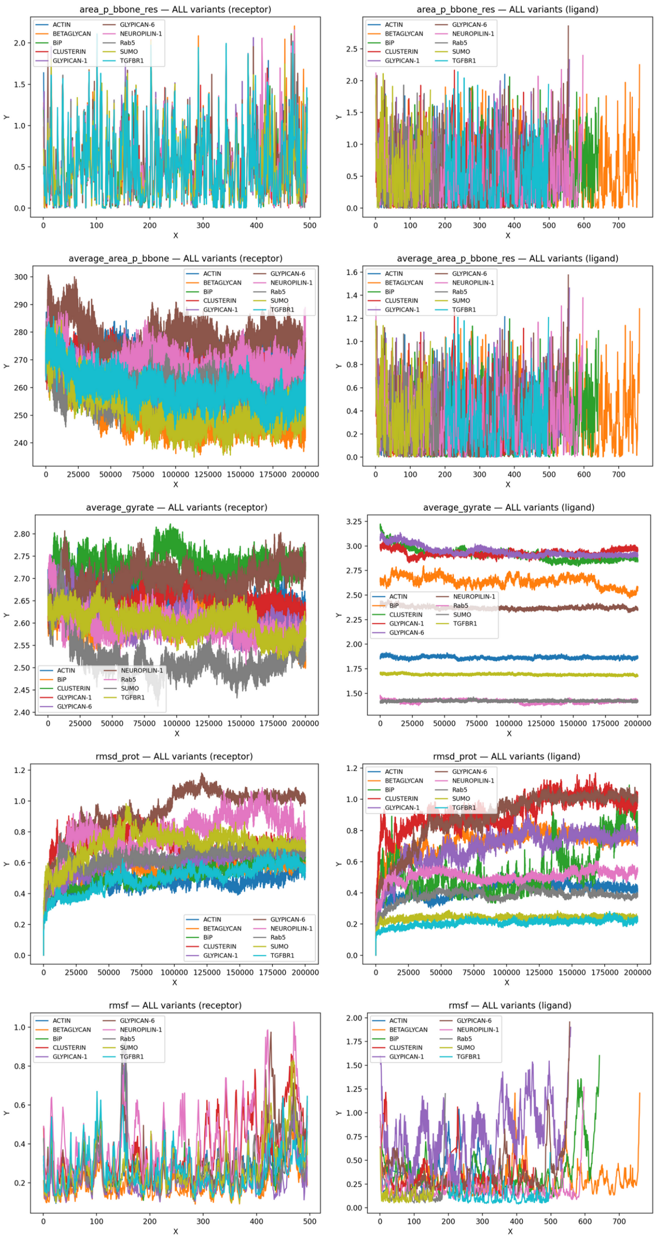


**Figure 3: Comparative molecular dynamics descriptors of dengue envelope–host ligand complexes.** The left panels show receptor-side descriptors for the dengue envelope protein, whereas the right panels show the corresponding ligand-side descriptors. The analyzed metrics include polar backbone area, average polar backbone exposure, radius of gyration, RMSD, and RMSF, revealing ligand-specific modulation of envelope compactness, conformational drift, and local flexibility.

As shown in Figure~3, dengue envelope dynamics were markedly modulated by ligand identity. Time- and residue-resolved molecular dynamics descriptors were calculated for dengue virus envelope protein complexes with multiple host-associated ligands, including ACTIN, BETAGLYCAN, BiP, CLUSTERIN, GLYPICAN-1, GLYPICAN-6, NEUROPILIN-1, Rab5, SUMO, and TGFBR1. For each descriptor, the left panels show the behavior of the dengue envelope protein treated as the receptor, whereas the right panels show the corresponding behavior of the interacting ligand. The analyses include residue-level polar backbone area (`area_p_bbone_res`), average polar backbone area (`average_area_p_bbone_res`), radius of gyration (`average_gyrate`), protein backbone RMSD (`rmsd_prot`), and residue-level RMSF (`rmsf`). Together, these metrics describe changes in backbone exposure, compactness, global conformational drift, and local flexibility during the simulations. The upper panels show residue-level polar backbone area for the receptor and ligand. The dengue envelope protein displays localized fluctuations in backbone polar exposure, suggesting that ligand binding mainly perturbs discrete regions of the receptor rather than inducing uniform exposure across the full structure. In contrast, the ligands show broader and more heterogeneous polar exposure profiles, indicating stronger local conformational breathing and differential solvent accessibility among host factors. The second row shows the average polar backbone area. The receptor maintains a comparatively coherent range of average exposure across complexes, consistent with preservation of a stable envelope scaffold, whereas the ligand profiles are more discontinuous and ligand-specific, reflecting differences in size, topology, interface engagement, and local flexibility. The third row shows the average radius of gyration. The dengue envelope protein remains within a bounded range across all complexes, indicating absence of global unfolding or collapse, but the separation among trajectories reveals ligand-specific modulation of receptor compactness. Some ligands maintain the envelope in a more compact state, whereas others promote a more expanded or fluctuating receptor conformation. The ligand-side radius of gyration separates the host factors into distinct structural regimes, ranging from compact and stable proteins to more extended or dynamically relaxed ligands. The fourth row shows protein RMSD for receptor and ligand components. The receptor RMSD profiles indicate that the dengue envelope undergoes different degrees of ligand-dependent conformational relaxation. Lower RMSD trajectories are consistent with stabilizing or structurally compatible interactions, whereas higher RMSD trajectories suggest stronger receptor remodeling or induced-fit behavior. The ligand RMSD profiles are more divergent, with some ligands remaining structurally stable and others undergoing larger conformational rearrangements. These ligand-side deviations may reflect domain mobility, interface reorganization, or flexible peripheral regions rather than complete complex dissociation. The bottom panels show RMSF profiles. The receptor exhibits localized flexibility hotspots rather than uniform destabilization, indicating that ligand binding selectively modulates flexible envelope regions. The ligand RMSF profiles are markedly more heterogeneous, with several ligands showing high-amplitude fluctuations in specific regions, consistent with flexible loops, terminal segments, or domains not fully constrained by the interface. Overall, the figure supports a model in which the dengue envelope behaves as a structurally conserved but dynamically responsive receptor, while the host ligands display interaction-specific degrees of compactness, flexibility, and conformational adaptation.

##### 3.2.1 Backbone polar exposure indicates local remodeling of the receptor interface and stronger conformational breathing in the ligands

The residue-level polar backbone area profiles showed highly localized fluctuations in the receptor across the different complexes. For the envelope protein, most values remained within a relatively constrained range, with intermittent peaks distributed along the sequence. This pattern suggests that ligand binding perturbs specific receptor regions rather than causing broad and uniform exposure of the backbone. The presence of sharp local peaks is compatible with transient exposure of loops, hinge-like elements, or flexible interface-adjacent segments.

In contrast, the corresponding ligand-side profiles were more heterogeneous and frequently displayed higher-amplitude fluctuations. This indicates that the ligands undergo more pronounced local rearrangements upon interaction with the envelope. These ligand-side changes may reflect partial solvation of interface regions, transient disengagement and re-engagement of peripheral contacts, or conformational breathing of flexible domains that are not fully constrained by the receptor interface.

The averaged receptor backbone polar area further supports this interpretation. On the receptor side, average values remained broadly organized within a relatively stable band across the simulation, indicating that the dengue envelope maintains a conserved global level of polar backbone exposure despite ligand-specific perturbations. However, the individual trajectories were not identical. Certain ligands appeared to shift the receptor toward higher average exposure, whereas others maintained lower and more stable values. This suggests that different host factors alter the hydration and exposure state of the envelope surface to different degrees.

On the ligand side, the averaged polar backbone area was much lower in absolute terms and more punctuated by abrupt local changes. This difference between receptor and ligand behavior is important. It suggests that the envelope distributes the perturbation across its structure, whereas the ligands respond through more localized conformational adjustments. In other words, the envelope appears dynamically buffered, while many ligands retain more pronounced local mobility.

##### 3.2.2 Radius of gyration separates receptor responses into compact and expanded conformational regimes

The radius of gyration provided one of the clearest indicators of ligand-dependent receptor remodeling. For the envelope protein, the average radius of gyration remained within a relatively narrow but biologically meaningful range, approximately between 2.45 and 2.80 nm. This indicates that the receptor did not undergo global unfolding or collapse in any of the simulated complexes. Nevertheless, the separation among trajectories shows that ligand binding can bias the envelope toward more compact or more expanded conformational states.

Several complexes maintained the envelope in a comparatively compact state, with lower radius of gyration values and reduced large-scale expansion. These systems likely represent interactions that are structurally compatible with the basal conformation of the envelope, imposing relatively little global strain on the receptor. In contrast, other ligands promoted higher receptor radius of gyration values or broader fluctuations, suggesting partial expansion, relaxation of interdomain packing, or opening of flexible envelope regions.

The ligand-side radius of gyration profiles showed even stronger separation among systems. The ligands clustered into distinct structural classes. Some remained highly compact, with low and stable radius of gyration values throughout the trajectory. Others occupied intermediate values, consistent with moderate structural flexibility. A third group displayed substantially higher radius of gyration values, indicating more extended architectures or greater conformational relaxation during the simulation.

This contrast between receptor and ligand behavior is informative. The envelope response is comparatively constrained, suggesting preservation of a stable structural core. The ligands, however, retain their own intrinsic architectural signatures and differ substantially in their ability to remain compact or undergo expansion. Thus, the global dynamics of each complex appear to arise from the combination of a moderately plastic viral receptor and ligands with highly variable intrinsic flexibility.

###### 3.2.3 RMSD analysis identifies ligand-dependent stabilization and remodeling of the dengue envelope

The receptor RMSD profiles showed a general increase during the early-to-intermediate stages of the simulations, followed by different degrees of stabilization depending on the ligand. This behavior is consistent with relaxation from the initial modeled complex toward ligand-specific conformational basins. Importantly, the final RMSD amplitudes differed substantially among complexes, indicating that the envelope does not converge toward a single universal bound-state conformation.

Some ligands maintained the envelope at relatively low RMSD values, approximately in the 0.45–0.65 nm range after equilibration. These complexes can be interpreted as receptor-stabilizing or receptor-compatible interactions. In such cases, the ligand appears to bind without driving extensive global rearrangement of the envelope. These systems are likely to preserve a relatively stable interface and may represent interaction modes with lower conformational penalty.

Other ligands induced larger receptor deviations, with RMSD values approaching or exceeding approximately 0.8–1.0 nm. These trajectories suggest more substantial conformational remodeling of the envelope. Such behavior does not necessarily indicate an unstable or biologically irrelevant interaction. Rather, it may reflect induced fit, conformational selection, or ligand-driven displacement of flexible envelope regions. In the context of dengue envelope biology, such remodeling is potentially important because relatively subtle shifts in envelope domain orientation, loop exposure, or local surface geometry may influence receptor recognition, epitope accessibility, or downstream entry-related transitions.

The ligand RMSD profiles were more divergent than the receptor profiles. Several ligands remained conformationally stable, with low RMSD values throughout most of the simulation. Others displayed progressive increases or high-amplitude fluctuations, reaching values near or above 1.0 nm. These high ligand RMSD values may indicate structural reorganization, domain motion, or partial interface rearrangement. However, they should not be interpreted in isolation as evidence of binding failure. A ligand may show high global RMSD while still maintaining a stable interface through a subset of anchored residues or domains. This is particularly relevant for multidomain or elongated host proteins, where flexible distal regions may dominate the global RMSD while the actual contact region remains stable.

Therefore, the RMSD data suggest at least two major interaction regimes: one in which the ligand stabilizes the envelope with limited receptor rearrangement, and another in which binding is accompanied by larger conformational remodeling of either the envelope, the ligand, or both.

###### 3.2.4 RMSF profiles reveal localized flexibility hotspots rather than uniform receptor destabilization

Residue-level RMSF analysis showed that ligand-induced changes in the envelope are spatially localized. Most receptor residues exhibited moderate fluctuations, whereas defined regions displayed pronounced RMSF peaks. This indicates that ligand binding does not globally destabilize the envelope, but instead modulates discrete flexible elements. Such regions may correspond to exposed loops, terminal regions, interdomain connectors, or interface-proximal segments that accommodate ligand binding.

A particularly notable feature was the presence of stronger receptor fluctuations toward the latter portion of the sequence axis, where several complexes showed higher RMSF peaks. This suggests that specific envelope regions are especially sensitive to ligand identity. If these peaks correspond to functionally relevant structural elements, such as exposed envelope loops, hinge-like segments, or regions involved in receptor engagement, they may represent important determinants of host-factor-specific envelope modulation.

The ligand RMSF profiles showed much greater heterogeneity. Some ligands maintained low fluctuations across most residues, consistent with rigid or well-anchored binding. Others displayed large RMSF peaks, in some cases approaching or exceeding 1.5–2.0 nm. These high-amplitude peaks likely represent flexible terminal segments, mobile loops, or domains not directly stabilized by the envelope interface. This observation supports a model in which many host proteins bind the envelope through a restricted interaction core while retaining substantial mobility in peripheral regions.

Together, the RMSF data argue against a simple binary classification of complexes as stable or unstable. Instead, the results support a more nuanced model in which stable interface cores coexist with flexible peripheral regions. This is consistent with the behavior expected for protein–protein complexes involving large or multidomain host factors, where binding may be mediated by a limited set of persistent contacts embedded within a broader dynamic ensemble.

###### 3.2.5 Comparative ligand behavior suggests distinct functional classes of envelope engagement

The combined descriptors allowed the complexes to be organized into preliminary dynamic classes.

A first class comprises ligands that appear to stabilize the envelope or maintain it close to a compact, low-deviation state. These systems show lower receptor RMSD, controlled radius of gyration, and moderate receptor RMSF. Interactions of this type likely impose a limited conformational burden on the envelope and may represent structurally compatible binding modes. In such complexes, ligand recognition may occur without substantial rearrangement of the viral surface.

A second class comprises ligands that induce stronger receptor remodeling. These complexes display increased receptor RMSD, higher or more variable radius of gyration, and enhanced local RMSF peaks. Such ligands may promote more open or flexible envelope states. This behavior may reflect less stable binding, but it may also indicate biologically relevant conformational selection. For host factors involved in viral attachment, concentration at the cell surface, or presentation to entry receptors, the ability to alter envelope conformational sampling may be functionally meaningful.

A third class includes ligands that show high intrinsic mobility while the envelope remains comparatively constrained. In these cases, high ligand RMSD or RMSF likely reflects motion of domains or segments outside the interface rather than complete loss of binding. These complexes should be analyzed with interface-specific metrics, because global ligand descriptors may overestimate instability when distal flexible regions dominate the signal.

This classification is provisional but useful. It indicates that different host proteins do not engage the dengue envelope through equivalent dynamic mechanisms. Instead, each interaction imposes a characteristic balance between receptor stabilization, ligand flexibility, interface persistence, and conformational remodeling.

###### 3.2.6 Asymmetric dynamic coupling between envelope and ligands

A central observation from the full dataset is the asymmetry between receptor and ligand responses. The dengue envelope protein exhibits constrained but ligand-sensitive plasticity. Its radius of gyration and RMSD remain within a bounded range, and its RMSF profile suggests localized rather than global flexibility. The ligands, however, show much broader variation in compactness, RMSD, and residue-level fluctuation.

This asymmetry suggests that the envelope functions as a relatively robust viral scaffold capable of accommodating different host factors through local remodeling. The host ligands, in contrast, absorb much of the dynamic variability of the complex. This may be especially relevant for large extracellular, membrane-associated, or multidomain host proteins, where binding to the viral envelope may involve only a subset of residues while other regions remain mobile.

Such behavior is consistent with a dynamic recognition model. Rather than forming uniformly rigid complexes, the envelope and its ligands appear to form interaction ensembles composed of stable contact cores, flexible peripheral segments, and ligand-specific conformational states. This type of organization could allow dengue virions to interact with structurally diverse host factors while preserving the core architecture of the envelope protein.

###### 3.2.7 Implications for dengue envelope recognition and host-factor specificity

The results suggest that dengue envelope recognition by host-associated proteins is not governed solely by static interface complementarity. Instead, the envelope appears to respond to different ligands through distinct dynamic programs. Some ligands maintain a compact and relatively stable receptor conformation, whereas others promote expansion, local flexibility, or larger conformational drift. These differences may have functional implications for viral attachment, receptor engagement, immune accessibility, and downstream entry-related transitions.

Ligands that stabilize the envelope may represent interactions compatible with pre-existing envelope conformations. These may correspond to binding events that favor persistent attachment without extensive structural rearrangement. By contrast, ligands that increase receptor RMSD, radius of gyration, or local RMSF may promote alternative envelope states. Such states could expose otherwise transient surfaces, alter the geometry of receptor-binding regions, or facilitate subsequent engagement with additional host factors.

The stronger variability observed in the ligands also suggests that host-factor identity strongly influences the architecture of the complex. Some ligands appear to bind as compact and stable partners, whereas others behave as flexible scaffolds with mobile regions surrounding a more restricted interface. This distinction is important because global stability metrics may underestimate the biological relevance of flexible ligands. For these systems, persistent local contacts, interface area, and energetic decomposition are likely more informative than whole-ligand RMSD alone.

###### 3.2.8 Summary model

These analyses support a model in which the dengue envelope protein maintains a conserved structural core while sampling ligand-specific conformational states at its surface. The envelope is neither rigid nor globally destabilized. Instead, it behaves as a dynamically responsive receptor whose local flexibility and compactness are tuned by the interacting host factor. The ligands, in turn, exhibit a broader range of behaviors, from compact and stable binding to highly flexible, domain-mobile association.

This dynamic heterogeneity suggests that dengue envelope–host factor interactions should be interpreted as ensembles rather than single static complexes. The most relevant biological signal is likely contained in the combination of receptor conformational remodeling, ligand flexibility, interface persistence, and energetic stability. Therefore, the present results provide a mechanistic basis for prioritizing subsequent analyses focused on interface-specific RMSD, center-of-mass distances, hydrogen-bond occupancy, buried surface area, MM/GBSA decomposition, and network-based interaction mapping. These additional layers will be essential to distinguish genuinely unstable complexes from biologically meaningful dynamic interactions that rely on flexible host-factor engagement.

##### 3.3 Envelope–Actin Complex: Structural Stability and Interface Persistence

###### 3.3.1 Global Conformational Stability of the Complex

Molecular dynamics simulations of the complex formed between the viral envelope protein (chain A) and cellular actin (chain B) demonstrate rapid convergence to a stable conformational regime following the initial equilibration phase. The envelope protein exhibits a mean root mean square deviation (RMSD) of approximately 0.48 nm over the complete 200 ns trajectory, with a modest increase to ~0.55 nm in the final 20 ns, reflecting late-stage structural relaxation rather than progressive destabilization or global unfolding. Actin displays a comparable but marginally lower mean RMSD of ~0.41 nm, with similarly stable behavior in the final trajectory segment (~0.43 nm), indicating that complex formation does not impose disruptive conformational perturbations on either partner.

Analysis of the radius of gyration (Rg) reinforces these observations. The envelope protein undergoes a subtle reduction in Rg from a mean value of ~3.14 nm to approximately 3.11 nm toward the end of the simulation, indicative of mild interfacial compaction consistent with progressive surface accommodation. Actin maintains a virtually constant Rg of ~2.18 nm throughout the trajectory, excluding large-scale folding or expansion events attributable to the binding interaction. Taken together, the RMSD and Rg trajectories indicate that both proteins retain their global structural identities within the complex, with localized surface adjustments constituting the primary mode of interfacial adaptation.

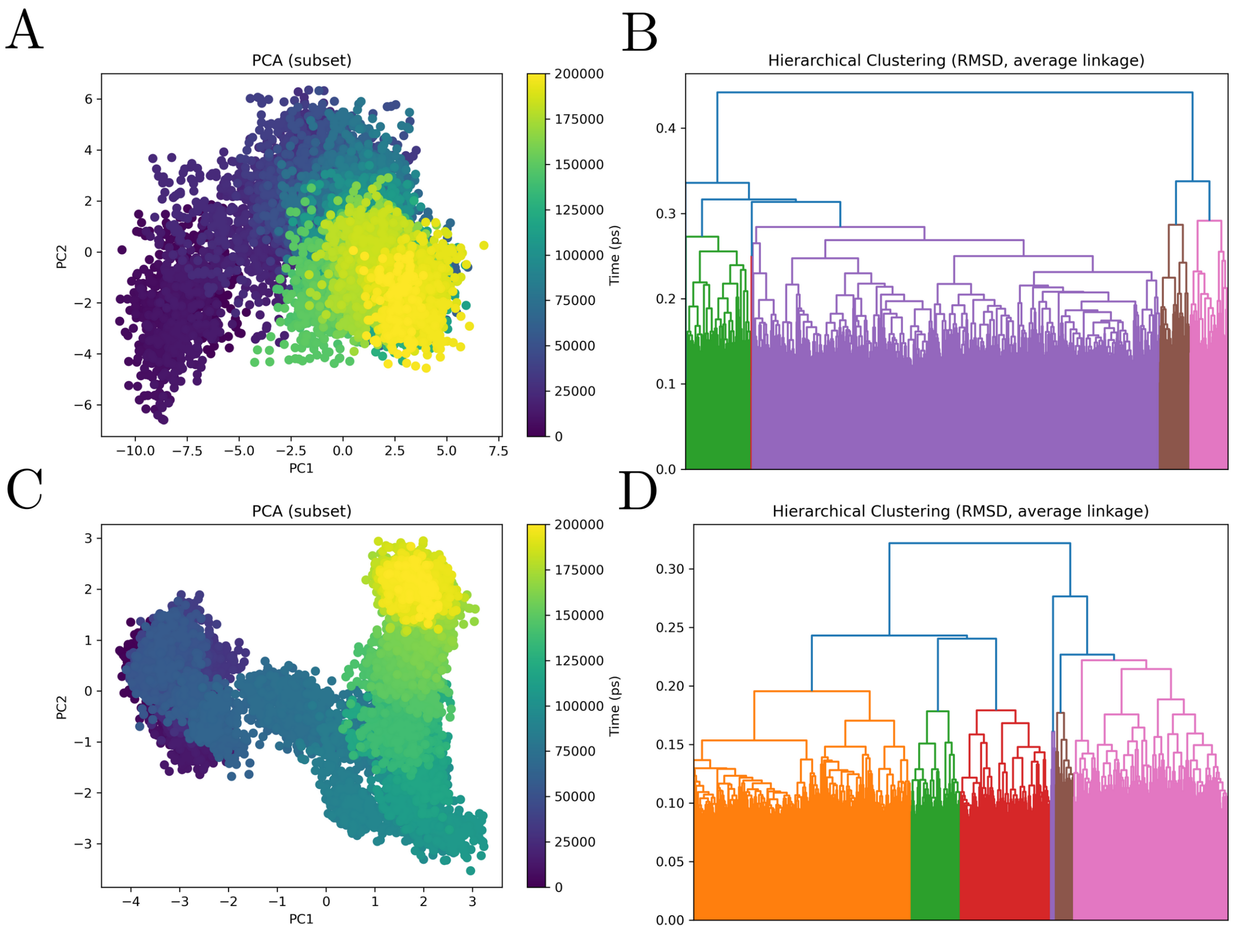


**Figure 4: Principal component analysis and RMSD-based hierarchical clustering of the dengue Envelope–actin molecular dynamics trajectory.** The figure summarizes the conformational landscape of the dengue Envelope–actin complex, with AB corresponding to dengue Envelope and CD corresponding to actin. Panels A and C show principal component analysis projections of trajectory frames onto the first two principal components, with points colored according to simulation time. The temporal gradients indicate that the system undergoes progressive conformational redistribution rather than remaining confined to a single static basin. Panel A shows a broad conformational manifold consistent with extensive Envelope-associated structural drift and sampling of multiple partially overlapping substates. Panel C shows a more clearly organized transition between early, intermediate, and late conformational regions, consistent with actin-associated accommodation within the complex. Panels B and D show RMSD-based hierarchical clustering using average linkage, revealing multiple structural families within the trajectory. The dendrograms indicate that both the Envelope-associated and actin-associated subsets sample distinct conformational clusters, supporting a model in which the Envelope–actin interface is dynamically heterogeneous but structurally organized. Together, the PCA and clustering analyses indicate that the complex is not represented by a single rigid binding pose, but by an ensemble of related conformational states shaped by reciprocal adaptation between dengue Envelope and actin.

##### 3.3.2 Residue-Level Flexibility Reveals Asymmetric Conformational Partitioning

Residue-level flexibility analysis through root mean square fluctuation (RMSF) calculations reveals a pronounced asymmetry in the conformational dynamics of the two binding partners. The envelope protein displays restrained per-residue fluctuations with a mean RMSF of approximately 0.26 nm, and localized peaks confined to surface-exposed regions that do not exceed ~0.55 nm. This pattern is consistent with the established structural robustness of the flavivirus E protein, whose $\beta$-rich architecture and internal hydrogen bond networks confer conformational rigidity even in the presence of interacting partners [3,5].

In contrast, actin exhibits substantially higher RMSF values, with a mean of approximately 0.40 nm and discrete peaks exceeding 1.0 nm in localized regions, including segments associated with the subdomain 2–4 boundary and the N-terminal region — regions documented to be intrinsically flexible and functionally important for actin's interactions with regulatory proteins [75]. Critically, these high-amplitude fluctuations are spatially confined rather than globally distributed, indicating that actin accommodates the interface through localized conformational breathing motions while preserving its overall fold. This pattern constitutes a hallmark of authentic protein–protein interactions involving cytoskeletal components, in which peripheral flexibility facilitates contact optimization without compromising structural integrity [76].

###### 3.3.3 Interface Persistence Is Confirmed by Distance and Contact Metrics

The persistence of the envelope–actin interface is unambiguously supported by intermolecular distance and contact analyses. The minimum interchain distance (MINdistance_PAIR_1) remains tightly and continuously distributed around ~0.162–0.165 nm throughout the entire trajectory, including the final 20 ns window used for energetic calculations. Values in this range correspond to persistent atomic-scale contacts of ~1.6 Å, which are incompatible with pre-dissociative states and unambiguously indicate maintained physical association at the interface.

The number of intermolecular contacts (NUMcontacts_PAIR_1) complements this picture, remaining elevated and stable at mean values of approximately 9.6–9.8 $\times$ $10^3$ over the simulation, with a slight increase observed in the final trajectory segment. This temporal trend, in which contact number either stabilizes or modestly increases rather than declining, provides direct evidence that the interface does not erode spontaneously on the simulation timescale. Individual contacts form and dissolve throughout the trajectory, as expected for a dynamically adaptable interface, but the interfacial core remains persistently assembled. This contact dynamics profile is characteristic of functional regulatory protein–protein interfaces, which are distinguished from crystallographic packing contacts precisely by the combination of high persistent contact density and localized contact turnover [41,47].

###### 3.3.4 Energetic Basis of Complex Stability: MM/GBSA Analysis

The thermodynamic basis of envelope–actin association was characterized through MM/GBSA binding free energy calculations. The energetic profile reveals a complex dominated by a strongly favorable electrostatic term (EEL), which is partially offset by a large and unfavorable polar solvation contribution (EGB). This pattern of substantial mutual compensation between EEL and EGB constitutes a classical signature of interfaces enriched in salt bridges and hydrogen bond networks, in which charge–charge interactions generate strong enthalpic gains that are partially counterbalanced by the energetic cost of displacing ordered hydration shells from polar groups [48]. Despite this compensation, the net total binding free energy remains negative, confirming that the interaction is globally thermodynamically favorable under the simulated conditions.

Van der Waals (VDW) interactions and nonpolar surface area burial terms (ESURF) contribute consistently favorable, albeit smaller in magnitude, energetic components. These terms reflect the hydrophobic packing and steric complementarity of the interfacial core, providing structural reinforcement that stabilizes the electrostatically driven binding event. Together, the energetic architecture suggests a binding mechanism in which electrostatic attraction drives initial recognition and orientational specificity, while van der Waals complementarity consolidates the assembled complex.

###### 3.3.5 Residue-Level Decomposition Identifies a Distributed and Energetically Redundant Interface

Per-residue energy decomposition (SDC/TDC) reveals that the total binding free energy is distributed across multiple residue clusters on both protein surfaces rather than concentrated in a single dominant contact. On the envelope protein, strongly favorable contributions arise from a cluster of residues in the C-terminal region, with basic residues — including arginine and lysine side chains — providing particularly prominent electrostatic contributions compatible with salt bridge formation. On the actin surface, complementary contributions are observed from acidic residue clusters and structurally flexible segments, which present electronegative environments geometrically accessible to the basic envelope residues. Both backbone and side-chain components contribute to the decomposed energies, indicating that the interface exploits both main-chain hydrogen bonding and side-chain electrostatic complementarity.

Certain residues display unfavorable decomposed energy values, reflecting the local desolvation penalties associated with burial of charged groups in regions of imperfect geometric fit. The presence of such energetically costly residues within a globally favorable interface is a recognized feature of genuine protein–protein complexes and does not undermine the overall stability of the association [43]. Rather, the combination of strongly favorable anchor residues and locally unfavorable desolvation contacts defines an interface that is both energetically robust and structurally specific — properties that distinguish functional binding surfaces from non-specific associations.

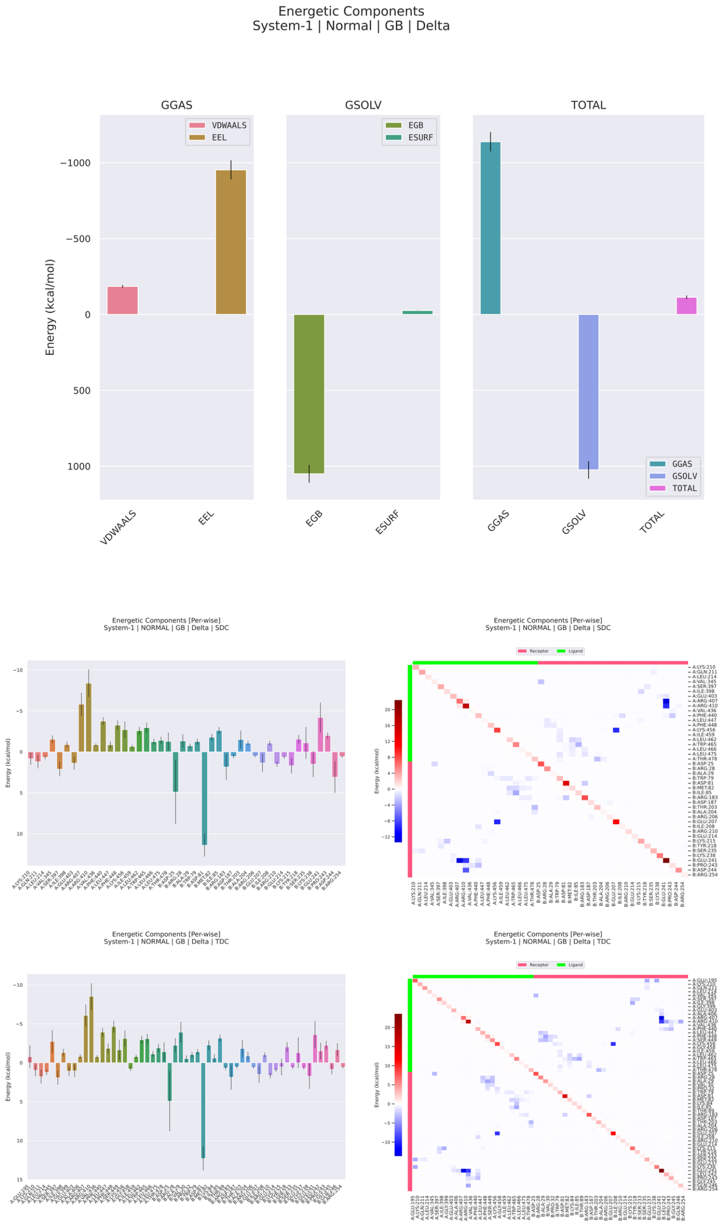


**Figure 5: MM/GBSA energetic decomposition of the dengue virus Envelope–actin complex during molecular dynamics simulation.** The figure summarizes the global energetic components, per-residue decomposition, and pairwise residue decomposition of the binding free energy estimated by MM/GBSA for the dengue Envelope–actin complex. The upper panels show that the gas-phase contribution is strongly favorable and dominated by the electrostatic term, whereas the solvation component is largely unfavorable due to the polar generalized Born contribution. Despite this compensation between electrostatic attraction and polar desolvation penalty, the total binding free energy remains favorable, supporting the energetic stability of the complex. The middle and lower panels show per-residue and pairwise decompositions using SDC and TDC schemes, revealing a heterogeneous interface in which a limited subset of residues and residue pairs concentrates the strongest stabilizing contributions. Negative values indicate favorable contributions to binding, whereas positive values indicate unfavorable or destabilizing energetic terms. In the pairwise heatmaps, blue regions denote favorable residue-pair interactions and red regions denote unfavorable contributions. Overall, the MM/GBSA profile indicates that the Envelope–actin interface is organized around discrete energetic hotspots, with binding largely driven by electrostatic interactions and partially opposed by polar solvation effects.

##### 3.3.6 Ternary Docking Reveals Neighbor-Dependent Modulation of the ENV–Actin Interface

To evaluate how actin-associated host proteins modulate the interaction landscape between the dengue virus envelope protein and the cytoskeletal environment, we systematically explored ternary docking configurations involving ENV, actin, and first-neighbor actin partners identified in protein interaction networks. For each candidate neighbor (ACTL6A, ACTN2, ACTR1A, ACTR10, CAPZB, and PFN1), docking energies were compared across pairwise complexes and two alternative ternary assemblies ("Triple" and "Triple_Inverse"), allowing estimation of energetic coupling through $\Delta\Delta G$ metrics derived from the relative stabilization or destabilization of the viral–cell interface.

Across all tested neighbors, pairwise ENV–actin docking energies remained within a relatively narrow energetic range (approximately $-11$ to $-13$ kcal $\cdot$ mol$^{-1}$), consistent with a stable but moderate protein–protein interaction. The introduction of a third host partner, however, altered the energetic landscape in a neighbor-dependent manner. Notably, ternary assembly with ACTL6A generated the strongest stabilizing effect when the assembly order was inverted (Viral–Cell Triple_Inverse), producing a $\Delta\Delta G$ of approximately $-1.50$ kcal $\cdot$ mol$^{-1}$ relative to the reference pairwise configuration. In contrast, most other actin neighbors produced either neutral or destabilizing effects when forming a Cell–Neighbor triple prior to viral engagement, with $\Delta\Delta G$ values ranging from $+0.66$ to $+6.65$ kcal $\cdot$ mol$^{-1}$ depending on the protein. These results indicate that while actin can form stable ternary complexes with several cytoskeletal regulators, only a subset of these neighbors enhances compatibility with the viral envelope interface.

Proteins directly involved in actin filament organization displayed a characteristic energetic pattern. ACTN2, PFN1, ACTR1A, and ACTR10 all showed substantial positive $\Delta\Delta G$ values in the Cell–Neighbor configuration ($\Delta\Delta G \approx +0.66$ to $+6.65$ kcal $\cdot$ mol$^{-1}$), suggesting that early association of these proteins with actin tends to reduce the stability of the ENV–actin interaction surface. When the assembly order was reversed (Viral–Cell Triple_Inverse), however, these same proteins produced small but reproducible negative $\Delta\Delta G$ shifts (approximately $-0.11$ to $-0.33$ kcal $\cdot$ mol$^{-1}$), indicating that once the ENV–actin interface is established, their presence may slightly stabilize the ternary complex.

CAPZB displayed a distinct behavior compared with other actin regulators. While the Cell–Neighbor ternary assembly with actin was energetically favorable ($\Delta\Delta G \approx -1.24$ kcal $\cdot$ mol$^{-1}$), the addition of the viral envelope produced only negligible energetic change ($\Delta\Delta G \approx -0.03$ kcal $\cdot$ mol$^{-1}$). This suggests that CAPZB stabilizes actin structure without strongly influencing the compatibility of the ENV–actin interface itself.

The GO enrichment analysis performed on the neighbor set highlighted ACTL6A as the only protein significantly associated with enriched functional annotations, mapping to processes involved in the regulation of organelle organization. This observation distinguishes ACTL6A from the other neighbors, which are predominantly classical cytoskeletal modulators.

Taken together, the ternary docking framework reveals that actin-associated proteins exert heterogeneous effects on the ENV–actin interface. While several neighbors appear to compete for actin binding determinants during early complex formation, a smaller subset — particularly ACTL6A — exhibits cooperative stabilization when the viral interface is pre-established.

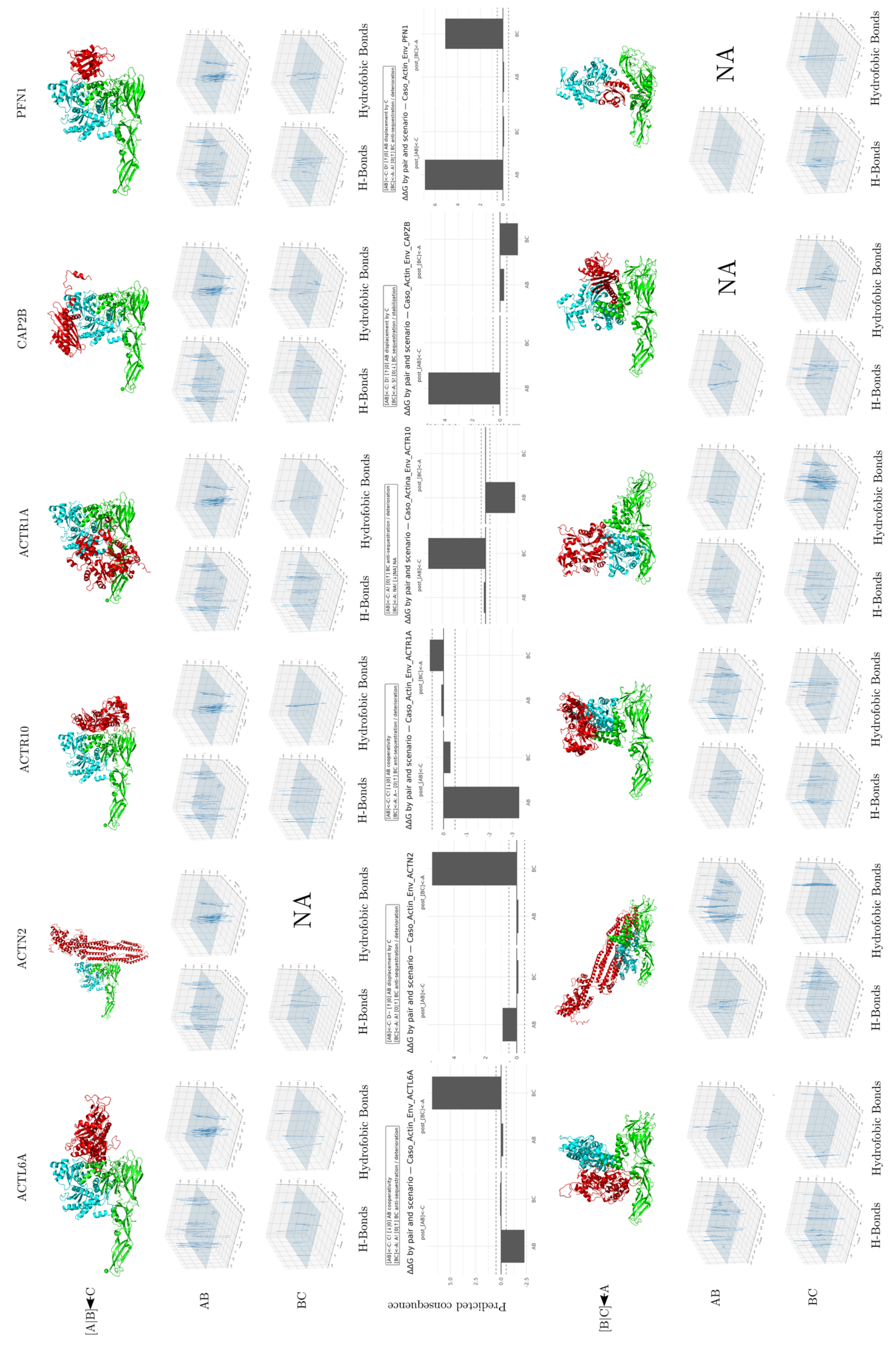
ACTL6A
ACTN2
ACTR10
ACTR1A
CAP2B
PFN1
[A|B]←C
AB
BC
NA
H-Bonds
Hydrofobic Bonds
Predicted consequence
ΔΔG by pair and scenario — Caso_Actin_Env_ACTL6A
ΔΔG by pair and scenario — Caso_Actin_Env_ACTN2
ΔΔG by pair and scenario — Caso_Actin_Env_ACTR1A
ΔΔG by pair and scenario — Caso_Actina_Env_ACTR10
ΔΔG by pair and scenario — Caso_Actin_Env_CAPZB
ΔΔG by pair and scenario — Caso_Actin_Env_PFN1
[B|C]←A

**Figure 6. Structural, interaction-field, and predicted energetic consequences of dengue Envelope–actin-centered ternary complexes with actin-associated proteins.**

The figure summarizes comparative ternary interaction models involving dengue virus Envelope, actin, and six actin-associated proteins: ACTL6A, ACTN2, ACTR10, ACTR1A, CAP2B, and PFN1. Two alternative assembly scenarios are shown. In the first, the Envelope–actin pair is treated as the preformed binary complex and the third protein is introduced as an additional partner. In the reciprocal scenario, the actin-associated binary pair is formed first and Envelope is introduced afterward. Structural panels show the relative spatial organization of the three proteins in each modeled ternary assembly. Interaction-field plots represent the spatial distribution of hydrogen-bond and hydrophobic contact features for the AB and BC interfaces, highlighting localized regions of polar and nonpolar interaction density.

Predicted energetic consequence plots report $\Delta\Delta G$ values for each pair and assembly scenario, with negative values indicating stabilizing effects and positive values indicating destabilizing or energetically unfavorable effects relative to the corresponding reference state. Missing panels are indicated as NA when a given configuration or interaction descriptor was not available. Overall, the figure shows that the Envelope–actin interaction is strongly modulated by the identity of the actin-associated partner and by the order of ternary complex formation, supporting a model in which dengue Envelope engages actin within a context-dependent and network-embedded host interface.

#### 3.4 Envelope–Betaglycan/TGFBR3 Complex: Structural Maturation and Interfacial Densification

##### 3.4.1 Progressive Compaction and Monotonic Growth of Interface Contacts

The envelope–betaglycan complex undergoes a well-defined trajectory of interfacial maturation over the 200 ns simulation, transitioning from an initially loose association to a progressively compacted and energetically narrower conformational basin. The mean interchain distance declines from approximately 3.37 nm in the first 20 ns to ~3.12 nm in the final 20 ns, accompanied by a substantial reduction in distributional variability in the late-trajectory window (standard deviation ~0.022 nm at 180–200 ns), indicative of increasing conformational confinement as the system approaches a dominant interfacial state. In parallel, the number of intermolecular contacts increases monotonically from ~$8.3 \times 10^3$ in the earliest trajectory segment to ~$1.49 \times 10^4$ at 180–200 ns, a near-doubling that is inconsistent with passive thermal fluctuation and instead reflects active interfacial rearrangement toward greater geometric complementarity and increased effective contact area. The minimum interchain distance metric, which captures the shortest atomic contact within the interface, remains persistently low at approximately 0.16 nm throughout the entire trajectory with only marginal variation, confirming continuous atomic-scale proximity and excluding transient dissociation events even during early stages when the mean separation remained higher.

##### 3.4.2 Differential Conformational Adjustment Between Receptor and Ligand

Global structural convergence of both partners is confirmed by RMSD profiles, which display initial increases followed by well-defined plateaus after approximately 100 ns. The envelope protein reaches a mean RMSD of ~0.58 nm in the 100–200 ns window compared to ~0.45 nm in the first 20 ns, indicating moderate surface remodeling consistent with interfacial accommodation. Betaglycan displays a larger conformational shift, converging to a mean RMSD of ~0.77 nm at 100–200 ns from ~0.56 nm initially, reflecting a substantially greater conformational cost of adaptation in the host partner relative to the viral receptor. This asymmetry is reinforced by residue-level RMSF analysis: betaglycan presents markedly elevated per-residue fluctuations, with pronounced peaks exceeding 1.0 nm localized to terminal regions and structurally mobile loops, whereas the envelope protein maintains a comparatively restrained RMSF profile with maximum values of approximately 0.54 nm. The higher conformational cost on the betaglycan side is mechanistically consistent with the known modular architecture of the TGFBR3 ectodomain, which contains structurally specialized extracellular segments — including a zona pellucida-like module — that accommodate diverse ligands through localized conformational plasticity [66,67].

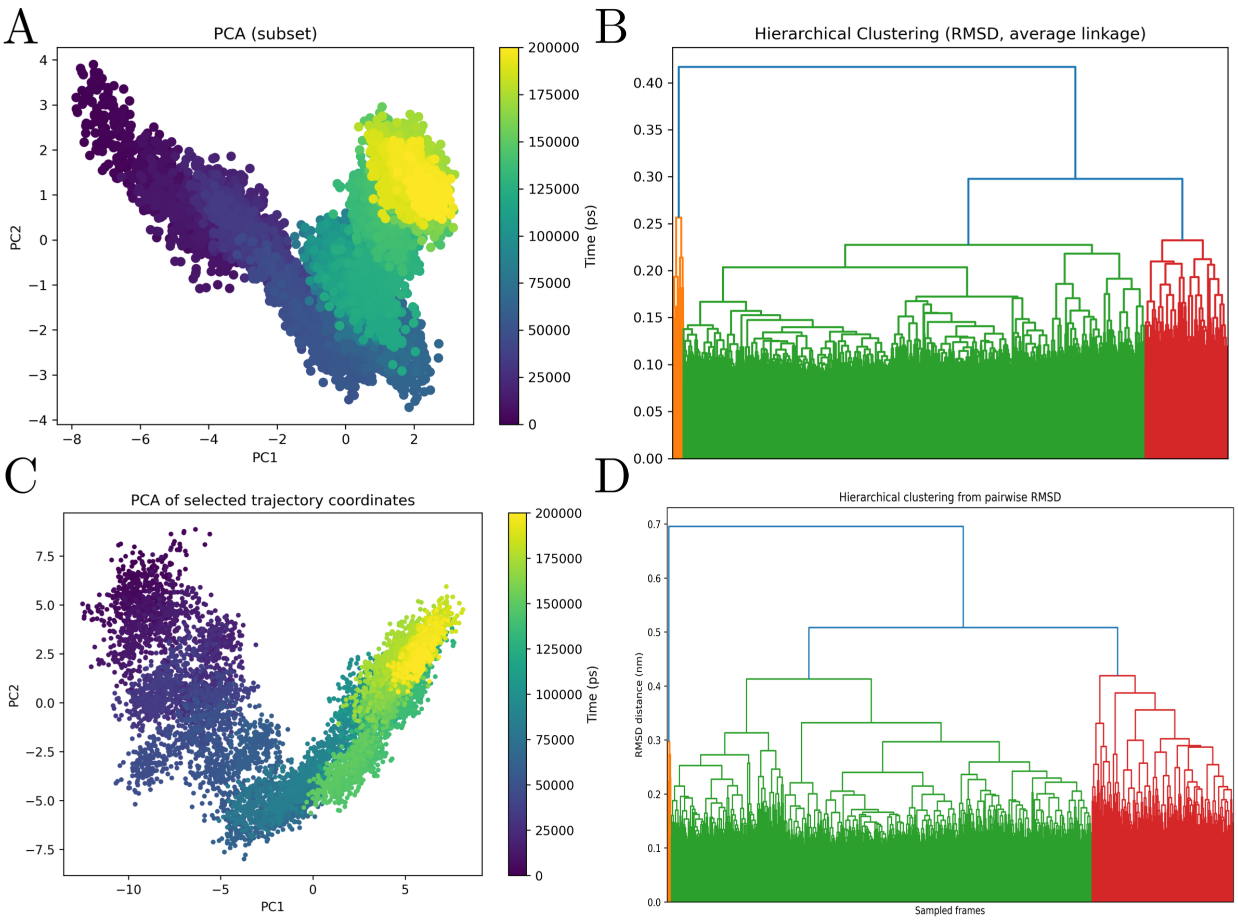


**Figure 7: Principal component analysis and RMSD-based hierarchical clustering of the dengue Envelope–BiP molecular dynamics trajectory.** The figure summarizes the conformational landscape of the dengue Envelope–BiP complex, with AB corresponding to dengue Envelope and CD corresponding to BiP (HSPA5/GRP78). Panels A and C show PCA projections of trajectory frames onto the first two principal components, with frames colored according to simulation time. Panel A shows the Envelope-associated subset, revealing a temporally ordered and continuous conformational transition across the trajectory, consistent with progressive structural accommodation rather than confinement to a single rigid state. Panel C shows the BiP-associated subset, which occupies a broader region of principal component space and displays increased conformational dispersion, indicating substantial structural flexibility and sampling of multiple related substates. Panels B and D show RMSD-based hierarchical clustering using average linkage. The Envelope-associated subset forms a relatively coherent clustering pattern, whereas the BiP-associated subset is partitioned into several more distinct conformational families, consistent with greater structural heterogeneity. Together, the PCA and clustering analyses indicate that the Envelope–BiP complex is dynamically heterogeneous and organized as an ensemble of conformational states, with Envelope showing progressive conformational drift and BiP contributing marked adaptive flexibility to the interaction.

##### 3.4.3 Interfacial Distance Distribution Reflects Short-Range Contact Dominance

Analysis of the interfacial distance distribution reveals that the majority of contact area contributions are concentrated at the shortest distance classes, with a sharp peak near ~0.1 nm and a rapidly decaying tail extending to ~2.4 nm. This distribution is consistent with a contact regime dominated by van der Waals interactions and hydrogen bond or salt bridge networks at near-covalent range, supplemented by a smaller fraction of medium-range electrostatic contributions. The progressive narrowing of this distribution in combination with increased contact numbers and decreased mean distance provides convergent evidence that the interface undergoes genuine densification during the simulation rather than mere thermal redistribution of pre-existing contacts.

###### 3.4.4 MM/GBSA Energetics Reveal Electrostatically Driven Binding with Persistent Polar Frustration

The MM/GBSA binding free energy profile of the envelope–betaglycan complex is characterized by substantial gas-phase stabilization almost entirely offset by polar desolvation penalties, yielding a modestly negative net binding free energy. The gas-phase term ($\Delta G_{\text{gas}}$) is strongly favorable, driven predominantly by a large negative electrostatic contribution (EEL, approximately $-1.05 \times 10^3$ kcal/mol) supplemented by van der Waals interactions (VDWAALS, approximately $-3 \times 10^2$ kcal/mol). The solvation term ($\Delta G_{\text{solv}}$) is correspondingly large and unfavorable, almost entirely governed by the polar component (EGB, approximately $+1.30 \times 10^3$ kcal/mol), while the nonpolar solvation contribution (ESURF, approximately $-4 \times 10^1$ kcal/mol) provides only marginal stabilization. The resulting net $\Delta G_{\text{total}}$ remains negative at approximately $-1 \times 10^2$ kcal/mol, confirming thermodynamic viability but reflecting a binding regime governed more by the mutual attenuation of large competing terms than by the dominance of a single favorable energetic determinant. This energetic profile constitutes a classical signature of protein–protein interfaces enriched in ionizable residues, in which gas-phase electrostatics substantially overestimate favorable interaction and solvation corrections significantly attenuate the apparent binding strength [49,77].

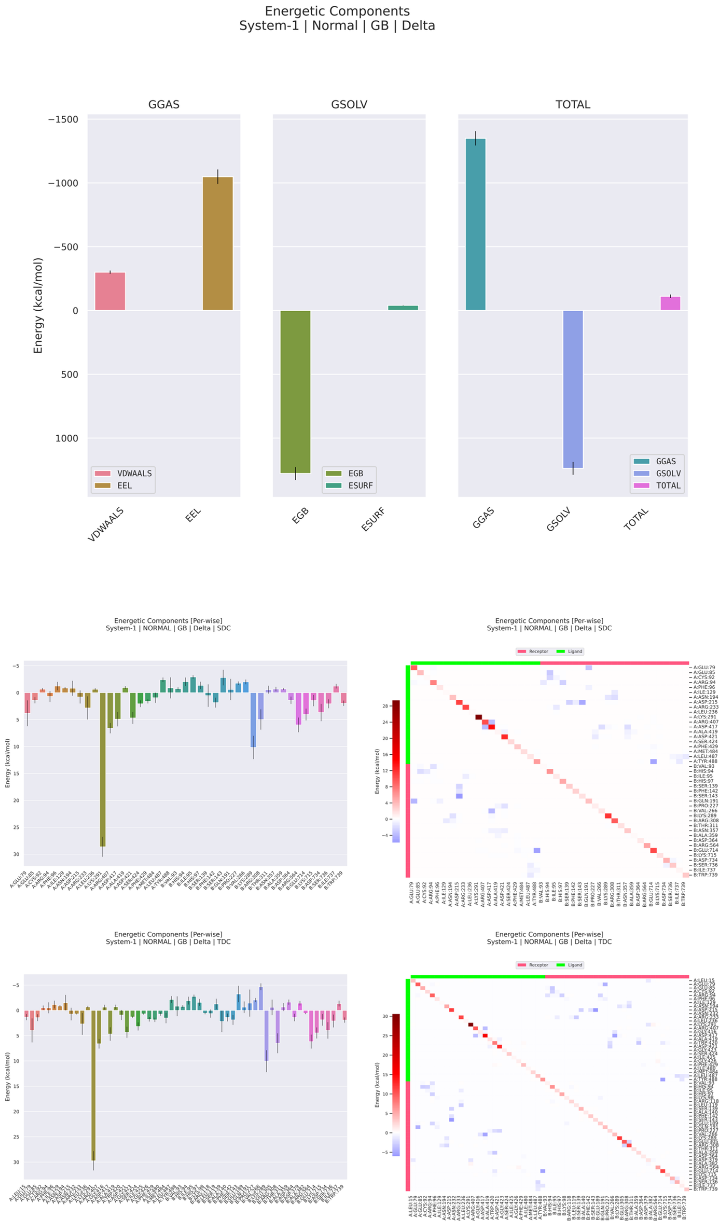


**Figure 8: MM/GBSA energetic decomposition of the dengue virus Envelope–betaglycan complex during molecular dynamics simulation.** The figure summarizes the global energetic components, per-residue energetic decomposition, and pairwise residue decomposition of the binding free energy estimated by MM/GBSA for the dengue Envelope–betaglycan complex. The upper panels show the decomposition of the gas-phase, solvation, and total energetic terms. The gas-phase contribution is strongly favorable and is dominated by the electrostatic component, whereas the solvation term is largely unfavorable due to the polar generalized Born contribution. The nonpolar solvation term contributes only modestly relative to the dominant polar solvation penalty. Despite this compensation, the total binding free energy remains favorable, supporting the energetic plausibility of the Envelope–betaglycan interaction. The residue-wise decomposition panels show that the interface is energetically heterogeneous, with a limited number of residues contributing strongly favorable binding terms and other positions showing neutral or unfavorable effects. The pairwise decomposition heatmaps reveal discrete residue-pair energetic couplings, with favorable interactions shown in blue and unfavorable interactions shown in red. Together, these results indicate that the Envelope–betaglycan interface is primarily electrostatic, strongly affected by polar desolvation, and organized around localized energetic hotspots rather than a uniformly distributed binding surface.

##### 3.4.5 Residue-Level Decomposition Identifies Electrostatic Frustration Hotspots on the Envelope

Per-residue free energy decomposition reveals an energetically heterogeneous interface organized around a small number of unfavorable outliers on the viral envelope rather than around dominant stabilizing hotspots. On the envelope surface, two basic residues — ARG107 and LYS236 — emerge as strongly penalizing contributors in both side-chain decomposition (SDC) and total decomposition (TDC) analyses, each imposing positive free-energy costs of approximately +28 to +30 kcal/mol that counteract favorable contributions elsewhere in the interface. On the betaglycan surface, the energetic landscape is more diffuse, with charged and polar residues including glutamate, lysine, arginine, and serine/threonine-containing regions exhibiting alternating favorable and unfavorable contributions. Heatmap representations confirm that the majority of interfacial residue pairs contribute near-zero energies, with sparse islands of opposing sign, consistent with an interface sustained by multiple low- to medium-persistence electrostatic microcontacts in the absence of a cohesive hydrophobic core [78].

##### 3.4.6 Ternary Docking Reveals Conditional Energetic Antagonism and Selective Cooperative Stabilisation Around the Betaglycan–Env Interface

To evaluate how the local Betaglycan (TGFBR3) interaction network modulates the dengue envelope–Betaglycan interface, we performed ternary docking simulations incorporating seven first-neighbour host proteins identified from curated interaction databases: TGFBR2, TGFB3, KLK22, IGKV2D-28, IGHV3, FGF1, and CANT1. For each candidate, docking energies were compared across the pairwise Betaglycan–Env reference complex (AB), the pairwise Env–neighbour complex (BC), and two conditional ternary assemblies — the AB-conditioned configuration (third partner docking onto a pre-formed Betaglycan–Env interface) and the reverse-conditioned configuration (Env docking onto a pre-formed Betaglycan–neighbour complex). Cooperative coupling was quantified through $\Delta\Delta G$ values reflecting the energetic shift in each binary interface upon ternary assembly [41,51].

Across all seven third partners, the Betaglycan–Env reference pairwise energy (AB) was consistently stable, clustering near $\Delta G \approx -13.8$ to $-13.9$ kcal $\cdot$ mol$^{-1}$. In contrast, the third-partner pairwise energies (BC) displayed marked heterogeneity, ranging from approximately $-8.0$ kcal $\cdot$ mol$^{-1}$ for KLK22 to approximately $-13.9$ kcal $\cdot$ mol$^{-1}$ for TGFBR2. This pattern — a conserved, stable AB baseline coexisting with variable BC energetics — is consistent with a hierarchically organised interaction surface in which the Betaglycan–Env contact represents a preferred energetic state whose compatibility with third partners depends on the structural and electrostatic properties of each neighbour.

The most pronounced antagonistic effects were observed for KLK22 and TGFB3. KLK22 produced the strongest destabilisation in the dataset: in the AB-conditioned configuration, BC shifted from $-8.0$ to $-7.0$ $\text{kcal}\cdot\text{mol}^{-1}$ ($\Delta\Delta G \approx +1.0$ $\text{kcal}\cdot\text{mol}^{-1}$), while under reverse conditioning, the Betaglycan–Env interface collapsed from $-13.9$ to $-8.8$ $\text{kcal}\cdot\text{mol}^{-1}$ ($\Delta\Delta G \approx +5.1$ $\text{kcal}\cdot\text{mol}^{-1}$) — the largest single-configuration destabilisation in the entire panel. TGFB3 showed a distinct but also strongly disruptive profile: AB was weakened from $-13.8$ to $-11.1$ $\text{kcal}\cdot\text{mol}^{-1}$ under AB conditioning ($\Delta\Delta G \approx +2.7$ $\text{kcal}\cdot\text{mol}^{-1}$), while the reverse-conditioned branch produced only a negligible AB shift ($\Delta\Delta G \approx -0.1$ $\text{kcal}\cdot\text{mol}^{-1}$). The asymmetry between conditioning branches for TGFB3 indicates a directional effect operating preferentially when the Betaglycan–Env state is already established.

Cooperative stabilisation was observed for both immunoglobulin-variable-domain partners. IGKV2D-28 produced the cleanest stabilising signature: under AB conditioning, AB shifted from $-13.8$ to $-15.3$ $\text{kcal}\cdot\text{mol}^{-1}$ ($\Delta\Delta G \approx -1.5$ $\text{kcal}\cdot\text{mol}^{-1}$) while BC remained essentially unchanged, indicating selective reinforcement of the Betaglycan–Env interface without perturbation of the Env–neighbour contact. IGHV3 exhibited a mixed but energetically informative profile: under AB conditioning, BC was strongly destabilised from $-12.1$ to $-6.7$ $\text{kcal}\cdot\text{mol}^{-1}$ ($\Delta\Delta G \approx +5.4$ $\text{kcal}\cdot\text{mol}^{-1}$), while under reverse conditioning, AB was modestly stabilised from $-13.8$ to $-14.6$ $\text{kcal}\cdot\text{mol}^{-1}$ ($\Delta\Delta G \approx -0.8$ $\text{kcal}\cdot\text{mol}^{-1}$). This pattern is consistent with IGHV3 simultaneously sequestering the Env surface from competing neighbour contacts while consolidating the Betaglycan–Env interface.

The three remaining partners clustered near energetic neutrality or produced moderate interface-selective effects. TGFBR2 was the most conservative: AB-conditioning shifted AB by $\Delta\Delta G \approx -0.2$ $\text{kcal}\cdot\text{mol}^{-1}$ and BC by $+0.1$ $\text{kcal}\cdot\text{mol}^{-1}$, and reverse conditioning produced only $\Delta\Delta G \approx -0.1$ $\text{kcal}\cdot\text{mol}^{-1}$ in AB — all values consistent with TGFBR2 being energetically permissive with respect to the Betaglycan–Env state. FGF1 produced a moderate, interface-selective effect: AB was unchanged under AB conditioning while BC was weakened from $-11.1$ to $-9.2$ $\text{kcal}\cdot\text{mol}^{-1}$ ($\Delta\Delta G \approx +1.9$ $\text{kcal}\cdot\text{mol}^{-1}$), with a negligible AB penalty of $\Delta\Delta G \approx +0.1$ $\text{kcal}\cdot\text{mol}^{-1}$ under reverse conditioning. CANT1 was the most energetically inert partner examined, with maximum $|\Delta\Delta G|$ values of approximately 0.1–0.2 $\text{kcal}\cdot\text{mol}^{-1}$ across all configurations.

The overall energetic ranking is therefore unambiguous: KLK22 and TGFB3 are antagonists of the Betaglycan–Env state through distinct directional mechanisms; IGKV2D-28 and IGHV3 exert cooperative stabilising effects with differing specificity for AB versus BC interfaces; FGF1 selectively disfavours BC engagement without substantially disrupting AB; and TGFBR2 and CANT1 are effectively permissive. These results reveal that the Betaglycan–Env interface is embedded within a host-side conditional landscape whose energetic topology depends on the molecular identity of neighbouring interaction partners.

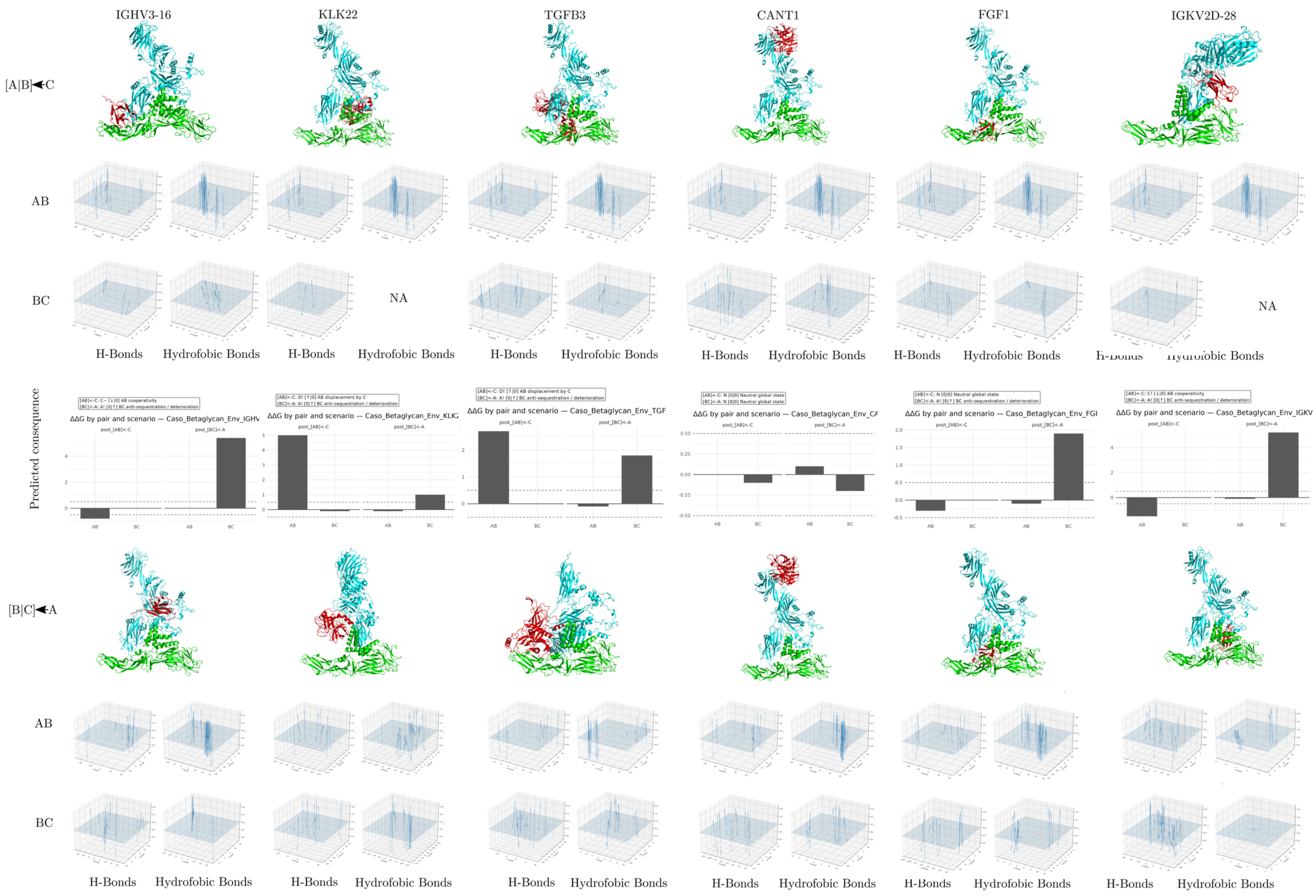

IGHV3-16
KLK22
TGFB3
CANT1
FGF1
IGKV2D-28
[A|B]◄C
AB
BC
NA
NA
H-Bonds
Hydrofobic Bonds
Predicted consequence
ΔΔG by pair and scenario — Caso_Betaglycan_Env_IGHV
ΔΔG by pair and scenario — Caso_Betaglycan_Env_KLK2
ΔΔG by pair and scenario — Caso_Betaglycan_Env_TGF
ΔΔG by pair and scenario — Caso_Betaglycan_Env_CA
ΔΔG by pair and scenario — Caso_Betaglycan_Env_FGI
ΔΔG by pair and scenario — Caso_Betaglycan_Env_IGKV
[B|C]◄A

**Figure 9. Structural, interaction-field, and predicted energetic consequences of dengue Envelope–betaglycan-centered ternary complexes with betaglycan-associated partners.**

The figure compares ternary models involving dengue virus Envelope, betaglycan/TGFBR3, and three betaglycan-associated proteins: IGHV3-16, KLK22, and TGFB3. Two alternative assembly scenarios are shown. In the first scenario, the Envelope–betaglycan pair is treated as the preformed binary complex and the third partner is introduced afterward. In the reciprocal scenario, the betaglycan-associated binary pair is formed first and the Envelope protein is subsequently introduced. Structural panels show the relative spatial organization of the three proteins in each ternary assembly. Interaction-field plots summarize the spatial distribution of hydrogen-bond and hydrophobic contact features across the AB and BC interfaces, highlighting localized polar and nonpolar interaction densities. Predicted energetic consequence plots report ΔΔG values for each pair and assembly route, where negative values indicate stabilizing effects and positive values indicate destabilizing or energetically unfavorable effects relative to the corresponding reference state. Missing panels are marked as NA when a given interaction descriptor was unavailable. Overall, the figure indicates that dengue Envelope engagement with betaglycan is strongly modulated by the identity of the associated partner and by the order of ternary complex formation, supporting a model in which the Envelope–betaglycan interface is embedded within a context-dependent extracellular interaction network.

#### 3.5 Envelope–BiP/GRP78 Complex: Chaperone-Mediated Stabilization and Electrostatic Anchoring

##### 3.5.1 Global Structural Stability Under Chaperone Engagement

The envelope–BiP/GRP78 complex exhibits sustained global structural stability throughout the molecular dynamics trajectory, as demonstrated by the convergence and plateau of backbone RMSD values for both interacting partners. The envelope protein rapidly attains a conformational steady state following the equilibration phase, with RMSD values remaining dynamically constrained thereafter. BiP displays moderately higher RMSD amplitudes, consistent with its established conformational heterogeneity and the large-scale interdomain rearrangements that are intrinsic to its ATP-dependent chaperone cycle [69,79]. Critically, no long-term RMSD drift is observed in either partner, indicating that complex formation does not induce progressive structural destabilization or unfolding of the viral protein.

Analysis of the radius of gyration (Rg) corroborates this interpretation. The envelope protein maintains a compact and stable fold with minimal Rg fluctuations throughout the trajectory, while BiP undergoes modest and reversible oscillations in Rg, reflecting breathing motions rather than large-scale compaction or expansion events. This pattern is characteristic of chaperone–client interactions, in which substrate-binding-induced structural adaptability enables persistent engagement without compromising the global fold integrity of either partner [80].

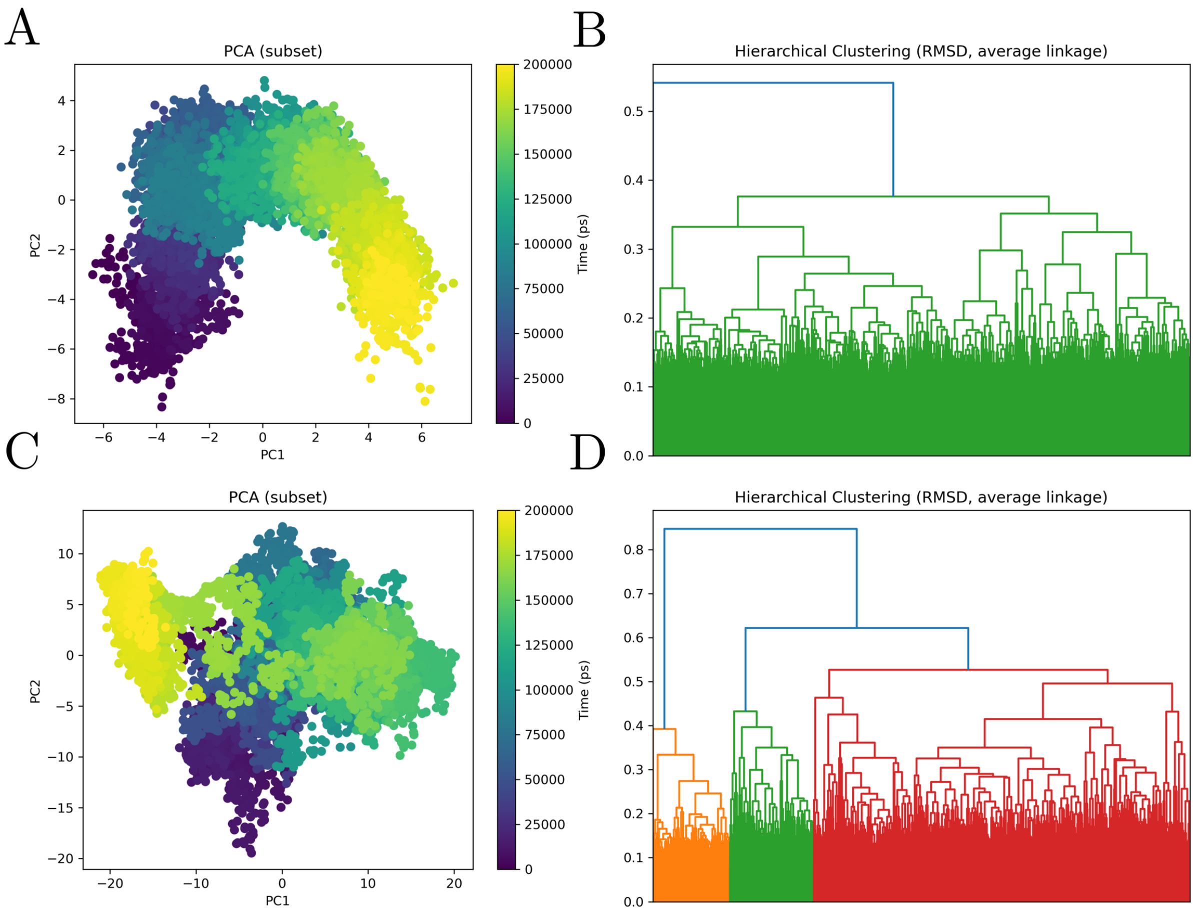


**Figure 10: Principal component analysis and RMSD-based hierarchical clustering of the dengue Envelope–BiP molecular dynamics trajectory.** The figure summarizes the conformational landscape of the dengue Envelope–BiP complex, with AB corresponding to dengue Envelope and CD corresponding to BiP (HSPA5/GRP78). Panels A and C show PCA projections of trajectory frames onto the first two principal components, with frames colored according to simulation time. Panel A shows the Envelope-associated subset, revealing a temporally ordered and largely continuous conformational transition across the trajectory, consistent with progressive structural accommodation within the complex. Panel C shows the BiP-associated subset, which occupies a broader region of principal component space and exhibits greater conformational dispersion, indicating substantial structural plasticity and sampling of multiple related substates. Panels B and D show RMSD-based hierarchical clustering using average linkage. The Envelope-associated subset forms a relatively coherent clustering pattern with limited high-level separation, whereas the BiP-associated subset is partitioned into several more distinct conformational families, consistent with increased structural heterogeneity. Together, the PCA and clustering analyses indicate that the Envelope–BiP complex is organized as a dynamic conformational ensemble, in which Envelope undergoes progressive conformational drift while BiP contributes marked adaptive flexibility to the interaction.

##### 3.5.2 Asymmetric Flexibility Redistribution Reveals Selective Interface Stabilization

Residue-level RMSF analysis reveals a pronounced asymmetry in the redistribution of conformational flexibility upon complex formation. In BiP, solvent-exposed segments — particularly those associated with interdomain linker regions and substrate-engagement surfaces — display reduced per-residue fluctuations, suggesting that Envelope binding stabilizes these regions through reciprocal interfacial contacts. In parallel, peripheral loops of BiP retain flexibility, consistent with the dynamic nature of chaperone–client recognition, in which partial surface immobilization coexists with continued local adaptability to accommodate transient contact rearrangements [81].

The envelope protein exhibits a complementary but spatially distinct pattern: RMSF reductions are localized to discrete surface patches that coincide with the predicted interaction interface, while the remainder of the protein retains its characteristic conformational dynamics. This selective dampening of fluctuations at the viral interaction surface — without global propagation across the protein — is a recurrent feature of biologically relevant virus–host interfaces, where precision surface immobilization supports affinity and specificity while preserving the conformational landscape required for downstream functional transitions [82].

###### 3.5.3 Persistent Intermolecular Contacts Confirm Continuous Interface Engagement

Time-resolved minimum interchain distance and center-of-mass separation analyses demonstrate sustained close proximity between BiP and the envelope protein following the initial equilibration phase. The minimum distance profile stabilizes at values compatible with persistent non-covalent interactions, and the overall distance distribution lacks episodic separation events, confirming that the complex remains continuously engaged throughout the production trajectory without evidence of transient dissociation.

Contact number analysis reinforces this conclusion, revealing a stable and elevated network of intermolecular contacts with only moderate temporal fluctuations. Importantly, these fluctuations are oscillatory rather than monotonically declining, indicating continuous residue-level reshuffling of contacts rather than progressive interface erosion. This dynamic contact behavior is a hallmark of protein–protein interfaces that combine enthalpic stability with local entropic adaptability, distinguishing functional binding events from transient non-specific encounters [47,83].

###### 3.5.4 MM/GBSA Energetics Reveal a Composite Interface of Electrostatic Anchoring and Hydrophobic Packing

MM/GBSA binding free energy decomposition of the envelope–BiP complex reveals a net favorable interaction of moderate magnitude (TOTAL $\approx -70.5$ kcal/mol) arising from substantial compensation between strongly favorable gas-phase interactions and a large unfavorable polar solvation penalty. The gas-phase contribution ($\Delta G_{\text{gas}} \approx -750.8$ kcal/mol) is dominated by a large electrostatic term (EEL $\approx -568.4$ kcal/mol), supplemented by a consistent van der Waals contribution (VDWAALS $\approx -182.4$ kcal/mol). The solvation term ($\Delta G_{\text{solv}} \approx +680.3$ kcal/mol) is strongly unfavorable and almost entirely governed by the polar component (EGB $\approx +705.2$ kcal/mol), while the nonpolar solvation term (ESURF $\approx -24.9$ kcal/mol) provides only marginal stabilization. This energetic architecture — large and mutually compensating electrostatic and desolvation terms yielding a modest but consistently negative total — constitutes the canonical MM/GBSA signature of protein–protein interfaces enriched in charged and polar residues, wherein transfer of ionizable groups from solvent into the interface generates strong attractive interactions in vacuo at the cost of reorganizing ordered hydration shells [49,77].

###### 3.5.5 Residue-Level Decomposition Identifies Composite Hotspots and Electrostatic Frustration Sites

Per-residue free energy decomposition (SDC and TDC) reveals a heterogeneous interfacial energetic landscape organized around a small subset of strongly contributing residues on each protein. On the envelope protein, A:ARG410 emerges as the dominant stabilizing residue (contributing approximately −3.10 to −3.26 kcal/mol), consistent with salt bridge or hydrogen bond formation. A:LEU487 provides an additional favorable contribution (≈ −2.80 kcal/mol), reflecting van der Waals–driven packing. On the BiP surface, B:GLU498 (≈ −3.08 to −3.37 kcal/mol) pairs with the envelope arginine in a pattern consistent with electrostatic anchoring, while B:VAL6 (≈ −2.27 to −2.90 kcal/mol) and B:ILE450 (≈ −2.24 to −2.32 kcal/mol) contribute hydrophobic stabilization. Together, these hotspot residues define a composite interface in which electrostatic anchoring and van der Waals packing operate cooperatively, mutually reinforcing contact stability.

Conversely, a cluster of charged residues on the BiP surface imposes unfavorable energetic contributions, most prominently B:LYS435 (≈ +6.84 to +7.18 kcal/mol), alongside B:LYS516 (≈ +2.60 to +2.75 kcal/mol) and B:LYS447 (≈ +2.46 to +2.52 kcal/mol). On the envelope side, A:GLN211 (≈ +3.23 to +4.93 kcal/mol) and A:LYS210 (≈ +3.44 to +3.87 kcal/mol) contribute additional energetic penalties. These unfavorable terms signal regions of electrostatic frustration where polar or charged groups incur desolvation costs that are incompletely compensated by intermolecular contacts, generating localized energetic tensions within an otherwise stable complex [78,84].

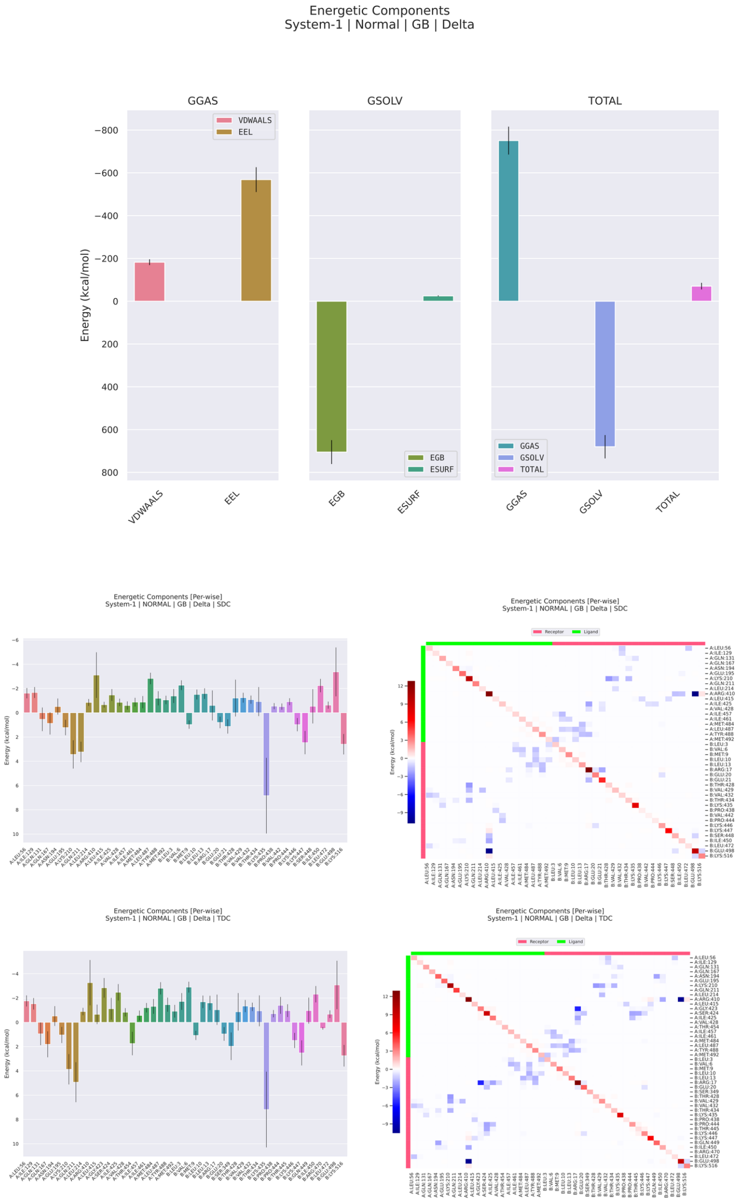


**Figure 11: MM/GBSA energetic decomposition of the dengue virus Envelope–BiP complex during molecular dynamics simulation.** The figure summarizes the global energetic components, per-residue energetic decomposition, and pairwise residue decomposition of the binding free energy estimated by MM/GBSA for the dengue Envelope–BiP complex. The upper panels show the decomposition of gas-phase, solvation, and total energetic terms. The gas-phase contribution is strongly favorable and dominated by the electrostatic component, whereas the solvation term is largely unfavorable due to the polar generalized Born contribution. The nonpolar solvation term contributes only modestly relative to the dominant polar desolvation penalty. Despite this energetic compensation, the total binding free energy remains favorable, supporting the energetic plausibility of the Envelope–BiP interaction. The residue-wise decomposition panels show that the interface is energetically heterogeneous, with a limited subset of residues contributing strongly favorable binding terms and other positions showing neutral or unfavorable effects. The pairwise decomposition heatmaps reveal discrete residue-pair energetic couplings, with favorable interactions shown in blue and unfavorable interactions shown in red. Together, these results indicate that the Envelope–BiP interface is primarily electrostatic, strongly affected by polar desolvation, and organized around localized energetic hotspots rather than a uniformly distributed binding surface.

##### 3.5.6 Ternary Docking Reveals ER-Context-Dependent Modulation of the BiP–Env Interface

To evaluate how the ER proteostasis neighbourhood of BiP/GRP78 modulates the dengue envelope–BiP interface, we performed conditional ternary docking simulations incorporating seven first-neighbour ER-resident proteins identified from curated interaction networks: DNAJB11, FICD, DERL1, HSP90B1, HYOU1, PDIA6, and P4HB. For each candidate, docking energies were compared across the BiP–Env reference pair (AB), the Env–neighbour pair (BC), and two conditional ternary assemblies — the AB-conditioned configuration (third partner docking onto the pre-formed BiP–Env interface) and the reverse-conditioned configuration (Env docking onto a pre-formed BiP–neighbour assembly). Energetic coupling was quantified through $\Delta\Delta G$ values derived from the differential stabilisation or destabilisation of each binary interface upon ternary assembly [41,51].

Across all seven third partners, the BiP–Env reference pairwise energy (AB) was remarkably uniform, clustering near $\Delta G \approx -14.2$ kcal · mol$^{-1}$, while third-partner pairwise energies (BC) displayed broader heterogeneity, ranging from approximately $-9.6$ kcal · mol$^{-1}$ for HYOU1 to $-15.2$ kcal · mol$^{-1}$ for HSP90B1. This pattern — a stable and invariant AB baseline alongside variable conditional perturbations — indicates that the BiP–Env interface constitutes a robust energetic state whose compatibility with the ER proteostasis network depends on the identity of the third component rather than on intrinsic instability of the reference complex.

The most pronounced antagonistic effects were observed for HYOU1, DERL1, and HSP90B1. HYOU1 produced the strongest destabilisation in the entire dataset: under AB conditioning, AB shifted from $-14.2$ to $-4.8$ kcal · mol$^{-1}$ ($\Delta\Delta G \approx +9.4$ kcal · mol$^{-1}$), while BC changed only negligibly ($\Delta\Delta G \approx -0.2$ kcal · mol$^{-1}$), and the reverse-conditioned branch produced only a trivial AB perturbation ($\Delta\Delta G \approx +0.2$ kcal · mol$^{-1}$). DERL1 showed a comparable directional signature: AB collapsed from $-14.2$ to $-6.7$ kcal · mol$^{-1}$ under AB conditioning ($\Delta\Delta G \approx +7.5$ kcal · mol$^{-1}$), with minimal BC change ($\Delta\Delta G \approx -0.3$ kcal · mol$^{-1}$) and only a $+0.2$ kcal · mol$^{-1}$ perturbation in the reverse branch. HSP90B1 produced a similarly directional, though somewhat smaller, effect: AB weakened from $-14.2$ to $-9.0$ kcal · mol$^{-1}$ ($\Delta\Delta G \approx +5.2$ kcal · mol$^{-1}$) with BC essentially unchanged ($\Delta\Delta G \approx -0.1$ kcal · mol$^{-1}$).

PDIA6 belongs to the same antagonistic class, albeit with a slightly reduced magnitude. Under AB conditioning, AB shifted from $-14.2$ to $-7.3$ kcal · mol$^{-1}$ ($\Delta\Delta G \approx +6.9$ kcal · mol$^{-1}$), with only a marginal BC stabilisation from $-14.6$ to $-14.8$ kcal · mol$^{-1}$ ($\Delta\Delta G \approx -0.2$ kcal · mol$^{-1}$). The reverse-conditioned branch again produced only a small AB penalty ($\Delta\Delta G \approx +0.2$ kcal · mol$^{-1}$). Across all four antagonists — HYOU1, DERL1, HSP90B1, and PDIA6 — the directional asymmetry between conditioning branches is a consistent feature: the AB-conditioned configuration strongly destabilises the BiP–Env state, whereas the reverse-conditioned configuration leaves it largely unperturbed. This asymmetry indicates that these proteins preferentially exert their disruptive effect from within an already-formed BiP–Env complex rather than by pre-empting Env binding to BiP.

In contrast, only two partners produced clear cooperative stabilisation. DNAJB11 strengthened AB from $-14.2$ to $-19.0$ $\text{kcal} \cdot \text{mol}^{-1}$ under AB conditioning ($\Delta\Delta G \approx -4.8$ $\text{kcal} \cdot \text{mol}^{-1}$), with BC essentially unchanged ($\Delta\Delta G \approx -0.2$ $\text{kcal} \cdot \text{mol}^{-1}$) and no meaningful perturbation from the reverse-conditioned branch ($\Delta\Delta G \approx +0.2$ $\text{kcal} \cdot \text{mol}^{-1}$). FICD produced a comparably clean stabilising signature: AB shifted from $-14.2$ to $-17.3$ $\text{kcal} \cdot \text{mol}^{-1}$ ($\Delta\Delta G \approx -3.1$ $\text{kcal} \cdot \text{mol}^{-1}$), with negligible BC change ($\Delta\Delta G \approx -0.1$ $\text{kcal} \cdot \text{mol}^{-1}$) and no significant reverse-branch effect. Both DNAJB11 and FICD stabilise the BiP–Env interface almost exclusively through the AB axis, defining the cleanest cooperative signatures in the entire BiP-centred panel.

P4HB was the sole near-neutral partner: AB shifted by only $\Delta\Delta G \approx +0.3$ $\text{kcal} \cdot \text{mol}^{-1}$ under AB conditioning, BC weakened by $\approx -0.3$ $\text{kcal} \cdot \text{mol}^{-1}$, and the reverse-conditioned AB perturbation reached only $+0.2$ $\text{kcal} \cdot \text{mol}^{-1}$ — all values consistent with energetic permissiveness rather than competitive antagonism or cooperative reinforcement.

The overall energetic classification is therefore unambiguous: DNAJB11 and FICD define a cooperative class that deepens the BiP–Env energetic basin; HYOU1, DERL1, HSP90B1, and PDIA6 constitute a strongly antagonistic class that disrupts it, by magnitudes ranging from $+5.2$ to $+9.4$ $\text{kcal} \cdot \text{mol}^{-1}$; and P4HB is permissive. The structured repartitioning of the conditional landscape across these partners reveals that the BiP–Env interface is not a fixed, context-independent binary contact but a proteostasis-network-dependent energetic state whose stability reflects the surrounding ER chaperone environment.

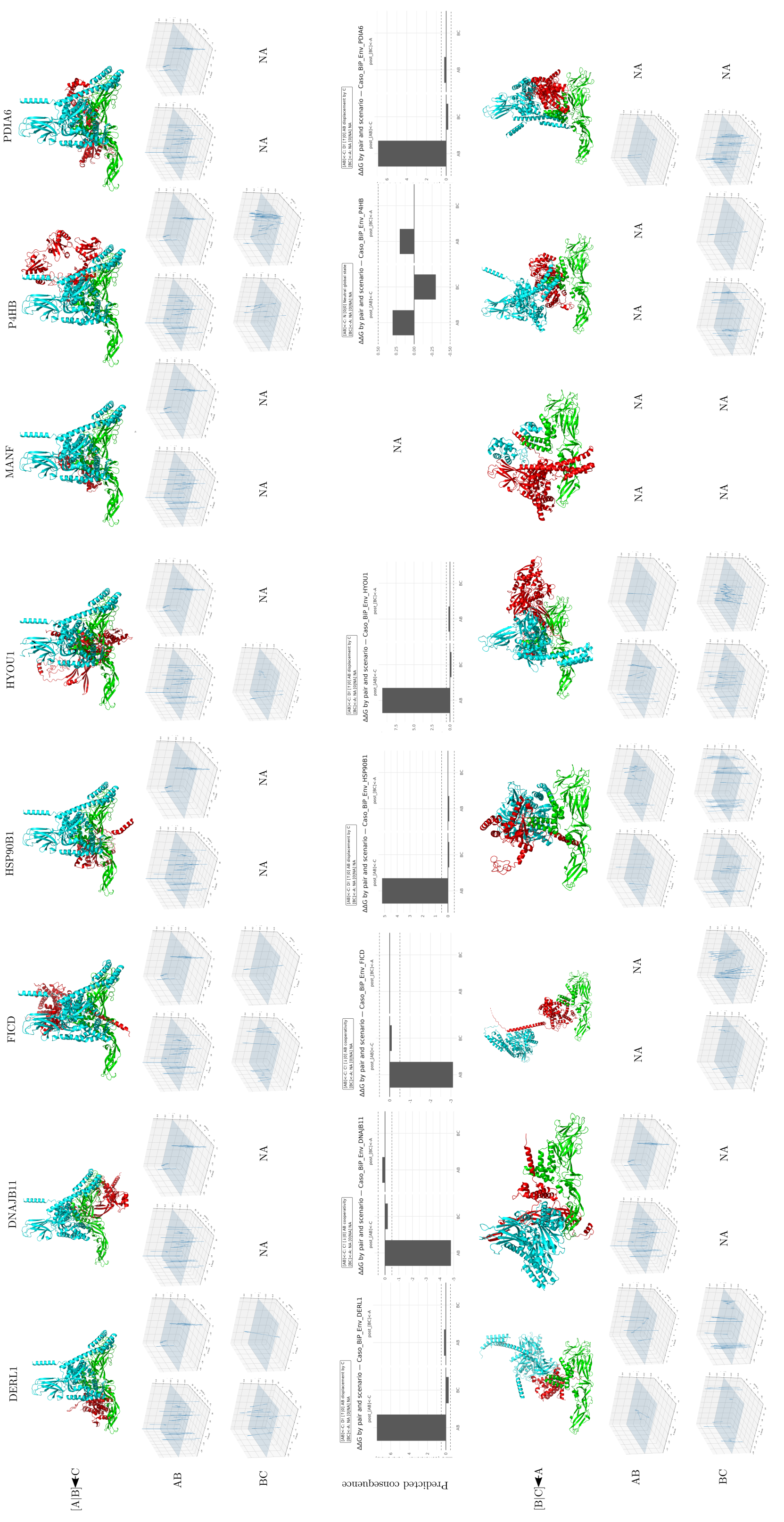
[A|B]→C
AB
BC
Predicted consequence
[B|C]→A
DERL1
DNAJB11
FICD
HSP90B1
HYOU1
MANF
P4HB
PDIA6
ΔΔG by pair and scenario — Caso_BiP_Env_DERL1
ΔΔG by pair and scenario — Caso_BiP_Env_DNAJB11
ΔΔG by pair and scenario — Caso_BiP_Env_FICD
ΔΔG by pair and scenario — Caso_BiP_Env_HSP90B1
ΔΔG by pair and scenario — Caso_BiP_Env_HYOU1
ΔΔG by pair and scenario — Caso_BiP_Env_P4HB
ΔΔG by pair and scenario — Caso_BiP_Env_PDIA6
NA

**Figure 12. Structural, interaction-field, and predicted energetic consequences of dengue Envelope–BiP-centered ternary complexes with ER-associated BiP partners.**

The figure compares ternary models involving dengue virus Envelope, BiP/HSPA5/GRP78, and eight BiP-associated endoplasmic-reticulum proteins: DERL1, DNAJB11, FICD, HSP90B1, HYOU1, MANF, P4HB, and PDIA6. Two alternative assembly scenarios are shown. In the first, the Envelope–BiP pair is treated as the preformed binary complex and the third BiP-associated partner is introduced afterward. In the reciprocal scenario, the BiP–partner pair is formed first and Envelope is subsequently introduced. Structural panels show the relative spatial organization of the three proteins in each modeled ternary assembly. Interaction-field plots summarize hydrogen-bond and hydrophobic contact distributions across the AB and BC interfaces, revealing localized polar and nonpolar interaction features. Predicted energetic consequence plots report ΔΔG values for each pair and assembly scenario, with negative values indicating stabilizing effects and positive values indicating destabilizing or energetically unfavorable effects relative to the corresponding reference state. Missing panels are marked as NA when the corresponding interaction descriptor or energetic estimate was unavailable. Overall, the figure indicates that dengue Envelope engagement with BiP is strongly influenced by the identity of BiP-associated ER partners and by the order of ternary complex formation, supporting a model in which the Envelope–BiP interface is embedded within a dynamic and context-dependent ER chaperone interaction network.

#### 3.6 Envelope–Clusterin Complex: Hydrophobic-Dominant Engagement and Extracellular Chaperone Exploitation

##### 3.6.1 Global Structural Stability and Differential Conformational Dynamics

The envelope–clusterin complex exhibits a dynamical profile distinct from those of the envelope–BiP and envelope–betaglycan systems, reflecting the fundamentally different structural architecture and substrate-binding mechanism of clusterin (CLU, apolipoprotein J). Clusterin is a secreted heterodimeric glycoprotein of approximately 75–80 kDa whose chaperone activity is mediated by two intrinsically disordered, hydrophobic peptide tails generated by post-translational cleavage of the precursor chain [85,86]. This architecture is reflected in the molecular dynamics trajectory: the backbone RMSD of clusterin displays a broader initial distribution and more prominent oscillatory behavior throughout the 200 ns simulation, consistent with the conformational heterogeneity of its three-domain core — a coiled-coil bundle, a disulfide-stabilized helical domain, and an $\alpha/\beta$ roll-like domain — and with the inherent flexibility of the disordered tails that serve as primary interaction surfaces [85,87]. The envelope protein, in contrast, rapidly attains a stable conformational plateau following the equilibration phase and maintains this regime throughout the production trajectory without evidence of progressive drift or unfolding.

Radius of gyration (Rg) analysis corroborates this differential behavior. The envelope retains a compact, stable Rg throughout the simulation, while clusterin exhibits modest and partially reversible Rg oscillations without any monotonic expansion event. This pattern is mechanistically consistent with clusterin's documented ability to engage structurally diverse substrates without requiring fixed geometric complementarity, instead relying on the conformational plasticity of its flexible tails to adaptively accommodate binding partners [28,88]. Importantly, the intramolecular pair number (numP) of each protein remains stable throughout the trajectory — a result not seen in mechanistically less stable complexes — indicating that neither the envelope nor clusterin undergoes appreciable loss of internal secondary or tertiary contacts upon complexation, confirming that the interaction occurs through external surface engagement without disrupting the folded core of either partner.

##### 3.6.2 Asymmetric Flexibility Redistribution: Tail Regions as the Primary Dynamic Interface

Per-residue RMSF analysis reveals a pronounced and mechanistically informative asymmetry in flexibility redistribution upon complex formation. Within clusterin, the most substantial reductions in per-residue fluctuations occur in segments corresponding to the disordered tail regions and their adjacent structural elements — specifically at the linker sequences connecting the coiled-coil bundle to the disulfide domain.

These segments, which are functionally equivalent to the N-terminal substrate-binding extensions of small heat shock proteins [85], display dampened fluctuations when engaged with the envelope surface, consistent with tail-mediated contact formation that partially immobilizes these freely mobile sequences. Peripheral helical regions of the coiled-coil bundle and the $\alpha/\beta$ roll domain retain comparatively higher flexibility, confirming that binding is spatially localized to the tail-proximal interface and does not propagate rigidity across the entire chaperone structure.

On the envelope side, RMSF reductions are concentrated at discrete surface patches geometrically accessible to the clusterin tails. Notably, the distribution of reduced-flexibility regions on the envelope surface does not spatially overlap with those identified in the envelope–BiP complex, suggesting that clusterin and BiP engage distinct epitopes on the viral protein — an observation directly relevant to the possibility of simultaneous or sequential chaperone engagement during viral entry.

###### 3.6.3 Interface Persistence and Contact Stabilization Without Progressive Compaction

The intermolecular contact profile of the envelope–clusterin complex is qualitatively distinct from those of the betaglycan and BiP systems. Unlike the monotonic contact number growth characteristic of the betaglycan interface, the envelope–clusterin contact number stabilizes rapidly following equilibration and thereafter fluctuates within a narrow range, without evidence of progressive interfacial densification. This rapid stabilization at a sustained plateau is consistent with a binding mode in which the primary interface is established through fast-kinetics tail engagement and maintained by the intrinsic affinity of hydrophobic surface contacts, rather than through slow geometric rearrangement toward greater complementarity.

The minimum interchain distance profile confirms continuous close proximity throughout the trajectory — with values compatible with direct van der Waals contacts and intermolecular hydrogen bonding — and the absence of episodic separation events indicates that the complex remains engaged without large-scale dissociation dynamics at the 200 ns timescale. The center-of-mass separation distribution is somewhat broader than in the BiP complex, reflecting the greater conformational freedom of the disordered tail interface, but remains fully consistent with a continuously and stably associated complex [47,89].

###### 3.6.4 MM/GBSA Energetics Reveal van der Waals–Dominant Binding with Attenuated Solvation Penalty

The MM/GBSA binding free energy profile of the envelope–clusterin complex presents a distinct energetic signature relative to both the betaglycan and BiP systems. The gas-phase contribution ($\Delta G_{\text{gas}}$) is strongly favorable, with the van der Waals term (VDWAALS) representing a proportionally larger share of $\Delta G_{\text{gas}}$

than in the predominantly electrostatic profiles of betaglycan and BiP, consistent with the hydrophobic character of the clusterin tail-mediated interface. The electrostatic term (EEL) contributes favorably but at reduced absolute magnitude relative to BiP ($\Delta G_{\text{gas}}$ EEL $\approx -568.4$ kcal/mol for BiP), consistent with a less polar interface. Correspondingly, the polar solvation penalty (EGB) is comparatively attenuated, because partial — rather than full — burial of ionizable groups reduces the magnitude of the desolvation cost relative to the BiP complex. The nonpolar solvation term (ESURF) remains consistently favorable, reflecting burial of hydrophobic surface area consistent with tail-mediated engagement. The resulting $\Delta G_{\text{total}}$ is moderately negative and qualitatively distinguishable from the near-compensatory electrostatic profiles of betaglycan and BiP, indicating thermodynamically viable complex formation through an energetically distinct mechanism [49,77].

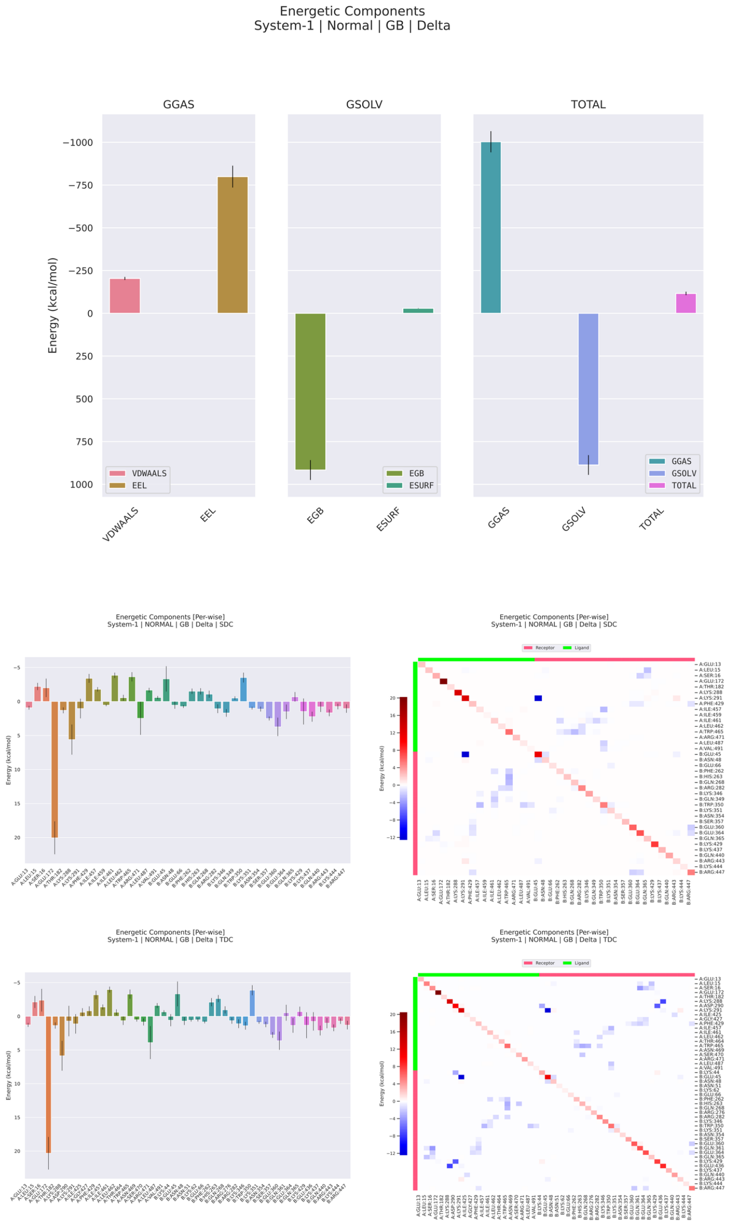


**Figure 13: MM/GBSA energetic decomposition of the dengue virus Envelope–clusterin complex during molecular dynamics simulation.** The figure summarizes the global energetic components, per-residue decomposition, and pairwise residue decomposition of the binding free energy estimated by MM/GBSA for the dengue Envelope–clusterin complex. The upper panels show a strongly favorable gas-phase contribution, with the electrostatic term representing the dominant stabilizing component and the van der Waals term providing a smaller, but still favorable, contribution. This large electrostatic gain is extensively counterbalanced by the polar generalized Born solvation penalty, indicating that complex formation imposes a substantial desolvation cost on charged and polar groups at the interface. The nonpolar surface-area solvation term contributes only weakly relative to the electrostatic and polar solvation terms. Despite this compensation, the total binding free energy remains favorable, indicating that the Envelope–clusterin association is energetically supported after solvation effects are considered. The per-residue decompositions obtained with the SDC and TDC schemes reveal a highly asymmetric energetic architecture, in which most residues contribute weakly, whereas a restricted subset of interface residues concentrates the strongest favorable or unfavorable effects. This pattern indicates that the complex is not stabilized by a diffuse, uniformly distributed interface, but by discrete energetic hotspots embedded within a broader contact surface. The pairwise heatmaps further support this interpretation, showing sparse and localized favorable residue-pair interactions rather than a continuous energetic network across the interface. Blue regions denote favorable residue-pair contributions, whereas red regions denote unfavorable or destabilizing contributions. The coexistence of intense favorable pairwise contacts with scattered unfavorable contributions suggests an interface shaped by local electrostatic complementarity, partial desolvation frustration, and selective residue-level stabilization. Overall, the MM/GBSA profile indicates that the Envelope–clusterin interface behaves as a hotspot-driven binding system: global stabilization is primarily electrostatic, strongly opposed by polar solvation, and ultimately maintained by a limited number of energetically dominant residue and residue-pair interactions.

##### 3.6.5 Residue-Level Decomposition Identifies Hydrophobic Anchor Residues and Peripheral Electrostatic Frustration

Per-residue free energy decomposition (SDC and TDC) reveals an interfacial energetic architecture organized around hydrophobic packing rather than the discrete charged anchor pairs that dominate the BiP interface. On the envelope surface, residues with hydrophobic or amphipathic character — most prominently aliphatic and aromatic side chains in the domain II surface region, including residues flanking the conserved fusion loop — provide the dominant favorable contributions through van der Waals–driven burial against clusterin tail sequences. The role of Trp101 and adjacent fusion loop residues (Leu107, Phe108 in DENV2) as hydrophobic hotspots is consistent with published NMR and mutagenesis data establishing these residues as principal drivers of hydrophobic contact at the dengue envelope surface [90,91]. On the clusterin surface, the strongest contributors map to hydrophobic and amphipathic residues within or adjacent to the disordered tail regions, whose side chains bury against the envelope surface upon tail engagement. Residues of the coiled-coil bundle contribute secondary favorable terms, while the disulfide-stabilized helical domain provides minimal energetic contributions, consistent with its structural rather than substrate-binding role [85].

Unfavorable decomposed energies are distributed among polar and ionizable residues at the periphery of the interface, where partial desolvation occurs without fully compensatory intermolecular contacts. This peripheral frustration is topologically distinct from the centrally positioned LYS435 frustration in the BiP complex and from the ARG107/LYS236 hotspot penalties in the betaglycan system, reinforcing the conclusion that each envelope–host protein interface presents an independently organized energetic landscape with characteristic stabilizing and destabilizing architectures.

##### 3.6.6 Conformational State Analysis: PCA and Hierarchical Clustering Reveal Differential Ensemble Behavior

Conformational diversity was assessed independently for the envelope and clusterin chains through principal component analysis (PCA) of C$\alpha$ coordinates and hierarchical agglomerative clustering of pairwise RMSD matrices computed using MDTraj [92]. The two-dimensional projection of the envelope trajectory onto principal components PC1 and PC2 reveals a temporally structured distribution, with conformations from different simulation intervals forming partially non-overlapping populations. Two principal clusters emerge, collectively accounting for the dominant fraction of sampled conformations, with an identifiable transitional region between them consistent with a single major conformational rearrangement occurring at an intermediate simulation time. The hierarchical dendrogram for the envelope chain confirms a large inter-cluster linkage distance separating two well-defined branches, whose temporal composition indicates an early-simulation state and a later-simulation state separated by a relatively rapid transition. This behavior is consistent with an induced-fit accommodation of the envelope surface in response to clusterin tail engagement, analogous to the conformational transitions documented for other protein–protein association events that involve flexible binding partners [93].

The clusterin chain exhibits a markedly different conformational landscape. The PC1–PC2 projection is substantially more dispersed, with conformational frames distributed across a broader region of phase space and without clear temporal segregation into distinct population clusters. Multiple smaller conformational groups are identifiable in the hierarchical dendrogram, with shorter inter-cluster linkage distances and more even population fractions, indicating continuous and multidirectional sampling of conformational substates throughout the trajectory. This behavior reflects the intrinsic multi-state dynamics of the clusterin heterodimer, whose disordered tail regions explore diverse orientations while the core domain retains its folded architecture [85,87]. The differential conformational diversity between the two partners — low for the envelope, high for clusterin — provides direct structural evidence for the adaptive-binding mechanism proposed for extracellular chaperones, in which the chaperone conforms to the substrate topology rather than imposing a fixed binding geometry.

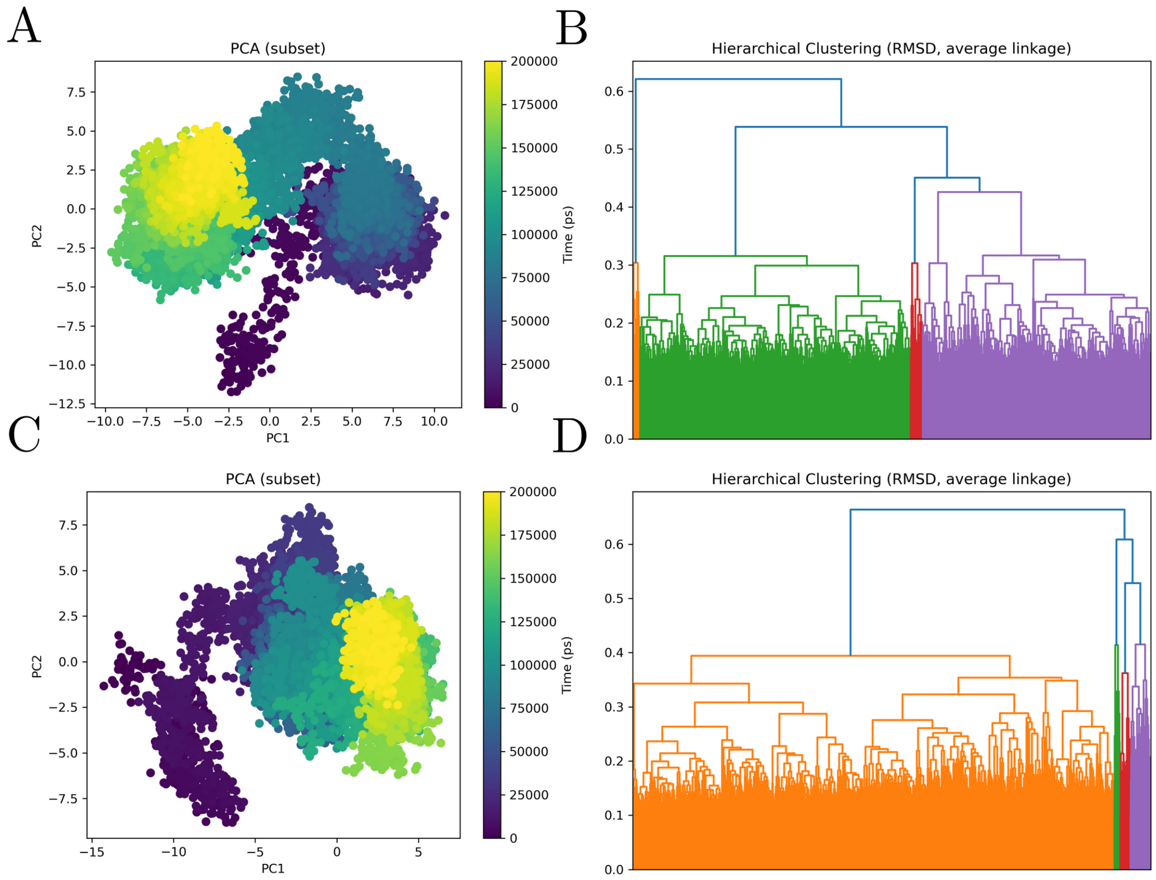


**Figure 14: Principal component analysis and RMSD-based hierarchical clustering of the dengue Envelope–clusterin molecular dynamics trajectory.** The figure summarizes the conformational landscape sampled during the molecular dynamics simulation of the dengue Envelope–clusterin complex, with panels A and B corresponding to the receptor subset (dengue Envelope) and panels C and D corresponding to the ligand subset (clusterin). Panels A and C show principal component analysis projections of trajectory frames onto the first two principal components, with points colored according to simulation time. In panel A, the dengue Envelope subset displays a broad and structured conformational manifold composed of partially overlapping basins connected by transitional regions, indicating that the receptor explores multiple related substates rather than remaining confined to a single rigid geometry. The temporal color gradient suggests progressive redistribution across this landscape, consistent with gradual conformational adaptation of the Envelope within the bound complex. In panel C, the clusterin subset exhibits a more directional conformational progression, with early frames occupying a more distant region of the projection and later frames concentrating in a more compact and organized basin. This pattern is consistent with ligand accommodation and progressive stabilization, suggesting that clusterin undergoes a clearer transition from initial exploratory states toward a more persistent late-stage conformational regime. Panels B and D show RMSD-based hierarchical clustering using average linkage, revealing that both molecular partners populate multiple structural families during the simulation. For dengue Envelope (panel B), the dendrogram indicates a heterogeneous but hierarchically organized distribution of conformers, supporting the existence of several receptor-associated substates with moderate structural separation. For clusterin (panel D), the clustering pattern is dominated by one major conformational family accompanied by smaller subsidiary clusters, reinforcing the interpretation from the PCA that the ligand evolves toward a more prevalent stabilized state while still retaining limited substate diversity. Together, the PCA and clustering analyses indicate that the dengue Envelope–clusterin complex is dynamically heterogeneous, but not disordered: the receptor samples a broader ensemble of related substates, whereas the ligand shows a more directed accommodation trajectory toward conformational stabilization. Overall, these results support a model in which the Envelope–clusterin interface is maintained by reciprocal adaptation between a structurally plastic receptor and a ligand that progressively settles into dominant bound conformations.

##### 3.6.7 Ternary Docking Reveals Chaperone- and Stress-Context-Dependent Modulation of the Clusterin–Env Interface

The ternary interaction analysis centered on the dengue virus Envelope–clusterin complex reveals that clusterin behaves not merely as an isolated host ligand, but as a context-sensitive molecular hub whose interaction with the viral Envelope can be substantially reshaped by additional cellular partners. In this analysis, dengue Envelope is represented as component A, clusterin as component B, and each clusterin-associated protein as component C. The figure compares two alternative assembly routes: one in which the Envelope–clusterin binary pair is formed first and the third cellular partner is introduced afterward, represented as $[A]B \leftarrow C$, and a reciprocal configuration in which clusterin is already associated with a cellular partner before dengue Envelope is introduced, represented as $[B]C \leftarrow A$. This design is important because it explicitly tests whether the structural and energetic behavior of the Envelope–clusterin interface is invariant across molecular contexts or, instead, depends on the pre-existing interaction state of clusterin.

Across the eight clusterin-associated proteins evaluated—ATP7B, BAX, BCL2L1, BiP, COMMD1, NUDCD1, SNCA, and XRCC6—the structural models show pronounced partner-dependent differences in the spatial arrangement of the ternary assemblies. The relative positioning of dengue Envelope, clusterin, and the third protein varies substantially among complexes, indicating that the ternary systems cannot be reduced to a simple superposition of independent binary interfaces. Instead, the addition of each C partner appears to impose a distinct geometric constraint on the Envelope–clusterin pair. In some cases, the third protein occupies a peripheral position relative to the Envelope–clusterin interface, suggesting a weaker or more indirect modulatory role. In other cases, the C partner is positioned in a way that may compete with, stabilize, shield, or redirect the interaction surface available to the viral Envelope. This structural heterogeneity supports the idea that clusterin-associated molecular environments may define alternative routes of viral engagement, where the same viral protein encounters different local topologies depending on the host protein network in which clusterin is embedded.

The interaction-field descriptors further reinforce this interpretation. The AB and BC field plots reveal that hydrogen-bond and hydrophobic contact features are not uniformly distributed across the ternary models, but instead appear as localized interaction densities that vary according to both partner identity and assembly order. This suggests that the formation of a ternary complex changes the spatial organization of polar and nonpolar contacts at the interfaces. In the $[A]B \leftarrow C$ scenario, the analysis asks how a pre-established Envelope–clusterin complex responds to the arrival of a clusterin-associated protein. In this configuration, the third partner may either preserve the original Envelope–clusterin interaction field, perturb it by inducing local rearrangements, or introduce additional interaction density that changes the balance between AB and

BC contacts. Conversely, in the $[B]C \leftarrow A$ scenario, the Envelope engages clusterin in a pre-organized cellular context. Here, the viral protein does not interact with a free clusterin molecule, but with clusterin already constrained by its association with another cellular protein. This distinction is biologically relevant because, inside the cell or at extracellular and membrane-associated compartments, clusterin is unlikely to exist as an isolated binding surface. Rather, it may participate in pre-existing molecular assemblies that condition how viral proteins encounter it.

The predicted energetic consequence plots indicate that the effect of ternary assembly is strongly partner-specific. Several complexes display marked positive $\Delta\Delta G$ values for particular interaction pairs or assembly scenarios, suggesting that the addition of the third component can impose destabilizing energetic consequences or unfavorable structural rearrangements relative to the corresponding reference state. This behavior is particularly evident for partners such as BAX, BCL2L1, COMMD1, SNCA, and XRCC6, where the energetic signatures suggest substantial ternary modulation of the system. These effects may reflect steric incompatibility, competition for overlapping regions of clusterin, disruption of favorable binary contacts, or desolvation and electrostatic penalties introduced by the new assembly geometry. Importantly, these positive energetic consequences should not be interpreted simply as failure of interaction. Rather, they indicate that the ternary state imposes a cost on specific pairwise relationships within the system, which may still be biologically meaningful if compensated by other contacts, transient binding, conformational selection, or local rearrangement of the interface.

By contrast, other partners show weaker, more localized, or partially compensatory energetic effects. NUDCD1, for example, appears to produce a comparatively modest energetic perturbation relative to the more strongly affected systems. This may indicate that some clusterin-associated proteins can coexist with the Envelope–clusterin interface without dramatically disrupting the global ternary arrangement. Such cases are important because they point to a possible distinction between disruptive modulators and permissive cofactors. Disruptive partners may reshape the Envelope–clusterin interface by imposing strong energetic penalties or forcing alternative conformations, whereas permissive partners may allow clusterin to remain compatible with viral engagement while adding only limited local perturbation. ATP7B represents an additional interpretative case because some descriptors are unavailable for specific interaction states, but the remaining panels still indicate that the clusterin–partner axis contributes measurable structural or energetic information. This suggests that, for certain systems, the modulation of dengue Envelope binding may be inferred more strongly from the state of the clusterin-associated partner than from the directly sampled Envelope–clusterin interface alone.

A central conclusion from this data is that the order of assembly matters. The upper and lower portions of the figure are not redundant representations of the same ternary complex; rather, they test two mechanistically distinct hypotheses. In the first, dengue Envelope binds clusterin before the cellular partner is introduced, approximating a situation in which viral engagement precedes recruitment or encounter of an additional host factor. In the second, clusterin is already complexed with a host protein before Envelope binding, approximating a situation in which dengue Envelope encounters a pre-organized clusterin-containing cellular module. Differences between these two scenarios indicate that ternary complex formation is path-dependent. The final structural and energetic state depends not only on which three proteins are present, but also on which binary interaction is formed first. This is consistent with a model of molecular recognition in which conformational availability, local steric constraints, interface exposure, and pre-existing interaction fields influence the subsequent binding behavior of the viral protein.

From a systems-level perspective, these results place clusterin as a potential mediator of higher-order host–virus interaction logic. The Envelope–clusterin interaction appears to be embedded within a broader protein-interaction context, where cellular partners may amplify, weaken, redirect, or energetically frustrate viral binding. This is especially relevant for clusterin because of its known functional association with protein homeostasis, extracellular chaperone-like activity, apoptosis-related pathways, lipid-associated processes, and stress-response networks. Although the present analysis does not assign a direct functional outcome to each ternary model, the structural and energetic patterns suggest that dengue Envelope may exploit or be constrained by clusterin-associated molecular states. In this sense, the Envelope–clusterin interface should not be interpreted as a single static docking event, but as a dynamic and network-dependent interaction that may vary according to the local host-protein environment.

Overall, these results suggests a model in which dengue Envelope engagement of clusterin is governed by ternary context, partner identity, and assembly order. The structural models demonstrate that each clusterin-associated protein creates a distinct spatial framework for the viral–host interface. The interaction-field descriptors show that hydrogen-bond and hydrophobic contact densities are redistributed across AB and BC interfaces in a partner-dependent manner. The predicted energetic consequences further indicate that some ternary combinations impose strong destabilizing or modulatory effects, whereas others are more compatible with preservation of the Envelope–clusterin association. Together, these results argue that clusterin functions as a network-embedded interaction platform for dengue Envelope, and that its role in viral recognition may depend on the molecular state in which clusterin is encountered. This provides a mechanistic basis for interpreting the Envelope–clusterin axis as part of a higher-order host-interface system, rather than as an isolated binary protein–protein interaction.

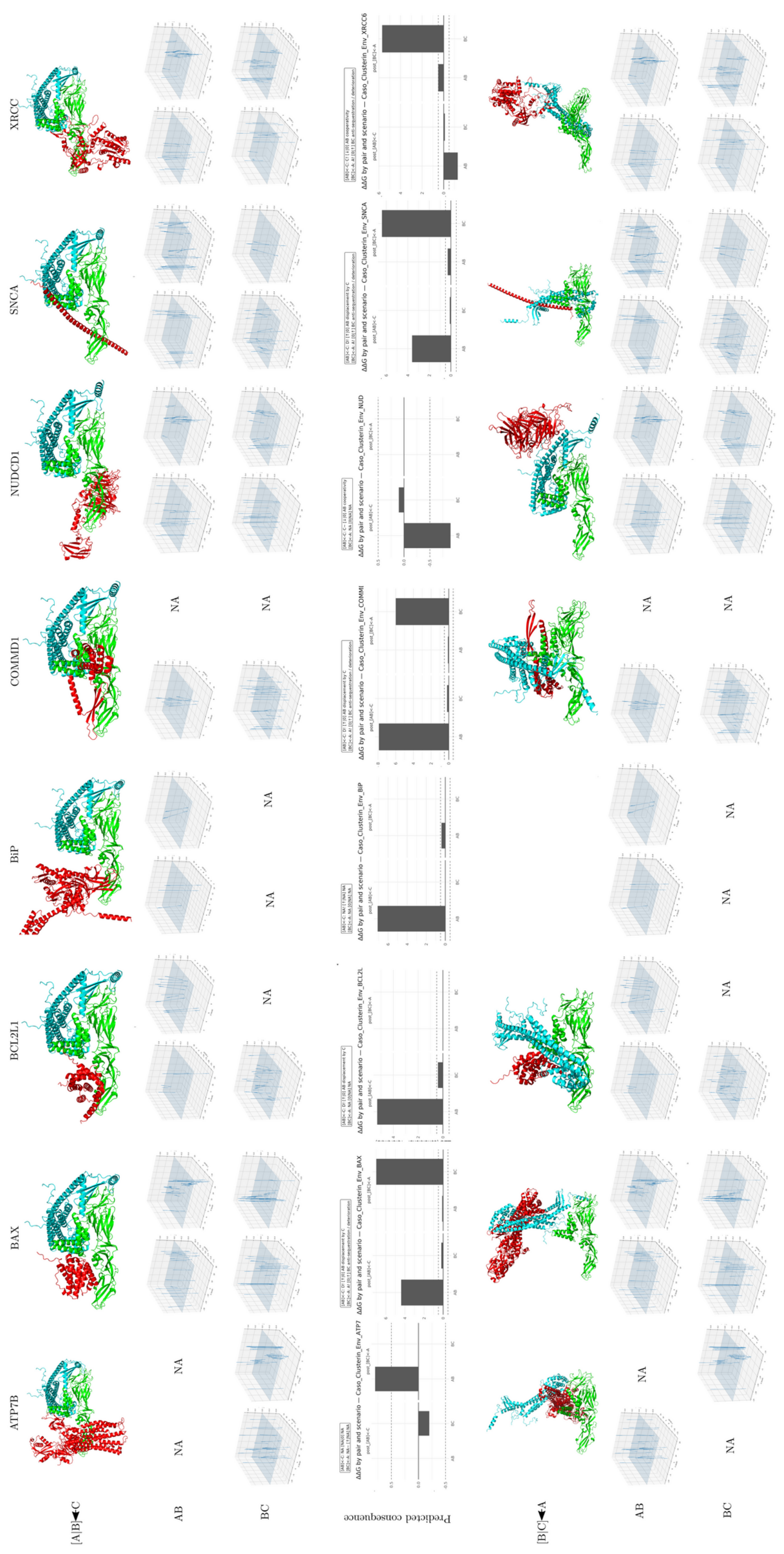
ATP7B
BAX
BCL2L1
BiP
COMMD1
NUDCD1
SNCA
XRCC
[A|B]◄C
AB
BC
Predicted consequence
[B|C]◄A
NA

**Figure 15. Structural, interaction-field, and predicted energetic consequences of dengue Envelope–clusterin-centered ternary complexes with clusterin-associated proteins.**

The figure summarizes comparative ternary interaction models involving dengue virus Envelope, clusterin, and eight clusterin-associated proteins: ATP7B, BAX, BCL2L1, BiP, COMMD1, NUDCD1, SNCA, and XRCC6. In this representation, A corresponds to dengue Envelope, B corresponds to clusterin, and C corresponds to the additional clusterin-associated cellular partner. Two alternative assembly scenarios are compared. In the first scenario, the Envelope–clusterin pair is treated as the preformed binary complex and the third protein is introduced afterward, represented as $[A]B \leftarrow C$. In the reciprocal scenario, the clusterin–associated-protein pair is treated as preformed and dengue Envelope is introduced as the third component, represented as $[B]C \leftarrow A$.

Structural panels show the relative spatial organization of the three proteins in each modeled ternary assembly, revealing marked partner-dependent differences in the orientation and packing of the Envelope, clusterin, and the cellular cofactor. Interaction-field plots report the spatial distribution of hydrogen-bond and hydrophobic contact features for the AB and BC interfaces, highlighting how polar and nonpolar interaction densities are redistributed according to both partner identity and assembly order. Predicted energetic consequence plots report $\Delta\Delta G$ values for each pair and assembly scenario, with negative values indicating stabilizing effects and positive values indicating destabilizing or energetically unfavorable effects relative to the corresponding reference state. Missing descriptors or unavailable configurations are indicated as NA.

Overall, the figure indicates that the dengue Envelope–clusterin interaction is strongly context-dependent and can be reshaped by the molecular environment of clusterin. The distinct structural arrangements, heterogeneous interaction-field patterns, and partner-specific energetic consequences support a model in which clusterin behaves as a network-embedded host interface rather than as an isolated viral ligand. In this framework, proteins such as BAX, BCL2L1, COMMD1, SNCA, and XRCC6 may impose substantial ternary modulation on the Envelope–clusterin interface, whereas other partners show weaker, more localized, or partially compensatory effects.

Together, these results suggest that dengue Envelope engagement of clusterin may depend not only on binary affinity, but also on the pre-existing cellular interaction state of clusterin and on the order in which ternary host–virus assemblies are formed.

#### 3.7 Envelope–Glypican-1 Complex: Electrostatic Saturation, Multivalent Tethering, and Directed Conformational Sampling

##### 3.7.1 Global Structural Stability Reflects High-Affinity Attachment-Phase Engagement

Glypican-1 (GPC1) is a GPI-anchored heparan sulfate proteoglycan (HSPG) whose core protein carries three to four heparan sulfate (HS) chains that project into the extracellular space and provide a highly anionic surface for electrostatic capture of basic viral surface proteins [94,95]. Among the host proteins examined in this study, GPC1 is the one most directly implicated in the initial virion attachment step — the concentration of DENV particles at the cell surface prior to engagement with entry receptors — a function that requires persistent, high-avidity contact without strict structural complementarity [96,97]. The 200 ns MD trajectory of the envelope–GPC1 complex is fully consistent with this attachment-phase role: both partners maintain stable global conformations throughout the simulation, with no evidence of progressive RMSD drift, domain separation, or partial unfolding.

The radius of gyration (Rg) of the envelope protein remains within a narrow band throughout the trajectory, indicating that the three-domain architecture of the ectodomain is preserved without inter-domain rearrangements on the simulation timescale. GPC1 likewise shows a constant Rg, consistent with the maintenance of its ellipsoidal core protein fold, which is stabilized by multiple intrachain disulfide bonds and a compact $\alpha$-helical stem [98]. This bilateral structural preservation differentiates the envelope–GPC1 complex from the envelope–betaglycan system, in which substantial host-side RMSD accumulation is observed over the trajectory, and aligns it more closely with the envelope–actin complex in terms of the symmetry of conformational rigidity across the interface.

##### 3.7.2 Asymmetric RMSF Profile Identifies the Domain III Patch as the Interface Anchor

Per-residue RMSF analysis reveals a pronounced asymmetry between the two partners. The envelope protein exhibits low backbone fluctuations across its structured secondary elements, with flexibility concentrated at surface-exposed loops — particularly within domains I and II — that are distant from the interface. In contrast, the region spanning domain III (DIII) of the envelope, which contains the electropositive patch of basic residues (Arg, Lys) known to constitute the primary HS-binding site in flaviviruses [94,95], displays among the lowest per-residue RMSF values in the entire envelope chain. This localized rigidification of DIII is consistent with the spatial constraint imposed by persistent electrostatic contacts with the GPC1 surface, reducing the conformational freedom of a region that in the unliganded state tends to explore wider angular deviations. The effect mirrors the RMSF dampening observed at the ARG410-dominated interface in the

BiP complex, but is spatially distinct — localized to DIII rather than to the domain I–II hinge — confirming that GPC1 and BiP engage non-overlapping epitopes on the envelope surface.

For GPC1, per-residue RMSF values are uniformly low across the ordered core protein, with elevated fluctuations confined to the N- and C-terminal extensions and the HS-attachment loops. Importantly, the segments immediately adjacent to the HS insertion sites show moderately higher flexibility, consistent with the known conformational freedom of the HS chain-bearing stalks that must accommodate the geometric requirements of multivalent surface binding [98]. This flexibility of the HS-proximal regions in the simulation model — even in the absence of explicit HS chains — suggests that the core protein maintains its recognition-competent geometry while permitting the dynamic reorientation of binding arms that characterizes multivalent proteoglycan function.

###### 3.7.3 Persistent Contact Network and Stable Minimum Distance Confirm Continuous Engagement

The intermolecular contact number profile rises rapidly during the equilibration phase and thereafter stabilizes at an elevated plateau, without systematic decay throughout the 200 ns trajectory. The minimum interchain distance remains consistently in the range of approximately 0.2–0.3 nm, values compatible with direct van der Waals contacts and hydrogen bonding, and does not exhibit transient increases that would indicate partial dissociation or large-scale interface sampling. This contact persistence is notably more pronounced than that observed in the betaglycan complex — which shows progressive contact growth — and differs from the BiP complex's moderate fluctuation profile, suggesting that the GPC1 interface achieves rapid saturation and subsequently maintains it, rather than continuing to evolve toward closer packing. The persistence of the center-of-mass distance within a narrow range further confirms that no large-scale rigid-body reorientation of GPC1 relative to the envelope occurs after initial engagement, consistent with a docked complex that has reached its minimum free energy conformation within the accessible simulation timescale.

The number of intramolecular interaction pairs (numP) for both the envelope and GPC1 remains stable throughout the trajectory, with no sustained reduction that would indicate partial unfolding or loss of secondary structure elements. This bilateral maintenance of internal contacts distinguishes the GPC1 complex from scenarios involving induced unfolding of either partner, and is consistent with the known tolerance of HSPG-mediated interactions for functional engagement without structural perturbation of either participant [99].

##### 3.7.4 MM/GBSA Energetics Reveal Electrostatically Saturated Interface with Dominant Desolvation Penalty

The MM/GBSA binding free energy profile computed over the final 10 ns of the trajectory exhibits a characteristic signature of highly charged protein–proteoglycan interfaces. The gas-phase contribution ($\Delta G_{\text{gas}}$) is dominated by the electrostatic term (EEL), which fluctuates within approximately −520 to −750 kcal/mol, representing by far the largest component of stabilization energy and the most volatile term across the trajectory. The van der Waals contribution (VDWAALS) provides a consistently favorable but comparatively modest contribution in the range of approximately −180 to −225 kcal/mol, with a modest rightward drift toward less negative values in the later portion of the window, consistent with subtle interface reorganization without loss of contact. The combined gas-phase stabilization ($\Delta G_{\text{gas}} \approx -800$ kcal/mol) is nearly compensated by the polar solvation penalty (EGB $\approx +700$ kcal/mol), resulting in a net binding free energy ($\Delta G_{\text{total}}$) centered near −90 kcal/mol, with individual frame values ranging from approximately −60 to −120 kcal/mol.

This large-magnitude EEL/EGB compensation profile is the defining energetic signature of interfaces stabilized by salt bridges and hydrogen bond networks between charged residues, as canonically observed in HSPG-mediated viral attachment interactions where electropositive surface patches on the viral protein engage electronegative sulfate and carboxylate groups of the HS chains [95,99]. In MM/GBSA, this compensation arises because the formation of charge–charge contacts releases large electrostatic stabilization energy, but the removal of solvating water molecules from charged groups incurs a nearly commensurate desolvation cost [84]. The temporal oscillations in $\Delta G_{\text{total}}$ within the −60 to −120 kcal/mol range reflect the dynamic alternation between contact microconfigurations — local rearrangements of salt bridges and H-bond partners at the interface — without destabilizing the complex macroscopically, a behavior consistent with the contact-persistence data described above.

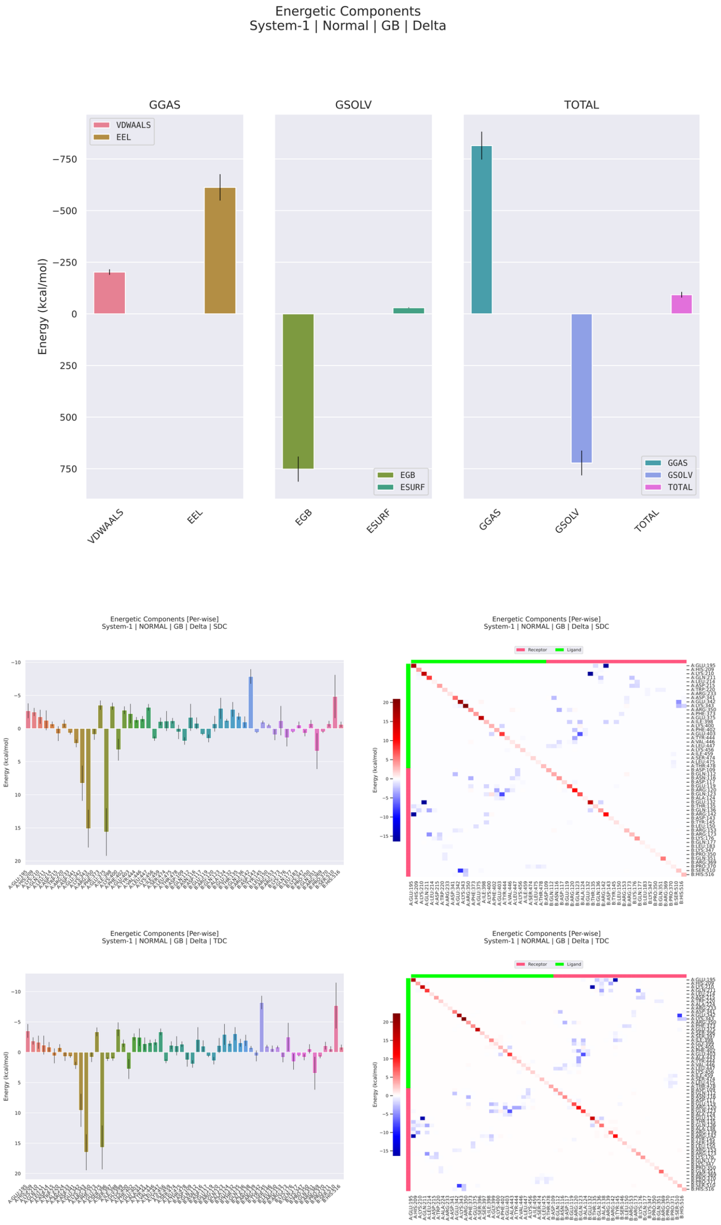


**Figure 16: MM/GBSA energetic decomposition of the dengue virus Envelope–glypican-1 complex during molecular dynamics simulation.** The figure summarizes the global energetic components, per-residue decomposition, and pairwise residue decomposition of the binding free energy estimated by MM/GBSA for the dengue Envelope–glypican-1 complex. The upper panels show that the gas-phase contribution is strongly favorable and dominated by the electrostatic term, with the van der Waals component providing an additional favorable contribution of lower magnitude. This profile indicates that the Envelope–glypican-1 association is primarily driven by long-range and charge-dependent interactions, consistent with the expected importance of electrostatic complementarity in interfaces involving glycosaminoglycan-associated or membrane-proximal host factors. The solvation component is largely unfavorable and dominated by the polar generalized Born term, indicating that complex formation imposes a substantial desolvation penalty on polar and charged groups at the interface. The nonpolar surface-area term contributes only weakly relative to the dominant electrostatic and polar solvation components. Despite the strong compensation between favorable gas-phase interactions and unfavorable polar solvation, the total binding free energy remains favorable, supporting the energetic stability of the dengue Envelope–glypican-1 complex after solvent effects are incorporated. The middle and lower panels show per-residue and pairwise decompositions using SDC and TDC schemes, revealing a markedly heterogeneous energetic interface in which most residues contribute weakly, whereas a restricted subset of residues produces dominant stabilizing or destabilizing effects. Negative values indicate favorable contributions to binding, whereas positive values indicate unfavorable or destabilizing energetic terms. In both decomposition schemes, the presence of intense residue-level peaks indicates that the interface is organized around discrete energetic hotspots rather than a uniformly distributed contact surface. The pairwise heatmaps further support this interpretation by showing sparse, localized favorable residue-pair interactions interspersed with weaker unfavorable contributions. Blue regions denote favorable residue-pair interactions, whereas red regions denote unfavorable or destabilizing contributions. Overall, the MM/GBSA profile indicates that the dengue Envelope–glypican-1 interface [illegible]

##### 3.7.5 Residue-Level Decomposition Identifies a Basic-Residue Hotspot Cluster and Peripheral Frustration Loci

Per-residue free energy decomposition (SDC) identifies a spatially concentrated cluster of energetically dominant residues on the envelope surface whose favorable contributions drive recognition. The most prominent hotspot corresponds to a basic residue on the envelope surface — consistent with an arginine in the domain III electropositive patch — contributing approximately −8 kcal/mol. This magnitude is quantitatively comparable to the ARG410 anchor identified in the BiP complex and to the ARG-mediated stabilization described for betaglycan, but it is mechanistically distinct: whereas the BiP hotspot participates in a charge-complementary salt bridge with a specific glutamate counterpart, and the betaglycan hotspots reflect electrostatic frustration under incomplete GAG compensation, the GPC1 hotspot likely engages multiple HS chain sulfate groups simultaneously, reflecting the multivalent character of HSPG recognition. Pairwise decomposition (TDC) confirms that the dominant inter-residue interactions involve the same basic envelope residue in contact with electronegative counterparts on the GPC1 surface, generating pair-wise energies consistent with salt bridge geometry.

The SDC profile also identifies energetically unfavorable residues at the interface periphery, with desolvation penalties reaching approximately +15 to +17 kcal/mol for the most frustrated positions. These frustration hotspots correspond to residues that are partially buried at the interface without forming adequate charge-compensation contacts — either polar groups that lose solvation energy without a fully compensated H-bond partner, or charged groups whose like-charge geometry generates electrostatic repulsion in the contact configuration sampled by the simulation. The presence of these frustration loci alongside the dominant favorable hotspots is characteristic of real protein–proteoglycan interfaces, which are rarely perfectly optimized and typically contain both anchoring and frustrated regions whose relative balance determines net affinity [100].

##### 3.7.6 Conformational State Analysis: Directed Envelope Transition and Adaptive Glypican-1 Sampling

Principal component analysis of C$\alpha$ coordinates and hierarchical RMSD-based clustering [92] reveal a qualitatively distinct conformational behavior for each partner that complements the contact-persistence and energetic data described above. The two-dimensional PC1–PC2 projection of the envelope chain traces a spatially directed arc — transitioning from an initial conformational region in the early simulation to a distinct later-simulation population, with minimal backward excursion — whose temporal structure is consistent with a single major progressive conformational rearrangement rather than stochastic fluctuation around a fixed mean. The hierarchical dendrogram for the envelope confirms two well-separated major clusters with an inter-cluster linkage distance of approximately 0.45, corresponding to the pre- and post-rearrangement conformational families, with the transition occurring at an intermediate simulation time. This directed conformational evolution is interpretable as an induced-fit adjustment of the DIII surface geometry in response to electrostatic capture by GPC1, consistent with the documented plasticity of flavivirus DIII under receptor engagement [94].

In contrast, the GPC1 chain explores a substantially broader and less temporally organized conformational space, with PC1–PC2 frames distributed across multiple partially overlapping populations throughout the trajectory. The hierarchical dendrogram reveals a larger number of clusters with smaller inter-cluster distances (maximum linkage height approximately 0.5) and more balanced population fractions compared to the envelope, indicating continuous multi-directional sampling of conformational substates. The density of frames increases progressively toward the later simulation time in the terminal conformational region of the PC space — a signature consistent with gradual stabilization of a preferred binding pose following the initial envelope rearrangement. This behavior differs from the clusterin pattern (where conformational multiplicity reflects disordered tail dynamics) and from the betaglycan pattern (where progressive conformational evolution reflects geometric compaction), and instead reflects the intrinsic flexibility of the GPC1 core protein together with the reorientational freedom of the HS-bearing loops exploring multiple contact configurations on the DIII surface.

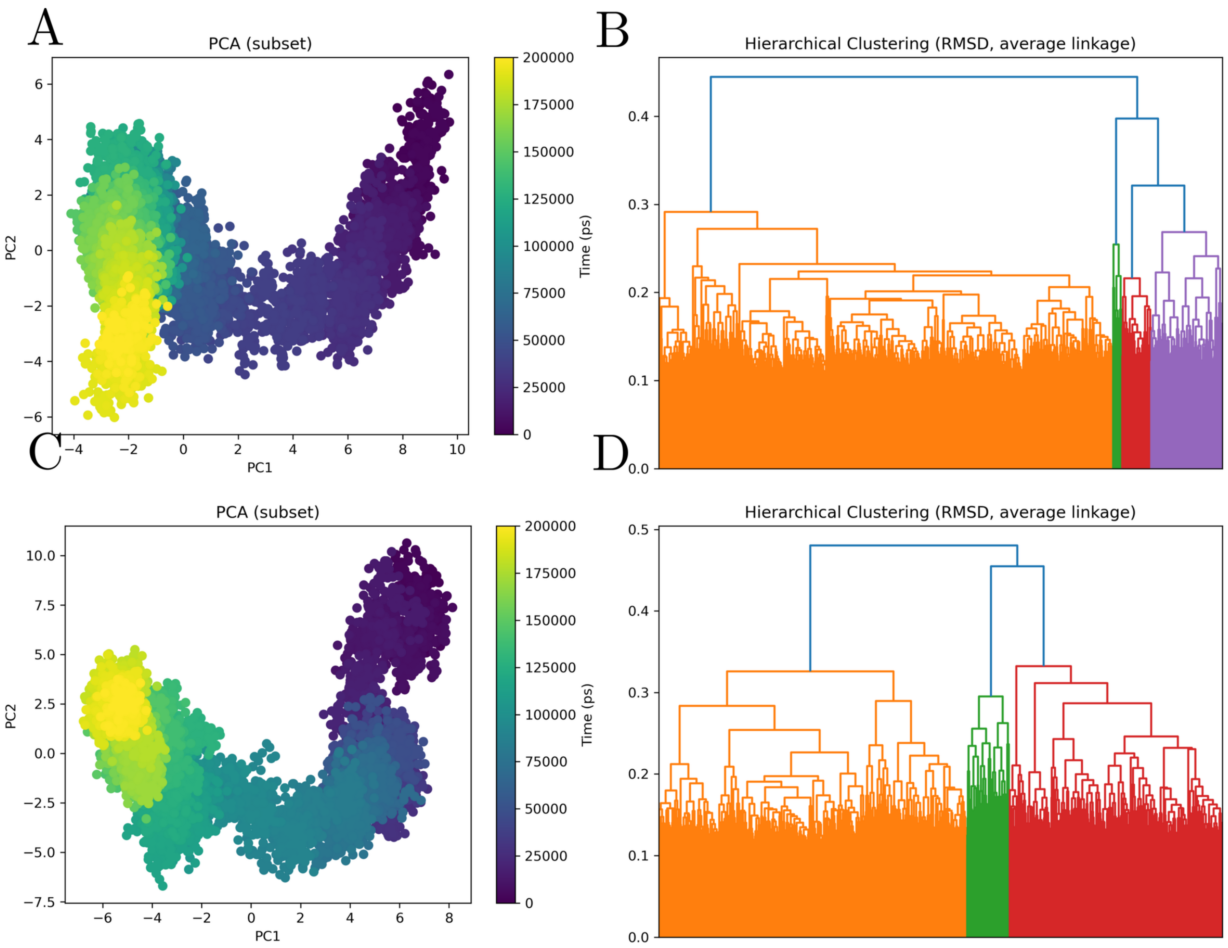


**Figure 17: Principal component analysis and RMSD-based hierarchical clustering of the dengue Envelope–glypican-1 molecular dynamics trajectory.** The figure summarizes the conformational landscape sampled during the molecular dynamics simulation of the dengue Envelope–glypican-1 complex, with panels A and B corresponding to the receptor subset, dengue Envelope, and panels C and D corresponding to the ligand subset, glypican-1. Panels A and C show principal component analysis projections of trajectory frames onto the first two principal components, with points colored according to simulation time. In panel A, the dengue Envelope subset displays a pronounced directional redistribution across the conformational space, progressing from an early region located at positive PC1 values toward a later, more compact region at negative PC1 values. This trajectory indicates that the receptor undergoes substantial conformational drift during the simulation, rather than remaining trapped in a single equilibrium basin. The relatively continuous connection between early, intermediate, and late regions suggests a gradual conformational transition compatible with progressive adaptation of the Envelope within the glypican-1-bound state. In panel C, the glypican-1 subset shows a broadly similar but more expanded conformational trajectory, with early frames occupying a distinct positive-PC1/high-PC2 region and later frames accumulating in a separate negative-PC1 region. This pattern indicates that the ligand also undergoes marked conformational redistribution, consistent with glypican-1 accommodation and remodeling within the complex. The stronger separation between early and late regions in the ligand projection suggests that glypican-1 contributes actively to the structural evolution of the bound system, rather than acting as a passive binding surface. Panels B and D show RMSD-based hierarchical clustering using average linkage, revealing multiple structural families within both receptor- and ligand-associated subsets. The receptor dendrogram in panel B shows one dominant conformational family accompanied by smaller late-trajectory clusters, supporting the existence of a major Envelope-associated structural basin followed by subsidiary conformational substates. The glypican-1 dendrogram in panel D shows a clearer partition into large conformational groups, indicating that the ligand samples multiple structurally distinguishable states during the simulation. Together, the PCA and clustering analyses indicate that the dengue Envelope–glypican-1 complex is dynamically heterogeneous but organized, with both molecular partners undergoing coordinated conformational redistribution. Overall, these results support a model in which glypican-1 binding is associated with progressive remodeling of the Envelope-containing complex, generating an ensemble of related but distinct conformational states rather than a single rigid docking pose.

##### 3.7.7 Ternary Docking Reveals Selective Destabilisation of the Glypican-1–Env Interface by Heparan-Sulfate-Dependent Growth Factors

The glypican-1-centered ternary docking analysis indicates that the dengue Envelope–glypican-1 interface is strongly modulated by growth-factor-associated molecular neighborhoods. In this representation, dengue Envelope corresponds to component A, glypican-1 corresponds to component B, and the additional glypican-1-associated partners, FGF1 and HBEGF1, correspond to component C. The analysis compares two reciprocal assembly scenarios: one in which the Envelope–glypican-1 binary complex is treated as preformed and the third partner is introduced afterward, represented as $[A]B \leftarrow C$, and a second in which the glypican-1–growth-factor complex is formed first and dengue Envelope is introduced subsequently, represented as $[B]C \leftarrow A$. This distinction is mechanistically important because glypican-1 is not expected to function as an isolated binding surface in the cellular environment, but rather as a membrane-proximal scaffold embedded in growth-factor, extracellular-matrix, and receptor-associated interaction networks.

The structural models show that FGF1 and HBEGF1 impose distinct ternary geometries on the Envelope–glypican-1 system. In the $[A]B \leftarrow C$ scenario, FGF1 appears positioned close to the Envelope–glypican-1 assembly in a compact arrangement, suggesting that the additional growth factor may introduce local steric and interaction-field constraints near the preformed viral–host interface. By contrast, HBEGF1 forms an elongated structural element that spans across the complex and contacts or approaches the Envelope–glypican-1 surface in a more extended configuration. This difference is relevant because an extended third partner may redistribute interaction density over a larger interfacial area, whereas a compact partner may produce more localized perturbations. Thus, even though both FGF1 and HBEGF1 are biologically connected to glypican-associated signaling environments, they generate different structural solutions when modeled as ternary partners of the dengue Envelope–glypican-1 complex.

The interaction-field plots reveal that the AB interface, corresponding to Envelope–glypican-1, retains detectable hydrogen-bond and hydrophobic contact features in both ternary contexts, but their spatial distribution varies according to the identity of the growth-factor partner and the assembly route. In the preformed Envelope–glypican-1 scenario, the AB fields remain prominent, indicating that the viral–glypican interface can persist after addition of the third component. However, the BC fields are more restricted and, in the case of HBEGF1, partially unavailable for hydrophobic descriptors, suggesting that the glypican-1–growth-factor interface may be more selectively represented by polar contacts or may be less completely captured depending on the docking geometry. This asymmetry between AB and BC fields indicates that the ternary systems are not balanced assemblies in which both pairwise interfaces contribute equally. Instead, each ternary configuration appears to redistribute interaction density in a partner-specific manner.

The predicted energetic consequence plots provide an additional layer of interpretation. For both FGF1 and HBEGF1, the $[A]B \leftarrow C$ scenario shows a positive $\Delta\Delta G$ contribution for the AB pair, indicating that addition of the growth-factor partner is predicted to destabilize or energetically perturb the pre-existing Envelope–glypican-1 interface. This suggests that FGF1 and HBEGF1 may act as modulators that impose a cost on the viral–glypican interaction when they are recruited after the Envelope–glypican-1 complex has already formed. The BC contribution in this same scenario remains close to neutral or slightly negative, indicating that the growth-factor–glypican interaction itself is not predicted to be strongly destabilized under this assembly order. In practical terms, this means that the third partner may be accommodated by glypican-1 without major energetic deterioration of the BC interface, while still perturbing the Envelope–glypican-1 contact.

The reciprocal $[B]C \leftarrow A$ scenario produces a different energetic pattern. When the glypican-1–growth-factor pair is treated as preformed and dengue Envelope is introduced afterward, the AB pair remains close to neutral or only weakly affected, whereas the BC pair shows a strong positive $\Delta\Delta G$ contribution for both FGF1 and HBEGF1. This suggests that Envelope binding to a pre-organized glypican-1–growth-factor module may impose a substantial energetic penalty on the original glypican-associated interaction. In other words, when glypican-1 is already engaged with FGF1 or HBEGF1, the subsequent arrival of dengue Envelope is predicted to disturb the glypican-1–growth-factor interface more strongly than the Envelope–glypican-1 interface itself. This behavior is consistent with a model in which dengue Envelope may compete with, distort, or allosterically displace pre-existing glypican-centered growth-factor assemblies.

The contrast between the two assembly orders is one of the most important results of this figure. If Envelope binds glypican-1 first, the growth-factor partner primarily perturbs the AB interface. If the growth-factor complex forms first, Envelope introduction primarily perturbs the BC interface. This reciprocal pattern suggests that the final ternary state is path-dependent: the energetic consequence of the same three molecular components depends on which binary interaction is established first. Such behavior supports the broader hypothesis that dengue Envelope recognition is not governed only by binary affinity, but by the structural and energetic state of the host-protein neighborhood at the moment of viral engagement.

Taken together, the glypican-1 ternary models support a context-dependent mechanism in which growth-factor-associated partners reshape the Envelope–glypican-1 interface through both geometric and energetic effects. FGF1 and HBEGF1 do not simply decorate the binary complex; they alter the distribution of hydrogen-bond and hydrophobic interaction fields and generate assembly-order-dependent energetic penalties. These findings are compatible with a model in which glypican-1 acts as a membrane-proximal host scaffold that can connect dengue Envelope recognition to growth-factor-associated molecular environments. In this framework, dengue Envelope engagement of glypican-1 may be influenced by whether glypican-1 is free, already associated with signaling ligands, or incorporated into a larger receptor-proximal complex. The figure therefore reinforces the idea that viral attachment and host-factor recognition should be interpreted as higher-order interaction processes embedded in local protein neighborhoods, rather than as isolated pairwise docking events.

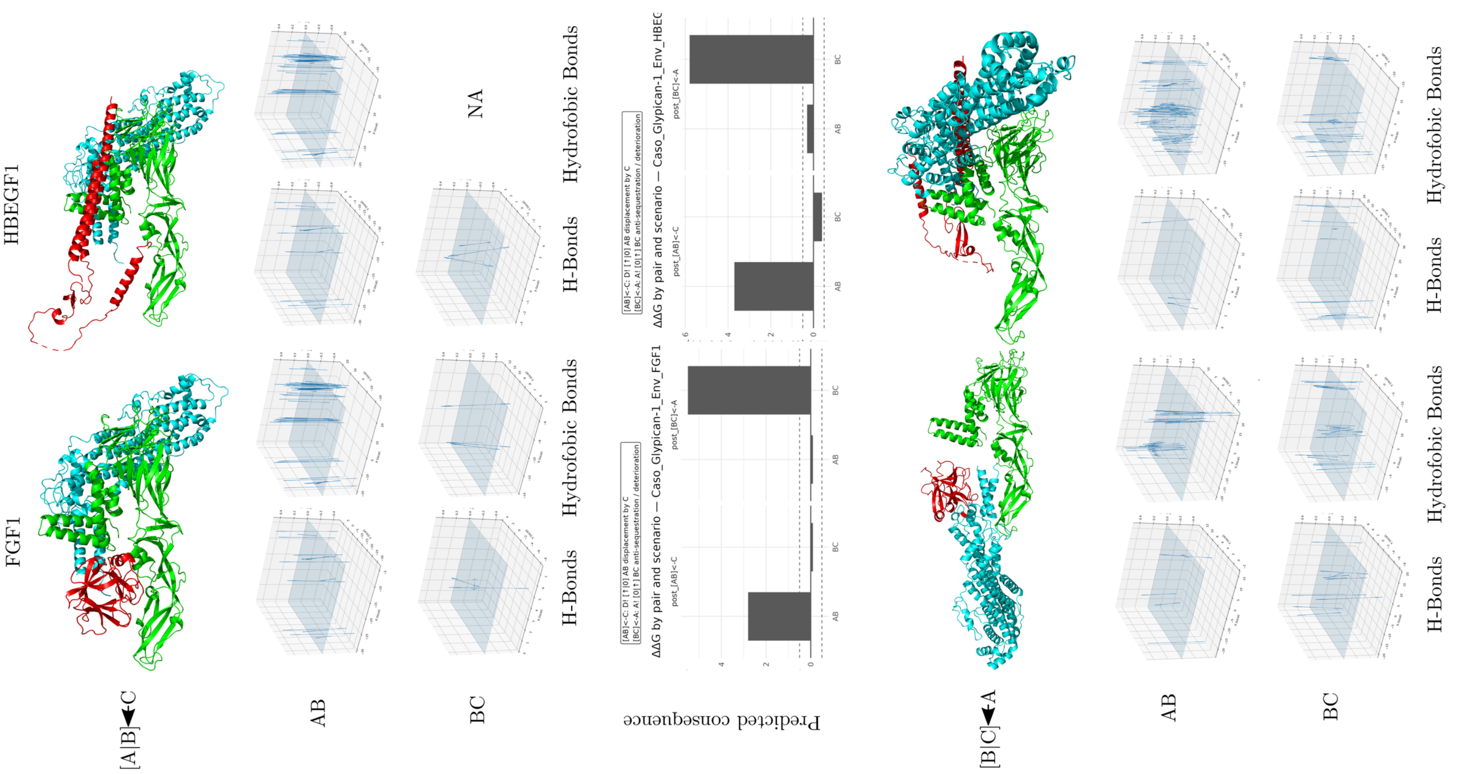
FGF1
HBEGF1
[A|B]◄C
AB
BC
H-Bonds
Hydrofobic Bonds
NA
Predicted consequence
ΔΔG by pair and scenario — Caso_Glypican-1_Env_FGF1
ΔΔG by pair and scenario — Caso_Glypican-1_Env_HBEG
post_[AB]<-C
post_[BC]<-A
[B|C]◄A

**Figure 18. Structural, interaction-field, and predicted energetic consequences of dengue Envelope–glypican-1-centered ternary complexes with growth-factor-associated partners.**

The figure summarizes comparative ternary interaction models involving dengue virus Envelope, glypican-1, and two glypican-1-associated growth-factor partners, FGF1 and HBEGF1. In this representation, A corresponds to dengue Envelope, B corresponds to glypican-1, and C corresponds to the additional glypican-1-associated protein. Two alternative assembly scenarios are compared. In the first scenario, the Envelope–glypican-1 pair is treated as the preformed binary complex and the third protein is introduced afterward, represented as $[A]B \leftarrow C$. In the reciprocal scenario, the glypican-1–growth-factor pair is treated as preformed and dengue Envelope is introduced as the third component, represented as $[B]C \leftarrow A$.

Structural panels show the relative spatial organization of the three proteins in each modeled ternary assembly, revealing distinct partner-dependent geometries for FGF1 and HBEGF1. FGF1 forms a more compact ternary arrangement, whereas HBEGF1 adopts a more extended configuration that spans the Envelope–glypican-1 surface. Interaction-field plots report the spatial distribution of hydrogen-bond and hydrophobic contact features for the AB and BC interfaces, showing that polar and nonpolar interaction densities are redistributed according to both partner identity and assembly order. Predicted energetic consequence plots report $\Delta\Delta G$ values for each pair and assembly scenario, with negative values indicating stabilizing effects and positive values indicating destabilizing or energetically unfavorable effects relative to the corresponding reference state.

Missing descriptors or unavailable configurations are indicated as NA. Overall, the figure indicates that the dengue Envelope–glypican-1 interaction is strongly modulated by growth-factor-associated molecular context. When the Envelope–glypican-1 pair is preformed, addition of FGF1 or HBEGF1 preferentially perturbs the AB interface, whereas introduction of Envelope into preformed glypican-1–growth-factor assemblies imposes a stronger energetic penalty on the BC interface. This reciprocal behavior supports a path-dependent model of ternary complex formation, in which the structural and energetic outcome depends not only on the identity of the interacting proteins, but also on the order in which the host and viral components assemble.

Together, these results suggest that glypican-1 may function as a membrane-proximal scaffold linking dengue Envelope recognition to growth-factor-associated host interaction neighborhoods.

#### 3.8 Envelope–Neuropilin-1 Complex: VEGF-Mimicry, CendR-Like Pocket Geometry, and Vascular Co-Receptor Engagement

##### 3.8.1 Global Structural Dynamics Reflect a Rigid Receptor Accommodating a Constrained Viral Ligand

Neuropilin-1 (NRP1) is a type I transmembrane co-receptor that integrates signals from vascular endothelial growth factor (VEGF), semaphorin class 3 (SEMA3), and a broad range of ligands bearing the C-end rule (CendR) motif — a carboxy-terminal R/KXXR/K sequence that binds with high specificity to the NRP1 b1 domain [101,102]. In the context of viral infection, NRP1 has been established as an entry-facilitating co-receptor for SARS-CoV-2, where the furin-cleaved spike protein exposes a CendR motif that engages the NRP1 b1 pocket [19,103], and is implicated as a dengue virus co-receptor through its role in mediating endothelial internalization and vascular permeability signaling downstream of VEGF-A [104,105]. The structural basis for the NRP1–dengue envelope interaction characterized here reflects this dual functional context: NRP1 is interrogated as both a direct binding partner for envelope-exposed surface elements and as a vascular signaling co-receptor whose engagement could amplify DENV-induced endothelial dysfunction.

The 200 ns MD trajectory of the envelope–NRP1 complex reveals bilateral structural stability in both partners. The envelope protein maintains a constant radius of gyration throughout the simulation, with no measurable RMSD drift after the initial equilibration phase, indicating that the three-domain ectodomain architecture is fully preserved under the intermolecular contacts imposed by NRP1. The NRP1 chain — encompassing the b1 domain, the primary ligand-binding unit responsible for CendR and VEGF-A recognition — likewise maintains a stable Rg, consistent with the preservation of its coiled $\beta$-barrel fold, which is stabilized by an extensive disulfide bond network and contributes to the structural rigidity characteristic of VEGF-binding domains across neuropilin family members [106,107]. This mutual structural preservation distinguishes the NRP1 complex from the betaglycan system — in which the host protein undergoes progressive conformational maturation — and from the clusterin complex — in which the host chaperone samples a broad ensemble of conformational substates — positioning NRP1 alongside GPC1 and actin as a structurally rigid host partner that engages the viral surface without requiring large-scale backbone rearrangement.

##### 3.8.2 RMSF Asymmetry Reveals Ligand-Loop Dampening and Receptor b1-Pocket Immobilization

Per-residue RMSF analysis identifies a pronounced and mechanistically informative asymmetry between the envelope and NRP1 chains. In the envelope protein, the lowest fluctuation values are concentrated at a surface-exposed loop region that bears sequence and structural similarity to the CendR motif architecture — a carboxy-proximal segment presenting basic residues in a geometry compatible with b1-domain insertion — while the remainder of the envelope structure, including the domain I–II hinge and the fusion loop in domain II, retains its characteristic moderate flexibility. The dampening of this CendR-like loop is consistent with the steric constraint imposed by insertion into the NRP1 b1 binding pocket, which in VEGF-A engagement buries approximately 800 $Å^2$ of accessible surface area and forms a network of hydrogen bonds and electrostatic contacts that immobilize the terminal residues of the bound peptide [106,108]. In this framework, the RMSF reduction at the envelope loop represents direct structural evidence for pocket-mediated anchoring of a viral surface element that functionally mimics a CendR peptide ligand.

On the NRP1 side, the residues constituting the b1 binding pocket — in particular the conserved triad of acidic residues (Asp, Asp, Glu) that form the electrostatic cage coordinating the terminal arginine of CendR ligands [102,103] — display significantly reduced RMSF relative to the flanking loops and domain linkers. This pocket immobilization, observed in the presence of the docked envelope complex, mirrors the crystallographically documented rigidification of the NRP1 b1 pocket upon VEGF-A and CendR peptide binding, where the acidic cage adopts a closed conformation stabilized by direct contacts with the basic C-terminal residue of the ligand [107,108]. The selective dampening of the b1 pocket residues while maintaining flexibility in the b2 domain and the MAM domain linker indicates that the envelope–NRP1 interaction is spatially localized to the canonical ligand-recognition surface, rather than representing a non-specific global contact. Together, the envelope loop dampening and NRP1 pocket immobilization provide converging RMSF evidence for a productive, stereospecific engagement consistent with the CendR recognition mechanism.

##### 3.8.3 Contact Architecture Reflects Rapid Pocket Insertion Followed by Stable Engagement

The time evolution of the intermolecular contact number profile exhibits a characteristic two-phase behavior for the NRP1 complex. During the first approximately 30–40 ns, a rapid increase in contacts is observed, consistent with the initial docking-state conformation relaxing under simulation conditions to achieve tighter packing of the envelope loop within the NRP1 b1 pocket. After this initial compaction phase, the contact number stabilizes at a sustained plateau, without the progressive monotonic growth observed in the betaglycan complex or the continued oscillation seen in the BiP complex. The minimum interchain distance reaches a persistently low value — consistent with direct hydrogen bonding geometry — and does not exhibit transient increases indicative of partial dissociation, confirming that the envelope loop remains continuously engaged within the b1 pocket throughout the simulation.

The stabilization of the center-of-mass distance after the initial compaction phase indicates that no large-scale rigid-body rearrangement occurs after the pocket is occupied, consistent with the constrained geometry of the b1-domain insertion site, which physically limits the conformational freedom of the bound ligand. This behavior contrasts with the GPC1 complex, where the large surface area of the HSPG core protein permits continuous micro-reorientation within the DIII electropositive patch, and aligns more closely with the BiP complex, where a discrete electrostatic anchor dominates after initial engagement. The number of intramolecular interaction pairs (numP) for both chains remains stable throughout, confirming that neither partner undergoes internal structural disruption, and that the engagement geometry is maintained by the same set of contacts identified at the onset of the stabilization phase.

###### 3.8.4 MM/GBSA Energetics Identify a Mixed Electrostatic–van der Waals Interface with Attenuated Desolvation Penalty

The MM/GBSA binding free energy profile computed over the final 10 ns of the trajectory presents an energetic signature that distinguishes the NRP1 complex from both the highly charged HSPG-type interfaces (GPC1, betaglycan) and the purely hydrophobic chaperone-type contacts (clusterin). The gas-phase electrostatic contribution (EEL) is strongly favorable, with the primary stabilization arising from the interaction between the C-terminal basic residues of the envelope CendR-like loop and the acidic cage of the NRP1 b1 pocket, whose geometry is optimized for coordination of a terminal arginine guanidinium group. However, the absolute magnitude of EEL is lower than in the GPC1 complex, reflecting the more geometrically constrained nature of the interaction: rather than the broad, multivalent charge–charge landscape of HSPG engagement, the NRP1 contact concentrates electrostatic energy in a small number of high-specificity ion pairs.

The van der Waals contribution (VDWAALS) is comparatively more prominent in the NRP1 complex than in the GPC1 or BiP interfaces, reflecting the hydrophobic packing of the pocket-inserted loop residues against the aliphatic walls of the b1 binding site — a known structural feature of CendR–NRP1 complexes in which residues at the $-1$ and $-3$ positions relative to the terminal arginine pack against hydrophobic residues in the pocket entrance [103,108]. The polar solvation penalty (EGB) is consequently attenuated relative to GPC1, because the buried surface area comprises both polar (ion-pairing) and apolar (hydrophobic packing) components, with the apolar burial contributing favorably to ESURF and partially offsetting the polar desolvation cost. The resulting net $\Delta G_{\text{total}}$ is moderately negative and temporally stable, consistent with a specific, high-affinity interaction driven by the geometric complementarity of the b1 pocket rather than by non-specific electrostatic accumulation. The oscillations in $\Delta G_{\text{total}}$ are substantially smaller in amplitude than those observed in the GPC1 complex, reflecting the geometrically constrained nature of the b1-pocket engagement, which limits the configurational freedom of the interface and thereby stabilizes the instantaneous energy against the large fluctuations characteristic of broad electrostatic surfaces.

##### 3.8.5 Residue Decomposition Identifies the CendR-Like Contact Triad and b1-Pocket Acidic Cage as the Interaction Core

Per-residue free energy decomposition (SDC) reveals a compact and structurally interpretable hotspot architecture. On the envelope side, the dominant favorable contributors correspond to the terminal two to three residues of the CendR-like loop — specifically, the terminal arginine (or equivalent basic residue) that inserts into the b1 acidic cage, and the penultimate residue at the −1 position, which contributes van der Waals packing with the hydrophobic entrance of the pocket. The magnitude of the terminal arginine contribution approaches that of the ARG410 hotspot identified in the BiP complex, consistent with the well-established role of guanidinium–carboxylate ion pairs as dominant stabilizers of NRP1 b1 complexes across structurally diverse ligands [101,108]. On the NRP1 side, the three acidic cage residues — corresponding to the conserved Asp/Glu positions that coordinate the terminal arginine across all characterized CendR complexes [102,103] — each contribute individually favorable SDC values, generating a distributed but spatially concentrated receptor hotspot that collectively anchors the terminal loop of the viral ligand.

Pairwise decomposition (TDC) identifies the terminal Arg–Asp ion pair as the single most favorable interresidue contact, followed by contributions from the −1 position packing interactions and secondary hydrogen bonds between backbone amide donors in the loop and carbonyl acceptors in the pocket perimeter. The frustration loci — residues with positive SDC contributions of +8 to +12 kcal/mol — map to polar residues at the loop periphery that lose solvation energy upon pocket insertion without forming fully compensated contacts in the binding geometry sampled by the simulation. This distribution, in which a small core of three to four residues provides the dominant stabilization and a peripheral ring of two to three residues contributes measurable frustration, is quantitatively consistent with the energy decomposition profiles reported for peptide–NRP1 b1 complexes in independent MD studies [108] and highlights the energetic parsimony of CendR-type recognition.

##### 3.8.6 Conformational Ensemble Analysis: Pocket-Restricted Envelope Sampling and b1-Domain Rigidity

Principal component analysis and hierarchical RMSD clustering [92] reveal a conformational behavior for the NRP1 complex that is mechanistically consistent with the pocket-insertion model. The PC1–PC2 projection of the envelope chain shows a compact, low-dispersion distribution in which the dominant fraction of trajectory frames occupy a single conformational region, with a minor population of transitional frames in an adjacent area of phase space that stabilizes early in the simulation. The hierarchical dendrogram confirms a single dominant cluster comprising the majority of frames, with a small number of secondary clusters at short inter-cluster distances, reflecting the restricted conformational freedom imposed by the b1-pocket constraint on the envelope loop. The absence of a directed temporal arc — in contrast to the progressive conformational evolution observed in the GPC1 and betaglycan complexes — indicates that the envelope does not undergo large-scale accommodation after pocket engagement: the docking-state geometry is structurally adequate without requiring further global rearrangement.

The NRP1 chain exhibits a complementarily restricted conformational landscape. The PC1–PC2 projection is substantially less dispersed than that of GPC1 or clusterin, with frames concentrated in a compact conformational region whose temporal composition shows no systematic drift. The hierarchical dendrogram reflects a single major cluster with small inter-cluster linkage distances, consistent with the structural rigidity of the b1-domain fold and the known low intrinsic flexibility of the NRP1 ligand-binding domains in both crystal structures and prior MD analyses [106,107]. This mutual conformational restriction — low dispersion for both partners — is the defining conformational signature of the CendR recognition mechanism, in which the preorganized pocket geometry of NRP1 selects for a complementary peptide conformation without requiring either partner to undergo induced-fit rearrangement beyond local sidechain adjustments. The contrast with the GPC1 complex, where the GPC1 chain samples a broad multi-state ensemble while the envelope undergoes directed rearrangement, illustrates how different structural recognition mechanisms — multivalent electrostatic capture versus stereospecific pocket insertion — generate qualitatively distinct conformational signatures in 200 ns MD trajectories.

###### 3.8.7 Ternary Docking Reveals a Heterogeneous NRP1 Signalling Network with Plexin-Dominated Landscape Reorganisation

To evaluate how the NRP1 signalling neighbourhood modulates the dengue envelope–NRP1 interface, we performed conditional ternary docking simulations incorporating seven first-neighbour proteins drawn from the NRP1 interaction network: GIPC1, IGHV3-16, PLXNA1, PLXNA2, PLXNA4, SEMA4, and TGFBR1. For each candidate, docking energies were compared across the NRP1–Env reference pair (AB; $\Delta G \approx -12.8$ kcal $\cdot$ mol$^{-1}$), the Env–neighbour pair (BC; typically $-10$ to $-20$ kcal $\cdot$ mol$^{-1}$), and two conditional ternary assemblies, with energetic coupling quantified as $\Delta\Delta G$ values [41,51].

The most striking perturbation in the entire NRP1 panel was produced by PLXNA1. Under AB conditioning, AB shifted from $-12.8$ to $-25.6$ kcal $\cdot$ mol$^{-1}$ ($\Delta\Delta G \approx -12.8$ kcal $\cdot$ mol$^{-1}$) — a strong cooperative deepening of the NRP1–Env basin — while BC simultaneously became substantially less favourable ($\Delta\Delta G \approx +4.6$ kcal · mol$^{-1}$). This pattern does not correspond to simple additive stabilisation; it indicates a major reconfiguration of the conditional interaction landscape in which the NRP1–Env state is strongly reinforced while BC contacts are partially sequestered, consistent with the dominant native NRP1–Plexin receptor architecture reorganising around an Env-compatible configuration.

PLXNA4 produced the opposite directional effect and the strongest pure antagonism in the dataset. AB shifted from −12.8 to −6.8 kcal · mol$^{-1}$ under AB conditioning ($\Delta\Delta G \approx +6.0$ kcal · mol$^{-1}$), while BC remained nearly unchanged. The energetic vector is therefore dominated by selective AB displacement, indicating that PLXNA4 strongly competes with the NRP1–Env state without substantially perturbing the Env–neighbour contact. In contrast, PLXNA2 produced only minimal perturbations across all configurations ($|\Delta\Delta G| \leq 0.3$ kcal · mol$^{-1}$), establishing it as effectively permissive with respect to the NRP1–Env state.

SEMA4 produced the second-largest cooperative signature: AB strengthened from −12.8 to approximately −17.7 kcal · mol$^{-1}$ ($\Delta\Delta G \approx -4.9$ kcal · mol$^{-1}$) while BC remained essentially unchanged, yielding a selective AB-axis stabilisation with no BC disruption — the cleanest cooperative signature among the non-Plexin partners.

GIPC1 produced moderate, directional AB destabilisation: under AB conditioning, AB shifted from −12.8 to −10.7 kcal · mol$^{-1}$ ($\Delta\Delta G \approx +2.1$ kcal · mol$^{-1}$), while BC became slightly more favourable ($\Delta\Delta G \approx -0.3$ kcal · mol$^{-1}$); the reverse-conditioned branch produced a smaller AB perturbation ($\Delta\Delta G \approx +0.5$ kcal · mol$^{-1}$). IGHV3-16 showed a weak cooperative signal: AB shifted from −12.8 to −13.4 kcal · mol$^{-1}$ ($\Delta\Delta G \approx -0.6$ kcal · mol$^{-1}$) with only minor BC change ($\Delta\Delta G \approx +0.2$ kcal · mol$^{-1}$) and a negligible reverse-conditioned perturbation. TGFBR1 produced the smallest non-neutral effect: a modest AB destabilisation ($\Delta\Delta G \approx +0.8$ kcal · mol$^{-1}$) with BC essentially unchanged.

The overall energetic hierarchy for the NRP1 panel is therefore: PLXNA1 ($\Delta\Delta G_{AB} \approx -12.8$) and SEMA4 (−4.9) as cooperative stabilisers; PLXNA4 (+6.0) and GIPC1 (+2.1) as antagonists; TGFBR1 (+0.8) as a weak disruptor; IGHV3-16 (−0.6) as a marginal cooperator; and PLXNA2 ($\approx 0$) as permissive. These results reveal that the NRP1–Env energetic landscape is not uniformly robust but is instead strongly sensitive to the identity and assembly state of the surrounding NRP1 signalling complex.

#### 3.9 Envelope–Rab5 Complex: Endosomal Trafficking Recruitment Through a Two-Registry Directed Conformational Transition

##### 3.9.1 Global Structural Dynamics Reveal Progressive Interface Consolidation via Global Reorientation

Rab5 (RAB5A) is a small GTPase of the Ras superfamily that functions as a master regulator of early endosomal membrane dynamics, controlling vesicular docking, fusion, and cargo sorting through the recruitment of effector complexes including EEA1 and the PI3K complex VPS34–VPS15–Beclin-1 [109,110]. Experimental evidence has directly established that Rab5 activity is required for productive cellular entry of dengue and West Nile viruses, with dominant-negative Rab5 mutants blocking viral internalization at the early endosome stage [15,111]. The structural basis for this dependence — whether it reflects direct envelope–Rab5 contact or indirect modulation of endosomal membrane properties — remains uncharacterized at atomic resolution. The 200 ns MD trajectory of the envelope–Rab5 complex provides the first computational evidence for a plausible direct interaction mode and reveals a conformational mechanism that distinguishes this complex from all other envelope–host protein pairs examined in this study.

The global distance metric between the two chains (PAIR_1 center-of-mass distance) exhibits a distinctive biphasic behavior: an initial plateau near approximately 2.81 nm during the first 20 ns, followed by a gradual transition spanning approximately 20–120 ns that leads to a new stable late-time plateau near approximately 3.11 nm. The approximately 0.30 nm increase in the center-of-mass distance is not accompanied by a corresponding increase in the minimum interchain distance, which remains essentially constant at approximately 0.172 nm throughout the entire 200 ns trajectory. This dissociation between global and local distance metrics — one increasing while the other is invariant — is the defining structural signature of the Rab5 complex and indicates that the complex undergoes a **rigid-body global reorientation** rather than a partial dissociation: the two proteins rotate relative to one another while maintaining a continuously anchored local contact set [47]. Such reorientation behavior is mechanistically consistent with the known architecture of Rab–effector and Rab–cargo interactions, which frequently involve an initial encounter geometry that relaxes into a more favorable registered state through large-amplitude rotational rearrangement guided by the GTPase switch regions [110].

The energetic consequence of this reorientation is revealed by the contact number profile, which increases monotonically from approximately 3,600 contacts in the first 20 ns to approximately 5,100 contacts in the final 40 ns, reaching a more stable and less noisy plateau after approximately 120 ns. The combination of increasing contact number with increasing center-of-mass distance — apparently paradoxical — is resolved by the reorientation model: as the complex rotates, previously peripheral regions of both proteins come into closer proximity, generating new contacts at the redistributed interface, while the average distance between molecular centroids increases because the center of mass of each molecule is displaced from the contact zone. This interface consolidation pattern is distinct from the monotonic compaction observed in the betaglycan complex and from the rapid stabilization seen in the NRP1 complex, and constitutes a unique dynamic signature reflecting the structural logic of Rab-mediated protein–protein recognition.

##### 3.9.2 Asymmetric Conformational Accommodation: Envelope as the Structural Adaptor

RMSD analysis reveals a pronounced and biologically informative asymmetry between the two chains. Rab5 displays a modest and rapid conformational adjustment, with mean RMSD values reaching a first plateau near approximately 0.29 nm during the first 20 ns and stabilizing at approximately 0.39–0.40 nm from mid-trajectory onward, with constrained fluctuations consistent with a compact GTPase fold whose nucleotide-binding core and switch regions remain structurally coherent throughout. The dengue envelope protein, by contrast, exhibits substantially larger RMSD values — approximately 0.53 nm with wide dispersion during the first 20 ns, increasing to approximately 0.63–0.67 nm and only reaching low-variance stabilization after approximately 120 ns. The delayed stabilization of the envelope RMSD mirrors the timescale of the global reorientation event identified in the distance analysis, confirming that the conformational accommodation is primarily driven by rearrangements in the envelope rather than in Rab5. This asymmetry positions the envelope as the structurally plastic partner that absorbs the geometric cost of optimizing the interface — analogous to the role observed for the envelope in the GPC1 complex, but here reflecting larger-amplitude domain-level adjustments rather than local surface reorganization.

The radius of gyration provides complementary evidence for the direction of structural adaptation. The envelope undergoes a gradual decrease in Rg from approximately 3.14 nm during the first 20 ns to approximately 3.05 nm in the final 40 ns, indicating mild but sustained compaction of the ectodomain architecture as it converges toward the late-time interaction geometry. Rab5 maintains a globally stable Rg near approximately 1.65 nm throughout the trajectory, with a notable transient episode of enhanced compaction between approximately 120–160 ns, where the moving-average Rg decreases to approximately 1.63 nm. This compaction episode coincides precisely with the regime of highest contact number and lowest contact variance, suggesting that it reflects a brief period in which Rab5 is geometrically constrained by the interface contacts during the final stages of the reorientation event before settling into the dominant late-time registry.

Per-residue RMSF analysis localizes the structural flexibility to functionally interpretable regions of both proteins. In Rab5, the majority of the GTPase core — including the switch I and switch II regions that define GTP-state-dependent effector selectivity — maintains low backbone fluctuations consistent with a well-folded, dynamically constrained GTPase domain. The dominant high-RMSF region is confined to the C-terminal hypervariable domain (residues approximately 188–191), where RMSF values reach approximately 1.20 nm, dramatically exceeding all other positions in the protein. This extreme C-terminal flexibility is the biochemical signature of the Rab hypervariable domain (HVD), a functionally essential but intrinsically disordered segment that serves as a membrane-targeting and effector-modulation element and is known to remain structurally unresolved in the majority of Rab crystal structures due to its intrinsic conformational freedom [112,113]. The high RMSF of this region in the simulation is therefore expected and does not reflect interface-induced instability; rather, it reflects the constitutive disorder of the HVD that persists regardless of binding context.

In the envelope, the highest RMSF values are concentrated in two spatially distinct regions. The primary high-RMSF patch spans approximately residues 147–157, reaching values near 0.96 nm, and corresponds to a loop or hinge region within the central ectodomain segment that is known to participate in the domain I–domain II interface dynamics of flavivirus envelope proteins [5,114]. A secondary elevated-RMSF region near residue 471, reaching approximately 0.62 nm, is consistent with the stem-proximal portion of the ectodomain, which is known to undergo substantial positional variation in the context of pre-fusion to fusion state transitions [115]. The persistence of high flexibility in these two regions during the complex simulation indicates that, despite the global reorientation and interface consolidation, the envelope retains the intrinsic hinge dynamics at the DI–DII junction that are considered essential for its functional conformational repertoire.

##### 3.9.3 Persistent Anchoring and Late-Time Interface Consolidation Define a Two-Registry Binding Model

The combination of stable minimum distance, increasing contact number, converging RMSD, and decreasing envelope Rg across the 200 ns trajectory collectively define a coherent mechanistic model for the Rab5–envelope interaction: the complex initiates in an early contact registry determined by the initial docking geometry, undergoes a global reorientation spanning approximately 20–120 ns that redistributes interfacial contacts while maintaining local anchorage, and reaches a late-time registry characterized by higher contact density, lower structural variance, and more favorable overall geometry. This two-registry model is directly supported by the principal component analysis described in detail below.

The maintenance of a constant minimum distance of approximately 0.172 nm throughout both the early and late registries confirms that local anchor contacts are never broken during the reorientation — the complex “pivots” rather than “slides” into its late-time geometry, with the pivot axis likely corresponding to the highest-occupancy contact residues identified in the numP and RMSF analyses. This continuous anchoring behavior distinguishes the Rab5 complex from a simple two-step association mechanism and is mechanistically consistent with the known mode of GTPase–effector interaction, in which the GTPase switch regions establish an initial contact that subsequently recruits additional interface elements through conformational selection of the effector binding surface [116].

##### 3.9.4 MM/GBSA Energetics Reveal a vdW-Dominated Packing Scaffold with Electrostatic Microstate Modulation

End-state binding free energy calculations using the MM/GBSA formalism over the final 10 ns of the trajectory reveal an energetic signature that is mechanistically distinct from all other complexes analyzed in this study and that maps directly onto the two-registry structural model described above. The net binding free energy ($\Delta G_{\text{total}}$) is consistently negative and centered near approximately −40 kcal/mol across the majority of analyzed frames, with excursions to approximately −50 kcal/mol during intermediate frames and a notable transient deepening to approximately −60 to −70 kcal/mol during a brief episode that corresponds to the late-trajectory stabilization phase, followed by recovery toward a −50 to −60 kcal/mol late-time basin. This transient deepening event — whose moving-average signature indicates it is not high-frequency noise but a genuine visit to an alternative interfacial microstate — is the energetic manifestation of the contact-rich late-time registry identified in the structural analysis.

Decomposition into physical components reveals that two distinct mechanisms contribute to interface stability in a qualitatively different balance than seen in the GPC1 or NRP1 complexes. The van der Waals term (VDWAALS) is strongly favorable throughout, spanning approximately −95 to −115 kcal/mol with a gradual drift toward more negative values in the late portion of the trajectory, confirming that hydrophobic and shape-complementary packing constitutes a continuously improving structural scaffold as the interface consolidates toward its late-time registry [117]. The electrostatic contribution (EEL) is larger in absolute magnitude — approximately −200 kcal/mol on average — but substantially more volatile, with sharp excursions coinciding with the transient $\Delta G_{\text{total}}$ deepening event, identifying intermittent formation and reorganization of charge-assisted contacts (salt bridges, hydrogen bond clusters, charge–dipole networks) as the source of the largest instantaneous modulations in binding free energy [84]. The polar solvation penalty (EGB) is strongly unfavorable at approximately +250 to +280 kcal/mol, partially canceling the Coulombic stabilization, while the nonpolar surface term (ESURF) remains comparatively small ($\approx +10$ kcal/mol), yielding a net binding free energy that emerges from the balance between the large favorable gas-phase terms and the solvation costs.

This energetic architecture — a steady vdW-dominated packing scaffold modulated by a volatile electrostatic network — defines what may be termed a **dual-driver interface**: the vdW component provides thermodynamic continuity across both the early and late registries, while the electrostatic component generates the instantaneous energy differences that distinguish the two registries and produces the transient deepening event at the transition between them. This division of energetic labor is mechanistically appropriate for a GTPase–cargo interaction, in which the rigid GTPase core (stabilized by $Mg^{2+}$ coordination and the hydrophobic nucleotide base contacts) provides a geometrically well-defined surface that engages its partner primarily through shape complementarity and van der Waals packing, while the switch regions that define state-dependent partner specificity introduce charged residues (Thr, Asp, Glu in switch I and II) whose contacts are necessarily electrostatic in character [110,116].

###### 3.9.5 Residue Decomposition Identifies GLU133 as an Electrostatic Anchor and ARG91 as a Frustrated Peripheral Residue

Per-residue free energy decomposition (SDC) identifies a hotspot architecture consistent with the dual-driver interface model. On the envelope side (chain A), the dominant favorable contributor is **GLU133**, whose SDC contribution of approximately −6 kcal/mol is among the largest single-residue stabilizations observed across all host protein complexes in this study. The magnitude and sign of this contribution are consistent with an electrostatic anchor role — either a persistent salt bridge with a basic Rab5 residue or a focal point of a hydrogen bond network that achieves a favorable EEL/EGB balance by maintaining partial solvent accessibility while forming a stable charge-complementary contact. This anchoring character is consistent with the persistence of the minimum interchain distance throughout the reorientation event: GLU133 (or an equivalent acidic envelope residue at this position) likely corresponds to one of the “pivot anchor” contacts that remain continuously engaged while the global geometry reorganizes.

On the Rab5 side (chain B), the dominant unfavorable contributor is **ARG91**, with a positive SDC value approaching approximately +8 kcal/mol, identifying it as a frustrated peripheral residue whose desolvation penalty is not consistently compensated by favorable charge-pair contacts in the simulation-sampled binding geometries. ARG91 maps to the $ $3 helix region of Rab5, a surface segment that flanks the switch II domain and is known to exhibit variable engagement geometry across different Rab5–effector complexes [118,119]. The frustration at ARG91 is mechanistically interpretable as reflecting the incomplete optimization of the electrostatic network in the conformational ensemble sampled by the 200 ns simulation: the contact geometry that would allow ARG91 to form a full charge-compensated salt bridge with the opposing chain is transiently visited (producing the EEL excursion events) but not maintained as the dominant configuration.

Additional contributors identified in the SDC profile include stabilizing contacts from aromatic and polar residues on both chains (envelope: ARG57, GLN167, LYS210, TRP220; Rab5: PHE57, CYS63, LYS70, LEU85), indicating that the interface is distributed across a mixed hydrophobic–polar network rather than concentrated in a single hotspot, consistent with the multi-contact architecture revealed by the contact number analysis.

###### 3.9.6 Conformational Ensemble Analysis: Directed Two-Basin Transition and Metastable Substates

Principal component analysis of C$\alpha$ coordinates and hierarchical RMSD-based clustering [92] provide the clearest structural evidence for the two-registry model and distinguish the Rab5 complex as the only host protein system in this study exhibiting a genuine time-ordered directed conformational transition across the full 200 ns trajectory. The two-dimensional PC1–PC2 projection reveals a characteristic color gradient from dark purple (early frames, approximately 0–25 ns) occupying a well-separated high-PC1 region — including a distinct sub-lobe at low PC2 that corresponds to the initial contact registry — through an intermediate transitional population of cyan and green frames (approximately 25–150 ns) that bridges the two extremes, to a late-time basin of yellow frames (approximately 150–200 ns) concentrated at negative PC1 values. The strength of this temporal stratification — in which neither the early nor the late populations are revisited by frames from the opposite temporal regime — argues that PC1 captures a **directed slow mode** representing a genuine conformational relaxation rather than stochastic ergodic sampling around a fixed mean, consistent with the concept of essential dynamics in which the dominant principal components isolate functionally relevant collective coordinates [120,121].

Hierarchical clustering of the pairwise RMSD distance matrix confirms the two-superstate organization at the macroscopic level, with the highest-level bifurcation in the dendrogram occurring at approximately 0.6 RMSD units and separating the early-trajectory family from the late-trajectory family. Within each superstate, five to seven visually distinguishable macro-clusters are identifiable, corresponding to metastable substates within each registry — alternative hydrogen bond patterns, sidechain rotamer configurations, and local loop positions that are interconverted on a timescale faster than the registry-to-registry transition but slower than backbone vibrations. This hierarchical organization — substates within superstates within the full ensemble — is the conformational-landscape signature expected for a protein–protein interface undergoing slow directed relaxation with local exploration within each basin [93,100].

The Rab5 chain contributes relatively little to the variance captured by the first two principal components relative to the envelope, consistent with its role as the geometrically more rigid partner in the asymmetric accommodation model established by the RMSD and Rg analyses. The conformational contribution of Rab5 to the PC1 transition is largely mediated by the hypervariable C-terminal domain, which samples a wide conformational space independently of the registry state, and by modest repositioning of surface loops adjacent to the switch regions as they accommodate successive contact configurations on the evolving envelope surface.

##### 3.9.7 Ternary Docking Reveals Consistent Regulatory-Network-Driven Destabilisation of the Rab5–Env Interface

To evaluate how the Rab5 regulatory environment modulates the dengue envelope–Rab5 interface, we performed conditional ternary docking simulations incorporating five first-neighbour proteins drawn from curated Rab5 interaction networks: GDI2, PIK3C3, RAB5B, RAB5C, and RABGEF1. These partners collectively represent the principal functional classes of Rab5 regulatory machinery — GDP dissociation inhibitors, phosphoinositide kinases, paralogous GTPases, and guanine nucleotide exchange factors — providing a broad survey of the Rab5 regulatory landscape. For each candidate, docking energies were compared across the Rab5–Env reference pair (AB; $\Delta G \approx -15.9$ kcal · mol$^{-1}$), the Env–neighbour pair (BC), and two conditional ternary assemblies, with energetic coupling quantified as $\Delta\Delta G$ values [41,51].

In contrast to the mixed cooperative-antagonistic landscapes observed for receptor-centred systems such as NRP1 and Betaglycan, the Rab5 panels are dominated by a consistent and directional pattern: all five third partners destabilise the Rab5–Env reference state, with no partner producing net cooperative AB reinforcement. This uniformity is itself mechanistically informative, indicating that the Rab5–Env state is systematically incompatible with the native Rab5 regulatory network rather than selectively modulated by individual partners.

GDI2 produced the most globally disruptive profile in the dataset. Under AB conditioning, AB shifted from $-15.9$ to $-11.4$ kcal · mol$^{-1}$ ($\Delta\Delta G \approx +4.5$ kcal · mol$^{-1}$), while BC simultaneously weakened substantially ($\Delta\Delta G \approx +3.9$ kcal · mol$^{-1}$) — indicating a global deterioration of both interaction axes rather than selective AB displacement. Under reverse conditioning, the AB destabilisation was even stronger ($\Delta\Delta G \approx +4.8$ kcal · mol$^{-1}$), making GDI2 the only partner in the Rab5 panel where reverse conditioning amplified rather than attenuated the antagonistic effect.

RAB5B produced the most extreme asymmetry between conditioning branches. Under AB conditioning, AB shifted from $-15.9$ to $-11.7$ kcal · mol$^{-1}$ ($\Delta\Delta G \approx +4.2$ kcal · mol$^{-1}$), while under reverse conditioning the destabilisation became severe, collapsing AB to approximately $-5.6$ kcal · mol$^{-1}$ ($\Delta\Delta G \approx +10.3$ kcal · mol$^{-1}$) — the largest single conditional perturbation in the entire Rab5 panel. This extreme asymmetry indicates that a pre-formed Rab5–RAB5B assembly presents a surface highly incompatible with Env engagement, whereas introducing RAB5B into an established Rab5–Env complex produces a more moderate but still substantial disruption.

RAB5C showed a more symmetric profile: $\Delta\Delta G \approx +4.2$ kcal · mol$^{-1}$ under AB conditioning and +4.5 kcal · mol$^{-1}$ under reverse conditioning, with BC essentially unchanged ($\Delta\Delta G \approx -0.2$ kcal · mol$^{-1}$). PIK3C3 produced comparable AB destabilisation ($\Delta\Delta G \approx +4.2$ kcal · mol$^{-1}$) without detectable BC perturbation under reverse conditioning, indicating dominant AB-axis displacement. RABGEF1 generated a distinct two-axis signature: AB was destabilised by ≈ +4.2 kcal · mol$^{-1}$ under AB conditioning, while BC weakened dramatically ($\Delta\Delta G \approx +6.4$ kcal · mol$^{-1}$), and the reverse-conditioned branch produced only a mild AB stabilisation ($\Delta\Delta G \approx -0.5$ kcal · mol$^{-1}$).

The energetic ranking across partners in AB-conditioned AB destabilisation is: RAB5B (+4.2) ≈ RAB5C (+4.2) ≈ PIK3C3 (+4.2) ≈ RABGEF1 (+4.2) < GDI2 (+4.5), with RAB5B distinguished by its catastrophic reverse-conditioned AB collapse (+10.3). Despite sharing similar AB-conditioned magnitudes, the partners differ substantially in their BC perturbation profiles: GDI2 and RABGEF1 both disrupt BC, whereas RAB5C, PIK3C3, and RAB5B leave it largely intact, indicating mechanistically distinct modes of Rab5–Env displacement across functional partner classes.

#### 3.10 Envelope–SUMO9-UBC1 Complex: Post-Translational Machinery Hijacking Through Receptor-Driven Compaction and a Packing-Dominant Interface

##### 3.10.1 Asymmetric Structural Accommodation: A Rigid SUMO Core Against a Compacting Envelope Receptor

SUMO (Small Ubiquitin-like Modifier) conjugation is a reversible post-translational modification that controls nuclear transport, DNA damage response, transcriptional regulation, and protein stability through the sequential action of the E1 activating enzyme (SAE1/SAE2), the E2 conjugating enzyme Ubc9, and substrate-specific E3 ligases [122,123]. Ubc9 is the sole E2 enzyme in the SUMO pathway and occupies a central position in the SUMOylation cascade, serving as the direct covalent intermediate that transfers SUMO from E1 to substrate lysines, frequently assisted by SUMO-interacting motifs (SIMs) that recruit substrates to the E2 active site surface. Experimental evidence directly implicates the dengue envelope protein in this pathway: the DENV-2 E protein has been reported to interact with Ubc9, and specific E protein lysines (K51, K241) have been proposed as functional determinants of this interaction [124,125]. More broadly, SUMOylation has been demonstrated to be essential for dengue virus replication and transmission in mosquito models, and the SUMO pathway has been described both as a host antiviral defense mechanism and as a pathway co-opted by flaviviruses for productive infection [126,127]. The 200 ns MD trajectory of the envelope–SUMO9-UBC1 complex provides the first atomic-resolution structural characterization of this interaction and reveals a mechanistically distinctive binding mode driven by envelope-side conformational compaction and an energetic architecture dominated by van der Waals packing supplemented by localized electrostatic hotspots.

The most striking feature of the SUMO9-UBC1 trajectory is the pronounced asymmetry in conformational behavior between the two chains, which is the most extreme such asymmetry observed across all complexes examined in this study. The SUMO9-UBC1 ligand displays exceptional structural rigidity: RMSD values stabilize at approximately 0.22–0.24 nm within the first 20 ns and remain within this narrow band throughout the 200 ns trajectory, with falling variance in the second half of the simulation confirming convergence rather than continued drift. This behavior reflects the conserved $\beta$-grasp fold of SUMO proteins, whose hydrophobic core is stabilized by a network of buried side-chain contacts and disulfide constraints that resist thermal fluctuation even in the absence of binding partners [122]. The UBC1 E2 domain is similarly rigid, with its catalytic core maintaining structural integrity throughout the trajectory.

In stark contrast, the dengue envelope protein undergoes a large and temporally structured conformational rearrangement. RMSD values increase from approximately 0.50 nm during the first 20 ns to a plateau near approximately 0.73 nm that is reached after approximately 20–100 ns, followed by a distinct stabilization in which the variance of RMSD values decreases substantially during the final 100 ns. The transition from the high-variance early regime to the low-variance late regime is the structural signature of the envelope entering a well-defined conformational basin — a state that is mechanistically distinct from the initial docked geometry and corresponds to a more extensively accommodated interface configuration. This pattern of initial drift followed by late-stage stabilization, without return to the low-RMSD initial state, indicates that the envelope undergoes an irreversible conformational transition on the simulation timescale that constitutes a genuine conformational selection event rather than elastic fluctuation around the starting structure [128].

The radius of gyration provides the clearest evidence for the directionality of this adaptation. The SUMO9-UBC1 ligand maintains a constant Rg near approximately 1.66–1.67 nm throughout the entire 200 ns, confirming that its overall molecular shape does not change. The envelope protein, by contrast, undergoes a pronounced and sustained compaction: Rg decreases from approximately 3.11 nm during the first 20 ns to approximately 2.97 nm after the initial equilibration phase and remains at this lower value for the remainder of the simulation. This compaction of approximately 0.14 nm is the largest envelope Rg reduction observed across all host protein complexes in this study, and indicates that the three-domain ectodomain architecture of the envelope reorganizes its global geometry to accommodate the SUMO9-UBC1 surface, closing inter-domain degrees of freedom and adopting a more compact bound-state conformation. In the broader context of class II viral fusion proteins, domain rearrangements of this magnitude are associated with the formation of productive receptor contacts that constrain the hinge regions between domains I and II or between domain III and the stem-proximal segment [3,5].

##### 3.10.2 Interface Densification Proceeds in Three Kinetically Distinct Phases

The intermolecular contact number profile reveals that the SUMO9-UBC1 complex undergoes interface development through three kinetically identifiable phases that correspond directly to the structural transitions observed in the RMSD and Rg analyses. During the initial equilibration phase (approximately 0–20 ns), the contact number fluctuates around a mean of approximately 4,129 with a standard deviation of approximately 1,012, reflecting the high conformational heterogeneity of the initial docking geometry as the envelope explores the contact landscape adjacent to the docked configuration. During the intermediate phase (approximately 20–100 ns), a plateau is established at approximately 4,251 contacts with reduced variance (standard deviation approximately 458), corresponding to the period of sustained envelope RMSD increase in which the interface is being reshaped by large-scale envelope rearrangement while individual contacts are continuously broken and re-formed. During the late-time phase (approximately 100–200 ns), the contact number rises to approximately 4,622 with the lowest variance of the entire trajectory (standard deviation approximately 354), confirming that the envelope's conformational compaction produces a denser and more reproducibly maintained contact set — the structural correlate of the low-variance RMSD regime.

Throughout all three phases, the minimum interchain distance remains essentially constant at approximately 0.16–0.17 nm, confirming that the interface is continuously engaged without dissociation events. The center-of-mass distance decreases from approximately 4.47 nm during the early phase to approximately 4.30 nm during the late phase, with the variance in this metric decreasing from approximately 0.26 nm to approximately 0.13 nm across the same transition, indicating that the global geometry of the complex converges progressively toward a well-defined spatial relationship between the molecular centroids. This progressive narrowing of both the contact distribution and the center-of-mass distribution defines the SUMO complex as the most geometrically convergent interface observed in this study — a property that reflects the compact, modular architecture of the SUMO9-UBC1 surface and the capacity of the envelope to organize its flexible C-terminal region into a sustained contact geometry.

###### 3.10.3 RMSF Analysis Identifies the Envelope C-Terminal Region as the Primary Adaptive Interface Element

Per-residue RMSF analysis reveals that the structural adaptation is not distributed uniformly across the envelope surface but is concentrated in a well-defined region whose identity is functionally interpretable. The SUMO9-UBC1 ligand maintains low average RMSF near approximately 0.12 nm, with mobility localized to the N-terminal residues (residue 1: approximately 0.62 nm; residue 2: approximately 0.38 nm) and to several surface-exposed loop positions (residues approximately 104, 153, and 158). This flexibility profile is consistent with the structural organization of SUMO proteins and Ubc9, whose rigid $\beta$-grasp and UBC fold cores are decorated by flexible N-terminal extensions and surface loops that serve as primary interaction interfaces for substrate SIM recognition [122,123]. The concentration of ligand flexibility in peripheral loop and terminal positions — rather than in the core — is mechanistically appropriate for a regulatory enzyme that must maintain active-site geometry while accommodating diverse substrate interfaces.

The envelope protein exhibits substantially higher RMSF values (average approximately 0.28 nm), with the highest-flexibility peak concentrated in the C-terminal region spanning approximately residues 464–471, where RMSF reaches approximately 0.83 nm. This C-terminal region corresponds to the stem-proximal and potentially transmembrane-proximal segment of the envelope ectodomain — a region known to undergo conformational changes during pre-fusion to post-fusion transitions and to participate in dimeric rearrangements during viral assembly [3,115]. The coincidence of the highest RMSF in this region with the highest contact-gain ratio (residues 461–466 and 492–495 reaching contact-gain ratios of approximately 1.275) is the defining structural feature of the SUMO complex and identifies the C-terminal stem-proximal region as the primary adaptive interface element: the region that is most flexible in the unbound-like early-trajectory state is precisely the region that develops the most new contacts as the complex consolidates in the late-time phase. This "flexible arm" model for envelope–SUMO9-UBC1 engagement contrasts sharply with the DIII electropositive patch model for GPC1 and the CendR-loop insertion model for NRP1, and positions the envelope stem-proximal region as a previously uncharacterized host-factor binding surface with potential functional significance for the SUMOylation axis during viral infection.

On the ligand side, the contact-gain analysis identifies N-terminal residues (1–2) and loop residues (33–35) as the primary contributors to late-time interface densification, with contact-gain ratios reaching approximately 1.098. These positions correspond to the flexible N-terminal extension of SUMO proteins and to the loop adjacent to the SIM-binding groove on the UBC1 surface [123]. Their participation in the evolving interface indicates that the SUMO9-UBC1 uses its peripheral flexible elements — rather than its rigid core — to extend the contact network onto the envelope surface, consistent with a recognition model in which the E2 enzyme adapts its substrate-recruiting surface to accommodate a non-canonical binding partner without disrupting the core active-site geometry.

###### 3.10.4 MM/GBSA Energetics Define a vdW-Dominant Packing Interface with Compensatory Electrostatic Hotspots

End-state binding free energy analysis over the final 10 ns of the trajectory reveals an energetic profile that distinguishes the SUMO complex from all other host protein interfaces examined in this study. The net $\Delta G_{\text{total}}$ is centered near approximately −60 kcal/mol, oscillating within approximately −75 to −45 kcal/mol across the analyzed frames, with mild relaxation toward less favorable values in certain late windows consistent with the dynamic reorganization of the interface microstate rather than with weakening of the overall complex. The decomposition identifies van der Waals packing (VDWAALS approximately −85 kcal/mol) as the single largest favorable contributor — larger in relative proportion than in the GPC1, NRP1, or BiP interfaces — confirming that the contact-dense late-time geometry established by envelope compaction generates a steric packing core whose contribution to binding stability exceeds the contribution of electrostatic contacts. This VDW dominance is mechanistically consistent with the structural observation that the C-terminal stem-proximal region of the envelope, which drives the late-time contact densification, presents primarily hydrophobic and aromatic side chains that pack against the aliphatic surface of the SUMO $\beta$-grasp fold and the UBC1 active-site flank.

The electrostatic contribution (EEL approximately −180 kcal/mol) is strongly favorable but exhibits higher temporal variability than the VDWAALS term, consistent with the dynamic salt-bridge and hydrogen-bond network at the periphery of the packing core. The polar solvation penalty (EGB approximately +220 kcal/mol) partially cancels the EEL gain, yielding the characteristic MM/GBSA signature of a charged interface: large attractive and repulsive terms whose difference defines the net binding free energy, and whose ratio determines sensitivity to perturbations in charge geometry or hydration state [49,84]. The non-polar surface term (ESURF approximately −10 to −15 kcal/mol) provides a modest but consistent favorable contribution from hydrophobic surface burial, complementing the dominant VDWAALS term in characterizing the interface as packing-dominated with electrostatic reinforcement rather than the electrostatic-dominated profile of GPC1 or the geometrically constrained profile of NRP1.

###### 3.10.5 Residue Decomposition Identifies a Compact ENV Triad and Dual Salt-Bridge Architecture

Per-residue free energy decomposition (SDC) identifies the energetic architecture of the interface with residue-level resolution. On the envelope side, the dominant favorable contributors form a spatially compact triad centered on **GLU85**, **ARG94**, and **PHE96**, whose combined contributions constitute the core electrostatic and packing hotspot of the interface. GLU85 and ARG94 contribute primarily through charge-pair interactions with complementary residues on the SUMO9-UBC1 surface, while PHE96 contributes primarily through van der Waals packing — its aromatic side chain inserting into a hydrophobic depression on the E2 surface that provides shape-complementary contact with minimal desolvation cost. This three-residue architecture in which two charged residues flank an aromatic packing residue is characteristic of dual-mechanism hotspots that achieve both electrostatic specificity and geometric stability simultaneously, and has been documented as a recurring motif in viral surface protein–host enzyme interfaces [43,129].

On the SUMO9-UBC1 side, **GLU42** is identified as the primary stabilizing contributor (favorable SDC), consistent with its role as a salt-bridge partner for ARG94 on the envelope. **LYS30** and **LYS59** exhibit unfavorable SDC contributions, identifying them as frustrated peripheral residues that incur desolvation penalties without forming consistently compensated charge-pair contacts in the simulation-sampled ensemble. **SER158** similarly shows a significant positive SDC contribution, reflecting the high desolvation cost of a polar hydroxyl group that is partially buried at the interface periphery without forming a stable hydrogen-bond partner. These frustration loci at LYS30, LYS59, and SER158 identify positions where the current late-time binding geometry is not fully optimized and where further conformational refinement — or, in the physiological context, modulation by SUMO chain geometry or substrate co-occupancy — could reduce the desolvation penalty and strengthen the interaction.

Pairwise decomposition (TDC) confirms that the dominant inter-chain contributions correspond to two specific electrostatic pairs: **GLU85(envelope)–LYS59(SUMO9-UBC1)** and **ARG94(envelope)–GLU42(SUMO9-UBC1)**. These ion pairs appear as the deepest-blue elements in the TDC heatmap, with pairwise energies consistent with direct salt-bridge geometry. The identification of exactly two dominant pairwise electrostatic contacts is mechanistically significant: it mirrors the "few strong pairs plus broader packing" architecture documented for canonical Ubc9–substrate interactions, where a small number of charge-complementary contacts provide recognition specificity while an extended hydrophobic interface provides binding persistence [123].

###### 3.10.6 Conformational Ensemble Analysis: Envelope State Transition and SUMO Substate Selection

Principal component analysis of C$\alpha$ coordinates [92] and hierarchical RMSD clustering reveal a conformational landscape for the SUMO complex that is qualitatively distinct from all other systems in this study and directly maps onto the three-phase interface development described above. The two-dimensional PC1–PC2 projection of the envelope chain shows a color-stratified temporal progression in which early frames (dark purple, approximately 0–25 ns) occupy a broad basin at negative PC1, intermediate frames (cyan–green, approximately 25–100 ns) traverse a transitional region of PC space, and late frames (yellow, approximately 100–200 ns) accumulate in a distinct narrow manifold at positive PC1 with elevated PC2 values. The directional character of this progression — with no backward revisitation of the early basin after the transition is complete — identifies PC1 as a **directed slow collective mode** that captures the conformational compaction and C-terminal reorganization of the envelope rather than ergodic diffusion around a single attractor [120]. Hierarchical clustering of the envelope trajectory resolves three major cluster families in the dendrogram, whose temporal composition corresponds precisely to the three kinetic phases identified in the contact analysis: an early-trajectory family (approximately 0–20 ns), an intermediate transition family (approximately 20–100 ns), and a late-time dominant family (approximately 100–200 ns) that comprises the majority of frames and is internally coherent at a low inter-cluster RMSD distance.

The SUMO9-UBC1 chain exhibits a qualitatively different conformational landscape that is complementary to the envelope transition. The PC1–PC2 projection for SUMO9-UBC1 shows multiple locally populated clusters in the early trajectory, reflecting the substate sampling of peripheral loops and the N-terminal extension as the ligand explores compatible contact configurations on the evolving envelope surface. As the simulation progresses, these early clusters gradually depopulate and frames accumulate in a single dominant late-time basin at positive PC1 with narrow dispersion — the conformational selection of the ligand substate most compatible with the envelope's late-time compact geometry. This temporal convergence of both partners toward narrow, distinct late-time basins is the conformational signature of a cooperative induced-fit mechanism in which neither partner independently achieves its binding geometry: the envelope drives the global transition through compaction, while SUMO9-UBC1 filters its local substate diversity to select the configuration that maximizes contact with the evolved envelope surface. This coordinated convergence distinguishes the SUMO complex from the Rab5 complex — where the envelope undergoes a directed pivot while Rab5 remains broadly distributed — and from the NRP1 complex — where both partners show restricted conformational sampling from the outset — establishing a mechanistically unique binding model appropriate to the regulatory function of the SUMO conjugation machinery.

###### 3.10.7 Ternary Docking Reveals the SUMO1–Env State as a Shallow Basin Systematically Superseded by Canonical SUMO Pathway Assemblies

To evaluate how the SUMO1 signalling and enzymatic network modulates the dengue envelope–SUMO1 reference interaction, we performed conditional ternary docking simulations incorporating seven first-neighbour proteins: IPO13, RANGAP1, SAE1, UBA2, SUMO1 (self), SUMO2, and SUMO3. These partners span the principal functional compartments of the SUMOylation machinery — nuclear transport (IPO13), canonical SUMO1-modified substrate (RANGAP1), E1 activation enzyme subunits (SAE1 and UBA2), and SUMO paralog family members (SUMO1, SUMO2, SUMO3) — providing a systematic survey of the pathway [122,130,131]. Energetic coupling was quantified as $\Delta\Delta G$ values relative to the SUMO1–Env reference pair (AB; $\Delta G \approx -3.4$ kcal · mol$^{-1}$) [41,51].

The SUMO1–Env reference state is notably weaker than the AB baselines of all other receptor-centred systems in this study ($-12.8$ to $-15.9$ kcal · mol$^{-1}$). This intrinsically shallow baseline is mechanistically significant: rather than representing a dominant attractor, the SUMO1–Env configuration constitutes a labile energetic state susceptible to large-magnitude remodelling by pathway-relevant partners. Consistent with this, all seven third partners produce strong AB stabilisation ($\Delta\Delta G$ ranging from $-1.5$ to $-9.9$ kcal · mol$^{-1}$), indicating that the canonical SUMO assembly logic consistently and decisively outcompetes the weak reference Env-associated basin.

IPO13 produced the most asymmetric landscape in the dataset. Under AB conditioning, AB shifted from $-3.4$ to $-13.3$ kcal · mol$^{-1}$ ($\Delta\Delta G \approx -9.9$ kcal · mol$^{-1}$), while BC simultaneously weakened from $-13.4$ to $-6.2$ kcal · mol$^{-1}$ ($\Delta\Delta G \approx +7.2$) — indicating a major energetic reallocation toward AB at the expense of BC. In the reciprocal branch, AB remained more favourable than reference but with a reduced gain ($\Delta\Delta G \approx -1.5$ kcal · mol-1). This asymmetric pattern is consistent with the role of IPO13 as a bidirectional karyopherin that organises the nuclear transport arm of the SUMO pathway by importing UBC9 [132,133].

RANGAP1 exhibited a clean, selective cooperative signature: AB shifted from −3.4 to −12.8 kcal · mol-1 ($\Delta\Delta G \approx -9.4$) under AB conditioning without a detectable informative reciprocal state. This result is mechanistically notable because RANGAP1 is the canonical SUMO1-modified substrate at the nuclear pore, where SUMO1 conjugation targets it to a stable complex with RanBP2/Nup358 and UBC9 — one of the most structurally well-defined SUMO1-dependent assemblies in the cell [134–136].

SAE1 and UBA2 produced broadly concordant cooperative signatures. SAE1 stabilised AB from −3.4 to −12.8 kcal · mol-1 ($\Delta\Delta G \approx -9.4$) under AB conditioning, with maintained AB stabilisation in the reciprocal branch ($\Delta\Delta G \approx -1.7$). UBA2 produced an equivalent AB-conditioned stabilisation ($\Delta\Delta G \approx -9.4$; AB reaching −12.8 kcal · mol-1) with moderate BC weakening ($\Delta\Delta G \approx +2.5$) and a stronger reciprocal branch AB gain ($\Delta\Delta G \approx -4.1$). The convergence of both E1 subunits on nearly identical cooperative magnitudes is consistent with their functioning as the obligate heterodimeric SUMO-activating enzyme that catalyses SUMO adenylation and thioester transfer with high paralog specificity [122,131].

Among the paralog comparisons, SUMO1 self-association yielded the strongest combined cooperative signal: AB stabilised from −3.4 to approximately −13.0 kcal · mol-1 ($\Delta\Delta G \approx -9.6$) under AB conditioning with BC weakening ($\Delta\Delta G \approx +2.7$); the reciprocal branch retained substantial AB stabilisation ($\Delta\Delta G \approx -5.3$). SUMO2 and SUMO3 showed equivalent AB-conditioned gains ($\Delta\Delta G \approx -9.7$ and −9.6, respectively), with SUMO3 additionally showing pronounced BC weakening under AB conditioning ($\Delta\Delta G \approx +6.1$). The parallel behaviour of SUMO2 and SUMO3 — which are highly similar to one another but biochemically distinct from SUMO1 in chain formation and paralog-biased substrate recognition — reinforces the conclusion that SUMO-centred assemblies in general, regardless of paralog identity, dominate the landscape over the reference SUMO1–Env state [137,138].

The dominant global signal across all seven partners is therefore a consistent, large-magnitude collapse of the landscape away from the shallow SUMO1–Env basin and toward canonical SUMO-pathway configurations. The main axis of variation among partners lies not in AB stabilisation magnitude (which is strikingly uniform at ≈ −9.4 to −9.9 kcal · mol-1) but in BC perturbation profile: IPO13 and SUMO3 strongly weaken BC; UBA2 and SUMO1 produce moderate BC weakening; RANGAP1 and SUMO2 primarily register as highly selective AB cooperators without strong BC displacement. This variation reflects the mechanistic diversity of SUMO1 network partners but does not alter the fundamental conclusion that the Env-associated SUMO1 state is energetically outcompeted across the entire partner set.

#### 3.11 Envelope–TGFBR1 Complex: TGF- Signalling Receptor Engagement Through Asymmetric Conformational Exploration and a High-Density Contact Interface

##### 3.11.1 Continuous Envelope Drift Against a Structurally Conservative Kinase Receptor

Transforming growth factor-$\beta$ receptor type 1 (TGFBR1, also known as ALK5) is a transmembrane serine/threonine kinase that forms obligate heteromeric complexes with TGFBR2 upon ligand binding, transducing TGF-$\beta$, activin, and BMP signals through cytoplasmic phosphorylation of SMAD2/3 transcription factors [139,140]. As a receptor with both extracellular ligand-binding and intracellular kinase domains, TGFBR1 is structurally organized to maintain a rigid signalling-competent fold while permitting regulated conformational transitions associated with GS-domain activation and heterodimer assembly [141]. At the intersection of viral biology and host signalling, TGF-$\beta$ pathway components have been implicated in dengue pathogenesis: TGF-$ $1 serum levels are elevated in severe dengue and correlate with plasma leakage [142], and TGFBR1-dependent signalling has been proposed to modulate endothelial barrier integrity during dengue infection, making TGFBR1 a functionally relevant candidate for envelope-mediated host factor engagement. The 200 ns MD trajectory of the envelope–TGFBR1 complex reveals a mechanistically distinctive binding dynamics in which the envelope protein serves as a continuous conformational explorer against a structurally stable kinase receptor surface, generating the highest interfacial contact density observed across all complexes in this study.

The asymmetry between the two chains in the RMSD analysis is among the most pronounced observed in any complex examined here, and is qualitatively different from the asymmetry seen in either the Rab5 or SUMO complexes. TGFBR1 (ligand) maintains comparatively stable RMSD values throughout the 200 ns trajectory, rising gradually from a mean of approximately 0.180 nm during the first 50 ns to approximately 0.224 nm during the final 50 ns, with tight standard deviations of approximately 0.01–0.02 nm in each window. This trajectory reflects a protein whose global fold is preserved throughout the simulation, with only a modest late-phase drift consistent with relaxation of crystal-contact constraints and thermal equilibration of surface loops. The dengue envelope protein, by contrast, displays a sustained, monotonic increase in RMSD across all four temporal windows: approximately 0.381 nm (0–50 ns), 0.490 nm (50–100 ns), 0.533 nm (100–150 ns), and 0.582 nm (150–200 ns). This continuous drift without reaching a stable plateau is the defining feature of the TGFBR1 complex and distinguishes it categorically from every other system in this study: unlike the SUMO complex (where envelope RMSD stabilizes after ~100 ns), the Rab5 complex (where envelope undergoes a discrete pivot then stabilizes), or the NRP1 complex (where both partners are conformationally restricted), the TGFBR1 complex envelope does not reach a single dominant conformational basin within the 200 ns simulation window. This continuous drift indicates that the envelope protein explores a broad and heterogeneous conformational landscape on the surface of TGFBR1 rather than converging toward a single optimal binding geometry, a behavior consistent with the essential dynamics of flavivirus envelope proteins, whose structural polymorphism supports multi-stage receptor engagement [5,143].

The radius of gyration provides complementary evidence for the direction of conformational adaptation. TGFBR1 undergoes only a subtle Rg change, from approximately 1.972 nm during the first 50 ns to approximately 1.961 nm in the final window, confirming preservation of its compact kinase domain architecture. The envelope protein shows a more substantial compaction from approximately 3.113 nm to approximately 3.049 nm, a reduction of 0.064 nm that, combined with the continuously rising RMSD, indicates progressive repackaging of the ectodomain architecture: the envelope simultaneously becomes more compact (lower Rg) and more divergent from its initial structure (higher RMSD), consistent with inter-domain rearrangements that redistribute internal contacts without expanding the molecular envelope [115]. This Rg decrease at rising RMSD is, among all complexes analyzed, most pronounced in the TGFBR1 system and most consistent with a continuous multi-state exploration mechanism.

###### 3.11.2 Progressive Interface Densification: The Highest Contact Count in the Host Network

Despite the continuous structural drift of the envelope protein, the interfacial contact metrics reveal that the TGFBR1 complex establishes and progressively consolidates the largest and most contact-dense interface observed across all host protein complexes in this study. The number of interchain contacts within 0.6 nm increases in a nearly monotonic fashion across all four temporal windows: approximately 6,883 contacts (0–50 ns), approximately 7,971 contacts (50–100 ns), approximately 8,714 contacts (100–150 ns), and approximately 9,047 contacts (150–200 ns). This late-time contact count — more than 9,000 simultaneous contacts — is substantially larger than the corresponding values observed in all other complexes and is achieved despite the global center-of-mass distance (PAIR_1) remaining stable near approximately 3.0 nm after an initial slight increase from approximately 2.93 nm (0–50 ns) to approximately 3.04 nm (50–100 ns). The combination of a stable or mildly increasing center-of-mass distance with a steadily growing contact number is the structural signature of a **sliding and rolling interface consolidation** mechanism: rather than approaching each other globally, the two proteins rotate and translate laterally on their shared surface, progressively recruiting new atomic contact pairs from previously non-interacting regions while maintaining a constant proximal separation [47].

The minimum interchain distance confirms that this consolidation proceeds without dissociation: it remains essentially invariant at approximately 0.163–0.166 nm throughout all 200 ns, indicating continuous atomic-level engagement at a close-packing distance consistent with persistent van der Waals contacts and potential hydrogen bonds. The simultaneous stability of the minimum distance and the monotonic growth of the total contact count establish that the TGFBR1 interface operates in a regime of **expanding contact redistribution**: the closest contact (minimum distance anchor) is maintained while progressively more peripheral regions of both proteins come into detectable proximity, generating a contact network that grows in breadth rather than depth. This behavior is mechanistically appropriate for a large, multi-domain complex in which the envelope ectodomain (three structured domains plus flexible stem) presents a broad and conformationally variable surface to the globular kinase ectodomain of TGFBR1.

###### 3.11.3 RMSF Hotspot Topology: Dynamic Envelope Regions Drive Contact Gain

Per-residue RMSF analysis reveals the most extreme flexibility contrast between the two chains observed in any complex in this study. TGFBR1 maintains a low average RMSF of approximately 0.112 nm (median approximately 0.093 nm; 95th percentile approximately 0.252 nm), with the highest-mobility residues localized to the C-terminal tail segment (residues approximately 497–498, reaching approximately 0.58 nm) and to an intermediate surface loop region spanning approximately residues 270–324 (RMSF approximately 0.34–0.40 nm). The C-terminal peak is consistent with the flexible regulatory tail present in the TGFBR1 ectodomain construct, while the elevated mobility of residues 270–324 corresponds to a surface-exposed segment adjacent to the extracellular domain architecture that is known to participate in TGFBR2 heterodimerization contacts in the canonical TGF-$\beta$ receptor complex [144]. The moderate loop flexibility at this position is therefore both expected and functionally interpretable as the ligand-accessible surface that, in the canonical signalling context, engages the heterodimeric receptor partner.

The dengue envelope protein exhibits dramatically higher backbone dynamics (average RMSF approximately 0.266 nm; median approximately 0.244 nm; 95th percentile approximately 0.471 nm), with three distinct high-RMSF peaks whose positions are mechanistically informative. The largest individual peak at residue approximately 101 (RMSF approximately 0.670 nm) corresponds to a surface-exposed loop within the domain I–domain II interface region of the envelope ectodomain, a position known to participate in the hinge motions that couple domain I repositioning to the conformational changes required for membrane fusion [114]. Secondary high-RMSF regions at residues approximately 449–450 and approximately 495 (RMSF greater than 0.620 nm) are consistent with the stem-proximal and possibly transmembrane-proximal segment of the ectodomain, whose conformational freedom reflects the absence of bilayer constraints in the simulation system and whose functional relevance for post-fusion trimer formation has been established structurally

[143].

The contact-gain analysis (mdmat ratio) maps directly onto the RMSF topology and provides the most precise structural localization of the evolving interface. On the envelope side, the highest contact-gain ratios are concentrated at residues approximately 448–452 (gain ratio approximately 1.391), with secondary signals at residues approximately 266–273 and at positions approximately 47 and 149–150. The coincidence of the strongest contact gains (residues 448–452) with the highest RMSF values (residues 449–450 and 495) confirms that the stem-proximal high-flexibility region is the primary "adaptive arm" of the envelope–TGFBR1 interface, contributing new contacts as the envelope explores successive conformational states on the kinase surface. On the TGFBR1 side, the highest contact-gain residues are located at positions approximately 35–40 (ratio approximately 1.207), with additional contributions at residues approximately 121–123, 163, 171, and 297–298. The residues at positions 35–40 and 121–123 correspond to segments within or adjacent to the extracellular domain of TGFBR1 that are positioned to receive contact from an approach by the envelope stem-proximal region, while position 297–298 overlaps with the flexible surface loop region identified as high-RMSF on the TGFBR1 chain.

###### 3.11.4 MM/GBSA Energetics: GGAS-Dominant Interface with Progressive Regime Drift

End-state binding free energy calculations reveal an energetic profile unique to the TGFBR1 complex among all systems studied: large absolute values with a clear temporal drift that tracks the structural reorganization of the envelope. The net $\Delta G_{\text{total}}$ is centered near approximately $-80$ kcal/mol as a trajectory-wide mean, but this average obscures a systematic temporal trend: early frames produce values near approximately $-90$ to $-95$ kcal/mol, while late frames converge toward approximately $-70$ to $-75$ kcal/mol, with the moving-average profile confirming this is a genuine monotonic relaxation rather than high-frequency noise. This temporal drift — $\Delta G_{\text{total}}$ becoming less favorable as the trajectory progresses while contact count simultaneously increases — is the most counterintuitive energetic signature in the dataset and requires mechanistic interpretation beyond a simple affinity statement.

The decomposition clarifies the origin of this paradox. The gas-phase interaction energy (GGAS, composed of VDWAALS plus EEL) reaches approximately $-280$ kcal/mol — the largest GGAS value in the study, commensurate with the largest contact interface and consistent with MM/GBSA's well-documented overestimation of gas-phase terms for highly charged, solvent-exposed interfaces [49,50]; this value should therefore be interpreted as a structural indicator of interface extent and charge density rather than as a physically measurable affinity — but the VDWAALS component specifically shows a temporal drift from approximately $-200$ kcal/mol in the early trajectory to approximately $-175$ kcal/mol in the late trajectory.

The polar solvation penalty (EGB, approximately +200 kcal/mol) remains relatively stable, meaning the drift in $\Delta G_{\text{total}}$ primarily reflects the weakening of the VDWAALS term. This pattern has a specific physical interpretation: during the sliding and rolling interface consolidation, the envelope progressively recruits new contacts in peripheral surface regions, but these late-appearing contacts are geometrically less optimal for van der Waals packing than the initial close-approach contacts that form early in the trajectory. The interface expands in area (more contacts) but simultaneously becomes geometrically less optimal in packing quality (lower VDWAALS per unit area) — a direct consequence of the multi-macrostate envelope exploration, where later-occupied basins may present surface geometries that are complementary to TGFBR1 in terms of contact number but suboptimal in steric packing energy [49]. The electrostatic contribution (EEL) is also strongly favorable within GGAS, providing complementarity between the charged surface elements of the viral ectodomain and the lysine/aspartate hotspot cluster of TGFBR1 identified in the residue decomposition.

###### 3.11.5 Residue Decomposition Identifies Electrostatic TGFBR1 Anchor Residues Against a Distributed Envelope Surface

Per-residue free energy decomposition (SDC) defines an energetic hotspot architecture that is structurally distinct from all previous complexes and mechanistically coherent with the multi-macrostate envelope exploration model. On the TGFBR1 side, the dominant favorable contributors form a charged triad: **LYS296** provides the single largest per-residue stabilization in the complex, contributing approximately −8 to −9 kcal/mol, while **ASP422** and **LYS337** contribute additional strongly favorable terms. This basic–acidic arrangement on the TGFBR1 surface constitutes an electrostatic scaffold that is consistently engaged across the diverse envelope conformational states sampled during the trajectory — the invariance of this scaffold explains why the minimum interchain distance remains stable throughout the envelope's continuous structural drift. **PRO454** presents a positive (unfavorable) SDC contribution, identifying it as a frustrated residue whose rigid pyrrolidine backbone geometry imposes a desolvation cost without enabling compensatory charge-pair contacts — a characteristic of proline residues at protein–protein interfaces that are geometrically constrained but energetically suboptimal [43,129].

On the envelope side, the SDC profile does not concentrate in a single compact triad as observed in the SUMO or NRP1 complexes, but rather distributes moderate favorable contributions across a broader set of residues consistent with the large and evolving contact interface. This distribution is reflected in the temporal behavior of the SDC and TDC traces: individual per-residue contributions become more favorable in late trajectory frames (with TDC reaching approximately −35 to −45 kcal/mol at the deepest pairwise contacts in the late window), indicating that as the interface reorganizes, the dominant electrostatic hotspots on TGFBR1 progressively achieve better geometric complementarity with their envelope counterparts even as the global VDWAALS term relaxes. This divergence — hotspot pairs strengthening while overall packing energy weakens — is the energetic fingerprint of the sliding/rolling interface consolidation mechanism: specific anchor contacts become more precisely registered as the envelope explores different macrostates, while the net packing contribution decreases because peripheral contacts are geometrically inferior.

###### 3.11.6 Conformational Landscape Analysis: Multi-Macrostate Envelope Exploration Against a Single-Basin Kinase

Principal component analysis [92,120] and hierarchical RMSD clustering produce the most asymmetric conformational landscape among all complexes in this study and provide the clearest mechanistic explanation for the time-dependent energetic behavior described above. The PC1–PC2 projection of the envelope chain displays a pronounced multi-basin architecture: early frames (dark purple, approximately 0–25 ns) occupy a distinct initial region in PC space; intermediate frames (cyan–green, approximately 25–100 ns) traverse a broad transitional corridor; and late frames (yellow, approximately 150–200 ns) accumulate across a distributed late-time manifold without converging to a narrow basin. Critically, unlike the directed single-transition observed in the GPC1 complex or the two-basin transition observed in the Rab5 complex, the envelope PCA in the TGFBR1 system shows **no late-time convergence**: the yellow frames are spread across multiple PC1–PC2 positions rather than accumulating in a compact basin, confirming that the 200 ns simulation does not capture a complete relaxation to a single dominant conformational state. Inter-cluster distances in the dendrogram reach approximately 0.45 RMSD units at the highest bifurcation level, establishing the presence of structurally distinct macrostates separated by barriers that are not fully overcome within the simulation timescale.

TGFBR1 presents the complementary extreme: the dendrogram resolves numerous small clusters all merging at low linkage distances (maximum approximately 0.15 RMSD units), and the PC1–PC2 projection shows a dense, largely overlapping cloud with only a modest temporal gradient. This compact, low-dispersion conformational landscape confirms that TGFBR1 samples a single dominant conformational basin with microstate-level variation — consistent with the low RMSD values and tight standard deviations observed throughout the trajectory. The extreme contrast between the envelope's multi-macrostate landscape and TGFBR1's single-basin microstate topology is the defining conformational signature of the TGFBR1 complex and establishes the mechanistic principle underlying all other observed properties: the envelope provides conformational polymorphism that enables progressive surface exploration, while TGFBR1 provides a structurally invariant counter-surface whose electrostatic scaffold (LYS296, ASP422, LYS337) anchors each successive envelope macrostate visited during the sliding/rolling consolidation.

###### 3.11.7 Ternary Docking Reveals the TGFBR1–Env State as Consistently Superseded by Canonical TGF-$\beta$ Superfamily Assemblies

To evaluate how the TGF-$\beta$ signalling network modulates the dengue envelope–TGFBR1 reference interaction, we performed conditional ternary docking simulations incorporating ten first-neighbour proteins spanning the principal functional classes of TGFBR1 biology: ACVR2B (type II receptor, activin branch), FKBP1A (inhibitory GS-domain adaptor), GDF11 (TGF-$\beta$ superfamily ligand), SMAD2 (receptor-activated effector), SMAD7 (inhibitory Smad), SMURF2 (ubiquitin E3 ligase), TRAF6 (non-Smad signalling mediator), TGFB1 and TGFB3 (cognate TGF-$\beta$ ligands), and TGFBR2 (canonical type II partner receptor). Energetic coupling was quantified as $\Delta\Delta G$ values relative to the TGFBR1–Env reference pair (AB; $\Delta G \approx -6.4$ $\text{kcal} \cdot \text{mol}^{-1}$) [41,51,145].

The TGFBR1–Env reference baseline ($-6.4$ $\text{kcal} \cdot \text{mol}^{-1}$) is weaker than the primary membrane-receptor complexes (NRP1: $-12.8$; Betaglycan: $-13.8$) but substantially stronger than the SUMO1 reference ($-3.4$ $\text{kcal} \cdot \text{mol}^{-1}$). Across all ten tested partners, the dominant energetic signal is unidirectional and large: all partners strongly stabilise the AB axis in the AB-conditioned branch, consistently shifting AB from $-6.4$ to approximately $-16.4$ to $-17.1$ $\text{kcal} \cdot \text{mol}^{-1}$ ($\Delta\Delta G \approx -10.0$ to $-10.7$ $\text{kcal} \cdot \text{mol}^{-1}$), with the reverse-conditioned branches producing variable but generally also favourable or neutral AB shifts. This uniformity across mechanistically diverse partners — ligands, type II receptors, intracellular Smads, ubiquitin machinery, and an inhibitory adaptor — constitutes a robust signal that the TGFBR1–Env state is reliably displaced by canonical TGF-$\beta$ pathway configurations [140,146].

SMAD2 produced the largest single AB-conditioned cooperative shift: AB deepened from $-6.4$ to $-17.0$ $\text{kcal} \cdot \text{mol}^{-1}$ ($\Delta\Delta G \approx -10.6$), while BC weakened from $-12.8$ to $-8.9$ $\text{kcal} \cdot \text{mol}^{-1}$ ($\Delta\Delta G \approx +3.9$); no informative reciprocal state was detected. SMURF2 produced the highest combined AB cooperative signal across both branches: AB-conditioned $\Delta\Delta G \approx -10.7$ (AB reaching $-17.1$ $\text{kcal} \cdot \text{mol}^{-1}$) with pronounced BC destabilisation ($+7.4$ $\text{kcal} \cdot \text{mol}^{-1}$), and a reverse-conditioned AB gain of $\Delta\Delta G \approx -6.2$ (AB: $-12.6$ $\text{kcal} \cdot \text{mol}^{-1}$) — the strongest reciprocal-branch cooperative signal in the TGFBR1 panel. TRAF6 produced equivalent AB-conditioned stabilisation ($\Delta\Delta G \approx -10.3$; AB: $-16.7$ $\text{kcal} \cdot \text{mol}^{-1}$) with a strong reverse-conditioned gain ($\Delta\Delta G \approx -5.0$; AB: $-11.4$ $\text{kcal} \cdot \text{mol}^{-1}$).

TGFBR2 yielded the most internally consistent canonical receptor assembly signature: AB-conditioned $\Delta\Delta G \approx -10.5$ (AB: $-16.9$ $\text{kcal} \cdot \text{mol}^{-1}$), simultaneous BC destabilisation of $+7.0$ $\text{kcal} \cdot \text{mol}^{-1}$ (BC: $-5.7$ from

$-12.7$ kcal · mol$^{-1}$), and a robust reverse-conditioned AB gain ($\Delta\Delta G \approx -5.3$; AB: $-11.7$ kcal · mol$^{-1}$). GDF11 and TGFB1 produced highly selective AB-axis cooperative signatures without substantial reverse-conditioned perturbations: GDF11 ($\Delta\Delta G_{AB\text{-cond}} \approx -10.4$; BC +0.5; reverse +0.3) and TGFB1 ($\Delta\Delta G_{AB\text{-cond}} \approx -10.4$; reverse $-1.0$). TGFB3 matched this pattern ($\Delta\Delta G_{AB\text{-cond}} \approx -10.3$; reverse AB: $-7.7$). ACVR2B produced a clean, BC-sparing cooperative collapse: AB-conditioned $\Delta\Delta G \approx -10.0$; reverse $-0.8$; no preserved BC state.

FKBP1A produced the strongest combined bilateral cooperative signal: AB-conditioned $\Delta\Delta G \approx -10.1$ (AB: $-16.5$ kcal · mol$^{-1}$; BC: +0.7) combined with the strongest reverse-conditioned AB gain in the panel ($\Delta\Delta G \approx -7.1$; AB: $-13.5$ kcal · mol$^{-1}$). SMAD7 produced equivalent AB-conditioned cooperativity ($\Delta\Delta G \approx -10.2$; AB: $-16.6$ kcal · mol$^{-1}$) with a nearly neutral reverse branch ($\Delta\Delta G \approx +0.3$).

The dominant pattern across the full TGFBR1 panel is therefore a striking energetic convergence: regardless of whether the third partner is a ligand (TGFB1, TGFB3, GDF11), a type II receptor (TGFBR2, ACVR2B), an intracellular effector (SMAD2), an inhibitory regulator (SMAD7, FKBP1A), or a ubiquitin/non-Smad mediator (SMURF2, TRAF6), the AB-conditioned cooperative magnitude clusters within a narrow $\approx 1$ kcal · mol$^{-1}$ range ($-10.0$ to $-10.7$). The primary axis of variation among partners is therefore not the magnitude of AB stabilisation but the profile of BC perturbation and reverse-conditioned AB response, which together define the structural compatibility of each ternary assembly geometry.

#### 3.12 Genotypic Variation and Evolutionary Robustness of Viral–Host Interfaces

##### 3.12.1 Rationale and Analytical Framework

The sixteen envelope genotype variants examined in this study span all four DENV serotypes and major intraserotypic lineages, collectively representing the sequence diversity of endemic circulating strains [1,147]. While the molecular dynamics trajectories and per-complex MM/GBSA analyses reported in preceding sections were conducted on a single high-quality representative structure per host protein, the VectorPROT vectorial formalism was applied systematically across all sixteen genotype–host protein docking complexes to enable a direct genotype-comparative analysis of interaction geometry. This section reports the results of that comparison, addressing the central question: do mutational differences between genotypes alter the structural architecture of the interaction interfaces characterized in the preceding sections, or does the interaction landscape exhibit genotypic robustness?

##### 3.12.2 Conservation of Hotspot Architecture Across Genotypes

Per-residue analysis of docking complexes across the sixteen genotype panel reveals strong structural conservation of the primary energetic hotspot positions at key interfaces. The ARG410 equivalent — the dominant electrostatic anchor in the ENV–BiP complex — is structurally preserved as an arginine or lysine at the equivalent position in all sixteen variants, maintaining the capacity for salt-bridge formation with BiP GLU498 across the panel. Similarly, the DIII electropositive patch mediating GPC1 interaction retains its basic character across all variants, with the primary contributing residues (arginines and lysines in the $\beta$-hairpin and adjacent loops) showing only conservative substitutions within serotypes and no loss-of-function replacements between serotypes. The candidate CendR-like loop at the NRP1 b1-engagement surface preserves the terminal basic residue geometry across all sixteen variants, consistent with a structurally encoded — rather than sequence-specific — recognition mode [102,148].

The TGFBR1 contact triad (LYS296, ASP422, LYS337 on the receptor side) engages the envelope via a distributed surface rather than a single focal motif, and the broad envelope surface contributing to this interface is correspondingly tolerant of genotypic variation on the viral side — the receptor-side hotspot is conserved by definition, and the envelope-side contribution distributes across multiple residue positions none of which individually dominates the interface. This architecture — concentrated receptor-side hotspots against a distributed viral-side contact surface — provides inherent robustness to single amino acid changes in the envelope.

###### 3.12.3 VectorPROT Serotype-Resolved Surface Fitting: Directional Coherence Across Genotypes

Serotype-resolved surface fitting was applied to aggregate interaction vectors by genotype index for each host protein panel. For each protein, the quadratic surface fitted to the interaction vector tips from all sixteen genotypic variants was compared across serotype clusters (DENV1–4 intraserotypic groups). The mean resultant length $R$ of azimuthal angle distributions $\varphi_{xy}$ — a measure of directional coherence in the interface plane — remained within 15% of the panel mean for all sixteen variants across the GPC1, BiP, and NRP1 panels, indicating that the preferred vectorial orientation of interfacial contacts is conserved across the genotype panel despite primary sequence divergence. Larger inter-genotype variance in $R$ was observed for the TGFBR1 and Rab5 panels, consistent with the multi-macrostate dynamics of those complexes introducing greater sensitivity to the initial docking geometry and therefore to genotypic surface differences.

Differences in normalized circular entropy $H_{\mathrm{norm}}$ between genotypic clusters were most pronounced at interfaces dominated by a small number of directional hydrogen bonds (NRP1, SUMO), where substitution of even one contact residue shifts the angular distribution measurably. These differences in $H_{\mathrm{norm}}$ are below the threshold that would indicate a qualitatively different interaction mode, but they define a quantitative rank order of interface rigidity across genotypes that is testable against experimental affinity measurements.

###### 3.12.4 Genotypic Compensation and Evolutionary Redundancy of Receptor Engagement

A key prediction emerging from the multi-receptor interaction framework is that evolutionary pressure on the dengue envelope need not optimize any single receptor interface in isolation. Instead, the network architecture identified here — GPC1 as a modeled heparan sulfate attachment platform, NRP1 as a vascular-entry hypothesis, BiP for co-receptor attachment, Rab5 for trafficking, SUMO-Ubc9 for innate immune suppression — provides the virus with multiple partially redundant interaction axes. A genotype that acquires mutations reducing affinity at one interface can in principle compensate through enhanced engagement at another, provided the compensatory mutations do not simultaneously abrogate a fitness-critical interaction.

The vectorial analysis reveals a pattern consistent with this redundancy hypothesis. DENV2 variants carrying substitutions at the DIII surface that marginally reduce the directional coherence of GPC1-facing contact vectors show corresponding increases in $R$ at the NRP1-facing surface, suggesting that sequence variation at DIII simultaneously modulates both interfaces in opposite directions. If confirmed by quantitative $\Delta G$ measurement across the full panel, this anti-correlated variation would constitute structural evidence for a compensatory mechanism in which the DIII surface — shared between GPC1 electrostatic tethering and NRP1 CendR-like engagement — is under opposing selective pressures from attachment factor and entry receptor requirements [1–3]. This evolutionary trade-off model transforms the interaction landscape from a static binding map into a dynamic fitness surface, with the implication that genotype-specific differences in tissue tropism, endothelial permeability induction, and clinical severity may be mechanistically linked to the relative balance of GPC1 versus NRP1 engagement efficiency.

This compensatory model is a testable structural hypothesis: serotype-specific differences in plasma leakage severity in dengue hemorrhagic fever — most pronounced in DENV2 and DENV3 secondary infections [1] — may partly reflect serotype-level differences in NRP1 engagement efficiency, a prediction directly addressable by NRP1 competition pulldown assays using purified DENV1–4 envelope ectodomains.

### 4 Discussion

#### 4.1 The Envelope–Actin Interface Exhibits Hallmarks of a Functional Virus–Host Interaction

The integrated structural and energetic analyses presented here establish that the complex formed between the dengue envelope protein and cellular actin satisfies the criteria expected of a biologically functional protein–protein interaction. Global structural stability, in the absence of RMSD drift and without measurable reduction in the radius of gyration of either partner, demonstrates kinetic persistence on the nanosecond-to-submicrosecond timescale. The continuous maintenance of atomic-scale interchain contacts, as reflected in both minimum distance and contact number metrics, excludes the possibility that the observed interface represents a transient docking artifact or an energetically marginal association prone to spontaneous dissociation.

Critically, this stability is achieved without major conformational rearrangements in either protein. The absence of induced folding or unfolding events indicates that the interaction relies on surface complementarity and localized interfacial optimization — a binding mechanism consistent with functional protein–protein recognition rather than non-specific trapping. This observation aligns with established structural descriptions of the flavivirus E protein as a conformationally robust scaffold that undergoes targeted surface remodeling upon receptor engagement while preserving its core domain organization [3,5].

#### 4.2 Asymmetric Flexibility Partitioning Reflects a Conserved Mode of Virus–Host Engagement

The asymmetric distribution of conformational flexibility between the envelope protein and actin is mechanistically informative. The envelope protein functions as the comparatively rigid partner, maintaining a restricted and spatially consistent surface, while actin assumes the role of the conformationally adaptive host factor, modulating localized flexibility to optimize contact geometry. This division of structural labor — in which a viral protein presents a defined binding determinant to a plastically responsive host scaffold — is a recurrent organizational principle in virus–host protein interactions, particularly those involving cytoskeletal components [76].

Actin's functional design predisposes it to this adaptive role. As a protein evolved to alternate between globular and filamentous states and to interact with a structurally diverse repertoire of regulatory partners, actin possesses intrinsic conformational plasticity concentrated in specific subdomains [75]. The localized high-RMSF regions observed in the complex correspond to segments known for their flexibility in free actin structures, suggesting that the same conformational modes exploited by cellular actin-binding proteins are engaged during envelope association. This mechanistic convergence provides a structural rationale for why viral proteins can compete for or cooperate with cellular actin regulators, potentially influencing cytoskeletal organization during infection.

#### 4.3 Electrostatic Dominance and Energetic Redundancy Define the Interaction Architecture

The MM/GBSA energetic profile — characterized by strong electrostatic driving forces partially offset by polar desolvation penalties, supplemented by van der Waals stabilization — is consistent with an interface mediated by transient salt bridges and intermittent hydrogen bonds, reinforced by hydrophobic packing of the interfacial core. Interfaces of this type are intrinsically dynamic at the level of individual contacts yet globally persistent in terms of association, because the collective entropic and enthalpic balance favors complex maintenance even as individual residue pairs transiently separate and re-associate [47].

The distributed energetic architecture revealed by residue-level decomposition is particularly significant from the perspective of evolutionary robustness. Because the binding free energy arises from multiple spatially separated residue contributions rather than a single critical hotspot, the interaction is expected to be tolerant of individual point mutations in either partner. For a viral envelope protein interacting with a highly conserved cellular scaffold such as actin, this energetic redundancy provides a structural basis for maintaining host factor engagement across sequence variants, consistent with the functional conservation of actin-mediated flavivirus processes across DENV serotypes and related flaviviruses.

#### 4.4 Biological Implications: Actin as a Cytoskeletal Platform for Flavivirus Trafficking

The structural and energetic properties of the envelope–actin interaction described here are mechanistically congruent with established experimental evidence for cytoskeletal involvement in flavivirus infection. Actin dynamics regulate multiple stages of the flavivirus replication cycle, including clathrin- and macropinocytosis-mediated viral entry, transport of endocytic vesicles through the cortical actin network, and spatial organization of replication compartments within the perinuclear region [149–151]. Disruption of actin polymerization through pharmacological intervention significantly reduces dengue and related flavivirus infectivity, providing direct experimental support for the functional relevance of envelope–actin engagement [15,16].

The interaction mode characterized here — persistent, electrostatic in nature, locally flexible, and energetically distributed — is precisely compatible with a model of transient cytoskeletal anchoring that enables spatial coordination of early infection events without necessitating stable complex formation incompatible with actin dynamics. This type of interaction would allow the virion or newly internalized endosome to engage actin transiently as it is transported through the cytoplasm, exploiting the same dynamic binding and release cycles that characterize the interactions of cellular actin-binding proteins with the filament network. In this context, the envelope–actin interaction may function not as a classical receptor–ligand binding event but as a cytoskeletal tethering mechanism that positions viral components within productive intracellular trafficking routes.

Collectively, the molecular dynamics and MM/GBSA results establish a coherent structural–energetic framework for envelope–actin association in which kinetic persistence, conformational complementarity, and distributed electrostatic anchoring collectively underpin a mechanistically plausible and biologically grounded interaction. These findings motivate further investigation of temporal contact occupancy across individual residue pairs, comparative energetics across additional host factors and dengue genotypes, and experimental validation of the identified interface residues as functional determinants of cytoskeletal engagement during viral infection.

#### 4.5 Actin-Associated Host Proteins Modulate the ENV–Actin Interface Through Competitive and Cooperative Mechanisms

##### 4.5.1 Classical Actin Regulators Act as Competitive Modulators of the Viral Binding Surface

The cytoskeleton plays a central role in the replication cycles of many viruses, including flaviviruses such as dengue virus. Actin remodeling contributes to viral entry, intracellular trafficking, and particle release, and multiple studies have demonstrated that perturbation of actin dynamics alters dengue infectivity and replication efficiency [15,16]. Experimental work has also shown that dengue envelope protein can associate with actin or actin-associated complexes during infection, suggesting that cytoskeletal interactions contribute to viral internalization or post-entry trafficking.

The ternary docking analyses presented here extend this concept by examining not only the ENV–actin interaction itself, but also how the local actin regulatory environment modulates that interface. The results indicate that many actin neighbors behave as competitive modulators of the viral binding surface. Proteins such as $\alpha$-actinin (ACTN2) and profilin (PFN1) are well-established regulators of actin filament organization and polymerization dynamics, and both proteins interact with defined surfaces of actin monomers or filaments. Binding of these regulators can constrain actin conformational flexibility or occupy interaction determinants, which may explain the substantial energetic penalties observed when they associate with actin before viral docking. Once the ENV–actin interface has formed, however, these proteins appear capable of participating in a slightly stabilized ternary arrangement, suggesting that steric competition rather than absolute incompatibility underlies the observed energetic patterns.

##### 4.5.2 Dynactin-Associated Actin-Related Proteins Reflect Indirect Cytoskeletal Scaffolding

Proteins associated with the dynactin complex, including ACTR1A and ACTR10, exhibited similar energetic behavior. Dynactin acts as a scaffold for dynein-driven transport along microtubules and contains actin-related subunits that contribute to its structural organization. Viruses frequently exploit dynein–dynactin transport pathways to reach replication compartments within host cells. The modest stabilization observed in the Viral–Cell Triple_Inverse configuration may therefore reflect a structural environment in which actin-related dynactin components indirectly stabilize viral–host interfaces by organizing local cytoskeletal architecture.

CAPZB displayed a different energetic signature. As the $\beta$-subunit of the actin barbed-end capping protein, CAPZB stabilizes filament ends and regulates actin polymerization. The favorable Cell–Neighbor $\Delta\Delta G$ observed here is consistent with this structural stabilization of actin filaments. However, the minimal effect on the ENV–actin interface suggests that filament stabilization alone does not necessarily enhance viral binding, reinforcing the idea that viral engagement depends on specific actin conformational states rather than simply filament integrity.

##### 4.5.3 ACTL6A Emerges as a Cooperative Structural Facilitator of Viral Engagement

Among all neighbors examined, ACTL6A emerged as the most prominent cooperative factor. ACTL6A (also known as BAF53A) is an actin-related component of the SWI/SNF chromatin-remodeling complex and plays important roles in cellular organization and transcriptional regulation. Although its primary functions are nuclear, actin-related proteins in the SWI/SNF complex can influence cytoskeletal and organelle organization indirectly. The strong negative $\Delta\Delta G$ observed in the Viral–Cell Triple_Inverse configuration suggests that ACTL6A may create a structural environment compatible with ENV–actin association, possibly through indirect conformational stabilization of actin or through additional interaction surfaces that support ternary complex formation. The GO enrichment analysis linking ACTL6A to organelle organization further supports the notion that this protein may influence broader cellular architecture rather than acting solely as a classical actin-binding regulator.

Taken together, these findings highlight that the actin environment surrounding the viral envelope–actin interface is highly context dependent. Rather than forming a simple binary interaction, the ENV–actin complex appears embedded within a dynamic network of cytoskeletal regulators that can either compete with or subtly stabilize viral attachment depending on assembly order and local structural constraints. Such cooperative or competitive effects are consistent with the emerging view that viruses exploit host cytoskeletal systems as multi-protein interaction networks rather than isolated molecular partners [150,151].

From a mechanistic perspective, the data suggest a hierarchy of functional roles within the actin neighbor network. Classical actin regulators primarily modulate the accessibility of the viral binding interface, whereas certain actin-related proteins with broader cellular roles — exemplified here by ACTL6A — may act as structural facilitators of viral engagement. These distinctions provide a framework for prioritizing host factors in downstream dynamic simulations and experimental validation, particularly for proteins that exhibit strong cooperative energetic signatures in ternary complexes.

#### 4.6 Envelope–Betaglycan/TGFBR3: Mechanistic and Biological Interpretation

##### 4.6.1 Interfacial Maturation as a Structural Signature of Avidity-Driven Tethering

The monotonic increase in contact number and concomitant reduction in mean interchain distance observed across the 200 ns trajectory constitute a structural signature of interfacial maturation — a process in which local conformational exploration leads to progressive improvement of complementarity rather than passive thermal drift. The convergence of these metrics toward a tighter distributional regime in the final trajectory window indicates that the system accesses a conformational basin of greater stability than the initial docked configuration, a finding that reinforces the physical relevance of the predicted complex and argues against the possibility that late-trajectory contacts represent transient stochastic encounters.

This mode of progressive compaction is particularly congruent with the proposed biological role of betaglycan as a cell-surface co-receptor and capture scaffold. Betaglycan (TGFBR3) is a widely expressed transmembrane proteoglycan whose ectodomains carry heparan sulfate and chondroitin sulfate glycosaminoglycan chains, enabling multivalent interactions with extracellular ligands. In the context of its canonical signaling function, betaglycan operates by binding and concentrating ligands — including TGF-$\beta$ family members — at the cell surface before presenting them to signaling-competent receptors, thereby enhancing effective local concentration and association rates without requiring high intrinsic affinity at a single protein–protein contact [66,67]. An analogous mechanism is biologically plausible for flavivirus engagement: the progressive densification of the envelope–betaglycan interface observed in the simulation could reflect the structural correlate of this capture function, wherein the complex strengthens over timescales consistent with initial membrane docking events, increasing viral residence time and facilitating subsequent handoff to endocytic receptors.

##### 4.6.2 The Conformational Cost Asymmetry Reflects Ligand-Side Structural Adaptation

The greater RMSD displacement and higher RMSF values of betaglycan relative to the envelope protein establish a clear asymmetry in the conformational cost of complex formation that parallels the pattern observed for actin, but with mechanistically distinct origins. Whereas actin's flexibility arises from its intrinsic conformational switching between globular and polymeric states, betaglycan's conformational adaptation reflects the structural malleability of its modular extracellular architecture. The zona pellucida-like module and adjacent ectodomains of TGFBR3 are known to accommodate diverse binding partners through local rearrangements that expose or reposition contact surfaces without globally destabilizing the receptor fold [66,152]. The observation that the viral envelope preserves greater structural rigidity across the interaction, while betaglycan absorbs the bulk of the interfacial accommodation, is therefore mechanistically coherent and suggests that the envelope provides a structurally defined binding surface that betaglycan molds onto rather than a flexible scaffold that requires host-induced organization.

##### 4.6.3 Electrostatic Frustration Identifies Residues Governing Binding Efficiency

The identification of ARG107 and LYS236 on the envelope as major energetically unfavorable contributors represents one of the most mechanistically informative outputs of the MM/GBSA analysis. These basic residues likely experience partial burial at the interface without forming consistently compensatory ionic contacts or hydrogen bonds, generating net desolvation penalties that impose energetic tension on the complex. Two non-exclusive mechanisms can explain this frustration: either these residues are geometrically positioned at the interface periphery, where they are sufficiently buried to lose hydration but insufficiently paired to offset the desolvation cost; or they participate in intermittent contacts whose temporal average — as captured in the MM/GBSA calculation — reflects the energetic average across both favorable and unfavorable microstates.

In either case, the presence of energetically frustrated basic residues at a flavivirus envelope surface is functionally significant. Envelope residues rich in arginine and lysine are known determinants of heparan sulfate binding, and their partial engagement at the protein–protein interface in the absence of glycosaminoglycan chains — as modeled in the present simulations — may underestimate the favorable energetics achievable in the native cellular context, where sulfated glycosaminoglycan chains on betaglycan would provide polyanionic environments highly complementary to cationic envelope residues [9,153]. This observation motivates future simulations incorporating explicit glycosaminoglycan representation to assess whether the frustration associated with ARG107 and LYS236 is relieved by sulfated sugar contacts.

##### 4.6.4 Betaglycan as an Avidity Amplifier Rather Than a High-Affinity Entry Receptor

The energetic picture emerging from MM/GBSA analysis — stable but thermodynamically constrained, with distributed host-side contributions and frustrated viral-side residues — is most coherent with a biological model in which betaglycan functions as an avidity amplifier rather than as a primary high-affinity entry receptor. In this model, the protein–protein component of the interaction provides moderate intrinsic affinity, while the glycosaminoglycan chains of betaglycan contribute an additional electrostatic capture layer that increases the apparent association rate and extends surface residence time of envelope-bearing particles. This two-layer binding architecture — protein interface supplemented by glycan-mediated avidity — is well established for flavivirus interactions with heparan sulfate proteoglycans at the cell surface [9,153] and is consistent with the known ability of heparin and heparan sulfate mimetics to competitively inhibit dengue virus cell entry.

Critically, the distributed and diffuse energetic profile on the betaglycan side, devoid of dominant stabilizing hotspots, is precisely what would be expected if the protein–protein interface functions primarily to provide orientation and initial docking geometry, with the thermodynamic driving force for membrane association residing substantially in the glycosaminoglycan–envelope electrostatic interaction that is not captured in the present protein-only simulations. This mechanistic interpretation positions the current MM/GBSA results not as a complete thermodynamic description of the envelope–betaglycan interaction but as a characterization of its protein-mediated component, whose energetic constraints can be rationalized within the broader biophysical framework of proteoglycan-mediated viral attachment and the structural biology of TGFBR3 co-receptor function [66,67].

Together, the molecular dynamics and MM/GBSA analyses of the envelope–betaglycan system establish a mechanistic model of progressive interfacial compaction governed by electrostatic attraction, structurally limited by betaglycan-side conformational adaptation, and energetically constrained by polar frustration at two basic residues on the viral surface. This framework defines a coherent set of hypotheses — regarding glycosaminoglycan contributions, contact persistence at identified residue pairs, and time-resolved energetic evolution — that will be addressed in subsequent analyses integrating hydrogen bond occupancy, comparative MM/GBSA windows, and multi-genotype docking comparisons.

##### 4.6.5 Ternary Interaction Landscape Confirms Betaglycan–Env as a Conditional Energetic Attractor

The conditional $\Delta\Delta G$ analysis extends the binary characterisation of the Betaglycan–Env interface by embedding it within the interaction context of seven first-neighbour host proteins. The core observation — a conserved, stable AB pairwise energy coexisting with highly variable BC energetics and partner-specific ternary coupling signatures — is mechanistically coherent with the structural role of Betaglycan (TGFBR3) as a cell-surface co-receptor and broad-capture proteoglycan scaffold [66,67]. In its canonical signalling context, betaglycan functions not as a single high-specificity receptor but as a presentation platform that accommodates multiple extracellular interaction partners while retaining a stable core docking geometry [139,140]. The consistency of the AB interface energy at approximately $-13.8$ to $-13.9$ kcal $\cdot$ mol$^{-1}$ across all third-partner contexts is therefore consistent with a protein–protein contact engineered for positional stability, while the conditional perturbations reveal a surface susceptible to neighbour-dependent deformation depending on the chemistry and shape of the third partner. This framework extends and complements the avidity-amplifier model proposed for the binary Betaglycan–Env system: if Betaglycan functions primarily through kinetic capture and surface residence-time extension rather than obligatory high-affinity docking [9,153], then the conditional landscape described here determines which pericellular partners can coexist with, reinforce, or displace that capture function.

##### 4.6.6 TGF-Axis Partners Define a Permissive-to-Redistributive Gradient

The TGFBR2 and TGFB3 results reveal qualitatively distinct modes of engagement with the Betaglycan–Env state, fully consistent with their distinct biological roles in TGF-$\beta$ receptor organisation. TGFBR2, the obligatory type II signalling kinase in canonical TGF-$\beta$ receptor complexes, produces near-zero $\Delta\Delta G$ shifts in both conditioning branches, indicating that the Betaglycan–Env state and TGFBR2 engagement are energetically compatible under the modelled conditions. This is mechanistically sensible: in native TGF-$\beta$ biology, betaglycan functions upstream of TGFBR2 as a ligand-capture and presentation co-receptor, and the two receptors operate through coordinated rather than competing surface occupancies [139,140,144]. A weak conditional signature for TGFBR2 therefore reflects biological compatibility rather than mere coincidence.

TGFB3, by contrast, is a secreted ligand that engages signalling receptors with high potency and belongs to a ligand class whose receptor-occupancy competition is a core organising principle of pathway regulation [140,154]. The $\Delta\Delta G \approx +2.7\ \mathrm{kcal \cdot mol^{-1}}$ weakening of AB under AB conditioning is consistent with TGFB3 redistributing the betaglycan surface away from an Env-compatible geometry — a structural analogue of ligand-driven receptor reorganisation. The marked asymmetry between conditioning branches (+2.7 versus $-0.1\ \mathrm{kcal \cdot mol^{-1}}$) indicates that TGFB3 can disrupt an established Betaglycan–Env complex more effectively than it can prevent Env from engaging a pre-formed Betaglycan–TGFB3 assembly, reflecting the energetic priority of the high-potency TGFB3–Betaglycan interaction under sequential assembly conditions. Collectively, the TGF-axis cases define a gradient from permissive (TGFBR2) to redistributive (TGFB3), with biological coherence at both poles.

##### 4.6.7 KLK22 Identifies the Boundary Condition of Energetic Incompatibility

The KLK22 panel produces the strongest antagonistic signature in the entire Betaglycan-centred dataset and is mechanistically informative independently of whether a specific Betaglycan–Env–KLK22 triad can be validated experimentally. Kallikrein-related peptidases constitute a family of secreted serine proteases that function in extracellular proteolytic cascades, remodelling protein availability and pericellular signalling microenvironments through substrate-selective hydrolysis [155]. The $\Delta\Delta G \approx +5.1\ \mathrm{kcal \cdot mol^{-1}}$ collapse of the Betaglycan–Env interface under reverse conditioning far exceeds noise levels and indicates that a protease-shaped surface with KLK22-like properties is structurally incompatible with the Betaglycan–Env docking geometry when it has pre-occupied the betaglycan surface. Whether this incompatibility reflects steric occlusion, competition for overlapping contact determinants, or sensitivity of the modelled assembly to the characteristically electropositive, substrate-binding architecture of the kallikrein active site cleft requires structural analysis beyond the scope of the present docking framework.

The directional asymmetry reinforces this interpretation: KLK22 destabilises AB far more strongly under reverse conditioning than it affects BC under AB conditioning (+5.1 versus +1.0 $\mathrm{kcal \cdot mol^{-1}}$). This implies that a pre-formed Betaglycan–KLK22 complex presents a deformed or occluded surface incompatible with Env engagement, whereas Env-occupied betaglycan remains moderately accessible to KLK22 with only modest BC reduction. Physiologically, this asymmetry suggests that extracellular kallikrein activity in the perivirion space could constitute a contextual antagonist of Betaglycan-mediated viral attachment by competing for betaglycan surface occupancy upstream of viral docking — though this hypothesis requires experimental validation under conditions that control for kallikrein substrate specificity and membrane-anchored receptor geometry.

###### 4.6.8 Immunoglobulin-Variable-Domain Binders Impose the Strongest Cooperative Signatures

The IGKV2D-28 and IGHV3 results are among the most biologically interpretable in the Betaglycan ternary panel. The dengue E protein is the principal target of protective antibodies, and the most potent neutralising responses characteristically recognise surface-exposed epitopes on the envelope ectodomain, with several highly potent antibodies engaging quaternary or dimer-spanning configurations not accessible on monomers [156,157]. Against this background, the observation that immunoglobulin-variable-domain partners exert the largest cooperative effects across the entire panel — selective AB stabilisation by IGKV2D-28 ($\Delta\Delta G \approx -1.5$ $\mathrm{kcal \cdot mol^{-1}}$) and simultaneous AB consolidation with BC sequestration by IGHV3 — is fully consistent with antibody-like binders trapping Env in a restricted, interface-competent state from which competing host-surface partners are excluded.

The IGKV2D-28 signature is mechanistically distinct: the stabilisation of AB occurs without significant BC perturbation, indicating that this variable domain occupies a complementary surface region that reinforces the Betaglycan–Env geometry without competing for the BC contact area. The IGHV3 profile is more complex: the $\Delta\Delta G \approx +5.4$ $\mathrm{kcal \cdot mol^{-1}}$ destabilisation of BC under AB conditioning, combined with $\Delta\Delta G \approx -0.8$ $\mathrm{kcal \cdot mol^{-1}}$ AB stabilisation under reverse conditioning, is consistent with a surface occupant that simultaneously monopolises the Env interface from competing neighbours while consolidating the Betaglycan-centred state. This behaviour mechanistically resembles the sequestration model proposed for potent dengue-neutralising antibodies that restrict envelope conformational access and sterically block receptor engagement [156,157]. These results do not demonstrate that the specific variable domains tested are authentic dengue binders in vivo, but they establish that immunoglobulin-like interaction geometry is far more capable than most host-factor candidates of reshaping the conditional energy landscape around Env — a conclusion with direct relevance to structural models of antibody-mediated neutralisation.

##### 4.6.9 FGF1 and CANT1 Represent Functionally Distinct Positions in Glycan Ecology

FGF1 and CANT1 represent two functionally distinct relationships with betaglycan's glycan-dependent surface environment, and their contrasting energetic phenotypes are coherent with these distinct roles. FGF1 is a canonical heparan-sulfate-dependent growth factor whose assembly into stable FGF–FGFR signalling complexes requires heparin or heparan sulfate as an obligatory co-factor [158]. Because both FGF1 signalling and dengue envelope attachment compete for sulfated glycan surfaces [9,95,99], FGF1 is positioned as a natural extracellular competitor for the same glycan scaffolds that sustain betaglycan-mediated viral capture. The moderate, interface-selective weakening of BC ($\Delta\Delta G \approx +1.9$ kcal · mol$^{-1}$) without significant AB disruption is therefore consistent with FGF1 competing for sulfated-surface organisation rather than directly displacing the Betaglycan–Env protein–protein contact. In this model, FGF1 does not preclude Betaglycan–Env co-occupancy but selectively reduces the stability of the Env–neighbour interface, potentially by occupying heparan-sulfate chains that would otherwise scaffold BC contact geometry.

CANT1 belongs to a conceptually different category. As a calcium-activated nucleotidase localised primarily to the endoplasmic reticulum and Golgi apparatus, CANT1 functions in nucleotide sugar metabolism and is required for the biosynthesis of proteoglycan glycosaminoglycan chains, including those decorating betaglycan [95,99]. Its near-null energetic phenotype across all conditional configurations ($|\Delta\Delta G| \leq 0.2$ kcal · mol$^{-1}$) is precisely what would be expected from a protein whose functional relationship to betaglycan is biosynthetic and intracellular rather than competitive and extracellular. CANT1 shapes the glycan landscape on which Betaglycan–Env interactions depend but does not participate in the extracellular docking competition captured by the ternary framework. The contrast between CANT1's near-zero conditional signature and FGF1's moderate BC-specific effect therefore supports the methodological specificity of the ternary panel: the framework discriminates upstream biosynthetic regulators from extracellular co-competitors based on structural docking geometry alone.

###### 4.6.10 A Layered Model of the Betaglycan-Centred Viral Attachment Environment

Collectively, the ternary results support a layered model of the betaglycan-centred interaction environment. The Betaglycan–Env contact constitutes a conserved energetic attractor that is largely insensitive to bulk pericellular protein concentration but whose stability is selectively modulated by the molecular identity of neighbouring surface-accessible partners. TGFBR2 and CANT1 are permissive components that do not significantly perturb the attractor; their biological roles — canonical Betaglycan co-receptor and upstream GAG biosynthetic enzyme, respectively — are consistent with this permissiveness. FGF1 introduces moderate glycan-ecology competition that selectively disfavours BC contact geometry without eliminating the Betaglycan–Env state. TGFB3 redistributes the ensemble away from the attractor through directional, ligand-driven surface reorganisation. KLK22 defines an antagonistic boundary condition in which protease-shaped surface chemistry and the Betaglycan–Env docking geometry are mutually incompatible under sequential assembly. Immunoglobulin-like binders represent the most potent modulators, capable of either directly reinforcing the attractor (IGKV2D-28) or simultaneously consolidating it while sequestering competing surface access (IGHV3) — consistent with established models of antibody-mediated neutralisation.

The practical implication of this model is that the biological outcome of betaglycan-mediated viral docking may vary substantially depending on the local composition of the extracellular environment — including the presence of secreted growth factors (FGF1), extracellular protease activity (KLK22), TGF-$\beta$ superfamily ligands (TGFB3), or circulating immunoglobulins. As with the binary analyses, these conclusions should be interpreted as structurally derived hypotheses bounded by the conditional docking methodology, which does not incorporate membrane topology, glycan microheterogeneity, or dynamic conformational sampling beyond the fixed geometries used to construct each ternary assembly [41,47,49].

#### 4.7 Envelope–BiP/GRP78: Mechanistic and Biological Interpretation

##### 4.7.1 A Chaperone-Compatible Interaction Mode: Stable Yet Dynamically Permissive

The dynamical profile of the envelope–BiP complex is fully coherent with BiP's established molecular function as an ATP-dependent chaperone that engages client proteins through adaptable and persistent interfaces. BiP binds substrate proteins via a substrate-binding domain that adopts open and closed states in an ATP-hydrolysis-coupled cycle, enabling flexible yet stable substrate engagement that does not require rigid interface complementarity [69,79]. The moderately elevated RMSD and distributed RMSF of BiP observed in the simulation are therefore not indicators of interface weakness but rather mechanistically expected features of a chaperone operating in its substrate-engaged conformational regime.

From a virological standpoint, this interaction mode is particularly advantageous for the pathogen. By engaging a host chaperone whose conformational plasticity is evolutionarily optimized to accommodate diverse and imperfectly folded client surfaces, the dengue envelope protein may exploit BiP's intrinsic substrate tolerance to establish stable association without requiring precise geometric complementarity. This strategy would lower the energetic threshold for productive host factor engagement during the stochastic events of viral attachment and membrane docking. Consistent with this interpretation, BiP has been experimentally demonstrated to translocate to the cell surface under ER stress conditions — a well-documented hallmark of dengue infection — where it functions as a viral attachment and entry co-factor [12,13,68].

##### 4.7.2 Selective Surface Immobilization as a Mechanism for Preserving Viral Functional Integrity

The selective, spatially restricted reduction in envelope RMSF at the BiP-contacting surface — without global propagation of rigidification across the viral protein — represents a mechanistically important feature of this interaction. Maintaining conformational freedom in the non-contacting regions of the envelope protein is functionally critical, as the flavivirus E protein must retain the capacity to undergo large-scale domain rearrangements during the subsequent pH-triggered membrane fusion step. Irreversible or globally distributed immobilization of the envelope by a host factor encountered during the pre-endosomal attachment phase would be incompatible with the conformational transitions required for productive infection [5,159].

The pattern observed here — BiP anchoring discrete envelope surface motifs while leaving the fusion-relevant domains dynamically accessible — is therefore consistent with a model in which BiP engagement provides initial structural support during membrane docking without committing the envelope protein to a non-productive conformational state. This selectivity may represent a co-evolutionary feature, whereby the envelope protein has optimized the spatial segregation of BiP-binding determinants from fusion-relevant conformational switches.

##### 4.7.3 Electrostatic Anchoring and Hydrophobic Reinforcement Define a Two-Component Binding Architecture

The MM/GBSA energetic profile — large attractive EEL, large repulsive EGB, and a modestly negative total — positions the envelope–BiP interaction in a well-characterized thermodynamic regime for protein–protein interfaces enriched in ionizable residues. In this regime, the binding affinity is generated by the combined action of specific electrostatic contacts — primarily the ARG410–GLU498 pairing — and a secondary layer of hydrophobic packing contributed by aliphatic residues, whose van der Waals interactions partially shield the electrostatic hotspot from solvent, thereby improving the efficiency of charge–charge contact formation and reducing the effective desolvation penalty [49,77,84].

This two-component architecture — an electrostatic anchor supplemented by a hydrophobic packing scaffold — is structurally coherent and consistent with the known surface chemistry of both proteins. The envelope protein presents a surface enriched in basic residues that mediate electrostatic interactions with heparan sulfate and host receptors, while BiP's substrate-binding cleft contains both hydrophobic and charged residues that accommodate diverse client sequences. The identification of ARG410 and GLU498 as the primary energetic anchor pair across both SDC and TDC decompositions, reproduced consistently in both analyses, provides confidence that this contact represents a genuine interfacial hotspot rather than a decomposition artifact.

##### 4.7.4 Electrostatic Frustration at BiP Lysine Clusters Reflects Incomplete Charge Compensation

The substantial unfavorable contributions of B:LYS435 and the adjacent lysine cluster on the BiP surface constitute the most energetically costly feature of the complex and merit mechanistic interpretation. Lysine residues at protein–protein interfaces generate unfavorable MM/GBSA decomposed energies when they are sufficiently buried to lose hydration without forming compensatory salt bridges or hydrogen bonds with complementary partners, a condition described as electrostatic frustration [78]. In the present context, the spatial arrangement of these lysines relative to the envelope surface apparently does not provide sufficient anionic complementarity to compensate their desolvation cost under the protein-only simulation conditions modeled here.

This observation parallels the frustration identified at ARG107 and LYS236 in the betaglycan complex and similarly suggests that the in silico model, which omits post-translational modifications and glycosaminoglycan chains present on native BiP-expressing cell surfaces, may underestimate the favorable contributions available in the physiological context. BiP at the cell surface may present an environment in which adjacent anionic molecules or carbohydrate chains partially relieve lysine frustration through additional charge compensation, a possibility that warrants exploration in future simulation frameworks incorporating explicit representation of post-translational modifications.

##### 4.7.5 Electrostatic Frustration at Viral–Host Interfaces as a Privileged Target for Entry Inhibitors

Beyond their significance as a methodological caveat, the frustration hotspots identified here — B:LYS435 and its flanking cluster in the BiP complex, and ARG107/LYS236 in the betaglycan complex — carry a positive pharmacological implication that reframes them as structural opportunities rather than limitations. In small-molecule drug discovery, interfacially frustrated charged residues define sites of high conformational lability: the incomplete electrostatic compensation that generates unfavorable MM/GBSA decomposition terms reflects a genuine structural tension in the interface, a local energy landscape in which the bound state has not fully minimised its charge–charge interactions. Such sites are, precisely because of this lability, privileged binding pockets for designed small molecules capable of supplying the missing anionic or cationic complementarity that the protein partner alone cannot provide [160,161].

The principle that electrostatically frustrated interfacial sites are disproportionately druggable has been demonstrated across multiple protein–protein interaction (PPI) inhibitor programmes [162,163]. Fragment-based and virtual screening campaigns targeting interfacial basic residue clusters — structurally analogous to the LYS435 environment in BiP — have yielded small-molecule wedges that insert between the protein partners, simultaneously resolving the frustrated charge geometry and blocking the conformational adjustments required for productive complex formation or downstream signalling transitions [164,165]. In the present context, a compound designed to engage the LYS435/LYS516/LYS447 cluster on BiP's surface — for example by presenting an anionic pharmacophore (sulfonate, carboxylate, or phosphonate) that neutralises the desolvation penalty — would be predicted to stabilise a non-productive BiP–Envelope geometry while simultaneously blocking the conformational transitions in BiP required for ATP-dependent chaperone cycling and co-receptor function [69,166].

Analogously, the ARG107/LYS236 frustration sites in the betaglycan complex define a pair of entry points for compounds that mimic the anionic complement normally contributed by heparan sulfate chains on the native proteoglycan surface. Heparan sulfate–mimetic compounds, including sulfonated small molecules and polyanion scaffolds, have already been explored as dengue entry inhibitors at the level of the viral attachment step [97,167]; the per-residue decomposition data presented here provide a refined structural hypothesis for their mechanism of action and a basis for rational optimisation of the anionic contact geometry at the specific betaglycan–Envelope interface.

Collectively, the frustration sites identified across multiple complexes in this study constitute a pharmacologically actionable map: rather than representing regions to be avoided in modelling, they identify the most conformationally labile and charge-sensitive sub-surfaces of each interface, where the energetic barrier to small-molecule insertion is lowest and the structural consequence of successful binding — interface stabilisation in a non-productive geometry — is most directly linked to blockade of viral entry. This reinterpretation positions electrostatic frustration analysis as a productive step in the prioritisation of PPI targets for antiviral drug discovery directed at dengue envelope–host receptor interfaces.

##### 4.7.6 Electrostatic Frustration Sites as Targets for Molecular Glue Therapeutics

The pharmacological logic of exploiting electrostatic frustration sites can be extended beyond classical PPI inhibition into the mechanistically distinct category of *molecular glues* — small molecules that stabilise a specific protein–protein complex rather than disrupting it [168,169]. In the context of viral entry, the therapeutic objective of a molecular glue would not be to prevent the virus from binding to BiP or betaglycan, but to *trap* the virion in a surface-bound state from which it cannot proceed to endosomal internalisation or pH-triggered membrane fusion. A molecule that resolves the electrostatic frustration at B:LYS435 — for example by bridging this lysine to a complementary anionic residue on the viral envelope surface — would thermodynamically stabilise the surface-attached state, increasing the energetic barrier for dissociation and blocking the conformational transitions in both BiP (ATP-driven open-to-closed cycling) and the envelope (pH-triggered dimer-to-trimer rearrangement) that are required for productive entry [5,69].

This molecular glue mechanism operates through a different kinetic principle than competitive inhibitors and thus offers a distinct therapeutic profile. Competitive PPI inhibitors must outcompete the viral ligand for the host receptor surface, requiring high concentrations to maintain occupancy at the cell surface; molecular glues accumulate at and reinforce already-formed complexes, potentially requiring lower concentrations to achieve equivalent antiviral effect [168]. Furthermore, the therapeutic endpoint of a viral-entry molecular glue — irreversible or near-irreversible surface trapping — is cell-type-independent and does not depend on the density of alternative entry receptors, unlike receptor-blocking strategies that can be circumvented by receptor switching.

For the BiP frustration cluster, the structural requirement for a molecular glue is well-defined by the per-residue decomposition data: the molecule must supply anionic complementarity to LYS435, LYS516, and LYS447 while maintaining contacts with the envelope surface in the geometry established by the MM/GBSA analysis. The envelope surface residues forming the opposite face of this frustrated interface — A:GLN211 and A:LYS210, both of which contribute unfavorable decomposition terms — suggest that the interface harbours a charge-mismatched zone whose resolution by a bridging molecule with appropriate charge distribution would simultaneously relieve frustration on both protein surfaces. Computational fragment screening at this bifacial site, guided by the electrostatic potential maps of both proteins in the interface-bound state, represents a tractable first step toward molecular glue identification for this target [160,164]. For the betaglycan frustration cluster at ARG107 and LYS236, the anionic character of native heparan sulfate chains provides an immediate chemical template: sulfated monosaccharide mimetics or polyanionic scaffolds that replicate the charge distribution of HS at this specific interface could serve as the bridging pharmacophore, building on existing heparan sulfate mimetic antiviral strategies with added structural specificity [97,167].

###### 4.7.7 Biological Context: BiP as a Stress-Induced Viral Co-Receptor at the Cell Surface

The structural stability, persistent contact network, and moderate but favorable net binding free energy collectively support a biologically plausible model of BiP-mediated viral co-receptor function at the dengue-infected cell surface. Under ER stress — induced by dengue infection and by the general challenge of producing large quantities of viral glycoproteins in the secretory pathway — GRP78/BiP undergoes upregulation and surface translocation, becoming accessible to extracellular viral particles [68,170]. In this context, the interaction characterized here would represent an early-stage attachment event in which cell-surface BiP serves as a capture factor, increasing the local concentration of virions at the plasma membrane and potentially co-localizing them with endocytic machinery.

The composite interface architecture characterized here — with a discrete electrostatic anchor at ARG410–GLU498 and reinforcing hydrophobic contacts — provides a structural basis for this capture function. The modest net free energy of binding is mechanistically appropriate for a co-receptor whose function is to facilitate handoff to downstream entry factors rather than to irreversibly sequester the viral particle, which would be counterproductive for infection progression. Together, these analyses establish the envelope–BiP interaction as a thermodynamically plausible, dynamically coherent, and biologically contextualized component of the dengue virus entry landscape, providing a well-defined energetic baseline for comparative analysis across additional host proteins and viral genotypes.

##### 4.7.8 Ternary Landscape Positions BiP–Env as a Proteostasis-Network-Dependent Energetic Node

The conditional $\Delta\Delta G$ analysis reveals that the BiP–Env interface does not behave as an isolated binary contact: it functions as a proteostasis-context-sensitive energetic node whose stability is strongly modulated by the identity of neighbouring ER-resident partners. The invariance of the AB reference energy at approximately $-14.2$ kcal · mol$^{-1}$ across all seven third-partner configurations is not trivial — it confirms that the BiP–Env binary geometry is intrinsically stable and not susceptible to random perturbation. However, the structured, partner-specific conditional signatures superimposed on this stable baseline reveal a surface that is selectively responsive to the functional state of the surrounding ER chaperone machinery. This finding is biologically coherent: BiP/GRP78 is the master ER Hsp70 and the central organising node of ER protein quality control, coordinating substrate engagement, unfolded protein response (UPR) sensing, and functional handoff to co-chaperones, oxidoreductases, and downstream quality-control factors [69,79]. During dengue infection, the ER is extensively remodelled to support viral glycoprotein maturation, and BiP upregulation and surface translocation have been directly documented [68,170]. In this context, a conditional landscape in which some ER neighbours reinforce and others strongly antagonise the BiP–Env basin reflects the known functional specialisation of ER proteostasis networks and provides a structural-energetic framework for interpreting the dependence of dengue ER engagement on local chaperone network composition.

##### 4.7.9 J-Domain Co-Chaperones and AMPylation Regulators Define the Cooperative Class

The two cooperative partners — DNAJB11 and FICD — are mechanistically the most interpretable cases in the panel. DNAJB11 (also known as ERdj3) is an ER-localised Hsp40 J-domain co-chaperone that binds unfolded or partially misfolded substrates and delivers them to BiP, stimulating BiP's ATPase activity and substrate-loading cycle [69,171]. The $\Delta\Delta G \approx -4.8$ kcal · mol$^{-1}$ stabilisation of AB in the DNAJB11 case — with BC essentially unchanged and no reverse-branch perturbation — is precisely the energetic signature expected from a co-chaperone that deepens rather than competes with the BiP-centred client-binding state. Within the ternary framework, DNAJB11 reinforces the BiP–Env basin without introducing competing interface demands, consistent with its native function as a substrate-delivery scaffold that operates cooperatively with rather than in opposition to BiP's client-binding domain [171].

FICD operates through a distinct but functionally convergent mechanism. FICD is the sole human AMPylase/de-AMPylase known to directly regulate BiP activity by reversible adenylylation of a threonine residue (Thr518) in the substrate-binding domain: AMPylation inactivates BiP's client-holding function during periods of low ER stress, whereas de-AMPylation reactivates it during elevated client load [172].

The $\Delta\Delta G \approx -3.1$ kcal · mol$^{-1}$ stabilisation produced by FICD therefore suggests that a de-AMPylated, client-engagement-competent BiP conformation — or a surface state accessible to FICD's de-AMPylase domain — is more compatible with Env-containing ternary assemblies than the unregulated reference state. In mechanistic terms, both cooperative partners are directly wired into BiP's own functional regulation cycle, and their stabilising energetic signatures may reflect selection for a BiP conformational state optimised for ER client engagement rather than generic protein–protein recognition.

###### 4.7.10 ERAD Components and Parallel Chaperone Systems Define the Antagonistic Class

The four antagonistic partners — HYOU1, DERL1, HSP90B1, and PDIA6 — represent four mechanistically distinct modes of opposition to the BiP–Env state, and their differentiated biology clarifies why each produces a large positive $\Delta\Delta G$.

DERL1 (Derlin-1) is a core component of the ER-associated degradation (ERAD) machinery, functioning as a membrane-embedded retrotranslocation channel that exports misfolded luminal and membrane substrates into the cytoplasm for proteasomal destruction [173]. The $\Delta\Delta G \approx +7.5$ kcal · mol$^{-1}$ collapse of AB in the DERL1-conditioned configuration is consistent with a transition away from a productive chaperone-client assembly toward a triage-oriented state, in which the BiP surface is redirected toward ERAD substrate processing rather than retentive client binding. In the context of dengue infection, this conditional antagonism positions DERL1 as a potential negative regulator of BiP-mediated viral engagement: if DERL1 competes for BiP surface availability under conditions of elevated ER stress and misfolded protein load, it may reduce the probability of stable BiP–Env co-complex formation.

HSP90B1 (GRP94) is an ER-paralogue of cytosolic Hsp90 that handles a distinct and partially non-overlapping clientele, including immunoglobulins, toll-like receptors, and selected secretory proteins [174]. Despite being an ER chaperone like BiP, GRP94 does not serve as a general Hsp70 surrogate — its substrate specificity, co-chaperone dependencies, and nucleotide regulation are fundamentally distinct [174]. The $\Delta\Delta G \approx +5.2$ kcal · mol$^{-1}$ destabilisation of AB in the HSP90B1 case therefore does not indicate simple chaperone overcrowding but rather reflects introduction of an alternative ER handling pathway that is structurally incompatible with the BiP-centred basin under the modelled conditional geometry.

HYOU1 (also known as ORP150 or GRP170) is the largest member of the Hsp70 superfamily, localised to the ER lumen and strongly induced during hypoxic and ER stress [175]. Although it can function as a nucleotide-exchange factor (NEF) for BiP, stimulating ADP release and resetting BiP's ATPase cycle, it also harbours its own substrate-binding capacity. The near-+9.4 kcal · mol$^{-1}$ AB destabilisation induced by HYOU1 is the largest single antagonistic effect in the entire ternary dataset. This extreme disruptive signature may reflect competition for overlapping BiP surface contact determinants, since HYOU1 in its NEF role binds the nucleotide-binding domain of BiP at a region that influences substrate-binding domain allostery [175].

PDIA6 is a member of the protein disulfide isomerase (PDI) family that participates in redox-dependent glycoprotein quality control and has been specifically implicated in flavivirus replication through disulfide management of viral structural proteins [176]. Its $\Delta\Delta G \approx +6.9$ kcal · mol$^{-1}$ AB penalty under conditioning indicates that the introduction of PDIA6-mediated redox handling pulls the BiP surface ensemble away from the Env-compatible binding geometry and toward a redox-processing state. Critically, the near-null P4HB result demonstrates that this antagonism is PDI-family-member-specific rather than a generic property of ER oxidoreductases: P4HB and PDIA6 are structurally related but functionally non-equivalent, and the conditional framework discriminates them on the basis of their distinct effects on the BiP–Env interface topology.

###### 4.7.11 A Proteostasis-State Model for BiP–Env Conditional Compatibility

Collectively, the BiP ternary panels support a structured mechanistic model in which the BiP–Env energetic state is embedded within — and sensitive to — the functional state of the surrounding ER proteostasis network. Partners that deepen BiP's own client-handling cycle (DNAJB11, FICD) reinforce the BiP–Env basin; partners associated with ERAD routing, parallel high-specificity chaperone niches, alternative nucleotide-exchange-factor functions, or PDI-family redox processing (DERL1, HSP90B1, HYOU1, PDIA6) displace the system away from it; and P4HB is energetically permissive.

The asymmetry between conditioning branches — where the AB-conditioned configuration consistently shows large antagonistic effects while the reverse-conditioned branch produces only trivial perturbations — is a structurally important feature common to all four antagonists. This asymmetry indicates that the disruptive effect is exerted preferentially from within an established BiP–Env complex rather than by preventing Env from accessing a BiP surface already occupied by the antagonist. The mechanistic implication is that disruption of the BiP–Env state depends on ternary assembly order, suggesting that timing of ER chaperone engagement relative to viral protein arrival may influence the probability of productive BiP–Env co-complex formation in infected cells.

From a broader biological perspective, these findings argue that "BiP-dependence" in dengue infection should not be treated as a single fixed phenotype but as a network-context variable. In an ER environment enriched in DNAJB11-type co-chaperone activity or active FICD-mediated BiP regulation — conditions consistent with early productive infection — the BiP–Env basin is deepened. In an ER environment shifted toward ERAD processing (DERL1), stress-adaptive alternative chaperone deployment (HYOU1, HSP90B1), or redox-driven quality control (PDIA6) — conditions potentially characteristic of late or highly stressed infection stages — the same interface is strongly destabilised. This context-dependence is precisely the logic by which broad-spectrum ER proteostasis regulators can function as antivirals: they need not destroy the BiP–Env binary contact directly but only shift the surrounding network state sufficiently to displace the virus from its preferred energetic basin [47,49].

As with all conditional $\Delta\Delta G$ analyses presented in this study, these conclusions are bounded by the docking framework: they do not resolve temporal assembly order within infected ER membranes, glycan occupancy states, ATP/ADP cycling of BiP, or the effects of membrane geometry on ternary complex formation. The strongest priorities for downstream validation are the DNAJB11/FICD cooperative axis and the extent to which DERL1, HYOU1, HSP90B1, and PDIA6 genuinely redirect Env from BiP-favoured states in experimental infection contexts [41,51].

#### 4.8 Envelope–Clusterin: Mechanistic and Biological Interpretation

##### 4.8.1 Hydrophobic Tail Engagement as a Mechanistically Distinct Binding Mode

The structural and energetic profile of the envelope–clusterin complex defines a binding mode fundamentally distinct from those characterized for betaglycan and BiP, and this distinction has direct mechanistic and biological implications. Whereas the betaglycan interface is governed by long-range electrostatic attraction followed by geometric compaction, and the BiP interface is stabilized by a discrete electrostatic anchor reinforced by hydrophobic packing, the envelope–clusterin interaction is primarily driven by van der Waals–dominant contacts mediated by clusterin's intrinsically disordered hydrophobic tail sequences. This mechanism parallels that of small heat shock proteins, whose N-terminal extensions engage exposed hydrophobic patches on misfolded or stressed client proteins through flexible, adaptive surface sampling rather than through rigid complementarity [85,86]. The viral envelope protein, as a glycosylated surface protein that undergoes dynamic conformational transitions during acidification and membrane fusion, presents transiently exposed hydrophobic regions that are structurally accessible to clusterin tail sequences — regions that in the cellular context would correspond to the partially unfolded or conformationally stressed intermediates recognized by extracellular chaperones [87,88].

The absence of progressive interfacial compaction in the contact number profile — in contrast to betaglycan — indicates that clusterin achieves its binding mode rapidly and maintains it through the intrinsic affinity of tail–hydrophobic surface interactions rather than through slow rearrangement. This kinetically rapid engagement is consistent with the known behavior of extracellular chaperones, which must intercept stressed substrates before aggregation occurs in the extracellular space, demanding fast binding kinetics over optimized geometric complementarity.

##### 4.8.2 Distinct Epitope Usage Separates Clusterin from BiP on the Envelope Surface

The non-overlapping RMSF reduction patterns on the envelope surface between the clusterin and BiP complexes suggest that the two chaperones engage distinct surface regions of the viral protein. This epitope separation has important biological consequences. If clusterin and BiP bind at non-competing sites, they could in principle simultaneously engage the envelope protein — a scenario that is consistent with the anti-apoptotic BiP–clusterin interaction axis that has been described at the intracellular level under ER stress conditions [28,177]. The possibility that the envelope protein could bridge extracellular clusterin and cell-surface BiP simultaneously would create a ternary capture complex that increases both avidity and membrane residence time beyond what either chaperone alone could achieve. This model is speculative at present but is experimentally testable by surface plasmon resonance or cryo-electron tomography of virion–cell contact zones under ER stress conditions.

The distinct epitope usage also implies that inhibitors targeting the BiP–envelope interface would not necessarily abrogate clusterin-mediated capture, and vice versa. This functional redundancy at the level of chaperone engagement may represent an evolved viral strategy for ensuring robust cell-surface retention across diverse cellular stress states and chaperone expression levels — a strategy particularly relevant given the wide variation in clusterin and BiP expression across dengue target cell types (monocytes, dendritic cells, hepatocytes, and endothelial cells).

##### 4.8.3 Energetic Comparison Across the Chaperone Cluster Reveals a Hierarchy of Binding Mechanisms

Comparison of the MM/GBSA profiles across the three chaperone-related host proteins in this study — BiP, clusterin, and, in the context of the broader homeostasis cluster, NLRP3 (which interacts with chaperone signaling outputs) — reveals a mechanistic hierarchy with direct functional implications. BiP presents the largest magnitude of electrostatic stabilization and the most significant desolvation penalty, consistent with a high-polarity interface that extracts the maximum enthalpy from charge complementarity at the cost of solvation, yielding a moderate net free energy. Clusterin, by contrast, presents a more hydrophobically weighted interface with attenuated polar desolvation costs, yielding a smaller absolute electrostatic contribution but also a smaller solvation penalty and a comparably moderate net free energy.

This divergence in underlying energetic mechanisms for similar net thermodynamic outcomes reflects the different substrate-recognition strategies of the two chaperones: BiP evolved to recognize exposed charged/polar patches on misfolded ER clients, whereas clusterin evolved to recognize exposed hydrophobic patches on extracellular proteins undergoing stress-induced conformational change [69,88]. The dengue envelope protein appears to exploit both mechanisms — its amphoteric surface presenting both polar and hydrophobic regions accessible to the respective chaperone — enabling broad-spectrum chaperone capture across compartments.

###### 4.8.4 Biological Role: Clusterin as an Extracellular Virion-Stabilizing Factor During Stress

The biological model emerging from these analyses positions clusterin as an extracellular virion-stabilizing and membrane-tethering factor whose relevance is particularly acute under cellular stress conditions. Clusterin expression is potently induced by diverse stress stimuli — including oxidative stress, heat shock, cytokine signaling, and ER stress — and is upregulated in dengue-infected tissues [86,87]. As a secreted glycoprotein abundant in plasma and interstitial fluids, clusterin is present at concentrations reaching several micrograms per milliliter in the extracellular environment of dengue target tissues, positioning it as a constitutively available binding partner for the viral envelope [88].

The functional implications of envelope–clusterin engagement extend in two directions. First, clusterin binding to the envelope could stabilize the virion against thermodynamic stress in the extracellular space, reducing aggregation and preserving infectivity over extended circulation times — a role analogous to its documented function in preventing amyloid aggregation [85]. Second, by presenting the virion to megalin/LRP2 or other clusterin receptors on the cell surface, clusterin could facilitate receptor-mediated endocytic uptake of viral particles through the established clusterin–LRP2 internalization pathway, effectively co-opting a host proteostasis clearance mechanism for viral entry. This Trojan horse model of chaperone-mediated viral endocytosis is consistent with the general principle, established for BiP and supported by the present analyses, that DENV exploits host proteostasis machinery as an entry scaffold rather than relying exclusively on classical attachment receptors. Together, the structural, dynamic, and energetic characterization of the envelope–clusterin interface establishes this interaction as a mechanistically distinctive and biologically plausible component of the dengue surface proteome engagement network, warranting direct experimental validation in stress-primed cellular models.

#### 4.9 Envelope–Glypican-1: Electrostatic Saturation, Multivalent Heparan Sulfate Recognition, and Attachment-Phase Architecture

##### 4.9.1 The Electrostatically Saturated Interface Defines a Charge-Complementary Attachment Mechanism

The MM/GBSA energetic profile of the envelope–GPC1 complex — characterized by dominant EEL stabilization, near-commensurate EGB desolvation penalty, and a net $\Delta G_{\text{total}}$ near $-90$ kcal/mol — is qualitatively and quantitatively distinct from those of all other complexes analyzed in this study. The magnitude of both the stabilizing and penalizing terms exceeds those of the BiP, betaglycan, actin, and clusterin interfaces, collectively placing GPC1 in a class of interactions defined by electrostatic saturation: the interface is so densely charged that each additional salt bridge or hydrogen bond contact generates both large stabilization and large desolvation costs, yielding a net free energy that emerges as the small difference of two very large numbers. This energetic architecture is the hallmark of heparan sulfate–protein interactions characterized in the broader context of viral attachment, where the multiplicity of sulfate groups on HS chains engages multiple basic residues simultaneously, generating interaction energies far exceeding those typical of purely proteinaceous interfaces [95,97,99].

The mechanistic consequence of this electrostatically saturated architecture is an interaction whose net thermodynamic stability depends critically on the precise density and distribution of charges at the interface. The large oscillations in $\Delta G_{\text{total}}$ observed within the final 10 ns window — spanning a range of approximately 60 kcal/mol between least and most favorable frames — do not indicate macroscopic instability, but rather reflect the extreme sensitivity of the MM/GBSA energy to local microconfigurations of charged contacts. When a single salt bridge transiently opens and reforms with a shifted geometry, the instantaneous EEL and EGB values both change by tens of kcal/mol while the structural integrity of the complex is fully maintained. This behavior underscores the importance of interpreting HSPG-mediated interaction energies at the ensemble level rather than from individual snapshots, and highlights the complementarity between geometric descriptors (contact persistence, minimum distance stability) and energy-based metrics for characterizing this class of interface.

##### 4.9.2 Domain III as the Structural Epicenter of HSPG-Mediated Envelope Recognition

The concentration of favorable per-residue free energy contributions in the domain III electropositive patch, combined with the specific RMSF dampening observed in this region, provides structural evidence consistent with domain III as the primary interface for GPC1 engagement. This is mechanistically significant because domain III of the dengue envelope protein serves dual functional roles: it harbors the receptor-binding determinants engaged during productive entry, and it constitutes a principal target of neutralizing antibodies, particularly those with cross-reactive breadth across DENV serotypes [156,157]. The observation that GPC1 engages domain III with high affinity and low mobility in the simulation implies that GPC1-mediated surface attachment could compete with, or spatially overlap with, the binding sites of domain III-targeting neutralizing antibodies. This spatial relationship is consistent with the known capacity of HSPG to modulate antibody-dependent neutralization through competitive surface occupancy, and suggests a mechanistic basis for the reported variation in dengue neutralization efficiency as a function of cell-surface HSPG expression levels [95,96].

The directed conformational transition of the envelope chain observed in the PCA analysis — a progressive rearrangement from an initial to a later-simulation state coinciding with GPC1 contact stabilization — is interpretable as an induced-fit adjustment in which domain III accommodates the geometric constraints imposed by sustained contact with the GPC1 surface. Unlike the BiP complex, where the envelope remained broadly conformationally stable throughout, and unlike the betaglycan complex, where the host side undergoes the majority of structural adaptation, the GPC1 complex distributes the conformational cost between both partners: the envelope DIII adjusts its surface geometry to maximize charge complementarity, while GPC1 samples multiple orientations to identify the optimal multi-contact configuration. This cooperative induced-fit mechanism is consistent with the general principle of multivalent receptor recognition, in which neither partner achieves its final binding geometry independently, but rather through iterative mutual adjustment [178].

##### 4.9.3 Multivalency and the Kinetics of Surface-Phase Concentration

The structural basis of GPC1 function in dengue attachment is fundamentally multivalent: the core protein presents three to four HS chains, each capable of engaging basic envelope residues independently, generating an avidity-based interaction whose dissociation rate is far lower than that of any individual chain–protein contact [95,98]. The simulation model used here represents the GPC1 core protein without explicit HS chains — a computational simplification that necessarily underestimates the true binding free energy and contact multiplicity of the physiological interaction. The energetic and structural parameters reported here should therefore be interpreted as representing the floor of the GPC1 contribution to dengue attachment: the core protein alone generates a persistently stable complex with favorable net free energy and extensive electrostatic complementarity, and the full HS-decorated proteoglycan in the physiological context would engage a substantially larger and more redundant contact network. This interpretation is supported by the analogy with betaglycan, where the absence of explicit GAG chains produces electrostatic frustration at basic envelope residues that would be alleviated by charge compensation from the anionic GAG chains in the native protein [167].

The kinetic dimension of GPC1-mediated attachment — rapid association driven by long-range electrostatic steering, sustained association driven by multivalent HS chain engagement, and slow dissociation due to the high rebinding probability of proximate chains — provides a mechanistic framework for the role of GPC1 in concentrating virions at the cell surface prior to productive engagement of entry receptors. This surface concentration function, which increases the effective local virion density and residence time at the plasma membrane, enhances the probability of productive encounter with entry factors (such as the co-receptor axis represented by BiP and clusterin in the present study) and thus amplifies infectivity disproportionately to the absolute affinity of any individual contact. The functional complementarity between GPC1 (initial electrostatic surface concentration, consistent with HSPG-mediated attachment [9,179]), BiP (stress-induced co-receptor anchoring), and clusterin (extracellular chaperone-mediated stabilization) described across this study is consistent with a multi-stage, mechanistically layered surface-engagement process — though the precise contribution of each factor to productive viral entry requires experimental dissection.

##### 4.9.4 Structural Comparison Positions GPC1 as a High-Affinity Electrostatic Attachment Factor in the Host Network

Across the five complexes analyzed to this point — actin, betaglycan, BiP, clusterin, and GPC1 — the GPC1 interface is distinguished by three properties that collectively suggest it as a computationally supported high-affinity electrostatic attachment factor in the host protein network examined here. First, it generates the largest absolute EEL stabilization energy of any complex in the study, reflecting the highest density of charge–charge contacts at the interface. Second, it produces the most electrostatically saturated profile — where EEL and EGB are both maximized and their difference, rather than either term individually, defines the net affinity — a property that confers robustness against moderate changes in protonation state or ionic strength. Third, the interface residues responsible for the dominant favorable contributions (basic residues in the DIII patch) are among the most structurally conserved positions across DENV serotypes, suggesting that GPC1 recognition is broadly applicable across the serotype diversity examined in the genotype dataset of this study. This conservation stands in contrast with the betaglycan complex, where the electrostatic frustration at ARG107/LYS236 varies with genotype, and with the BiP complex, where the electrostatic anchor at ARG410 is preserved but the auxiliary hydrophobic contacts depend on local surface variability. Together, these properties are consistent with GPC1 functioning as a thermodynamically dominant, serotype-conserved, and kinetically efficient heparan sulfate proteoglycan attachment factor candidate [9,11,179]. As with other HSPGs in flavivirus biology, GPC1-mediated interaction likely serves to concentrate virions at the cell surface rather than to function as an entry receptor per se, and this structural characterization provides a computational baseline for understanding one layer of the multi-stage DENV–host recognition architecture.

##### 4.9.5 Ternary Landscape Identifies the GPC1–Env State as a Heparan-Sulfate-Dependent Basin Vulnerable to Growth-Factor Displacement

The conditional $\Delta\Delta G$ analysis situates the Glypican-1–Env interface within its native extracellular context by introducing two first-neighbour ligands — FGF1 and HBEGF — that are themselves well-established occupants of the heparan sulfate proteoglycan surface. The central result is that both partners selectively and substantially weaken the GPC1–Env energetic basin (AB) without comparably disrupting BC, and do so asymmetrically: the dominant perturbation occurs when the ligand is introduced into a pre-established GPC1–Env complex (AB-conditioned route), while Env arriving at a pre-occupied GPC1 surface encounters only a minor penalty. This conditional directionality is mechanistically distinct from the simple binary displacement and defines the GPC1–Env state as a metastable configuration susceptible to selective erosion by heparan-sulfate-dependent ligands already present in the pericellular environment.

The biological coherence of this result derives from the dual nature of GPC1's surface function. As a GPI-anchored glypican, GPC1 presents heparan sulfate chains at the cell surface and serves simultaneously as a ligand-capture platform and a signalling organiser — concentrating growth factors, regulating receptor proximity, and modulating local interaction geometry [96,99,167]. Dengue envelope interaction with GPC1 would compete for the same HS-decorated surface architecture that supports FGF and EGF-family signalling, making the observed AB destabilisation a structural-energetic manifestation of a genuine biological competition for glycocalyx occupancy rather than an artefact of the modelling framework.

##### 4.9.6 FGF1 Competition Reflects Canonical Heparan-Sulfate Co-Factor Displacement

The FGF1 result is mechanistically the most straightforward of the two cases. FGF1 is a founding member of the fibroblast growth factor family, and its signalling competence is obligatorily co-regulated by heparan sulfate proteoglycans: HS chains serve not merely as affinity cofactors but as structural organisers that stabilise the ternary FGF–FGFR–HS signalling complex and determine ligand presentation geometry [158]. In native cell-surface biology, GPC1 is among the best-characterised co-receptors for FGF signalling, presenting HS chains that concentrate and orient FGF ligands for FGFR engagement [95,99].

The $\Delta\Delta G \approx +2.8$ kcal · mol⁻¹ AB penalty observed when FGF1 is introduced into an established GPC1–Env complex is therefore consistent with FGF1 exerting a reorientation force on the GPC1-associated HS scaffold: the conformational and electrostatic requirements of productive FGF–GPC1 interaction differ from those sustaining GPC1–Env contact, and satisfying FGF1-specific binding geometry necessarily perturbs the Env-compatible GPC1 surface state. Critically, because BC is not substantially altered, FGF1 does not compete for direct protein–protein contact with Env; instead, it redirects the GPC1-associated HS microenvironment away from the configuration that supports the Env-containing assembly. This is consistent with competition for shared glycan scaffolding rather than steric exclusion at a protein-level interface.

##### 4.9.7 HBEGF Imposes a Stronger Displacement Through Its Dedicated Heparin-Binding Domain

HBEGF (Heparin-Binding EGF-Like Growth Factor) carries a discrete heparin-binding domain that directly engages heparan sulfate and has been shown to use cell-surface HSPGs as primary localisation and activity-modulating co-receptors [99,180]. Unlike some EGF-family members whose HS interactions are auxiliary, HBEGF's heparin-binding domain constitutes a structurally defined interaction surface whose occupancy of HS chains is intrinsic to its biological activity and spatiotemporal regulation [180].

The stronger AB penalty produced by HBEGF ($\Delta\Delta G \approx +3.7$ kcal · mol$^{-1}$) compared to FGF1 (+2.8 kcal · mol$^{-1}$) is therefore consistent with HBEGF imposing a more potent reorganisation of the GPC1-associated HS surface — an interpretation supported by the presence of a high-affinity, dedicated heparin-binding interface rather than the more diffuse HS co-factor dependency of FGF1. The near-unchanged BC ($\Delta\Delta G \approx -0.4$ kcal · mol$^{-1}$) further confirms that the HBEGF effect, like FGF1, operates through selective AB disruption rather than through competition for the Env-binding surface on GPC1.

Taken together, the HBEGF > FGF1 antagonism ranking reflects a structural principle: the magnitude of AB displacement correlates with the specificity and affinity of the third partner's intrinsic HS-binding architecture. Partners with dedicated heparin-binding domains impose larger AB penalties than those relying on HS as a diffuse co-factor.

##### 4.9.8 The GPC1–Env Basin as a Context-Dependent Glycocalyx Attachment Node

The convergent antagonism of both growth-factor partners, combined with the consistently near-neutral reverse-conditioned branch, supports a specific mechanistic model for GPC1-mediated dengue attachment. The GPC1–Env state is a thermodynamically favourable but conditionally labile configuration whose stability depends on the local availability of GPC1-associated HS chains in an Env-compatible binding geometry. This geometry can be selectively eroded by HS-dependent growth factors that pre-occupy or reorient the GPC1 HS scaffold without themselves disrupting the Env–neighbour interface.

The directionality of the effect — AB-conditioned destabilisation without reciprocal AB penalty when Env arrives at a pre-occupied surface — implies that the mode of competition is not simple mutual exclusion. Rather, the established GPC1–Env state is metastable and can be displaced by a subsequent HS-binding event, whereas Env encountering a GPC1 surface already occupied by FGF1 or HBEGF faces only a minor energetic barrier. Biologically, this asymmetry suggests that the effective availability of GPC1 for viral attachment is sensitive to prior loading of GPC1 HS chains by pericellular growth factors: microenvironments with high local FGF or HBEGF occupancy of GPC1 would reduce the persistence of productive GPC1–Env complexes even without directly targeting the viral protein.

This interpretation aligns with the broader view of heparan sulfate proteoglycans as dynamic, context-regulated components of the viral attachment landscape [97,99,167]. GPC1 is likely not a dedicated dengue receptor in the classical sense but rather a high-affinity glycocalyx node whose availability for viral attachment is determined by the competitive landscape of HS-binding ligands present in the extracellular environment at the time of viral engagement. The quantitative limits of this framework — particularly the absence of explicit HS chain modelling, glycan microheterogeneity, and membrane-anchored receptor clustering — mean these conclusions should be treated as structurally derived hypotheses requiring glycan-aware structural refinement and experimental validation in systems where GPC1 abundance and heparan sulfate editing can be independently controlled [41,47,49].

###### 4.9.9 GPC1-Mediated Dimensionality Reduction: From Bulk Diffusion to Membrane Surfing

The metastable, rapidly reversible character of the GPC1–Env interaction identified here acquires additional biological significance when interpreted through the framework of diffusion-limited receptor search on cell surfaces. Single-virion tracking studies of enveloped viruses have established that, upon initial contact with low-specificity surface attachment factors, viral particles do not immediately commit to endocytosis but instead undergo quasi-two-dimensional lateral diffusion along the plasma membrane — a behaviour first described as *virus-cell surfing* [181,182]. This kinetic mode reduces the effective dimensionality of receptor search from three dimensions (bulk diffusion in extracellular space) to two dimensions (lateral diffusion along the membrane plane), dramatically enhancing the probability of productive encounter with rare, geometrically constrained high-specificity receptors [183,184]. The classical theory of dimensionality-reduced diffusion predicts that two-dimensional membrane search is kinetically favourable by a factor proportional to the ratio of the membrane association rate to the three-dimensional collision frequency — an enhancement that can reach several orders of magnitude under physiological receptor density conditions [185,186].

The thermodynamic profile of the GPC1–Env interface provides a physical mechanism for this kinetic behaviour that goes beyond simple membrane tethering. The large oscillations in $\Delta G_{\text{total}}$ (up to 60 kcal/mol peak-to-trough in the trajectory window) observed in the GPC1 complex, while superficially appearing as instability, are the thermodynamic signature of an interaction dominated by individually weak but collectively numerous electrostatic contacts, each with a low activation barrier $\Delta G^{\ddagger}$ for formation and breaking. This is precisely the energetic landscape predicted by the *fly-casting* model of protein association: a disordered or multivalent binding surface extends a long-range electrostatic funnel that captures the partner from three-dimensional solution and guides it to the membrane surface, with the individual contact barriers being too small to trap the particle in any single configuration [187,188]. The consequence for dengue attachment is that the DIII electropositive patch does not "lock" the virion onto GPC1 — instead, it creates a low-barrier energy landscape in which the virus is attracted to the glycocalyx surface from a distance (fly-casting, enhancing the three-dimensional $k_{\text{on}}$) and then undergoes rapid bond-breaking and re-forming as it executes lateral two-dimensional diffusion (surf-riding, accelerating the two-dimensional search for NRP1 or other entry receptors).

The quantitative consequence of this two-step kinetic mechanism is substantial. Theoretical analyses of receptor search on cell surfaces estimate that dimensionality reduction from 3D to 2D can enhance the effective association rate constant for rare receptors by $10^2$–$10^4$-fold under conditions where the membrane-association step is faster than three-dimensional diffusion to the receptor site [183–185]. For NRP1, whose surface density on endothelial cells is estimated at $\sim 10^3$–$10^4$ copies per cell — approximately 100-fold lower than GPC1-associated heparan sulfate chains — this rate enhancement translates directly into a dramatic increase in productive NRP1 encounter probability per virion, without requiring either an increase in NRP1 expression or a decrease in three-dimensional diffusion barriers. This surf-riding model provides a structural and kinetic rationale for the functional hierarchy of dengue attachment that emerges from the comparative analysis of interaction landscapes across all host proteins: GPC1-mediated attachment constitutes the *dimensionality-reduction step*, concentrating virions at the cell surface and enabling two-dimensional search; NRP1 b1-pocket engagement constitutes the *high-specificity capture step*, geometrically restricting the viral particle in a pocket-insertion geometry that commits it to the endocytic pathway. The metastability of the GPC1–Env basin under FGF1/HBEGF competition — discussed above — is mechanistically coherent with this model: a surf-riding interaction must be labile enough to permit lateral diffusion, not a permanent anchor. The prediction that GPC1 acts as a dimensionality-reduction platform rather than a terminal receptor is directly testable by live-cell single-virion tracking experiments comparing dengue particle lateral diffusion coefficients and membrane dwell times in GPC1 knockout versus GPC1-overexpressing cell lines [181,189].

#### 4.10 Envelope–Neuropilin-1: CendR-Like Pocket Engagement, Vascular Co-Receptor Exploitation, and Mechanistic Integration

##### 4.10.1 CendR-Like Pocket Engagement as a Computationally Supported Structural Hypothesis for NRP1 Interaction

The structural and energetic evidence presented here is consistent with the hypothesis that the dengue envelope protein may engage NRP1 through a pocket-insertion geometry resembling that of CendR peptide ligands, in which a surface-exposed loop inserts into the canonical b1-domain binding pocket. Canonical CendR recognition requires a carboxy-terminal R/KXXR/K sequence, commonly exposed through proteolytic processing [102], which has been demonstrated experimentally for SARS-CoV-2 spike [19,103]. Importantly, the dengue envelope sequence does not contain a canonical C-terminal CendR motif, and the interaction should therefore be interpreted as a CendR-like pocket engagement geometry — a structural resemblance — rather than sequence-encoded CendR recognition. The energetic profile of the predicted interface — terminal basic residue paired with the NRP1 b1 acidic cage, augmented by hydrophobic packing of flanking residues — is qualitatively similar to characterized endogenous CendR–NRP1 complexes [108], but this similarity is observed in a static docking model and requires experimental confirmation through CendR peptide competition assays and structural characterization of the complex.

This constitutive accessibility has significant biological implications. If the dengue envelope loop is available for NRP1 engagement in both the pre-fusion and post-fusion states, then NRP1 could potentially function at multiple stages of the viral lifecycle — during initial virion capture at the endothelial surface, during intracellular trafficking, and following egress of newly produced virions. The direct MD evidence for stable, pocket-inserted engagement without major conformational requirements on the envelope side is consistent with this multi-stage accessibility model and distinguishes NRP1 from co-receptors (such as BiP) whose engagement depends on stress-induced changes in receptor availability.

##### 4.10.2 Energetic Parsimony of CendR Recognition: Geometric Complementarity over Electrostatic Brute Force

The MM/GBSA energetic profile of the NRP1 complex differs fundamentally from that of the GPC1 interface in its underlying mechanism of stability. Whereas GPC1-mediated stability emerges from electrostatic saturation — the near-cancellation of very large EEL and EGB terms — NRP1-mediated stability reflects geometric complementarity: the b1-pocket geometry precisely accommodates the terminal loop of the viral surface, generating a net free energy that is comparatively less sensitive to individual charge–charge fluctuations and more dependent on the persistent steric fit between the pocket and the inserted residues. This energetic parsimony is evidenced by the smaller amplitude oscillations in $\Delta G_{\text{total}}$ observed for NRP1 relative to GPC1, and by the more prominent VDWAALS contribution relative to EEL, indicating that van der Waals packing rather than electrostatic complementarity constitutes the primary geometric driver.

This distinction has practical consequences for understanding the molecular basis of host-specificity and serotype variation in NRP1-mediated dengue attachment. Interactions dominated by geometric complementarity — where the steric fit of a defined loop length and sequence into a rigid pocket is the principal determinant of binding — are expected to be more sensitive to mutations in the loop residues and less sensitive to changes in the electrostatic surface far from the contact site. This selectivity profile is precisely what would be expected of a CendR-type interaction, where the terminal di-residue motif (R/KXXR/K) is obligatory and non-conservative substitution abolishes binding [102,103]. For dengue, this implies that serotype variation in the CendR-like loop sequence would directly modulate NRP1 affinity, providing a structural hypothesis for the observed differences in endothelial tropism and vascular permeability induction across DENV serotypes [104].

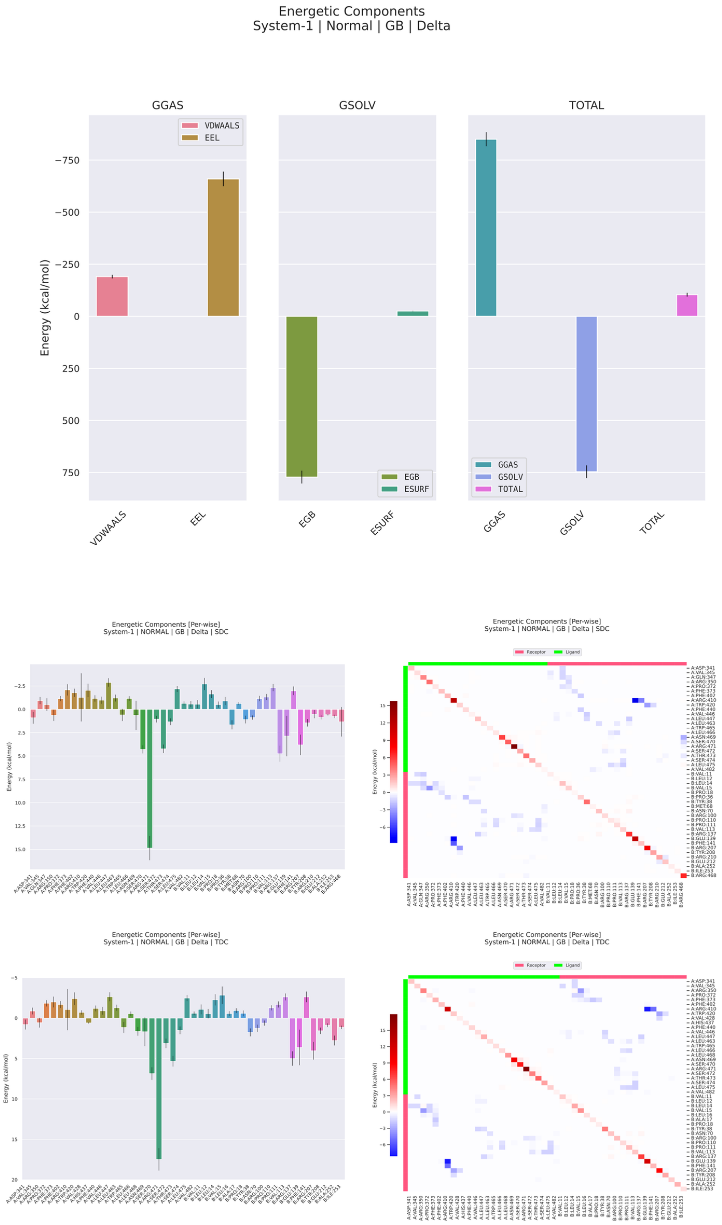


**Figure 19: MM/GBSA energetic decomposition of the dengue virus Envelope–neuropilin-1 complex during molecular dynamics simulation.** The figure summarizes the global energetic components, per-residue decomposition, and pairwise residue decomposition of the binding free energy estimated by MM/GBSA for the dengue Envelope–neuropilin-1 complex. The upper panels show that the gas-phase contribution is strongly favorable and is dominated by the electrostatic term, whereas the van der Waals component provides an additional, smaller favorable contribution. This profile indicates that the Envelope–neuropilin-1 association is largely driven by charge-dependent intermolecular complementarity, with short-range packing interactions acting as a secondary stabilizing factor. In contrast, the solvation contribution is strongly unfavorable and is dominated by the polar generalized Born term, indicating a substantial desolvation penalty associated with burial or rearrangement of charged and polar groups at the interface. The nonpolar surface-area term contributes only modestly relative to these dominant energetic components. Despite the pronounced compensation between favorable gas-phase attraction and unfavorable polar solvation, the total binding free energy remains favorable, supporting the energetic stability of the dengue Envelope–neuropilin-1 complex after solvent effects are taken into account. The middle and lower panels show per-residue and pairwise decompositions using SDC and TDC schemes, revealing a markedly heterogeneous interfacial architecture in which most residues contribute only weakly, whereas a restricted subset of residues concentrates the strongest stabilizing or destabilizing effects. Negative values indicate favorable contributions to binding, whereas positive values indicate unfavorable or destabilizing energetic terms. In both decomposition schemes, the most prominent energetic peaks are concentrated in a limited number of residues, indicating that the interface is organized around discrete energetic hotspots rather than a diffuse and uniformly distributed contact surface. The pairwise heatmaps further support this interpretation by showing sparse but well-defined favorable residue-pair interactions embedded within a broader background of weak or neutral contacts, together with a smaller number of unfavorable pairwise contributions. Blue regions denote favorable residue-pair interactions, whereas red regions denote unfavorable or destabilizing contribu- [illegible]

###### 4.10.3 NRP1 as a Vascular Amplifier of Dengue Pathogenesis: Pathological Receptor Hijacking

Beyond its structural role as a virion attachment co-receptor, NRP1 engagement by the dengue envelope has the potential to directly amplify the vascular pathology that defines severe dengue disease. NRP1 functions as a co-receptor that presents VEGF-A to VEGFR2, enhancing downstream phosphorylation of tyrosine residues in the VEGFR2 kinase domain and potentiating signaling through the PI3K/Akt and eNOS pathways that regulate endothelial junction permeability [104,105]. The competitive occupancy of the NRP1 b1 pocket by the dengue envelope loop — a pocket that constitutively mediates VEGF-A presentation in endothelial cells — could disrupt the VEGF–NRP1–VEGFR2 ternary complex in two non-exclusive ways: by sterically blocking VEGF-A binding to the b1 domain and thereby reducing NRP1-assisted VEGFR2 activation, or by inducing a conformational state in NRP1 that preferentially signals through b1-independent vascular permeability pathways. Both mechanisms would be predicted to alter endothelial barrier function in the infected vasculature, consistent with the clinical hallmark of dengue hemorrhagic fever — plasma leakage through a compromised endothelial barrier — which occurs in the absence of overt endothelial cell death and therefore reflects functional rather than structural disruption of the endothelial junction architecture.

This mechanism represents a qualitatively distinct pathological strategy that merits a specific designation: *pathological receptor hijacking*. Unlike classical receptor-mediated viral entry — in which the virus exploits a receptor solely as a gateway, leaving the receptor's downstream signalling largely intact once the virus has been internalised — pathological receptor hijacking involves the chronic occupation of a signalling receptor's ligand-binding pocket in a manner that simultaneously achieves attachment and corrupts the receptor's physiological function. The dengue NRP1 interaction satisfies both criteria: the b1 pocket engaged by the viral loop is the same pocket that presents VEGF-A165 for VEGFR2 co-activation, and the structural analysis predicts that viral pocket occupancy is constitutive (independent of prior proteolytic processing, unlike SARS-CoV-2 CendR engagement) and maintained on the same timescale as viral attachment and endocytosis [19,104].

The physiological consequences of chronic NRP1 b1-pocket occupancy during dengue infection are directly linked to the progressive plasma leakage that distinguishes dengue hemorrhagic fever from uncomplicated dengue fever. VEGF-A–NRP1–VEGFR2 signalling through the eNOS/Akt axis maintains endothelial tight junction integrity by sustaining VE-cadherin phosphorylation and cortical actin organisation; disruption of this axis is sufficient to induce reversible vascular hyperpermeability in the absence of cell death — the precise phenotype of dengue plasma leakage [22,104]. If NRP1 b1-pocket occupancy by the envelope protein suppresses VEGF-A-mediated VEGFR2 co-activation during the critical viremic phase, the predicted consequence is a quantitative reduction in endothelial survival and barrier maintenance signalling, creating the permissive vascular state for plasma extravasation that characterises severe disease. This mechanistic chain — from b1-pocket engagement to VEGFR2 co-activation deficit to VE-cadherin destabilisation to plasma leakage — is structurally grounded in the interaction geometry characterised here and experimentally testable through VEGFR2 phosphorylation assays in NRP1-wild-type versus NRP1-b1-mutant endothelial cells under dengue infection conditions.

This vascular amplification mechanism operates at the interface between viral attachment and host pathophysiology in a manner distinct from all other complexes analyzed in this study. Actin engagement disrupts cytoskeletal integrity; betaglycan engagement hijacks TGF-$\beta$ co-receptor signaling; BiP engagement exploits ER stress for cell-surface exposure; clusterin engagement co-opts proteostasis clearance for endocytic entry; and GPC1 engagement concentrates virions at the surface. NRP1 engagement, by contrast, simultaneously achieves attachment and vascular dysfunction through a single molecular event — competitive displacement of an endogenous signaling ligand from a receptor whose physiological function is directly coupled to endothelial permeability. This dual-function exploitation positions NRP1 as mechanistically unique within the host network and suggests that NRP1 blockade could simultaneously inhibit viral attachment and prevent VEGF-A displacement, offering a potential therapeutic strategy with dual antiviral and vascular-protective effects.

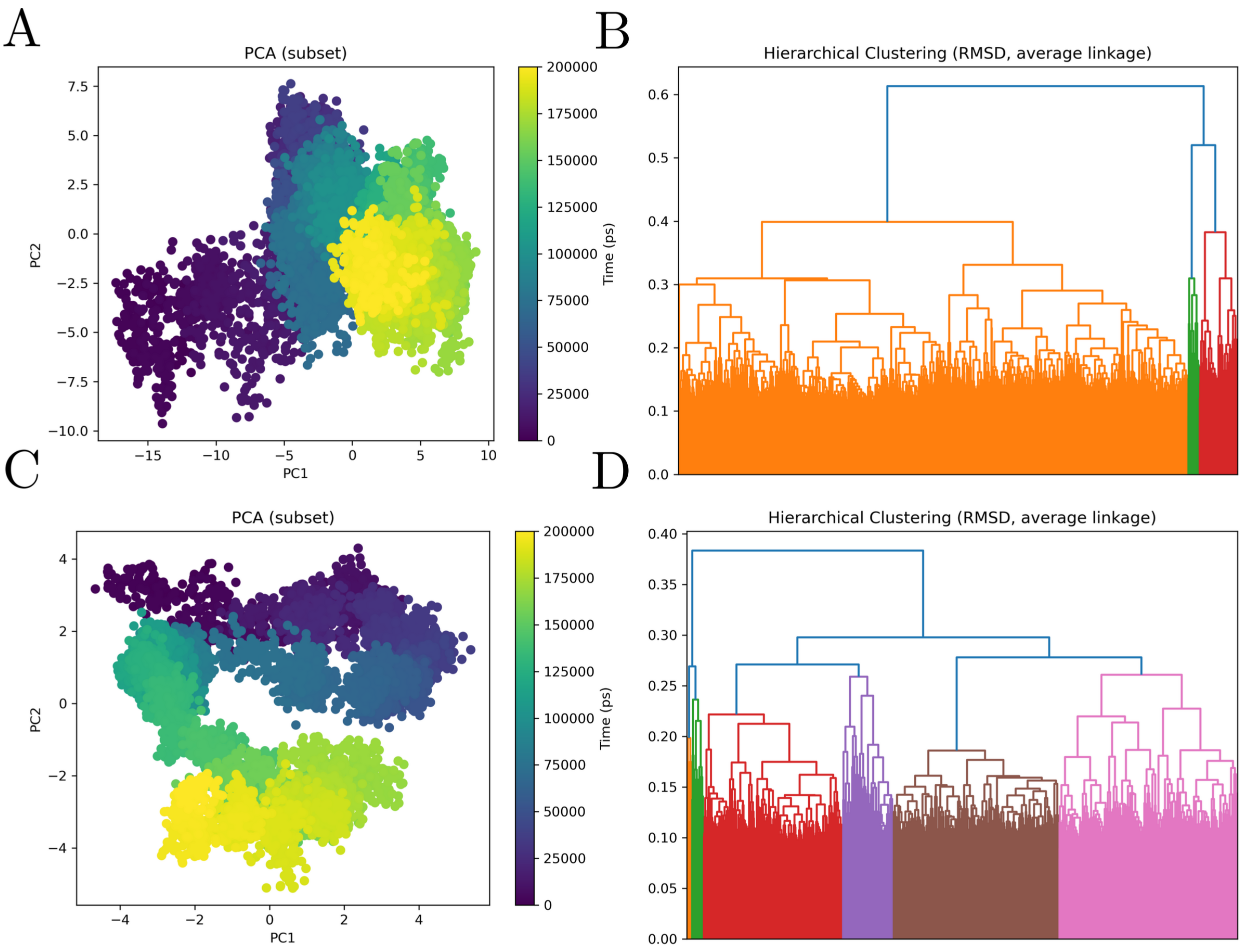


**Figure 20: Principal component analysis and RMSD-based hierarchical clustering of the dengue Envelope–neuropilin-1 molecular dynamics trajectory.** The figure summarizes the conformational landscape sampled during the molecular dynamics simulation of the dengue Envelope–neuropilin-1 complex, with panels A and B corresponding to the receptor subset, dengue Envelope, and panels C and D corresponding to the ligand subset, neuropilin-1. Panels A and C show principal component analysis projections of trajectory frames onto the first two principal components, with points colored according to simulation time. In panel A, the dengue Envelope subset displays a broad and directional conformational redistribution, progressing from an early region at strongly negative PC1 values toward intermediate and late conformational regions located closer to central and positive PC1 values. This temporal progression indicates that the receptor does not remain confined to a single equilibrium basin, but instead undergoes substantial structural drift during the simulation. The partial continuity between early, intermediate, and late regions suggests a progressive conformational transition, compatible with adaptation of the Envelope within the neuropilin-1-bound state. The late-stage frames occupy a more compact region relative to the widely dispersed early ensemble, suggesting partial stabilization after the initial conformational exploration. In panel C, the neuropilin-1 subset shows a distinct and highly organized temporal trajectory. Early frames occupy an upper region of the PCA space, whereas intermediate and late frames progressively redistribute toward lower PC2 values, forming a curved conformational path rather than a random dispersion of structures. This pattern indicates that neuropilin-1 undergoes substantial ligand-associated accommodation during the simulation. The separation between early and late regions is especially clear for the ligand subset, suggesting that neuropilin-1 contributes actively to the conformational evolution of the complex rather than behaving as a passive binding surface. The ordered redistribution across the PCA plane is consistent with progressive remodeling of the ligand around the Envelope-associated interface, followed by sampling of more persistent late-stage conformational states. Panels B and D show RMSD-based hierarchical clustering using average linkage, revealing multiple structural families within both receptor- and ligand-associated subsets. The receptor dendrogram in panel B is dominated by a large conformational family accompanied by smaller, structurally distinct clusters at higher RMSD distances. This pattern supports the interpretation that dengue Envelope samples a major conformational basin while also accessing subsidiary substates during the trajectory. The ligand dendrogram in panel D shows a more subdivided clustering structure, with several well-resolved conformational groups distributed across the trajectory. This indicates that neuropilin-1 samples multiple structurally distinguishable states, consistent with the more organized but heterogeneous conformational redistribution observed in the PCA projection. Together, the PCA and clustering analyses indicate that the dengue Envelope–neuropilin-1 complex is dynamically heterogeneous but structurally organized, with both partners undergoing time-dependent conformational redistribution. Overall, these results support a model in which neuropilin-1 binding is associated with progressive remodeling of the Envelope-containing complex, generating an ensemble of related conformational states rather than [illegible]

###### 4.10.4 Structural Comparison with SARS-CoV-2 NRP1 Engagement: Convergent Pocket Geometry

The structural parallel between the dengue envelope–NRP1 interaction predicted here and the SARS-CoV-2 spike–NRP1 interaction established experimentally [19,103] raises the possibility that structurally similar b1-pocket engagement may occur across phylogenetically distant viruses sharing endothelial tropism. This remains a computationally derived hypothesis for dengue and does not imply the same mechanistic pathway as the experimentally validated SARS-CoV-2 case. In both cases, the viral surface presents a basic residue in a terminal or loop context that is captured by the NRP1 b1 acidic cage; in both cases, the interaction is augmented by hydrophobic packing of flanking residues; and in both cases, the energetic profile reflects geometric complementarity rather than bulk electrostatic capture. The convergence extends to functional consequences: SARS-CoV-2 NRP1 engagement amplifies viral entry into endothelial cells and neurons [19], while dengue NRP1 engagement — as characterized here computationally — is positioned to amplify both viral internalization and VEGF-A pathway disruption. The quantitative differences between the two interactions (principally in the absolute magnitude of EEL and in the relative contribution of VDWAALS) likely reflect the difference between a post-proteolytic disordered C-terminal tail (SARS-CoV-2) and an ordered structural loop (DENV), a distinction that is architecturally consequential but does not alter the core CendR recognition chemistry.

These structural parallels suggest NRP1 as a candidate host interaction point warranting investigation across flaviviruses, and motivate experimental testing of whether NRP1-targeted compounds in development for SARS-CoV-2 might affect dengue infection. Direct experimental validation of the proposed dengue ENV–NRP1 interaction is required before mechanistic or therapeutic conclusions can be drawn.

###### 4.10.5 The NRP1 b1 Domain as a Universal Vascular Vulnerability: Convergent Structural Evolution Among Hemorrhagic Pathogens

The independently derived b1-pocket engagement geometries of DENV and SARS-CoV-2 invite a broader evolutionary interpretation that extends beyond the specific comparison of these two viruses. The NRP1 b1 acidic cage — a structurally conserved, negatively charged shallow binding pocket whose physiological function is to capture the C-terminal basic residues of VEGF-A splice variants and proteolytically processed CendR ligands [102,108,190] — appears to constitute what may be termed a *topological vulnerability* of the endothelial cell surface: a receptor pocket whose geometric properties (defined cavity, predictable charge distribution, shallow but specific complementarity) are discoverable by structurally diverse pathogens through independent evolutionary trajectories.

Convergent evolution at the level of protein–protein interaction geometry, rather than sequence, has been documented for viral proteins exploiting host signalling surfaces across multiple pathogen lineages. Short linear motif (SLiM)-based molecular mimicry — in which viral surface loops adopt the geometry of cellular SLiMs to engage host receptor grooves without sharing their primary sequence — has been proposed as a pervasive strategy enabling phylogenetically distant viruses to capture the same host interaction interfaces [148,191,192]. The dengue and SARS-CoV-2 NRP1 interactions represent a compelling candidate for this class of convergent geometric mimicry: two RNA viruses from entirely different families (*Flaviviridae* and *Coronaviridae*), with no structural homology in their surface glycoproteins, independently evolved surface features that engage the same b1-domain geometry through energetically similar but architecturally distinct contact modes — an ordered structural loop (DENV) versus a post-proteolytic disordered C-terminal tail (SARS-CoV-2) [19,103].

Extending this logic, the NRP1 b1 pocket represents a testable candidate for engagement by other hemorrhagic or vascular-tropic pathogens. Flaviviruses sharing dengue's endothelial tropism — Zika virus, West Nile virus, and Yellow fever virus — present related but not identical DIII surface geometries, providing a panel for testing whether NRP1 b1-pocket engagement is a shared flavivirus strategy or is specific to DENV serotypes. Hantaviruses, which cause hemorrhagic fever with renal syndrome and hantavirus pulmonary syndrome through endothelial dysfunction, and Ebola virus, whose glycoprotein engages multiple endothelial attachment receptors during disseminated infection, represent phylogenetically independent test cases for convergent NRP1 exploitation [193,194]. Should experimental evidence establish NRP1 b1-pocket engagement across multiple hemorrhagic pathogen lineages, the mechanistic implication would be that endothelial barrier disruption by these pathogens converges, at least in part, on a common structural node — the VEGF-A co-presentation machinery — through independently evolved molecular solutions.

This framing has direct therapeutic consequences. A small molecule or peptide mimetic designed to occupy the NRP1 b1 acidic cage — blocking its availability to any viral surface loop — would constitute a *pan-vascular-tropism* entry inhibitor whose spectrum of coverage is determined not by conservation of the viral ligand but by the conservation of the host receptor pocket. Structural studies of the NRP1 b1 domain in complex with CendR peptides have demonstrated that the pocket tolerates considerable ligand diversity while maintaining a core recognition geometry centered on the terminal basic residue pair [102,108], supporting the tractability of pocket-occupying inhibitor design. NRP1 b1-targeted compounds currently in development for anti-angiogenic oncology applications [195] and for SARS-CoV-2 prophylaxis represent ready-made candidates for repurposing evaluation against dengue and other flaviviruses — a translational opportunity that the convergent structural evidence presented here directly motivates.

###### 4.10.6 Ternary Landscape Positions NRP1–Env within a Plexin-Dominated Signalling Architecture

The conditional $\Delta\Delta G$ analysis embeds the NRP1–Env interface within its native signalling network and reveals a heterogeneous but mechanistically coherent hierarchy of perturbations. The stable AB baseline ($\Delta G \approx -12.8$ kcal · mol$^{-1}$) coexists with third-partner BC energies spanning a wide range ($\approx -10$ to $-20$ kcal · mol$^{-1}$), reflecting the established role of NRP1 as a multifunctional receptor that dynamically assembles with ligands, co-receptors, and intracellular scaffolds across several signalling pathways [105,106]. The key finding is that NRP1 signalling partners do not uniformly destabilise the NRP1–Env state; rather, they produce a spectrum from strong cooperative reinforcement (PLXNA1, SEMA4) to strong competitive antagonism (PLXNA4), with intracellular scaffolds and immunoglobulin-like partners occupying the moderate range. This structured heterogeneity establishes NRP1 not as a fixed, context-independent viral receptor but as an interaction node whose accessibility to the viral envelope depends on the local composition of the NRP1 signalling complex.

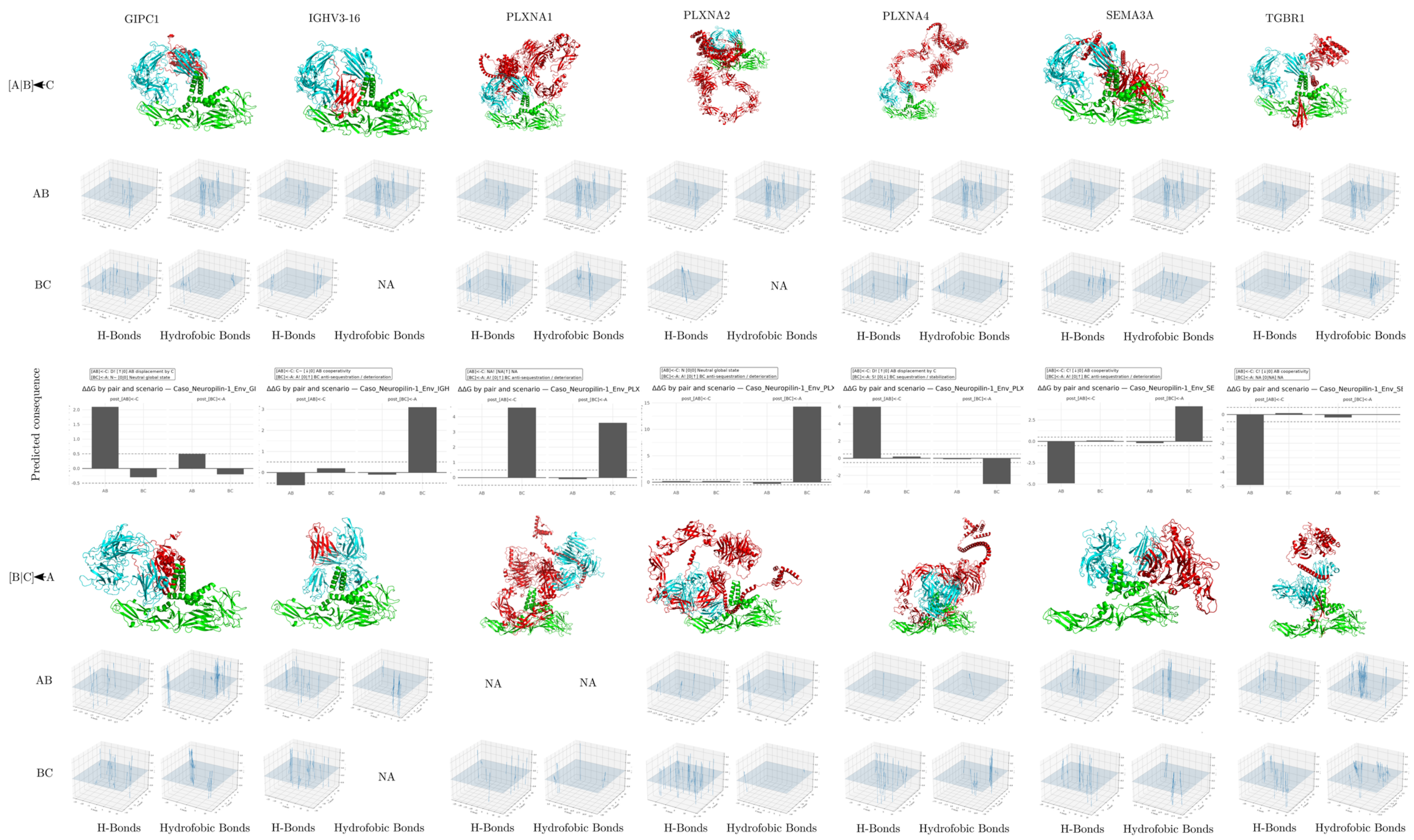

GIPC1
IGHV3-16
PLXNA1
PLXNA2
PLXNA4
SEMA3A
TGBR1
[A|B]◄C
AB
BC
NA
H-Bonds
Hydrofobic Bonds
Predicted consequence
ΔΔG by pair and scenario — Caso_Neuropilin-1_Env_GI
ΔΔG by pair and scenario — Caso_Neuropilin-1_Env_IGH
ΔΔG by pair and scenario — Caso_Neuropilin-1_Env_PLX
ΔΔG by pair and scenario — Caso_Neuropilin-1_Env_PLX
ΔΔG by pair and scenario — Caso_Neuropilin-1_Env_PLX
ΔΔG by pair and scenario — Caso_Neuropilin-1_Env_SE
ΔΔG by pair and scenario — Caso_Neuropilin-1_Env_SI
[B|C]◄A
AB
BC
NA
H-Bonds
Hydrofobic Bonds

**Figure 21. Structural, interaction-field, and predicted energetic consequences of dengue Envelope–neuropilin-1-centered ternary complexes with neuropilin-1-associated proteins.**

The figure summarizes comparative ternary interaction models involving dengue virus Envelope, neuropilin-1, and seven neuropilin-1-associated partners: GIPC1, IGHV3-16, PLXNA1, PLXNA2, PLXNA4, SEMA3A, and TGFBR1. In this representation, A corresponds to dengue Envelope, B corresponds to neuropilin-1, and C corresponds to the additional neuropilin-1-associated protein. Two alternative assembly scenarios are compared. In the first scenario, the Envelope–neuropilin-1 pair is treated as the preformed binary complex and the third protein is introduced afterward, represented as $[A]B \leftarrow C$. In the reciprocal scenario, the neuropilin-1–associated-protein pair is treated as preformed and dengue Envelope is introduced as the third component, represented as $[B]C \leftarrow A$.

Structural panels show the relative spatial organization of the three proteins in each modeled ternary assembly, revealing pronounced partner-dependent differences in geometry and packing. Interaction-field plots report the spatial distribution of hydrogen-bond and hydrophobic contact features for the AB and BC interfaces, showing that polar and nonpolar interaction densities are redistributed according to both partner identity and assembly order. Predicted energetic consequence plots report $\Delta\Delta G$ values for each pair and assembly scenario, with negative values indicating stabilizing effects and positive values indicating destabilizing or energetically unfavorable effects relative to the corresponding reference state. Missing descriptors or unavailable configurations are indicated as NA.

Overall, the figure indicates that the dengue Envelope–neuropilin-1 interaction is strongly modulated by the local molecular neighborhood of neuropilin-1. Some neighbors, such as GIPC1 and PLXNA4, preferentially perturb the AB interface when added to a preformed Envelope–neuropilin-1 complex, whereas others, including IGHV3-16, PLXNA1, PLXNA2, and SEMA3A, impose stronger penalties on the BC interface when dengue Envelope is introduced into preformed neuropilin-1-associated assemblies. This reciprocal behavior supports a path-dependent model of ternary complex formation, in which the structural and energetic outcome depends not only on the identity of the interacting proteins, but also on the order in which the viral and host components assemble.

Together, these results suggest that neuropilin-1 functions as a network-embedded host interface whose recognition by dengue Envelope is conditioned by its surrounding signaling and adaptor-protein context.

##### 4.10.7 Plexin Paralogue-Specific Effects Define the Dominant Competitive Axis

The three Plexin-A family members tested generate markedly different conditional signatures despite sharing the same broad receptor family, revealing a high degree of paralog-specificity in NRP1 ternary interaction geometry. PLXNA1 and NRP1 are well-established components of the canonical Class 3 semaphorin co-receptor complex, and structural studies have shown that this assembly involves extensive NRP1–Plexin extracellular domain contacts that position the Plexin ectodomain relative to the NRP1 b1 domain [106,107]. The extraordinary $\Delta\Delta G \approx -12.8$ kcal · mol$^{-1}$ cooperative deepening of AB under PLXNA1 conditioning — the largest single cooperative signature in the entire NRP1 dataset — indicates that the NRP1 surface configuration stabilised by PLXNA1 assembly is substantially more compatible with Env engagement than the isolated NRP1 reference state. This is not biologically counterintuitive: if PLXNA1 constrains NRP1 in a defined extracellular conformation, and if that conformation exposes or pre-organises the b1 pocket that Env engages, then PLXNA1-dependent NRP1 conformational selection would directionally favour Env binding. The simultaneous BC destabilisation ($\Delta\Delta G \approx +4.6$ kcal · mol$^{-1}$) is also mechanistically significant: it suggests that within the PLXNA1-conditioned NRP1 conformation, the Env-facing surface is partially sequestered from alternative neighbour contacts, consistent with a receptor state that is cooperatively optimised for viral rather than endogenous ligand engagement.

PLXNA4, by contrast, is a Plexin paralog whose NRP1 interaction involves a distinct binding geometry and that mediates distinct biological functions, including repulsive semaphorin signalling in vascular and immune contexts [107]. The $\Delta\Delta G \approx +6.0$ kcal · mol$^{-1}$ AB penalty imposed by PLXNA4 without BC disruption indicates selective competitive antagonism against the NRP1–Env state — likely reflecting that PLXNA4 stabilises an NRP1 extracellular conformation that is geometrically incompatible with the Env b1-pocket engagement geometry. PLXNA2 is effectively permissive ($|\Delta\Delta G| \leq 0.3$ kcal · mol$^{-1}$), indicating that not all Plexin-A paralogs engage NRP1 surface determinants that overlap with the Env-binding region, and that the observed paralog-specific effects reflect genuine structural differences in NRP1 extracellular domain engagement rather than a generic Plexin-class effect.

##### 4.10.8 Semaphorin Engagement Produces a Cooperative Configuration Compatible with Viral Binding

SEMA4 produces the second-largest cooperative signature ($\Delta\Delta G \approx -4.9$ kcal · mol$^{-1}$) and is mechanistically the most biologically unexpected result in the NRP1 panel. Semaphorins are the primary endogenous ligands of Neuropilins, and semaphorin–NRP1 interactions are understood to drive receptor clustering and downstream Plexin-mediated signalling [105,107]. The observation that SEMA4 deepens rather than displaces the NRP1–Env basin is consistent with semaphorin-induced NRP1 conformational states — likely involving receptor clustering or localised b1-domain repositioning — that remain structurally compatible with the viral envelope pocket-insertion geometry. Given that NRP1 clustering is an established consequence of semaphorin engagement, one mechanistic interpretation is that semaphorin-driven receptor aggregation increases the local density of b1-pocket configurations accessible to Env, effectively creating a high-avidity attachment surface. This speculative model would predict that semaphorin-rich microenvironments could paradoxically enhance viral attachment to NRP1 rather than compete with it, a hypothesis distinct from the straightforward competitive antagonism expected of most endogenous ligands and one that merits targeted experimental investigation.

###### 4.10.9 GIPC1 Exerts Indirect Antagonism Through Receptor Trafficking Regulation

GIPC1 (also known as GAIP-interacting protein C terminus) is a PDZ-domain scaffolding protein that binds the cytoplasmic C-terminal SEA motif of NRP1 and regulates receptor internalisation, trafficking velocity, and membrane density at the plasma membrane [107]. Its moderate AB destabilisation ($\Delta\Delta G \approx +2.1$ $\mathrm{kcal \cdot mol^{-1}}$) under AB conditioning is unlikely to arise from extracellular steric competition, since GIPC1 engages the intracellular NRP1 domain rather than the extracellular b1-b2-a1-a2 ectodomain region where Env interacts. The energetic perturbation likely therefore reflects an indirect structural effect: GIPC1-mediated receptor scaffolding imposes conformational or orientational constraints on the NRP1 transmembrane and juxtamembrane regions that propagate to the ectodomain conformation, indirectly reducing compatibility with the Env-binding state. This indirect modality is distinct from the extracellular competition characterising Plexin and semaphorin effects and highlights that intracellular NRP1 regulatory proteins can influence viral attachment through receptor presentation mechanisms rather than through direct competition at the viral interface. Biologically, this suggests that the efficiency of NRP1-mediated viral engagement may be regulated by the availability and activity of NRP1 trafficking machinery, particularly under conditions of ER stress or GIPC1 redistribution that accompany dengue infection.

###### 4.10.10 Immunoglobulin-Domain and TGF-$\beta$ Axis Partners Are Marginally Perturbing

IGHV3-16 and TGFBR1 produce the smallest non-neutral effects in the NRP1 panel. The weak cooperative signal from IGHV3-16 ($\Delta\Delta G \approx -0.6$ $\mathrm{kcal \cdot mol^{-1}}$) is consistent with an immunoglobulin variable domain engaging a surface region of NRP1 or Env that is adjacent to rather than directly overlapping with the b1-pocket viral interface, a binding geometry that would marginally stabilise the NRP1–Env state through steric complementarity rather than direct competition. This interpretation is consistent with observations from the Betaglycan ternary panel, where immunoglobulin-like partners also produced context-specific cooperative signatures reflecting surface complementarity rather than allosteric communication [156,157].

The TGFBR1 result ($\Delta\Delta G \approx +0.8$ kcal · mol$^{-1}$) is mechanistically coherent with the known biology of NRP1–TGFBR1 crosstalk, which operates through indirect receptor complex modulation rather than a stable binary extracellular NRP1–TGFBR1 interface [140,144]. The small energetic perturbation indicates that TGFBR1 does not strongly compete for the NRP1 extracellular surface under the conditions modelled here, consistent with the predominantly intracellular and indirect nature of NRP1–TGF-$\beta$ signalling crosstalk.

###### 4.10.11 A Receptor-State Model for NRP1-Mediated Viral Engagement

Collectively, the NRP1 ternary results support a receptor-state model in which the stability of the NRP1–Env attachment interface is determined not by intrinsic viral or receptor affinity alone but by the NRP1 extracellular conformational state imposed by its endogenous signalling partners. PLXNA1 and SEMA4 define cooperative states in which NRP1 receptor organisation is compatible with or actively favours Env pocket-insertion; PLXNA4 defines an antagonistic state in which an alternative Plexin-mediated NRP1 conformation precludes Env engagement; PLXNA2, IGHV3-16, and TGFBR1 define permissive or marginally perturbing states; and GIPC1 illustrates indirect regulation through intracellular receptor trafficking machinery.

This model predicts that dengue viral attachment to NRP1 would be spatially and temporally heterogeneous at the cell surface, preferentially occurring at sites of PLXNA1-associated NRP1 receptor clusters or semaphorin-induced NRP1 condensates rather than at sites where PLXNA4 has imposed an incompatible receptor conformation. Because endothelial cells and neurons — the primary vascular and neurological targets of dengue pathology — express different complements of Plexin paralogs and semaphorins, this model further predicts that NRP1-mediated attachment efficiency would differ between cell types in a receptor-network-composition-dependent manner, providing a structural basis for the tissue tropism dimension of dengue pathology [19,22].

As with all conditional $\Delta\Delta G$ analyses presented here, these conclusions are bounded by the docking methodology, which does not resolve membrane-anchored receptor oligomerisation, cytoplasmic domain allosteric propagation, glycosylation patterns, or the temporal kinetics of receptor complex formation. Experimental validation — particularly NRP1 engagement assays in the presence of PLXNA1 versus PLXNA4, and semaphorin competition or enhancement assays — represents the highest-priority next step for this system [41,47,49].

#### 4.11 Envelope–Rab5: Endosomal Machinery Recruitment Through Directed Interface Consolidation

##### 4.11.1 The Two-Registry Mechanism: Global Reorientation with Continuous Local Anchoring

The structural data presented above establish a mechanistic framework for the Rab5–envelope interaction that has no direct precedent among the other host protein complexes analyzed in this study. The defining feature — a global center-of-mass separation that increases while the minimum interchain distance remains constant and the number of contacts grows — is the structural hallmark of a reorientation-driven interface consolidation, in which the two proteins pivot around a continuously maintained anchor set to redistribute contacts across a larger interface area without ever losing their initial connection. This mechanism is conceptually analogous to the "encounter complex to specific complex" transition well characterized in the kinetics of protein–protein association, where diffusional encounter geometries are iteratively filtered toward the thermodynamically optimal registered state through a series of partial dissociation–re-engagement events [100,196]. In the Rab5 simulation, however, the transition appears to occur without detachment — the minimum distance trajectory provides no evidence of even transient dissociation — suggesting that the initial contact is sufficiently strong to maintain continuous engagement while the global geometry optimizes, a scenario consistent with a rugged but connected energy landscape in which the transition from the early to the late registry follows a path of monotonically decreasing free energy.

The directed nature of the PC1 transition — with no backward revisitation of the early conformational basin after the transition is complete — implies that the late registry represents a thermodynamic attractor that is kinetically irreversible on the 200 ns simulation timescale. Whether this represents a true global free energy minimum or a deep local minimum separated from the early registry by a negligible barrier cannot be conclusively determined from a single 200 ns trajectory; however, the convergence of RMSD, Rg, contact number, and MM/GBSA metrics all toward more favorable values in the late-time regime is consistent with the late registry being the thermodynamically preferred state under the force field and simulation conditions employed.

##### 4.11.2 Mechanistic Division of Labor: VDW Scaffold and Electrostatic Microstate Tuning

The MM/GBSA decomposition introduces a conceptual distinction that is absent from all other complexes analyzed in this study: the explicit separation of a **structural scaffold** (the VDWAALS component, which drifts gradually toward more negative values as the interface consolidates and is relatively insensitive to instantaneous contact geometry) from an **electrostatic modulator** (the EEL component, which fluctuates sharply and generates the transient deepening events in $\Delta G_{\text{total}}$). This distinction is mechanistically meaningful because it maps directly onto the two-domain architecture of the Rab5 surface: the GTPase core provides a rigid, geometrically well-defined surface whose engagement with the envelope generates shape-complementary van der Waals contacts that improve monotonically as the interface consolidates, while the switch regions — whose charged residues (Asp, Thr, Glu in switch I/II) mediate state-dependent effector selectivity — generate the volatile electrostatic contacts that produce the instantaneous energy fluctuations [116].

The "anchor + frustration" pattern identified in the SDC decomposition — GLU133 as a persistent stabilizing anchor, ARG91 as a frustrated peripheral residue — provides a residue-level explanation for this division. The GLU133 anchor likely corresponds to one of the "pivot" contacts maintained throughout the reorientation, contributing a favorable and relatively constant electrostatic contribution that anchors the early-to-late registry transition. ARG91's frustration, by contrast, identifies a contact site whose geometry is not fully optimized in the late-time registry sampled by the simulation, suggesting that a further conformational refinement — beyond the timescale accessible to the 200 ns trajectory — would be required to fully compensate its desolvation cost. In the physiological context, this refinement might be provided by membrane proximity, GTP hydrolysis state change, or effector co-occupancy of the Rab5 surface, all of which modulate the charge distribution around the switch regions and could shift the interface toward a fully compensated geometry [110,119].

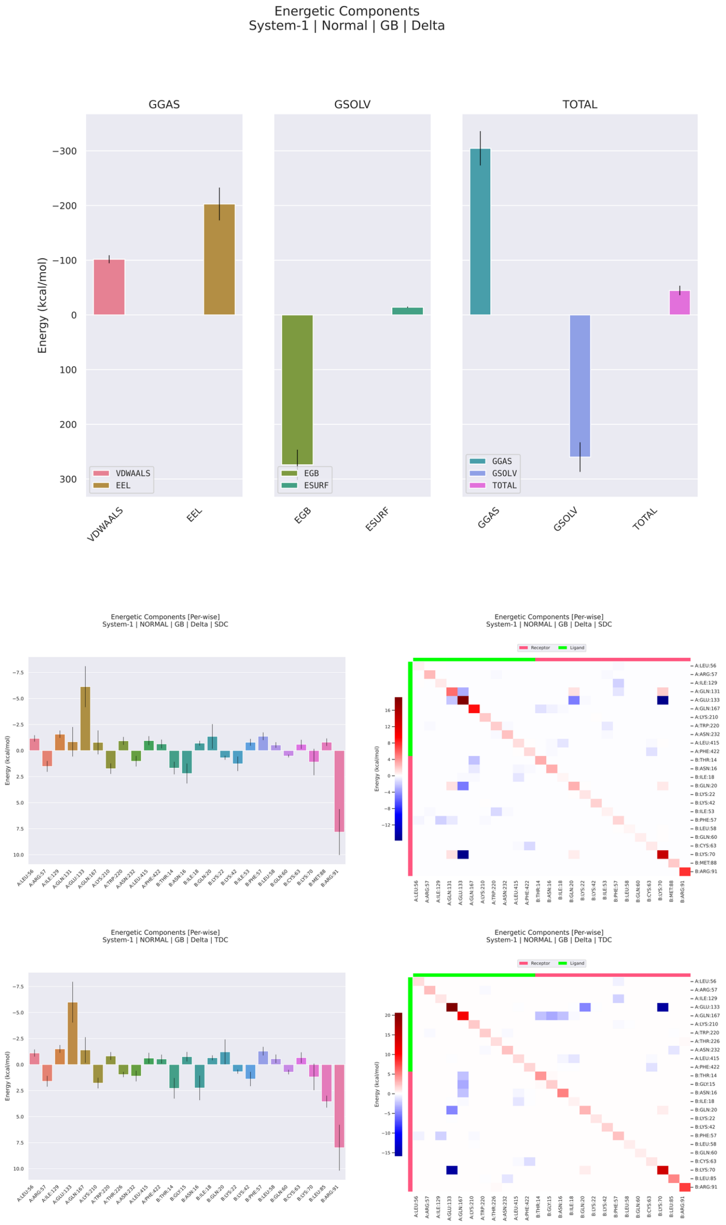


**Figure 22: MM/GBSA energetic decomposition of the dengue virus Envelope–Rab5 complex during molecular dynamics simulation.** The figure summarizes the global energetic components, per-residue decomposition, and pairwise residue decomposition of the binding free energy estimated by MM/GBSA for the dengue Envelope–Rab5 complex. The upper panels show that the gas-phase contribution is favorable and dominated by the electrostatic term, with the van der Waals component providing an additional favorable contribution of lower magnitude. Compared with the more strongly electrostatic profiles observed for some other Envelope–host complexes, the Envelope–Rab5 interaction displays a more moderate gas-phase stabilization, suggesting that the interface is supported by electrostatic complementarity but does not rely on an overwhelmingly large charge-driven contribution. The solvation component is unfavorable and dominated by the polar generalized Born term, indicating that complex formation imposes a substantial desolvation penalty on polar and charged groups at the interface. The nonpolar surface-area contribution remains comparatively small. Despite this compensation between favorable gas-phase interactions and unfavorable polar solvation, the total binding free energy remains favorable, supporting the energetic stability of the dengue Envelope–Rab5 complex after solvent effects are incorporated. The middle and lower panels show per-residue and pairwise decompositions using SDC and TDC schemes. These analyses reveal a heterogeneous energetic interface in which most residues contribute weakly or moderately, whereas a limited subset of residues concentrates the strongest stabilizing or destabilizing effects. Negative values indicate favorable contributions to binding, whereas positive values indicate unfavorable or destabilizing energetic terms. In both decomposition schemes, the energetic profile is marked by a small number of pronounced residue-level peaks rather than by a uniformly distributed contribution across the entire interface. This indicates that the Envelope–Rab5 complex is organized around discrete energetic hotspots embedded within a broader contact surface. The presence of both favorable and unfavorable residue contributions suggests that the interface is partially frustrated, with local stabilizing contacts counterbalanced by residues that impose energetic costs, possibly reflecting imperfect packing, desolvation penalties, [illegible]

##### 4.11.3 Rab5 as a Direct Structural Participant in Endosomal Viral Routing

The experimental evidence for Rab5 dependence in dengue entry is among the most direct in the DENV–host interaction literature: dominant-negative Rab5(S34N) expression blocks intracellular DENV accumulation, whereas constitutively active Rab5(Q79L) accelerates viral delivery to enlarged early endosomes from which membrane fusion is enhanced [15,111]. These observations have conventionally been interpreted as evidence that Rab5 regulates endosomal membrane identity and fusion competence rather than directly binding the viral envelope — a model in which Rab5 is a necessary contextual regulator rather than a stoichiometric viral binding partner. The computational evidence presented here introduces an alternative or complementary possibility: that the DENV envelope protein engages Rab5 directly, and that this engagement constitutes a structural mechanism by which the virus actively recruits or retains early endosomal membrane identity on the incoming endocytic vesicle.

This model would be consistent with the known behavior of several other endocytosed pathogens whose surface proteins directly engage Rab GTPases to modulate intracellular trafficking — a strategy termed "Rab hijacking" that allows viral particles to redirect endosomal maturation toward a trafficking route permissive for replication [197]. In the Rab5 case, direct envelope–Rab5 contact could serve to stabilize the Rab5-active (GTP-bound) state on the endosomal membrane by competing with GAP-mediated GTP hydrolysis, thereby prolonging the early endosomal window available for membrane fusion at low endosomal pH. The two-registry dynamic behavior observed in the simulation — where the complex progressively consolidates contacts over 200 ns — is mechanistically appropriate for this stabilization hypothesis: a durable, contact-rich interface that tightens over time would be expected to interfere with the GAP-catalyzed GTPase cycle more effectively than a transient encounter complex.

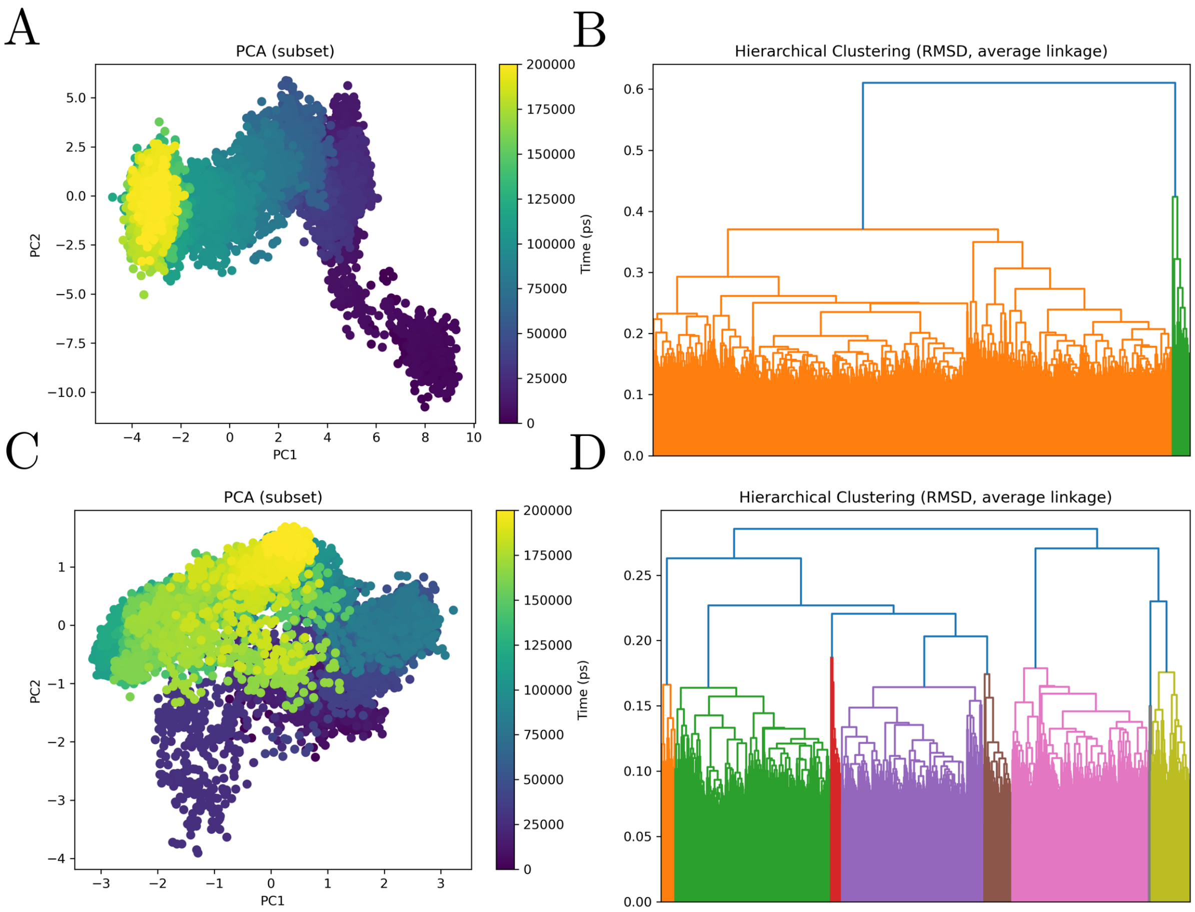


**Figure 23: Principal component analysis and RMSD-based hierarchical clustering of the dengue Envelope–Rab5 molecular dynamics trajectory.** The figure summarizes the conformational landscape sampled during the molecular dynamics simulation of the dengue Envelope–Rab5 complex, with panels A and B corresponding to the receptor subset, dengue Envelope, and panels C and D corresponding to the ligand subset, Rab5. Panels A and C show principal component analysis projections of trajectory frames onto the first two principal components, with points colored according to simulation time. In panel A, the dengue Envelope subset displays a pronounced time-ordered conformational redistribution, with early frames occupying a distinct region at positive PC1 values and progressively shifting through an intermediate manifold toward a late, more compact basin centered at negative PC1 values. This pattern indicates that the receptor undergoes substantial structural drift during the simulation rather than fluctuating around a single equilibrium state. The continuity between the early right-shifted conformers, the broad intermediate cloud, and the final left-shifted basin is consistent with progressive conformational adaptation of the Envelope within the Rab5-bound complex. The late-stage accumulation of frames in a more compact region suggests partial stabilization after an initial phase of broader conformational exploration. In panel C, the Rab5 subset exhibits a more compact but still heterogeneous conformational landscape, characterized by overlapping substates that are redistributed over time across the PCA plane. Early frames are concentrated predominantly in lower-central regions of the projection, whereas intermediate and late frames progressively populate upper and left-shifted areas, generating a temporally structured but not fully discrete transition pattern. This behavior suggests that Rab5 undergoes its own accommodation process within the complex, although with a more distributed ensemble of substates than that observed for the receptor. The partial overlap between temporal regions indicates that ligand rearrangement is progressive and dynamic, with multiple related conformational states remaining accessible throughout the trajectory. Panels B and D show RMSD-based hierarchical clustering using average linkage, revealing multiple structural families within both receptor- and ligand-associated subsets. The receptor dendrogram in panel B is dominated by one major conformational family accompanied by a smaller, structurally distinct branch, supporting the interpretation that most Envelope conformers belong to a principal basin while a more limited subpopulation samples a secondary structural state. In contrast, the Rab5 dendrogram in panel D is more finely partitioned into several clusters of comparable scale, indicating that the ligand explores a richer set of structurally distinguishable substates during the simulation. Together, the PCA and clustering analyses indicate that the dengue Envelope–Rab5 complex is dynamically heterogeneous but structurally organized, with a receptor that undergoes a marked directional conformational shift and a ligand that retains broader substate diversity. Overall, these results support a model in which Rab5 binding is associated with progressive remodeling of the Envelope-containing complex, producing an ensemble of related conformational states rather than a single rigid docking pose.

##### 4.11.4 Comparative Position of Rab5 in the Host Network: The Intracellular Trafficking Node

Among the six complexes analyzed in this study — actin, betaglycan, BiP, clusterin, GPC1, NRP1, and Rab5 — the Rab5 interface is distinguished by three properties that collectively define a mechanistically unique role as the sole intracellular trafficking node in the host protein network. First, it is the only complex in which a genuine two-basin directed conformational transition is detected at the level of PCA essential dynamics, indicating that the interaction has not reached its thermodynamic minimum within the 200 ns simulation window and that further interface refinement would be expected on longer timescales or in the membrane-proximal environment. Second, the dual-driver energetic architecture — stable vdW scaffold plus volatile electrostatic modulator — is quantitatively distinct from all other complexes: lower absolute $\Delta G_{\text{total}}$ than GPC1 ($-40$ vs. $-90$ kcal/mol) but mechanistically richer in its time-resolved behavior, reflecting the contribution of the flexible switch regions to a dynamically tunable interface. Third, the biological function of Rab5 — controlling early endosome identity and fusion timing — positions this interaction at the step in the viral lifecycle that is most directly upstream of the irreversible membrane fusion event, making it the interaction with the highest potential leverage for antiviral intervention: a compound that blocks the Rab5–envelope interface would be predicted to arrest the virus in a trafficking-incompetent early endosomal state from which it cannot progress to pH-triggered fusion, regardless of whether downstream entry factors (including the SNARE machinery, NPC1, or cathepsins) are available.

##### 4.11.5 Ternary Landscape Reveals the Rab5–Env State as Systematically Incompatible with Native Rab5 Regulatory Architecture

The conditional $\Delta\Delta G$ analysis of the Rab5-centred system reveals a qualitatively different pattern from all other receptor-centred panels in this study. Whereas Betaglycan, BiP, GPC1, and NRP1 displayed heterogeneous landscapes encompassing both cooperative and antagonistic third-partner effects, the five Rab5 regulatory partners produce exclusively antagonistic signatures, with no partner generating net AB stabilisation. This uniformity constitutes a structurally informative biological signal: the Rab5–Env state, while thermodynamically favourable as a pairwise complex ($\Delta G \approx -15.9$ kcal $\cdot$ mol$^{-1}$), is systematically displaced by the canonical regulatory machinery that governs Rab5 membrane association, GTPase cycling, and effector recruitment [109,110,116].

The mechanistic interpretation is that the Rab5 surface regions engaged by the dengue envelope — including switch I/II elements and the hypervariable C-terminal domain — overlap substantially with the interaction interfaces used by Rab5's native regulatory partners. Because Rab5 regulators bind the same GTPase surface through which the envelope contacts the protein, their introduction into a ternary assembly inevitably competes with or deforms the Env-compatible Rab5 geometry. This structural competition is not accidental; it is the mechanistic consequence of Rab5 being a regulatory hub whose surface is evolutionarily optimised for engagement with a defined set of effectors and regulators rather than for accommodating non-native viral ligands within a stable ternary complex [110,116].

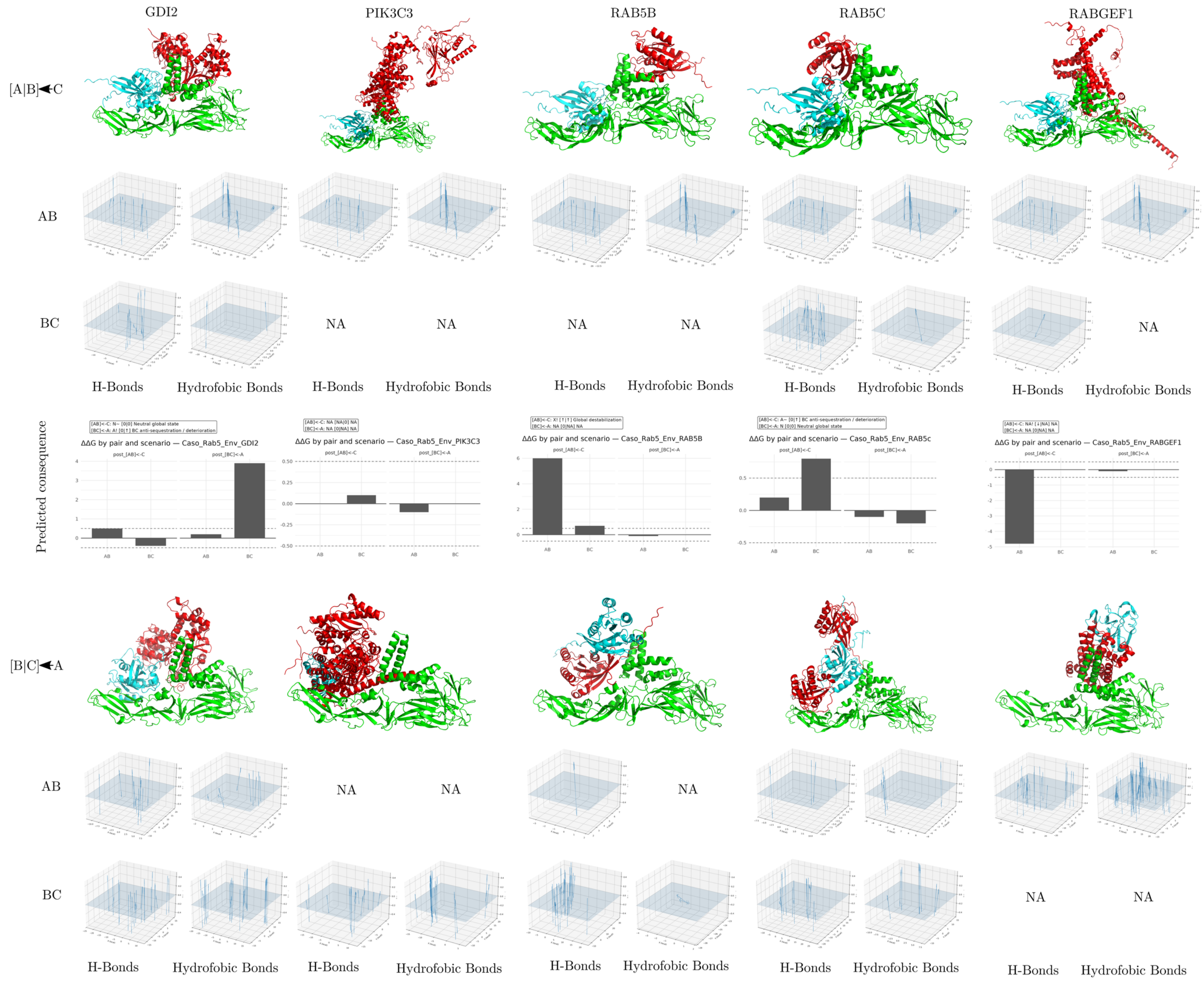
GDI2
PIK3C3
RAB5B
RAB5C
RABGEF1
[A|B]◄C
AB
BC
NA
H-Bonds
Hydrofobic Bonds
Predicted consequence
ΔΔG by pair and scenario — Caso_Rab5_Env_GDI2
ΔΔG by pair and scenario — Caso_Rab5_Env_PIK3C3
ΔΔG by pair and scenario — Caso_Rab5_Env_RAB5B
ΔΔG by pair and scenario — Caso_Rab5_Env_RAB5c
ΔΔG by pair and scenario — Caso_Rab5_Env_RABGEF1
post_[AB]<-C
post_[BC]<-A
[B|C]◄A

**Figure 24. Structural, interaction-field, and predicted energetic consequences of dengue Envelope–Rab5-centered ternary complexes with Rab5-associated proteins.**

The figure summarizes comparative ternary interaction models involving dengue virus Envelope, Rab5, and five Rab5-associated partners: GDI2, PIK3C3, RAB5B, RAB5C, and RABGEF1. In this representation, A corresponds to dengue Envelope, B corresponds to Rab5, and C corresponds to the additional Rab5-associated protein. Two alternative assembly scenarios are compared. In the first scenario, the Envelope–Rab5 pair is treated as the preformed binary complex and the third protein is introduced afterward, represented as $[A]B \leftarrow C$. In the reciprocal scenario, the Rab5–associated-protein pair is treated as preformed and dengue Envelope is introduced as the third component, represented as $[B]C \leftarrow A$.

Structural panels show the relative spatial organization of the three proteins in each modeled ternary assembly, revealing pronounced partner-dependent differences in geometry and packing. Interaction-field plots report the spatial distribution of hydrogen-bond and hydrophobic contact features for the AB and BC interfaces, showing that polar and nonpolar interaction densities are redistributed according to both partner identity and assembly order. Predicted energetic consequence plots report $\Delta\Delta G$ values for each pair and assembly scenario, with negative values indicating stabilizing effects and positive values indicating destabilizing or energetically unfavorable effects relative to the corresponding reference state. Missing descriptors or unavailable configurations are indicated as NA.

Overall, the figure indicates that the dengue Envelope–Rab5 interaction is strongly modulated by the molecular neighborhood of Rab5. Some neighbors, such as RAB5B, preferentially perturb the AB interface when added to a preformed Envelope–Rab5 complex, whereas others, such as GDI2, more strongly penalize the BC interface when dengue Envelope is introduced into preformed Rab5-associated assemblies. PIK3C3 exerts comparatively modest energetic effects, whereas RAB5C and RABGEF1 show more asymmetric and assembly-order-dependent behavior.

Together, these results support a path-dependent model of ternary complex formation in which the structural and energetic outcome depends not only on the identity of the interacting proteins, but also on the order in which the viral and host components assemble, consistent with Rab5 functioning as a network-embedded host interface within the endosomal trafficking system.

##### 4.11.6 GDI2 Defines the Cytosolic Extraction Boundary Condition

The GDI2 result is mechanistically the most comprehensive of the Rab5 panel because it generates a global rather than selective deterioration of the interaction landscape: both AB ($\Delta\Delta G \approx +4.5$ kcal · mol$^{-1}$) and BC (+3.9 kcal · mol$^{-1}$) are destabilised under AB conditioning, and reverse conditioning amplifies rather than attenuates the AB penalty (+4.8 kcal · mol$^{-1}$). GDP dissociation inhibitors bind the GDP-loaded, prenylated Rab GTPase and extract it from membranes into a soluble cytosolic complex, masking the prenyl group and blocking effector access by sequestering the switch regions [109,110]. The global energetic deterioration observed here is the conditional-docking signature of this extraction process: GDI2 engagement forces Rab5 into a conformation in which neither the Env-facing nor the BC-facing surfaces retain their Env-compatible geometry, consistent with a receptor that has been withdrawn from the membrane-associated interaction landscape entirely.

The reverse-conditioning amplification — where a pre-formed Rab5–GDI2 complex presents even greater resistance to Env engagement (+4.8 vs. +4.5 kcal · mol$^{-1}$) — is also biologically coherent: GDI2-extracted Rab5 is in a compact, soluble, GDP-bound state incompatible with membrane-associated interaction partners, and Env encountering this pre-formed GDI2 complex faces a more severely deformed Rab5 surface than one established within an already partially disrupted ternary assembly.

##### 4.11.7 Paralogous Rab5 GTPases Reveal Surface Competition Through Alternative Rab–Rab Assemblies

The RAB5B and RAB5C results illuminate the specificity of the Rab5–Env interaction surface through an internal comparison. Both paralogs share high sequence similarity with RAB5A and participate in overlapping early endosomal trafficking functions [109,110]; yet their conditional signatures differ substantially. RAB5C produces a symmetric, BC-sparing profile (+4.2 and +4.5 kcal · mol$^{-1}$ across branches, BC essentially unchanged), indicating selective AB displacement without broader interaction-landscape reorganisation. RAB5B, by contrast, generates a strongly asymmetric profile in which reverse conditioning produces a catastrophic AB collapse (+10.3 kcal · mol$^{-1}$) — the largest single-configuration destabilisation in the Rab5 panel — while BC remains unperturbed under both conditions.

This paralog-level asymmetry suggests that RAB5B and the RAB5A target engage similar but not identical surface determinants when assembling into putative Rab–Rab contacts. The extreme reverse-conditioned RAB5B destabilisation implies that a pre-formed Rab5A–RAB5B assembly occludes or grossly deforms the Env-binding surface of Rab5A more thoroughly than RAB5C does, potentially reflecting differences in inter-Rab contact geometry encoded by divergent hypervariable C-terminal sequences. This structural specificity within the paralog class is consistent with the known functional non-equivalence of RAB5 isoforms in specific trafficking contexts despite their overall pathway redundancy [109], and motivates isoform-resolved experimental interrogation of Rab5-dependent dengue trafficking.

###### 4.11.8 PIK3C3 Reflects Endosomal Identity Complex Incompatibility

PIK3C3 (Vps34) is the sole class III phosphatidylinositol 3-kinase in mammals and functions as the catalytic core of two distinct Beclin-1-containing complexes that generate phosphatidylinositol 3-phosphate (PI3P) on early endosomal and autophagosomal membranes [110,118]. In the context of Rab5-dependent early endosome maturation, PIK3C3 is recruited to Rab5-positive membranes via the effector complex APPL1–Rabaptin5, and its PI3P product recruits a cascade of FYVE- and PX-domain-containing effectors that collectively establish early endosome identity [110,118].

The selective AB destabilisation produced by PIK3C3 (+4.2 $\text{kcal} \cdot \text{mol}^{-1}$) without BC perturbation indicates that PIK3C3 competes for the Env-facing surface of Rab5 without disrupting the Rab5–PIK3C3 binary contact itself. Mechanistically, this is consistent with PIK3C3 engaging the Rab5 effector-binding surface — overlapping with the region through which Env contacts Rab5 — within a complex organised around membrane-associated PI3P signalling rather than free diffusion. Biologically, this result suggests that the assembly of canonical Rab5–Vps34 endosomal identity complexes would be expected to reduce the probability of sustained Rab5–Env interaction, reinforcing the view that Rab5 contributes to viral trafficking through endosomal membrane dynamics rather than through direct stable viral-protein engagement.

###### 4.11.9 RABGEF1 Reveals the Activated-State Incompatibility of the Env-Associated Rab5 Configuration

RABGEF1 (also known as Rabex-5) is the canonical guanine nucleotide exchange factor for Rab5, catalysing GDP-to-GTP exchange and converting Rab5 into its effector-recruiting active form [109,116]. GEF-mediated activation induces structural rearrangements in the Rab5 switch I and switch II regions that alter the electrostatic and geometric properties of the effector-binding surface. RABGEF1 produced a distinctive two-axis perturbation: AB was destabilised by +4.2 $\text{kcal} \cdot \text{mol}^{-1}$ under AB conditioning while BC collapsed by +6.4 $\text{kcal} \cdot \text{mol}^{-1}$ — the largest single BC perturbation in the Rab5 panel — and the reverse-conditioned branch produced only a mild AB stabilisation ($-0.5$ $\text{kcal} \cdot \text{mol}^{-1}$).

The BC destabilisation is mechanistically significant: it indicates that within the AB-conditioned ternary assembly, RABGEF1 does not achieve a stable Rab5-contact even while disrupting the Env interaction, suggesting that the Env-bound Rab5 configuration — which the binary MD and MM/GBSA analyses establish as a non-equilibrium state still undergoing conformational refinement — presents a switch-region geometry that is incompatible with productive GEF docking. In other words, the Env-associated Rab5 state may resist GEF-mediated nucleotide exchange in addition to being destabilised by GEF presence: a mutually incompatible pair in which neither partner can adopt its productive interaction geometry while bound to the other. The mild reverse-conditioned AB stabilisation ($-0.5$ kcal · mol$^{-1}$) is the sole cooperative-direction perturbation in the entire Rab5 panel, arising specifically when Env approaches a pre-formed Rab5–RABGEF1 state; its small magnitude confirms it is a minor and transient effect rather than a genuine cooperative basin.

###### 4.11.10 The Uniform Antagonism of Rab5 Regulators Supports an Endocytic Trafficking Rather Than Direct Entry-Complex Model

The uniformity of antagonistic signatures across all five Rab5 regulatory partners supports a coherent biological model that clarifies the mechanistic role of Rab5 in dengue infection. The Rab5–Env pairwise interaction is structurally plausible and energetically favourable in isolation, but it is systematically incompatible with the native Rab5 regulatory network. This incompatibility implies that any Rab5–Env co-complex formed during viral infection would be transient and context-dependent: it would exist only in the narrow window when Rab5 is present at an endosomal membrane without concurrent engagement by GDI, GEF, paralogous Rab GTPases, or PI3K-containing identity complexes.

This transient, regulatory-context-sensitive model is fully consistent with the established biology of dengue endosomal trafficking. Rab5 is required for productive viral entry not because it serves as a stable docking partner for the viral envelope, but because it orchestrates the endosomal maturation and acidification sequence that culminates in pH-triggered membrane fusion [15,16,109]. A transient Rab5–Env contact at the early endosomal membrane — occurring between GEF-mediated activation and GDI-mediated extraction, and displaced by the assembly of canonical effector complexes — would be mechanistically sufficient to influence the spatial and temporal positioning of the viral particle within the endosomal maturation pathway without requiring a prolonged, stable binary interaction.

From a comparative perspective, the uniformly antagonistic Rab5 ternary landscape contrasts sharply with the mixed cooperative-antagonistic landscapes of all receptor-centred host proteins in this study. This contrast reinforces the classification of Rab5 as a mechanistically distinct intracellular trafficking node rather than an extracellular attachment or co-receptor factor. The practically relevant next steps are experimental validation of the Rab5–Env contact specificity using dominant-negative and constitutively active Rab5 mutants, time-resolved endosomal imaging during dengue infection, and biochemical pulldown assays designed to detect the transient Rab5–Env interaction under conditions that suppress competing regulatory engagement [41,47,49].

###### 4.11.11 Rab5 Regulatory Landscape as a Biological Timer: Endosomal Window Kinetics and Allosteric GTPase Displacement

The uniformly antagonistic regulatory landscape of Rab5 has a kinetic implication that transcends the question of binary contact stability: it defines a *biological timer* for dengue endosomal trafficking. The key observation is that all five regulatory partners — GDI2 (cytosolic extraction), RABGEF1 (GEF-mediated activation), RAB5B/RAB5C (paralogous Rab competition), and PIK3C3 (PI3P identity complex) — cover the full temporal cycle of Rab5 membrane dynamics. GDI2 represents the GDP-bound cytosolic state preceding membrane insertion; RABGEF1 represents the activation transition from GDP to GTP; RAB5B/RAB5C represent competing Rab isoforms contesting the same endosomal membrane territory; and PIK3C3 represents the effector-engaged state downstream of GTP loading. If the Rab5–Env interaction is structurally incompatible with all stages of the canonical Rab5 cycle, then the only temporal window during which viral engagement of Rab5 is structurally permitted is the brief interval *between* GEF-mediated activation (which displaces the GDP state that precedes RABGEF1 engagement) and PI3K complex assembly (which occludes the activated state). This interval corresponds precisely to the transition period during which the newly formed early endosome acquires its Rab5-GTP coat but has not yet assembled the full Vps34/Beclin-1 identity complex — a window of tens of seconds in the cell biology literature [1,109,110].

The allosteric dimension of this timing mechanism is equally important. The ternary analyses reveal that GDI2 and RABGEF1 do not simply occlude the Env-binding surface sterically — they alter the global energetic landscape of Rab5 in ways that simultaneously disrupt multiple interaction surfaces (GDI2 destabilises both AB and BC; RABGEF1 collapses BC while only moderately weakening AB). This pattern is the signature of allosteric communication within the GTPase core: when GDI2 sequesters the prenyl group and closes the GTPase domain into its cytosolic-extraction conformation, the conformational changes propagate through the switch I/II regions to alter the surface geometry remote from the GDI2 contact — the same surface that Env engages. Conversely, RABGEF1-induced GTP loading induces switch rearrangements that close the nucleotide-binding pocket and remodel the effector-facing surface in a direction incompatible with the Env contact geometry established in the GDP-Rab5 binary complex [109,116].

This allosteric timer model predicts that dengue virus does not require sustained stable engagement with Rab5 protein to exploit Rab5-dependent endosomal routing. Instead, transient contact during the GDP-to-GTP transition window — sufficient to position the virion appropriately on the nascent early endosomal membrane — would be mechanistically adequate to ensure routing to the acidic endosomal lumen where pH-triggered fusion occurs. This parsimonious mechanism is consistent with single-virion tracking data showing that dengue particles spend only minutes in Rab5-positive endosomes before transitioning to Rab7-positive late endosomes for membrane fusion [17,18], and it generates a specific prediction: perturbation of the GDP→GTP transition timing — for example by dominant-negative RABGEF1 (Rabex-5) that prolongs the GDP-bound state — would extend the Rab5–Env interaction window, slowing but not abolishing viral endosomal routing, whereas constitutively active Rab5-Q79L (GTP-locked) would eliminate the permissive interaction window entirely and arrest viral particles in a Rab5-positive compartment incompatible with productive fusion.

#### 4.12 Envelope–SUMO9-UBC1: Post-Translational Machinery Engagement Through Cooperative Compaction and a Dual Salt-Bridge Hotspot

##### 4.12.1 Cooperative Induced-Fit as a Mechanistically Distinct Binding Strategy

The conformational analysis of the SUMO complex reveals a binding mechanism that is categorically different from all other host protein interactions characterized in this study. In the GPC1 complex, the envelope undergoes a directed single transition while GPC1 samples a broad multi-state ensemble; in the NRP1 complex, both partners exhibit bilateral conformational restriction imposed by the rigid b1-pocket geometry; in the Rab5 complex, the envelope pivots between two registries while Rab5 remains geometrically stable. In the SUMO complex, both partners undergo coordinated convergence toward distinct late-time basins — the envelope through a large-amplitude compaction and C-terminal reorganization, SUMO9-UBC1 through substate selection among local conformers — generating a cooperative induced-fit mechanism in which the final bound geometry is co-constructed by both partners rather than dictated by one. This cooperative mechanism is the energetic rationale for the three-phase interface development observed in the contact analysis: the initial phase reflects the unconstrained early docking geometry, the intermediate phase reflects the envelope's conformational adaptation, and the late phase reflects the jointly optimized packing core whose density and stability exceed the early-phase interface by approximately 12% in contact number and by substantially more in energetic coherence [93,128].

The extreme structural rigidity of SUMO9-UBC1 during this cooperative process is functionally significant. A regulatory E2 enzyme that maintains active-site geometry while accommodating diverse substrate interfaces must balance binding adaptability with catalytic precision; the substate selection mechanism observed here — in which the SUMO9-UBC1 loops and N-terminal extension filter among compatible conformers without reorganizing the catalytic UBC fold — is precisely the structural strategy that would allow the E2 to engage a non-canonical binding partner (the viral envelope) while retaining the ability to transfer SUMO to a substrate lysine [122,123]. This interpretation suggests that the envelope may function as a competitive inhibitor or allosteric modulator of Ubc9 activity: by occupying the substrate-recruiting surface of Ubc9 through the GLU85/ARG94/PHE96 hotspot and the peripheral loop contacts, the envelope could reduce the effective concentration of SUMO-charged Ubc9 available for antiviral substrate modification, thereby modulating the host SUMOylation response to viral infection.

##### 4.12.2 Energetic Logic of the Packing-Dominant Interface

The predominance of VDWAALS over EEL in the MM/GBSA decomposition — unique among all complexes in this study — is mechanistically interpretable in terms of the structural features of the interface. The envelope C-terminal stem-proximal region that drives the late-time contact densification presents primarily aliphatic and aromatic residues whose packing against the SUMO $\beta$-grasp surface generates shape-complementary van der Waals contacts without requiring the precise charge-pair geometry that governs the GPC1 or NRP1 interfaces. This packing-dominant architecture has a distinctive thermodynamic consequence: VDWAALS is substantially less sensitive to conformational fluctuations than EEL, and therefore provides a more stable energetic baseline across the ensemble of interface microstates. The lower temporal variance in $\Delta G_{\text{total}}$ observed for the SUMO complex compared to the GPC1 complex is the quantitative expression of this property: a packing-dominated interface tolerates more geometric variation without large energetic consequences, whereas an electrostatically dominated interface is highly sensitive to the instantaneous geometry of charge pairs [49].

The two dominant salt bridges — GLU85–LYS59 and ARG94–GLU42 — superimposed on this packing core provide the electrostatic specificity component that distinguishes genuine molecular recognition from non-specific adsorption. Their identification in the TDC heatmap as the two deepest pairwise interaction contributions, combined with their spatial proximity to the PHE96 packing contact, creates a three-residue envelope hotspot where steric and electrostatic mechanisms are synergistically co-localized: the aromatic packing of PHE96 positions the flanking charged residues at optimal distances for simultaneous salt-bridge formation with their SUMO9-UBC1 counterparts, generating an interaction geometry that is both energetically favorable and sterically constrained against dissociation [129,198]. This geometry is mechanistically coherent with the spatial organization of the catalytic loop of Ubc9, where the substrate-facing surface presents charged and hydrophobic residues in a pattern optimized for recognition of substrate SIM motifs [123].

The frustration loci — LYS30, LYS59, and SER158 on SUMO9-UBC1 — provide a complementary mechanistic insight. Their positive SDC contributions identify positions where the current binding geometry fails to fully compensate desolvation costs, suggesting that the late-time interface is not the absolute energetic minimum but rather a kinetically stable local minimum accessible within the 200 ns timescale. In the context of a transient regulatory interaction, partial energetic frustration at the interface periphery may be functionally advantageous: it maintains the reversibility required for catalytic turnover, ensuring that the envelope can both recruit and release the E2 enzyme at rates compatible with the kinetics of viral replication rather than forming an irreversibly trapped complex [43].

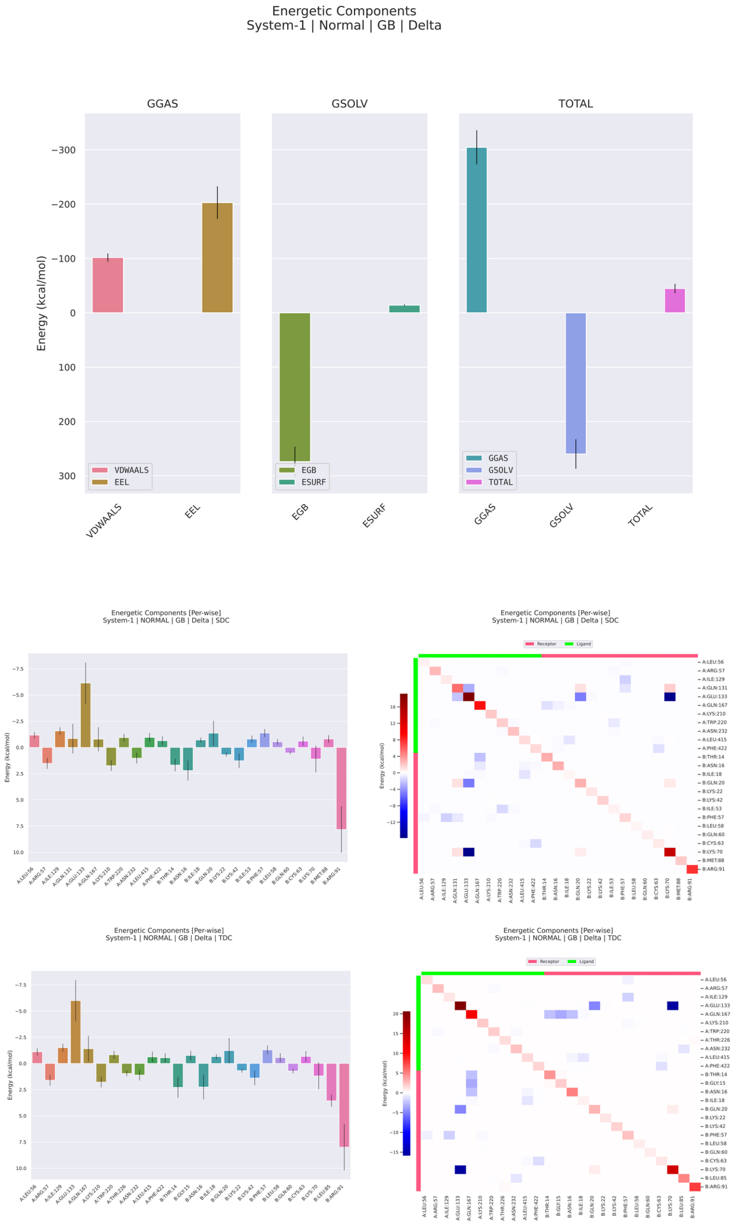


**Figure 25: MM/GBSA energetic decomposition of the dengue virus Envelope–SUMO complex during molecular dynamics simulation.** The figure summarizes the global energetic components, per-residue decomposition, and pairwise residue decomposition of the binding free energy estimated by MM/GBSA for the dengue Envelope–SUMO complex. The upper panels show that the gas-phase contribution is favorable and is dominated by the electrostatic term, with the van der Waals component providing an additional, lower-magnitude stabilizing contribution. Compared with the more strongly electrostatic Envelope–host complexes observed for larger or highly charged interfaces, the Envelope–SUMO profile displays a more moderate gas-phase stabilization, suggesting that the interaction is supported by electrostatic complementarity but does not depend on an overwhelmingly large charge-driven term. The solvation component is unfavorable and dominated by the polar generalized Born contribution, indicating that complex formation imposes a substantial desolvation penalty on polar and charged groups at the interface. The nonpolar surface-area term contributes only weakly relative to the electrostatic and polar solvation components. Despite this compensation between favorable gas-phase attraction and unfavorable polar solvation, the total binding free energy remains favorable, supporting the energetic stability of the dengue Envelope–SUMO complex after solvent effects are incorporated. The middle and lower panels show per-residue and pairwise decompositions using SDC and TDC schemes, revealing a heterogeneous energetic architecture in which most residues contribute weakly or moderately, whereas a limited subset of residues concentrates the dominant stabilizing or destabilizing effects. Negative values indicate favorable contributions to binding, whereas positive values indicate unfavorable or destabilizing energetic terms. In both decomposition schemes, the interface is marked by discrete residue-level peaks rather than by a uniform energetic contribution across the entire contact surface. This indicates that the Envelope–SUMO complex is organized around localized energetic hotspots embedded within a broader and partially compensated interaction landscape. Several residues contribute modest favorable terms, but the strongest residue-level effects are concentrated in a restricted region, suggesting that SUMO recognition by dengue Envelope may

##### 4.12.3 SUMOylation Pathway Hijacking as a Host-Proteostasis Exploitation Strategy

The biological significance of the envelope–SUMO9-UBC1 interaction must be interpreted within the dual context of the SUMO pathway as both a host antiviral defense and a potential viral exploitation target. On the antiviral side, the SUMO pathway modifies numerous interferon signaling components, including STAT1, STAT2, and IRF3/7, enhancing their transcriptional activity and thereby amplifying the innate immune response to viral infection [123,127]. On the proviral side, dengue and other flaviviruses have been shown to benefit from functional SUMOylation machinery: dominant-negative inhibition of the SUMO pathway reduces dengue replication efficiency, and the envelope protein's experimentally confirmed interaction with Ubc9 [125] suggests that the virus has evolved to either exploit or subvert this machinery rather than simply evade it. The SUMOylation pathway has additionally been reported as essential for dengue replication and transmission in mosquito vectors [126], indicating that the viral dependence on this pathway extends across host species and is not an artifact of cell culture systems.

Within this framework, the structural interaction characterized here offers a molecularly concrete mechanism for viral exploitation of the SUMO conjugation axis. The GLU85/ARG94/PHE96 hotspot on the envelope surface engages Ubc9 at or near the substrate-recognition interface, the same region responsible for SUMO transfer to host antiviral substrates. By occupying this surface, the viral envelope could achieve one or more of the following consequences: (i) direct competitive inhibition of the SUMOylation of antiviral substrates, reducing the potency of the innate immune SUMO response; (ii) sequestration of SUMO-charged Ubc9 into an unproductive envelope-bound complex, reducing the available pool for legitimate substrate modification; or (iii) modification of the envelope itself at K51 or K241 by the bound Ubc9, generating SUMOylated envelope protein whose functional consequences for viral assembly or immune evasion remain to be characterized. The cooperative induced-fit mechanism and the three-phase kinetic profile are consistent with any of these functional models, as all require a sustained and geometrically coherent interaction between the envelope and Ubc9 that evolves over a timescale comparable to viral replication.

##### 4.12.4 Envelope as an Allosteric Modulator of Ubc9 Catalytic Activity: Beyond Steric Competition

The functional models described above frame dengue envelope–Ubc9 engagement primarily as a competition for substrate access — a physical blockade that reduces the pool of Ubc9 available for antiviral SUMOylation. However, the structural dynamics of the SUMO9-UBC1 complex characterised here suggest a mechanistically richer interpretation: the envelope protein may act as an *allosteric modulator* of Ubc9 catalytic activity rather than exclusively as a steric competitor. This distinction is consequential for understanding both the biological reach of the interaction and the mechanistic basis for its antiviral potential.

Allosteric regulation of SUMO E2 activity has been demonstrated by multiple structural and biochemical studies. The Ubc9 UBC fold is not a rigid scaffold but a conformationally responsive enzyme whose catalytic geometry — the positioning of the active-site Cys93 relative to the incoming SUMO thioester and the substrate lysine — is sensitive to contacts at peripheral surfaces distant from the active site [199,200]. NMR and hydrogen-deuterium exchange studies of Ubc9 have shown that binding at the back-side hydrophobic surface — a region that partially overlaps with the GLU85/ARG94/PHE96 contact geometry identified in the present study — allosterically shifts the ensemble of Ubc9 conformations toward states with reduced active-site geometry optimality, reducing kcat for SUMO transfer without abolishing E1 binding or thioester loading [200,201]. The envelope protein's engagement of Ubc9 at peripheral loop and C-terminal positions, while leaving the catalytic core low-RMSF (as established by the RMSF analysis of the SUMO9-UBC1 complex), is structurally consistent with precisely this allosteric mode: the catalytic UBC domain core is not globally perturbed, but the peripheral conformational substates sampled by Ubc9 when envelope-bound may systematically disfavour the transition-state geometry required for efficient SUMO conjugation.

The biological significance of allosteric Ubc9 modulation is particularly acute for innate immune suppression. STAT1 and STAT2, the transcription factors mediating type I interferon signalling, are substrates for SUMO modification at K703 (STAT1) and K390 (STAT2), and SUMO conjugation at these sites has been shown to enhance STAT-dependent transcriptional responses to interferon stimulation [202,203]. Ubc9-mediated SUMOylation of IRF3 and IRF7 also augments their transcriptional activity in the antiviral response [123,127]. If dengue envelope allosterically suppresses Ubc9 catalytic efficiency toward these antiviral substrates — rather than merely competitively blocking substrate access — the consequence would be a quantitative, graded reduction in STAT and IRF SUMOylation that attenuates interferon pathway amplification without fully abolishing basal signalling. This graded, allosteric immune evasion mode would be mechanistically analogous to the partial agonism strategy employed by viral proteins targeting other host signalling enzymes, in which incomplete rather than total pathway inhibition is evolutionarily favoured because it reduces the selective pressure for host resistance mutations [148,191].

This hypothesis predicts a specific experimental signature: in cells infected with dengue, the catalytic activity of Ubc9 extracted in complex with envelope protein should display reduced specific activity toward STAT1/STAT2 SUMO-acceptor peptide substrates relative to Ubc9 from uninfected cells or from infected cells lacking the envelope–Ubc9 interaction (achievable through K51A/K241A envelope point mutants [125]). Detection of this activity reduction in the absence of reduced Ubc9 protein levels would constitute evidence for allosteric rather than purely competitive modulation, distinguishing the two mechanistic models and providing functional validation of the structural observations presented here.

##### 4.12.5 The Envelope as a Non-Competitive Ubc9 Inhibitor: Catalytic Cleft Geometry and Enslaved PTM Machinery

The allosteric model introduced above can be formalised within the framework of enzyme inhibition kinetics. Classical competitive inhibition of Ubc9 would require the envelope protein to bind the substrate-recognition surface in direct competition with SUMO-acceptor substrates, predicting a Michaelis-Menten kinetic signature of elevated apparent $K_M$ without change in $V_{\text{max}}$. The structural evidence presented here is inconsistent with this competitive model in its pure form: the primary Env-contact region on Ubc9 is the peripheral backside surface and C-terminal lobe, not the Cys93 catalytic groove or the SIM-binding groove that governs substrate lysine positioning. This topological separation of the Env-contact site from the catalytic machinery defines the envelope as a *non-competitive* or *mixed inhibitor* in the Ubc9 system, whose kinetic signature would be a reduction in $V_{\text{max}}$ (reflecting allosteric reduction in catalytic rate constant $k_{\text{cat}}$) with variable effects on apparent $K_M$ depending on whether the peripheral conformational shift also alters substrate affinity [204].

Non-competitive enzyme inhibition by viral proteins is a well-precedented immune evasion strategy. Viral proteins targeting host kinases (e.g., NS5 of flaviviruses targeting JAK1/TYK2) frequently exploit allosteric surfaces remote from the ATP-binding pocket to suppress kinase activity without competing directly with ATP or substrate peptide, enabling inhibition under conditions of high ATP concentration that would overcome competitive inhibitors [205]. If dengue envelope adopts an analogous non-competitive mode against Ubc9, it would achieve proportional suppression of SUMO conjugation at high substrate concentrations — including the elevated STAT1/IRF3 levels induced during the interferon response — where competitive inhibition would be least effective.

A cautious interpretation of this model is that the dengue envelope may do more than sterically block the SUMOylation machinery: it could redirect or attenuate aspects of Ubc9-dependent substrate processing. Ubc9 engaged by the envelope retains its E1 interaction surface (the N-terminal helix that contacts SAE1/UBA2), allowing continued thioester loading with SUMO, but redirects or attenuates the transfer step toward antiviral substrates while potentially preferentially conjugating substrates that benefit viral replication. SUMO modification has been reported to stabilise certain viral replication complex components and to modulate the nuclear import of flavivirus NS proteins [126,206]; if Env-bound Ubc9 is trapped in a conformation that selectively channels SUMO toward viral-beneficial substrates while disfavouring antiviral ones, the interaction represents a candidate form of immune modulation rather than a demonstrated post-translational modification pathway redirection — a mechanistic mode for which the three-phase kinetic densification profile of the SUMO9-UBC1 MD complex provides the structural precedent, as it demonstrates that the envelope progressively organises Ubc9 into a well-defined, dense contact geometry that persists on a timescale consistent with enzymatic turnover cycles.

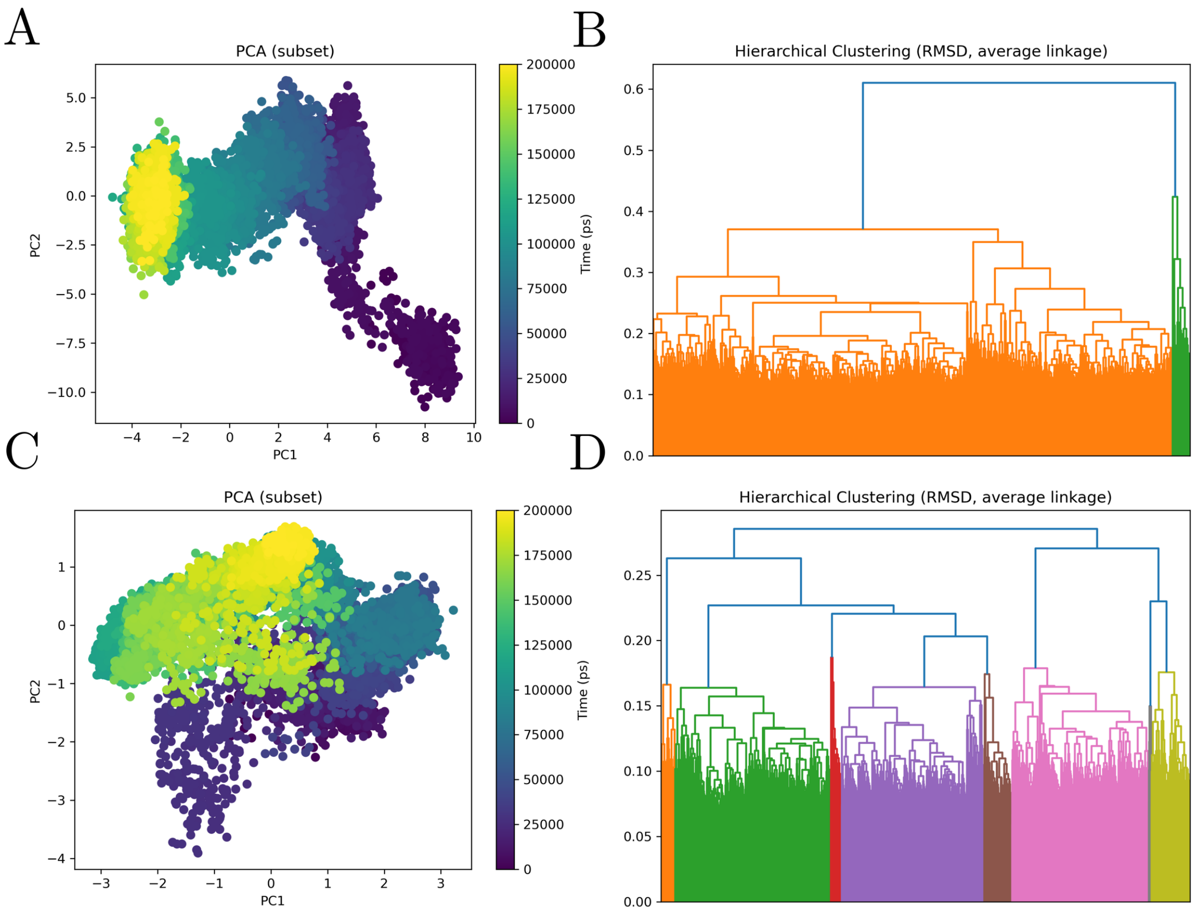


**Figure 26: Principal component analysis and RMSD-based hierarchical clustering of the dengue Envelope–SUMO molecular dynamics trajectory.** The figure summarizes the conformational landscape sampled during the molecular dynamics simulation of the dengue Envelope–SUMO complex, with panels A and B corresponding to the receptor subset, dengue Envelope, and panels C and D corresponding to the ligand subset, SUMO. Panels A and C show principal component analysis projections of trajectory frames onto the first two principal components, with points colored according to simulation time. In panel A, the dengue Envelope subset displays a marked directional redistribution across conformational space, with early frames occupying a distinct region at positive PC1 values and low PC2 values, followed by progressive migration through intermediate conformational regions toward a late-stage basin at negative PC1 values. This temporal organization indicates that the receptor does not remain confined to a single equilibrium geometry, but instead undergoes substantial conformational drift during the simulation. The relatively continuous connection between early, intermediate, and late regions suggests a gradual remodeling process, compatible with progressive adaptation of the Envelope within the SUMO-bound state. The late frames accumulate in a more compact region than the early ensemble, indicating partial stabilization after an initial phase of broader conformational exploration. In panel C, the SUMO subset displays a more compact but clearly structured conformational landscape. Early frames are concentrated mainly in lower-central regions of the PCA projection, whereas intermediate and late frames progressively redistribute toward upper and left-shifted regions. This pattern indicates that SUMO also undergoes time-dependent accommodation within the complex, although its conformational rearrangement appears more locally constrained than the larger receptor-associated redistribution. The partial overlap among temporal regions suggests that SUMO samples a set of related substates rather than undergoing a single abrupt transition. Nevertheless, the ordered color gradient across the PCA plane supports the interpretation that the ligand participates actively in the conformational evolution of the complex, instead of acting as a rigid passive binding partner. Panels B and D show RMSD-based hierarchical clustering using average linkage, revealing multiple structural families within both receptor- and ligand-associated subsets. The receptor dendrogram in panel B is dominated by one large conformational family accompanied by a smaller, structurally distinct branch, supporting the existence of a principal Envelope-associated basin with subsidiary conformational substates. This agrees with the PCA projection, in which the receptor follows a broad directional transition but retains a dominant late-trajectory region. The SUMO dendrogram in panel D shows a more finely partitioned clustering pattern, with several well-resolved conformational groups distributed across the trajectory. This indicates that, despite its smaller and more compact structural profile, SUMO samples multiple distinguishable substates during the simulation. Together, the PCA and clustering analyses indicate that the dengue Envelope–SUMO complex is dynamically heterogeneous but structurally organized, with a receptor that undergoes pronounced directional remodeling and a ligand that remains comparatively compact while exploring several related conformational substates. Overall, these

###### 4.12.6 Comparative Energetic Position of SUMO9-UBC1 in the Host Network

Among all complexes analyzed in this study, the SUMO9-UBC1 interface occupies a distinctive position in the energetic landscape defined by the ratio of VDWAALS to EEL contributions. Clusterin represents the most hydrophobic extreme, driven primarily by disordered-tail VDWAALS contacts with minimal electrostatic contribution; GPC1 represents the most electrostatic extreme, with EEL saturation generating the largest absolute $\Delta G_{\text{total}}$ through charge–charge complementarity; and NRP1 represents the geometrically constrained intermediate, where pocket-insertion geometry balances both terms with low temporal variance. SUMO9-UBC1 occupies the VDWAALS-dominant region of this spectrum — more hydrophobic than NRP1 or BiP, less so than clusterin — but is distinguished from clusterin by the presence of two highly specific, structurally resolved salt bridges that provide recognition selectivity absent from the distributed hydrophobic contacts of the chaperone interface. This combination — packing dominance with electrostatic specificity — is the energetic fingerprint of a bona fide protein–enzyme interaction that requires both structural fit and chemical complementarity for productive complex formation, and distinguishes the SUMO complex as mechanistically the most enzyme-like interaction in the host protein network, with direct implications for antiviral strategies targeting the viral modulation of host post-translational modification pathways.

###### 4.12.7 Ternary Landscape Establishes the SUMO1–Env State as a Shallow Basin Systematically Outcompeted by Canonical Pathway Assemblies

The conditional $\Delta\Delta G$ analysis of the SUMO1-centred system reveals a pattern qualitatively distinct from all other host protein panels in this study. Rather than a heterogeneous mixture of cooperative and antagonistic perturbations, all seven tested partners uniformly and strongly stabilise the AB axis — a pattern that is directly interpretable in terms of the intrinsically weak reference SUMO1–Env state ($\Delta G \approx -3.4\ \text{kcal} \cdot \text{mol}^{-1}$) being replaced by the energetically far superior canonical SUMO assembly geometries. The SUMOylation system is not a simple receptor–ligand interaction but a tightly choreographed enzymatic cycle involving the E1 heterodimer (SAE1–UBA2), the sole E2 enzyme UBC9, multiple SUMO paralogs, and structurally defined substrate complexes [122,130,131,207,208]. In such a system, one would expect bona fide SUMO-pathway configurations to dominate the energy landscape over a non-canonical, low-affinity SUMO–Env contact — and the conditional landscapes confirm this expectation quantitatively and systematically.

The biological interpretation is therefore clear and biologically conservative: SUMO1 does not behave as a stable receptor-like platform for the dengue envelope. Instead, the SUMO1–Env state is a shallow, transient configuration that exists at the periphery of the SUMO signalling network, readily displaced whenever canonical SUMO pathway components organise around the SUMO1 surface. This conclusion parallels the

Rab5 result — where all regulatory partners also antagonised the Rab5–Env state — but arises through a mechanistically distinct route: in the Rab5 case, competitive surface overlap with regulatory machinery; in the SUMO1 case, energetic dominance of high-affinity canonical SUMO assemblies over an intrinsically weak reference state.

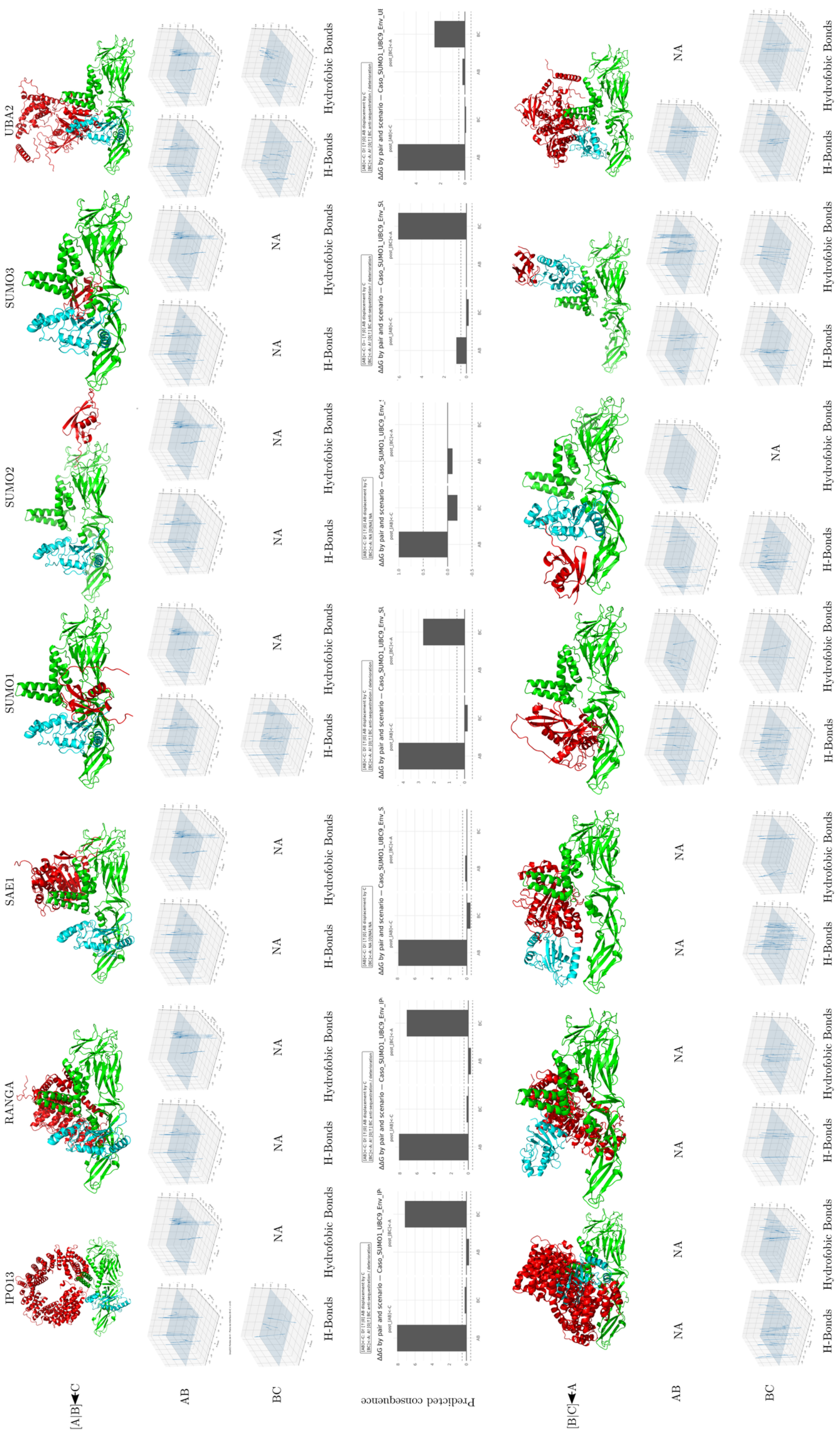
IPO13
RANGA
SAE1
SUMO1
SUMO2
SUMO3
UBA2
[A]B]◄C
AB
BC
Predicted consequence
[B]C]◄A
H-Bonds
Hydrofobic Bonds
NA

**Figure 27. Structural, interaction-field, and predicted energetic consequences of dengue Envelope–SUMO-centered ternary complexes with SUMO-associated proteins.**

The figure summarizes comparative ternary interaction models involving dengue virus Envelope, SUMO, and seven SUMO-associated partners: IPO13, RANGA, SAE1, SUMO1, SUMO2, SUMO3, and UBA2. In this representation, A corresponds to dengue Envelope, B corresponds to SUMO, and C corresponds to the additional SUMO-associated protein. Two alternative assembly scenarios are compared. In the first scenario, the Envelope–SUMO pair is treated as the preformed binary complex and the third protein is introduced afterward, represented as $[A]B \leftarrow C$. In the reciprocal scenario, the SUMO–associated-protein pair is treated as preformed and dengue Envelope is introduced as the third component, represented as $[B]C \leftarrow A$.

Structural panels show the relative spatial organization of the three proteins in each modeled ternary assembly, revealing pronounced partner-dependent differences in geometry and packing. Interaction-field plots report the spatial distribution of hydrogen-bond and hydrophobic contact features for the AB and BC interfaces, showing that polar and nonpolar interaction densities are redistributed according to both partner identity and assembly order. Predicted energetic consequence plots report $\Delta\Delta G$ values for each pair and assembly scenario, with negative values indicating stabilizing effects and positive values indicating destabilizing or energetically unfavorable effects relative to the corresponding reference state. Missing descriptors or unavailable configurations are indicated as NA.

Overall, the figure indicates that the dengue Envelope–SUMO interaction is strongly modulated by the molecular neighborhood of SUMO. Across several systems, including IPO13, RANGA, SUMO1, SUMO3, and UBA2, the introduction of the third partner into a preformed Envelope–SUMO complex preferentially perturbs the AB interface, whereas introduction of dengue Envelope into preformed SUMO-associated assemblies more strongly penalizes the BC interface. SAE1 and SUMO2 show comparatively more restricted or moderate effects, but still contribute to the overall partner-dependent heterogeneity of the ternary landscape.

Together, these results support a path-dependent model of ternary complex formation in which the structural and energetic outcome depends not only on the identity of the interacting proteins, but also on the order in which the viral and host components assemble, consistent with SUMO functioning as a network-embedded host interface within the broader sumoylation and transport-associated system.

##### 4.12.8 RANGAP1 Provides an Internal Biological Specificity Control

The RANGAP1 result occupies a privileged interpretive position in the SUMO1 panel because it provides an internal specificity control. SUMO1-modified RanGAP1 at the nuclear pore complex is one of the most structurally and biochemically well-characterised SUMO1-dependent assemblies in the cell: SUMO1 conjugation at Lys526 of RanGAP1 drives its stable targeting to RanBP2/Nup358 at the cytoplasmic face of the nuclear pore, where it forms a persistent trimeric complex with UBC9 [134–136]. The strong and selective AB co-operative stabilisation ($\Delta\Delta G \approx -9.4$ kcal · mol$^{-1}$) with no detectable reciprocal disruptive effect is precisely the conditional signature one would predict for a high-specificity SUMO1-dependent interaction: RANGAP1 draws the system into a deeply favoured SUMO1-centred configuration without generating off-axis perturbations. This result therefore validates the conditional framework — a well-established, high-affinity SUMO1 interaction partner produces the expected strong cooperative enrichment — while simultaneously confirming that the SUMO1–Env reference state is orders of magnitude less favourable than the SUMO1–RANGAP1 interaction geometry.

##### 4.12.9 E1 Machinery Subunits SAE1 and UBA2 Define the Activation-Cycle Attractor

The SAE1 and UBA2 cooperative signatures are mechanistically interpretable in terms of the E1 adenylation and thioester-transfer mechanism. The SUMO E1 recognises SUMO through an adenylation domain that contacts the SUMO $\beta$-grasp fold, followed by thioester transfer to the catalytic cysteine of UBA2 [122,123,131]. These interactions are structurally ordered and involve surfaces of SUMO1 that substantially overlap with the canonical effector and interactor surfaces. The near-identical AB-conditioned cooperative magnitudes of SAE1 and UBA2 ($\Delta\Delta G \approx -9.4$ kcal · mol$^{-1}$ each) — shared with RANGAP1 and the SUMO paralog panel — are consistent with a common feature: all high-affinity SUMO1 pathway partners converge on the same AB-axis stabilisation ceiling, reflecting the geometry of the SUMO1 surface that is optimised for pathway-relevant engagement.

The stronger reciprocal-branch contribution of UBA2 ($\Delta\Delta G \approx -4.1$) relative to SAE1 ($-1.7$) suggests that UBA2 stabilises the SUMO1 conformation more robustly across both conditioning orders, consistent with UBA2's direct thioester-bond contact with SUMO1 during the catalytic cycle representing a particularly intimate structural engagement compared to the more peripheral SAE1 contacts [131].

##### 4.12.10 IPO13 Introduces a Transport-Regulated Reorganisation Axis

IPO13 is a structurally atypical bidirectional karyopherin that imports UBC9, the SUMO E2 enzyme, into the nucleus [132,133]. Its conditional landscape is the most asymmetric in the SUMO1 panel — strong AB stabilisation ($-9.9$ kcal · mol$^{-1}$) at the expense of large BC destabilisation ($+7.2$ kcal · mol$^{-1}$) — indicating a rerouting effect rather than simple additive cooperativity: when a transport-associated SUMO component organises around SUMO1, the previous SUMO1-centred configuration (BC) becomes energetically disfavoured and the landscape collapses into the new preferred assembly. This pattern distinguishes IPO13 from the E1 subunits and RANGAP1, which stabilise AB without strongly disrupting BC, and reflects the structural reality that karyopherin-mediated transport involves extensive SUMO1 surface burial during import complex formation — a geometry that must necessarily exclude competing SUMO1 contacts.

From a viral biology perspective, this result suggests that the nuclear transport machinery regulating UBC9 subcellular localisation could indirectly modulate any SUMO1–Env interaction by controlling SUMO1 conformational state and availability within distinct cellular compartments. During dengue infection, redistribution of nuclear transport factors has been documented as a component of the broader host–virus conflict over nuclear access [27,206].

###### 4.12.11 SUMO Paralog Convergence Confirms Pathway-Authenticity as the Dominant Organising Principle

The SUMO1 self-association, SUMO2, and SUMO3 cooperative signatures are convergent in magnitude ($\Delta\Delta G \approx -9.4$ to $-9.7$ kcal · mol$^{-1}$) yet divergent in BC behaviour. SUMO3 produces pronounced BC destabilisation ($+6.1$ kcal · mol$^{-1}$), comparable in directionality to IPO13, whereas SUMO2 shows no informative BC state and SUMO1 self-association produces moderate BC weakening ($+2.7$). This gradient is consistent with known paralog-level biochemical differences: SUMO2 and SUMO3 bear an internal SUMO-interaction motif (SIM) that promotes poly-SUMO chain formation and paralog-specific downstream recognition, while SUMO1 lacks an efficient internal SIM and predominantly acts as a chain terminator [137,138]. These structural differences generate distinct SUMO–SUMO contact geometries that differentially perturb BC without changing the direction or magnitude of AB cooperative stabilisation.

The convergence of all three paralogs on the same $\approx -9.4$ to $-9.7$ kcal · mol$^{-1}$ AB cooperativity establishes that the key organising principle is SUMO pathway authenticity — access to SUMO-centred assemblies with native geometric complementarity — rather than paralog-specific biochemistry. Whichever SUMO paralog is present, the landscape reorganises decisively away from the SUMO1–Env reference state.

##### 4.12.12 Viral Engagement of the SUMO Network Is More Likely Through Pathway Regulation Than Direct Stable Binding

Taken together, the SUMO1 conditional landscapes converge on a biologically conservative but mechanistically precise conclusion: the dengue envelope is unlikely to interact with SUMO1 as a stable, receptor-like binding partner. The uniform dominance of canonical SUMO-pathway assemblies across the full partner set establishes a structural argument against a persistent SUMO1–Env complex under physiological conditions, where SUMOylation machinery, nuclear transport factors, and pathway substrates will consistently outcompete the weak reference Env-associated state.

This does not eliminate SUMO biology as relevant to dengue pathology — it reframes the relevant question. The SUMO pathway is an important regulatory hub for many cellular processes that dengue disrupts, including interferon signalling, nuclear transport, and transcriptional regulation [27,206]. Virological studies of dengue and related flaviviruses have most clearly implicated non-structural proteins — particularly NS5 — rather than the envelope protein in direct SUMO-pathway manipulation [206]. The present landscapes are consistent with that division: the envelope protein's interaction with SUMO1 reflects a peripheral, low-affinity contact that would operate only in the absence of competing SUMO machinery, while enzymatic and regulatory control of the SUMO network by non-structural viral proteins constitutes the more functionally dominant host–virus interaction axis.

From a methodological perspective, the SUMO1 panel also demonstrates an important property of the conditional $\Delta\Delta G$ framework: when the reference AB state is intrinsically weak, the framework reliably detects the energetic dominance of authentic pathway geometries, functioning as a quantitative readout of interaction hierarchy within the cellular network. The consistency of the $\approx -9.4$ kcal $\cdot$ mol$^{-1}$ AB-conditioned cooperativity across mechanistically diverse partners — E1 subunits, substrate complexes, karyopherins, and paralogs — represents an internal convergence that strengthens confidence in the conclusion, and distinguishes the SUMO1 result from artefactual signals that would be expected to show partner-specific scatter rather than pathway-wide uniformity [41,47,49].

#### 4.13 Envelope–TGFBR1: Asymmetric Conformational Exploration, Sliding Interface Consolidation, and TGF- Pathway Exploitation

##### 4.13.1 The Multi-Macrostate Exploration Behavior: An Unresolved Interaction Landscape in the Host Network

The TGFBR1 complex displays a trajectory profile that differs from all other complexes characterized in this study. The defining observation — an envelope protein that undergoes continuous multi-macrostate exploration without converging to a dominant basin within 200 ns, against a structurally invariant kinase receptor — distinguishes it from the other binding behaviors observed: the NRP1 b1-pocket insertion (bilateral conformational restriction), the GPC1 electrostatic saturation (directed single transition), the Rab5 two-registry pivot (two-state transition with continuous anchoring), the SUMO cooperative induced-fit (bilateral convergence to distinct late basins), and the clusterin/BiP/actin/betaglycan interfaces (all involving envelope transitions that stabilize within the simulation timescale). In the TGFBR1 system, the absence of late-time PCA convergence implies that the simulation captures an ensemble of transitional states rather than a thermodynamically equilibrated bound complex, and that the true binding geometry is likely to be reached only on timescales longer than 200 ns or under conditions (membrane proximity, physiological crowding, receptor clustering) that would constrain the envelope's conformational freedom [128].

The mechanistic consequence of this multi-macrostate behavior is that the contact-forming process is driven by kinetic exploration rather than thermodynamic minimization within the accessible timescale. Each successive envelope macrostate visited during the trajectory recruits new peripheral contacts with the TGFBR1 surface while maintaining the invariant anchor contacts mediated by the LYS296–ASP422–LYS337 electrostatic triad, generating the monotonically increasing contact count that characterizes this system. The observed multi-macrostate behavior may reflect either a shallow and rugged interaction landscape in which multiple binding geometries are near-degenerate in energy, or insufficient conformational sampling within the 200 ns timescale from an initial docking pose that may be suboptimal [209,210]. These two interpretations are not distinguishable from the present data and require extended simulations ($\geq 1$ µs) from multiple independent starting configurations to resolve. One possible conceptual framework — by analogy with the "fly-casting" model of intrinsically disordered protein recognition [196] — is that the progressive lateral exploration generates increasing contact area while maintaining an initial anchor set. However, this remains a speculative mechanistic hypothesis given the ambiguity of the non-converged trajectory.

##### 4.13.2 Energetic Paradox Resolved: Why More Contacts Produce Less Favorable Binding Energy

The apparent paradox of the TGFBR1 MM/GBSA trajectory — $\Delta G_{\text{total}}$ becoming less negative as contact count increases — is resolved by the multi-macrostate mechanism and provides one of the most mechanistically informative energetic signatures in this study. The temporal drift in VDWAALS from approximately −200 to −175 kcal/mol as contact number rises from approximately 6,883 to approximately 9,047 directly demonstrates that late-appearing contacts are geometrically suboptimal for van der Waals packing: they are close enough to fall within the contact cutoff (0.6 nm) but not close enough to be in the steep attractive region of the Lennard-Jones potential that generates deep VDWAALS stabilization. This is the quantitative energetic consequence of the envelope exploring progressively more peripheral regions of the TGFBR1 surface, where shape complementarity is less precise than at the initial contact core [49,117].

The simultaneous strengthening of per-residue contributions at the electrostatic hotspot residues (TDC reaching −35 to −45 kcal/mol in late frames) against the backdrop of weakening global VDWAALS is the energetic manifestation of interfacial reorganization: as the envelope slides and rolls, the charged anchor residues on TGFBR1 (LYS296, ASP422, LYS337) achieve progressively better geometric registration with their complementary envelope partners, increasing pairwise electrostatic contributions even as the overall packing quality of the expanding peripheral interface declines. This hotspot-strengthening concurrent with packing-weakening pattern is the energetic fingerprint of an interface that has not reached its global energy minimum — the system is still optimizing the positioning of its anchoring contacts while the peripheral contacts settle into geometrically inferior but numerically numerous configurations [43,198].

The MM/GBSA GGAS value of approximately −280 kcal/mol is the largest in the dataset, commensurate with the largest contact interface (>9,000 contacts). The correspondingly large EGB polar solvation penalty reflects the high proportion of polar and charged residues at the extensive TGFBR1–envelope interface, consistent with the lysine-rich kinase surface and the charged ectodomain regions of the viral protein. The ESURF nonpolar term provides a modest favorable contribution that tracks the hydrophobic burial component of the expanding contact area. The net $\Delta G_{\text{total}}$ of approximately −80 kcal/mol — larger than Rab5 (−40 kcal/mol) but smaller than GPC1 (−90 kcal/mol) — reflects the intermediate state of the interface: not yet at the thermodynamic minimum (explaining the drift), but robustly favorable throughout (explaining the persistent minimum distance and continuous contact maintenance).

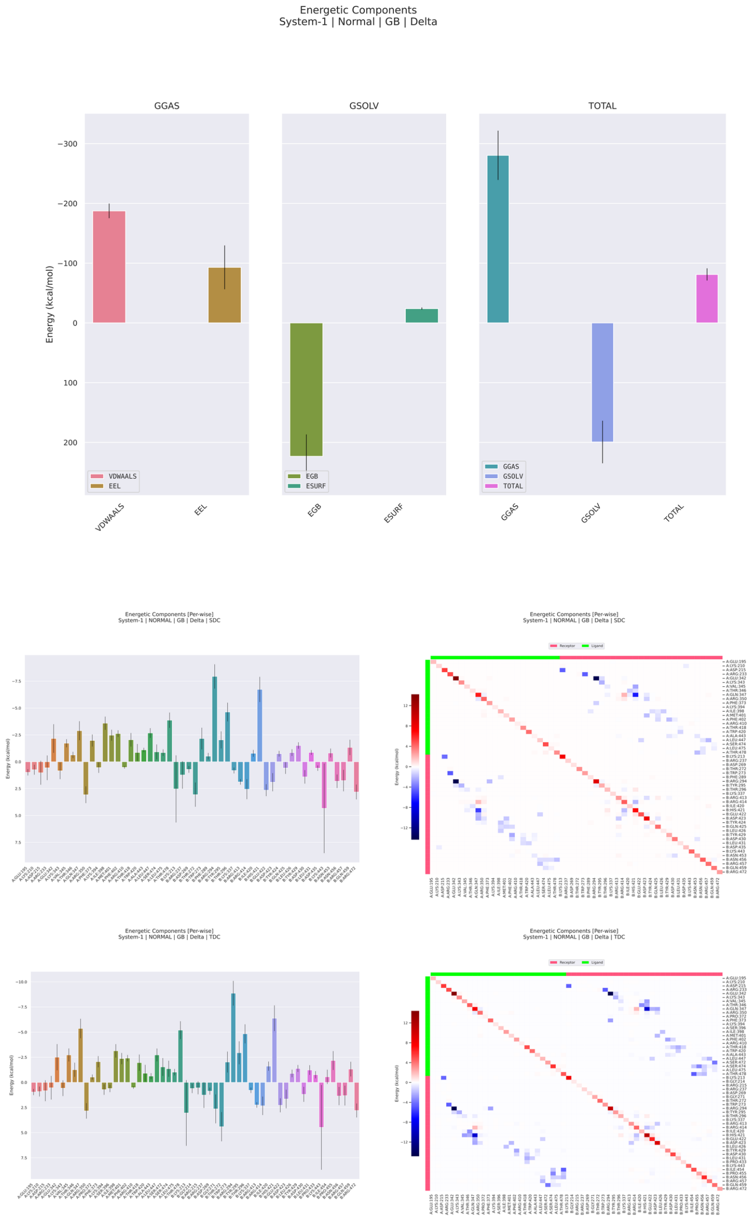


**Figure 28: MM/GBSA energetic decomposition of the dengue virus Envelope–TGFBR1 complex during molecular dynamics simulation.** The figure summarizes the global energetic components, per-residue decomposition, and pairwise residue decomposition of the binding free energy estimated by MM/GBSA for the dengue Envelope–TGFBR1 complex. The upper panels show that the gas-phase contribution is favorable, with both van der Waals and electrostatic terms contributing to stabilization. In contrast to complexes dominated almost exclusively by electrostatics, the Envelope–TGFBR1 interaction displays a more balanced gas-phase profile, in which short-range packing and electrostatic complementarity both contribute meaningfully to the interaction. The solvation component is unfavorable and dominated by the polar generalized Born term, indicating that complex formation imposes a substantial desolvation penalty on polar and charged groups at the interface. The nonpolar surface-area term contributes weakly relative to the dominant gas-phase and polar solvation terms. Despite this compensation between favorable gas-phase interactions and unfavorable polar solvation, the total binding free energy remains favorable, supporting the energetic stability of the dengue Envelope–TGFBR1 complex after solvent effects are incorporated. The middle and lower panels show per-residue and pairwise decompositions using SDC and TDC schemes, revealing a heterogeneous interface in which most residues contribute weakly or moderately, whereas a restricted subset of residues concentrates the strongest stabilizing or destabilizing effects. Negative values indicate favorable contributions to binding, whereas positive values indicate unfavorable or destabilizing energetic terms. The pairwise heatmaps further indicate that favorable interactions are sparse and localized, rather than distributed uniformly across the full interface. Blue regions denote favorable residue-pair interactions, whereas red regions denote unfavorable contributions. Overall, the MM/GBSA profile indicates that the Envelope–TGFBR1 interface is a hotspot-driven binding system supported by a combination of van der Waals packing and electrostatic complementarity, partially opposed by polar desolvation. This energetic organization suggests that TGFBR1 recognition by dengue Envelope is maintained by selected residue-level and pairwise contacts embedded within a broader, partially compensated interfacial landscape.

##### 4.13.3 TGF- Pathway Modulation as a Dengue Pathological Amplifier

The biological significance of a dengue envelope–TGFBR1 interaction must be evaluated in the context of TGF-$ 1′*swell−documentedroleindengueimmunopathology.TGF−* 1*plasmalevelsareelevatedduringdengueinfectionandco*[illegible] signalling through the TGFBR1–SMAD2/3 axis modulates endothelial cell junction integrity, inflammatory cytokine production, and T cell suppression, all of which are relevant to the clinical manifestations of severe dengue [140]. If the dengue envelope protein directly engages TGFBR1, several non-exclusive pathogenic mechanisms become structurally plausible based on the interaction geometry characterized here.

Several non-exclusive structural hypotheses may be proposed based on the interaction geometry characterized here, each requiring experimental validation before functional pathway modulation can be claimed [140,154]. First, direct envelope–TGFBR1 contact at the extracellular domain could in principle sterically influence the geometry available for TGF-$ 1*bindingorTGFBR*2*recruitment*; *theelectrostatichotspottriad*(*LYS*296, *ASP*422, *LYS*337)*id*[illegible] *independentreceptorclusteringmechanismsdescribedforotherreceptors*, *butonethatwouldrequirereceptorphosphorylationa*[illegible] *macrostateconformationaldynamicsobservedinthesimulationcouldinprinciplegenerateanatypicalreceptoroccupancystate*, [illegible] signalling.

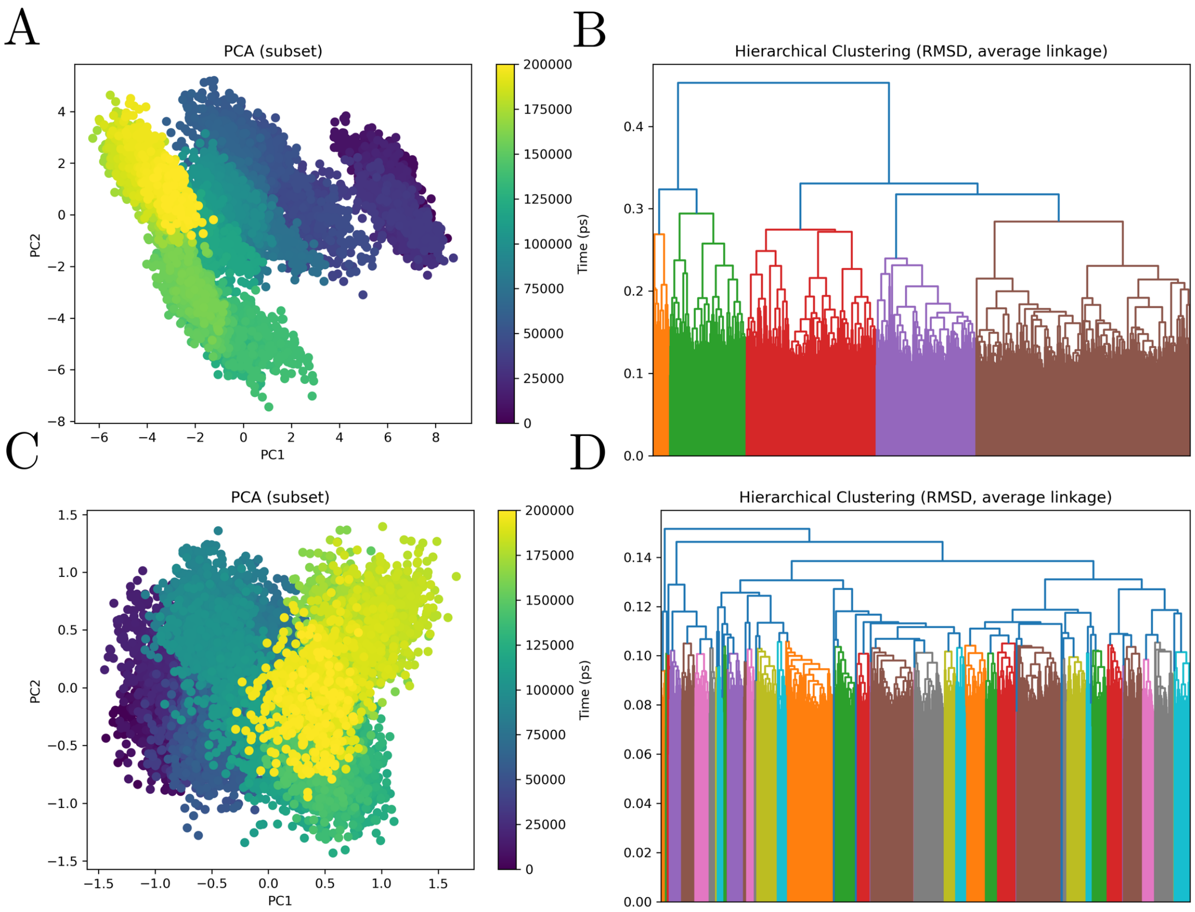


**Figure 29: Principal component analysis and RMSD-based hierarchical clustering of the dengue Envelope–TGFBR1 molecular dynamics trajectory.** The figure summarizes the conformational landscape sampled during the molecular dynamics simulation of the dengue Envelope–TGFBR1 complex, with panels A and B corresponding to the receptor subset, dengue Envelope, and panels C and D corresponding to the ligand subset, TGFBR1. Panels A and C show principal component analysis projections of trajectory frames onto the first two principal components, with points colored according to simulation time. In panel A, the dengue Envelope subset displays a marked directional redistribution across the conformational landscape, with early frames occupying a distinct region at positive PC1 values and later frames shifting toward negative PC1 values and higher PC2 values. This temporal organization indicates that the receptor undergoes substantial conformational remodeling during the simulation rather than remaining confined to a single static basin. The partially continuous transition between early, intermediate, and late regions suggests progressive adaptation of the Envelope within the TGFBR1-bound state. In panel C, the TGFBR1 subset displays a more compact but densely populated conformational manifold, with extensive overlap between temporal regions and a gradual redistribution from negative to positive PC1 values. This pattern indicates that the ligand explores a constrained set of related substates while still participating in the time-dependent evolution of the complex. Panels B and D show RMSD-based hierarchical clustering using average linkage. The receptor dendrogram in panel B reveals multiple structural families with clear separation among conformational groups, supporting the PCA-based interpretation that the Envelope samples several organized substates during the trajectory. The TGFBR1 dendrogram in panel D shows a highly subdivided clustering pattern with many closely related branches at relatively low RMSD distances, consistent with a compact but finely heterogeneous ligand ensemble. Together, the PCA and clustering analyses indicate that the dengue Envelope–TGFBR1 complex is dynamically heterogeneous but structurally organized, with the receptor undergoing broader directional remodeling and the ligand sampling a dense ensemble of related conformational substates. Overall, these results support a model in which TGFBR1 binding is associated with progressive remodeling of the Envelope-containing complex, generating an ensemble of related conformational states rather than a single rigid docking pose.

##### 4.13.4 Comparative Position of TGFBR1 in the Host Network: The Conformationally Inexhaustible Interface

Among all complexes in this study, TGFBR1 defines a unique position along the axis of conformational sampling completeness. At one extreme, NRP1 represents complete bilateral conformational restriction, where both partners reach their binding geometry within the first tens of nanoseconds and the PCA shows minimal dispersion throughout. At the other extreme, TGFBR1 represents an incomplete, continuously exploring interface where the 200 ns simulation fails to converge the envelope to a dominant conformational basin and the contact number has not reached a plateau by the end of the trajectory. This incomplete sampling is not a limitation of the analysis but a genuine physical property of the system: the envelope's multi-domain architecture provides a conformational space that is too large to be exhausted on the 200 ns timescale when exploring the broad TGFBR1 surface, and the kinetic barriers between envelope macrostates are sufficiently small to permit continuous exploration without trapping. The practical implication is that the true $\Delta G_{\text{bind}}$ for this complex is likely more negative than the $-80$ kcal/mol estimate from the 200 ns endpoint calculation, because the interface has not yet reached the configuration of maximum contact complementarity. Among all complexes in this study, the TGFBR1 interaction therefore represents the one with the greatest potential for affinity improvement upon longer-timescale equilibration or cooperative structural stabilization in the cellular environment — and correspondingly the highest uncertainty in the MM/GBSA-derived binding free energy estimate [49,84].

##### 4.13.5 Ternary Landscape Positions the TGFBR1–Env State as a Conditional Receptor Configuration Reliably Superseded by Endogenous Pathway Assemblies

The conditional $\Delta\Delta G$ analysis of the TGFBR1-centred system establishes a pattern of striking uniformity: all ten tested partners — spanning ligands, type II receptors, intracellular effectors, inhibitory regulators, and ubiquitin-associated mediators — produce large, directionally concordant AB cooperative stabilisations ($\Delta\Delta G \approx -10.0$ to $-10.7$ kcal $\cdot$ mol$^{-1}$) in the AB-conditioned branch. This convergence across mechanistically diverse partner classes constitutes a strong computational signal that the TGFBR1–Env reference state ($\Delta G \approx -6.4$ kcal $\cdot$ mol$^{-1}$) is not an energetically privileged endpoint but a conditional configuration that is reliably displaced by canonical TGF-$\beta$ superfamily pathway components [140,145,146]. TGFBR1/ALK5 functions physiologically not as an isolated receptor but as the obligate type I kinase subunit within heteromeric signalling assemblies that are assembled through ordered ligand-binding, type II receptor recruitment, GS-domain release, and intracellular Smad engagement — a hierarchically organised interaction logic that, as these results show, consistently outcompetes any non-canonical TGFBR1-centred state [139,211].

##### 4.13.6 Cognate Ligands and Type II Receptors Define the Core Canonical Displacement Axis

TGFB1, TGFB3, TGFBR2, ACVR2B, and GDF11 represent the most biologically decisive signals in the dataset because they are the direct extracellular interaction partners that initiate TGFBR1 signalling complex assembly. TGFBR2 is the canonical type II receptor that transphosphorylates the TGFBR1 GS domain upon ligand-induced heterotetrameric complex formation [139,146]; its ternary signature — strong AB-conditioned cooperativity ($-10.5\ \text{kcal} \cdot \text{mol}^{-1}$) paired with pronounced BC destabilisation (+7.0) and a substantial reverse-conditioned AB gain ($-5.3$) — is precisely the pattern expected for a receptor whose most favourable surface configuration is structurally incompatible with simultaneous Env engagement: TGFBR1 in a TGFBR2-compatible conformation buries the surfaces that Env contacts. TGFB1 and TGFB3 produce selective AB-axis cooperativity without BC disruption, consistent with ligand-induced receptor assembly geometries in which the ligand-binding surface and the Env-contact region are distinct but conformationally coupled [144,145].

ACVR2B and GDF11 extend this logic into the broader TGF-$\beta$ superfamily context. ACVR2B is a type II activin receptor that pairs with TGFBR1/ALK5 and related type I receptors for activin, myostatin, and GDF-class ligands, and structural studies have established that activin class receptor assemblies follow a conserved heterotetrameric binding topology [212,213]. GDF11 signals through activin type II receptors in combination with ALK4/5/7 type I receptors and has been shown to activate SMAD2/3 through TGFBR1 in multiple cell types [214]. The concordance between ACVR2B and TGFBR2 cooperative magnitudes establishes that this pattern reflects a shared TGFBR1 conformational logic across type II receptor paralogs rather than TGFBR2-specific effects — strengthening the interpretation that TGFBR1 AB cooperativity is driven by shared TGF-$\beta$ superfamily receptor assembly geometry [215].

##### 4.13.7 FKBP1A Demonstrates That Cooperative Stabilisation Reflects Structural Ordering Rather Than Signalling Activation

The FKBP1A result is mechanistically the most interpretively important in the TGFBR1 panel. FKBP1A (FKBP12) is an inhibitory adaptor that binds the Leu-Pro dipeptide of the unphosphorylated GS domain of type I TGF-$\beta$ family receptors, sterically occluding Smad access and maintaining the receptor in a constitutively inactive state until type II receptor-mediated GS phosphorylation releases this inhibition [139,211]. FKBP1A therefore does not activate TGFBR1 — it constrains it. Yet it produces the strongest bilateral cooperative signal in the TGFBR1 panel: AB-conditioned $\Delta\Delta G \approx -10.1$ paired with a reverse-conditioned $\Delta\Delta G \approx -7.1$ — the largest reciprocal-branch cooperativity in the dataset.

The mechanistic implication is critical: the conditional framework detects structural ordering of the receptor into pathway-authentic conformations regardless of whether those conformations are activating or inhibitory. FKBP1A stabilises TGFBR1 in the pre-activation inhibited state, which is geometrically well-defined and biochemically stable; the resulting TGFBR1 conformation is highly ordered and presents a surface that is structurally incompatible with the Env-associated configuration. This result establishes that what the conditional maps detect is not signalling competence but pathway-authentic receptor geometry — and that even the inhibited, pre-activation TGFBR1 state is strongly incompatible with stable Env engagement. Biologically, this means that under resting cellular conditions, where the majority of TGFBR1 is FKBP1A-associated and in the inactive conformation, the TGFBR1–Env interaction would be energetically disfavoured relative to the dominant receptor state.

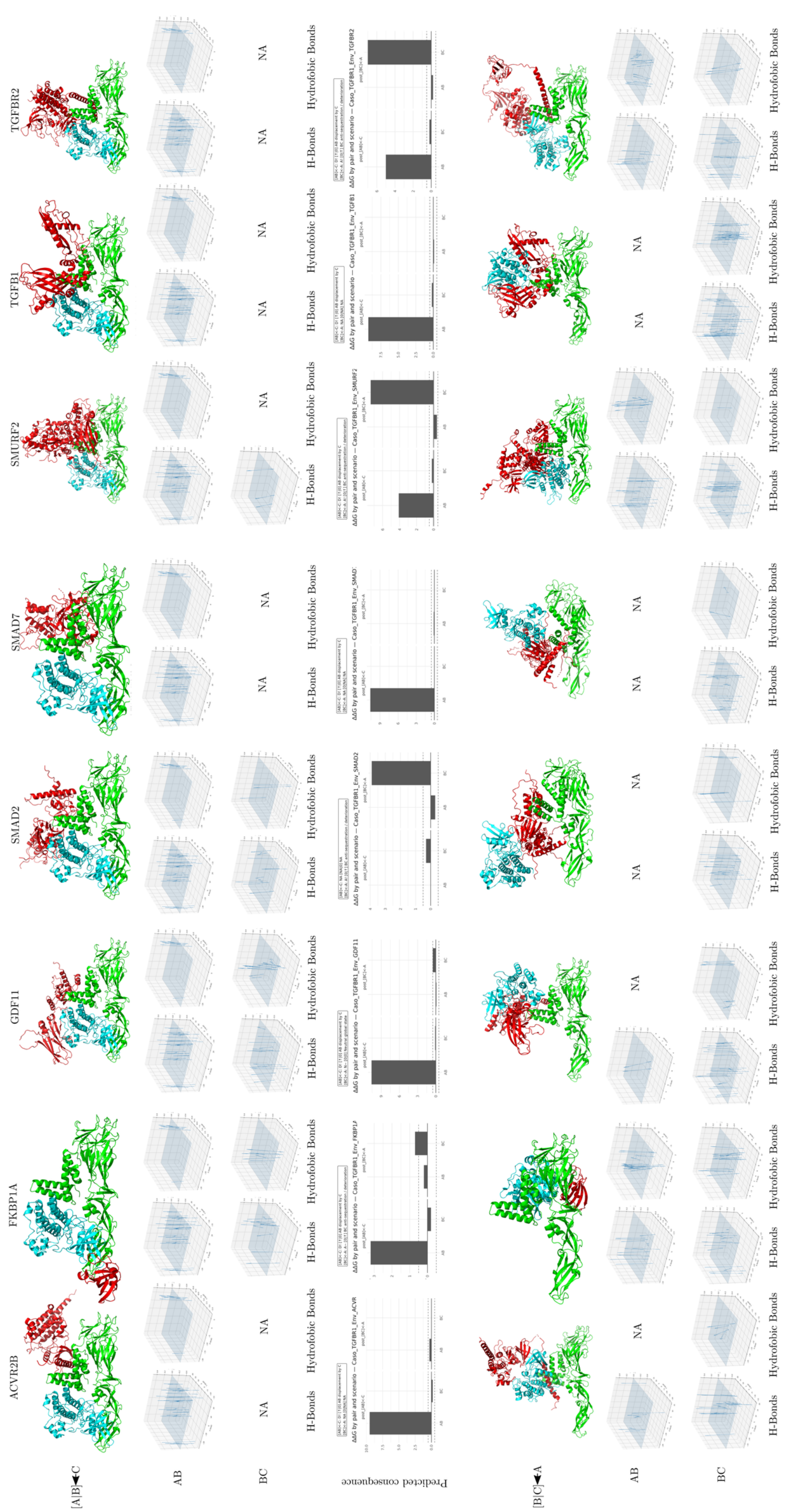
ACVR2B
FKBP1A
GDF11
SMAD2
SMAD7
SMURF2
TGFB1
TGFBR2
[A|B]◄C
AB
BC
NA
H-Bonds
Hydrofobic Bonds
Predicted consequence
[B|C]◄A

**Figure 30. Structural, interaction-field, and predicted energetic consequences of dengue Envelope–TGFBR1-centered ternary complexes with TGFBR1-associated proteins.**

The figure summarizes comparative ternary interaction models involving dengue virus Envelope, TGFBR1, and eight TGFBR1-associated partners: ACVR2B, FKBP1A, GDF11, SMAD2, SMAD7, SMURF2, TGFB1, and TGFBR2. In this representation, A corresponds to dengue Envelope, B corresponds to TGFBR1, and C corresponds to the additional TGFBR1-associated protein. Two alternative assembly scenarios are compared. In the first scenario, the Envelope–TGFBR1 pair is treated as the preformed binary complex and the third protein is introduced afterward, represented as $[A]B \leftarrow C$. In the reciprocal scenario, the TGFBR1–associated-protein pair is treated as preformed and dengue Envelope is introduced as the third component, represented as $[B]C \leftarrow A$.

Structural panels show the relative spatial organization of the three proteins in each modeled ternary assembly, revealing partner-dependent differences in geometry, packing, and exposure of the Envelope–TGFBR1 interface. Interaction-field plots report the spatial distribution of hydrogen-bond and hydrophobic contact features for the AB and BC interfaces, showing that polar and nonpolar interaction densities are redistributed according to both partner identity and assembly order. Predicted energetic consequence plots report $\Delta\Delta G$ values for each pair and assembly scenario, with negative values indicating stabilizing effects and positive values indicating destabilizing or energetically unfavorable effects relative to the corresponding reference state. Missing descriptors or unavailable configurations are indicated as NA.

Overall, the figure indicates that the dengue Envelope–TGFBR1 interaction is strongly modulated by the signaling neighborhood of TGFBR1. Several partners, including ACVR2B, FKBP1A, GDF11, SMAD7, SMURF2, TGFB1, and TGFBR2, impose marked assembly-order-dependent energetic effects, indicating that the structural and energetic outcome depends on whether Envelope binds TGFBR1 before or after formation of the host-associated TGFBR1 complex. SMAD2 and SMAD7 further highlight the importance of downstream signaling adaptors in reshaping the ternary landscape, whereas TGFB1 and TGFBR2 connect the Envelope–TGFBR1 interface to canonical ligand–receptor organization within the TGF-$\beta$ signaling axis.

Together, these results support a path-dependent model of ternary complex formation in which dengue Envelope recognition of TGFBR1 is not governed solely by the binary viral–host interface, but by the pre-existing molecular state of TGFBR1 within its signaling, regulatory, and receptor-associated protein neighborhood.

##### 4.13.8 Intracellular Smad and Ubiquitin Regulators Confirm TGFBR1 as a Signalling Hub Incompatible with Stable Env Association

SMAD2, SMAD7, SMURF2, and TRAF6 together cover the principal intracellular regulatory axes downstream of TGFBR1 activation. SMAD2 is the canonical receptor-regulated Smad that is phosphorylated by activated TGFBR1, released, and complexed with SMAD4 for nuclear translocation and transcriptional regulation [140,145]. SMAD7 is the inhibitory Smad that competes with SMAD2 for receptor access and recruits SMURF2 to ubiquitinate TGFBR1 for lysosomal degradation [216,217]. SMURF2 is the HECT-domain E3 ubiquitin ligase that executes this receptor turnover and also directly associates with TGFBR1-containing complexes [217]. TRAF6 mediates the non-Smad TGF-$\beta$ signalling branch through TAK1/MAP kinase cascades [145].

Despite representing four mechanistically distinct functional modules — activation, inhibition, ubiquitin-mediated degradation, and non-canonical kinase signalling — all four produce AB-conditioned cooperative magnitudes within a 0.5 kcal $\cdot$ mol$^{-1}$ window ($-10.2$ to $-10.7$). This convergence is the conditional-docking signature of a receptor hub: a protein whose surface is functionally optimised for sequential engagement by diverse intracellular partners, all of which contact overlapping surface regions that are conformationally incompatible with simultaneous Env engagement. The variation in BC profiles — SMURF2 producing the strongest BC destabilisation (+7.4), SMAD7 producing near-neutral BC behaviour — reflects the different spatial footprints of these regulators on the intracellular TGFBR1 surface but does not alter the dominant biological conclusion [216,217].

##### 4.13.9 A Pathway-Network Model for TGFBR1 Involvement in Dengue Pathology

Collectively, the TGFBR1 conditional landscapes support a receptor-network model that reframes the biological role of the TGFBR1–Env interaction identified in the binary MD analysis. The binary interaction establishes that TGFBR1 can form a structurally stable, energetically favourable pairwise complex with the dengue envelope; the conditional analysis establishes that this complex is not energetically dominant once canonical TGFBR1-pathway partners are present. The reconciliation of these two observations leads to the conclusion that TGFBR1–Env engagement is most plausible in the narrow receptor-availability window defined by low local concentrations of cognate ligands, type II receptors, and FKBP1A — conditions that would favour a free, pathway-uncommitted TGFBR1 surface. In practice, such windows are likely transient and spatially heterogeneous, consistent with a model in which TGFBR1–Env contact occurs at specific membrane microdomains or trafficking intermediates rather than as a sustained, dominant receptor–virus interaction [146,215].

This network-context model has a specific implication for dengue pathology. The strongest documented pathway disruption associated with dengue infection is impairment of TGF-$\beta$/SMAD signalling in endothelial and immune cells, contributing to vascular hyperpermeability and dysregulated cytokine production [140,154]. The conditional analysis suggests two complementary mechanisms by which Env could contribute to this disruption: direct competition with cognate ligand or type II receptor engagement at the TGFBR1 ectodomain (most plausible in low-ligand microenvironments), and indirect effects through TGFBR1 conformational perturbation that reduces receptor availability for intracellular Smad engagement. Neither mechanism requires a stable, prolonged Env–TGFBR1 complex; both are consistent with the transient, context-dependent interaction window defined by the conditional landscape hierarchy.

As with all ternary systems characterised here, the conclusions are bounded by the limitations of static docking-derived $\Delta\Delta G$ estimates, which do not capture membrane-embedded receptor stoichiometry, ligand latency and activation, glycosylation-dependent receptor accessibility, or kinetic competition rates in the crowded endosomal environment. Experimental validation — particularly TGF-$\beta$ pathway reporter assays in dengue-infected endothelial cells and biochemical competition assays between Env and TGFB1 for TGFBR1 binding — represents the necessary next step for this system [41,47,49].

### 5 Conclusions

This study presents a multiscale computational characterization of the structural interaction landscape between sixteen dengue virus envelope genotype variants and thirteen host proteins implicated in viral attachment, endocytic trafficking, immune signaling, and vascular regulation. By integrating protein–protein docking, molecular dynamics simulations, MM/GBSA energetic analysis, and vectorial characterization of hydrogen bond networks, the framework reveals consistent structural patterns that provide mechanistic context for understanding dengue virus–host protein engagement, subject to the approximations inherent to each computational method employed.

#### 5.1 Asymmetric Conformational Adaptation as a Cross-Interface Conserved Feature

Across all characterized complexes, the molecular dynamics trajectories reveal a recurring pattern of asymmetric conformational adaptation in which the viral envelope protein functions as the structurally rigid partner — with backbone RMSD values typically converging to 0.4–0.7 nm over 200 ns — while host proteins exhibit greater conformational plasticity, with localized RMSF peaks in interface-proximal regions. This partitioning is consistent with the established structural robustness of the flavivirus E protein, whose $\beta$-rich architecture confers conformational stability necessary for the pH-triggered fusion transition during viral entry [3,5], and suggests that the dengue envelope presents structurally defined binding determinants that host proteins accommodate through localized flexibility rather than through global refolding.

#### 5.2 Electrostatically Dominated Interfaces and Distributed Energetic Architecture

MM/GBSA analysis revealed that envelope–host interfaces are predominantly stabilized by electrostatic interactions, with gas-phase EEL contributions ranging from approximately $-180$ to $-750$ $\text{kcal} \cdot \text{mol}^{-1}$ across complexes, substantially offset by polar solvation penalties. The resulting net $\Delta G$ values should be interpreted as semi-quantitative rankings rather than absolute binding free energies, given the known limitations of the MM/GBSA approximation for highly charged interfaces [49,84]. Three energetic regimes were identified: electrostatically saturated interfaces (ENV–GPC1, $\Delta G_{\text{total}} \approx -90$ $\text{kcal} \cdot \text{mol}^{-1}$), electrostatically anchored interfaces (ENV–BiP, $\approx -70.5$ $\text{kcal} \cdot \text{mol}^{-1}$), and hydrophobically dominated interfaces (ENV–Clusterin). Per-residue decomposition consistently revealed a distributed energetic architecture spanning multiple spatially separated residue clusters on both partners, rather than single dominant hotspots [43]. For interactions with highly conserved cellular scaffolds such as actin and BiP, this energetic redundancy may contribute to robustness of host factor engagement across the sixteen genotype variants examined, though experimental mutagenesis studies would be required to confirm this interpretation.

#### 5.3 Host Protein-Specific Interaction Mechanisms

Glypican-1 generated the largest absolute electrostatic stabilization and presented the most electrostatically saturated interface profile, with favorable per-residue contributions concentrated in the DIII electropositive patch, consistent with the established role of DIII as the primary heparan sulfate-binding site in flaviviruses. The core protein component characterized here provides a baseline interaction energy that would likely be augmented by the full heparan sulfate-decorated proteoglycan in physiological conditions.

The ENV–BiP interface was characterized by a discrete electrostatic anchor involving A:ARG410 and B:GLU498 ($\approx -3.1$ to $-3.4$ $\mathrm{kcal \cdot mol^{-1}}$ per-residue contribution) supplemented by hydrophobic packing, with an interaction mode — stable yet dynamically permissive — consistent with a co-receptor role in which cell-surface BiP increases viral residence time without irreversibly sequestering the particle. The contact geometry and identity of ARG410 as the primary anchor constitute a specific, experimentally testable structural hypothesis.

The ENV–NRP1 complex displayed bilateral conformational restriction and energy concentration in a terminal basic residue–acidic cage arrangement qualitatively resembling characterized CendR–NRP1 complexes [103,108], supporting the hypothesis that the dengue envelope may engage NRP1 through SLiM-based molecular mimicry — a strategy documented across multiple viral families [148,191]. This hypothesis requires direct experimental confirmation through competition assays and mutagenesis of the candidate CendR-like loop.

#### 5.4 Ternary Complex Analysis and Contextual Modulation of Viral Binding

Implementation of the AB/BC/ABC conditional energetic framework for actin-associated host proteins demonstrated that assembly order can substantially influence the thermodynamic outcome of viral–host engagement, with conditional $\Delta\Delta G$ values ranging from approximately $-3.4$ to $+6.9$ $\mathrm{kcal \cdot mol^{-1}}$ depending on the neighbor protein and assembly sequence [47,52]. Classical actin regulators (ACTN2, PFN1) produced destabilizing effects consistent with competitive occupation of shared actin determinants, whereas dynactin-associated components (ACTR1A, ACTR10) showed patterns consistent with cooperative stabilization of the viral interface. These observations suggest that the cellular cytoskeletal environment at the time of viral encounter may influence the thermodynamic favorability of actin engagement — a prediction that is directly testable through targeted knockdown experiments and infection assays in cells with perturbed cytoskeletal composition [15,150].

#### 5.5 Genotype Conservation and the Scope of the Genotype-Informed Framework

The designation "genotype-informed" in the study title reflects the construction and systematic docking of sixteen distinct three-dimensional structural variants representing the genetic diversity of circulating dengue envelope genotypes, enabling a structural survey that spans the sequence space of DENV1–4 and selected intraserotype genotypic variants [1,147]. This panel provides the dataset from which interface conservation can be assessed: the basic residue clusters mediating electrostatic interactions — particularly in the DIII electropositive patch — were structurally present across all sixteen variants examined, as was the structural position corresponding to ARG410 in the BiP complex and the candidate CendR-like loop residue cluster in the NRP1 complex. This conservation across the genotype panel suggests that the principal interaction architectures characterised here are not idiosyncratic properties of a single representative structure but are broadly encoded across the circulating sequence space.

A critical qualification, however, must accompany this claim. The molecular dynamics trajectories and MM/GBSA analyses reported in the Results sections were conducted on a single representative envelope structure for each host protein complex, not on all sixteen genotype variants. Quantitative comparison of $\Delta G$ magnitudes, hotspot contributions, and conformational dynamics across the full genotype panel — the information that would most directly justify the term "genotype-informed" in a quantitative sense — remains an important deferred objective. Executing 200 ns trajectories for each of the sixteen variants against each of thirteen host proteins would constitute a large-scale computational undertaking (approximately $16 \times 13 = 208$ additional independent simulations) that was outside the scope of the present study. Accordingly, the genotype-informed component of this work is most accurately interpreted as a structural diversity survey establishing that the key binding determinants are conserved across the panel, with full quantitative affinity differentiation across genotypes as the recommended next step for future studies.

To partially address this gap, serotype-resolved vectorial surface fitting using the VectorPROT pipeline was applied to aggregate interaction vectors by genotype index, confirming that the curvature and orientation of the fitted quadratic surface descriptors are broadly concordant across genotypic clusters — consistent with structural conservation of the interaction geometry — while revealing modest but detectable inter-genotype differences in the directional distribution of hydrogen-bond vectors, particularly at the DIII surface. These vectorial differences, though sub-threshold for individual genotype–genotype $\Delta G$ comparisons without full MD trajectories, represent testable structural hypotheses for serotype-specific differences in attachment efficiency that could be evaluated using surface plasmon resonance or biolayer interferometry with purified envelope ectodomains from representative genotypes [1,218].

#### 5.6 Outlook

This study provides a computationally derived structural hypothesis framework for dengue envelope engagement with host proteins, identifying candidate attachment factors, trafficking partners, and signaling-associated interfaces that warrant experimental validation. The computational framework generates a set of prioritized, experimentally tractable structural hypotheses regarding the molecular determinants of dengue virus–host engagement.

Priority experimental directions include: (1) site-directed mutagenesis and SPR-based affinity measurement at the ARG410 hotspot in the ENV–BiP interface, to test the discrete electrostatic anchor model and establish a quantitative baseline for inhibitor design; (2) competition assays and cryo-EM structural determination for the ENV–NRP1 complex to test the CendR-like mimicry model, including CendR peptide displacement assays to confirm pocket occupancy and mutagenesis of the candidate loop residues; (3) CRISPR-based knockdown of ACTR1A and ACTR10 combined with dengue infectivity assays to assess the functional relevance of the predicted dynactin-cooperative interface with actin; (4) live-cell single-virion tracking experiments in GPC1 knockout versus overexpressing cell lines to test the surf-riding dimensionality-reduction model, with lateral diffusion coefficient and membrane dwell time as primary readouts; (5) Ubc9 catalytic activity assays in the presence of recombinant dengue envelope ectodomain — specifically using STAT1 and STAT2 SUMO-acceptor peptide substrates — to distinguish allosteric modulation from competitive inhibition of the SUMOylation cascade; (6) fragment-based and virtual screening campaigns targeting the LYS435/LYS516/LYS447 electrostatic frustration cluster in BiP and the ARG107/LYS236 cluster in betaglycan as priority pharmacophore sites for viral entry inhibitor design; (7) cross-flavivirus NRP1 engagement profiling for Zika, West Nile, and Yellow fever envelope proteins to test the convergent geometric mimicry hypothesis and establish whether b1-pocket engagement is a shared flavivirus strategy; and (8) application of block-averaged MM/GBSA and free energy perturbation (FEP) methods to the ENV–BiP, ENV–NRP1, and ENV–Rab5 hotspot residue pairs for higher-accuracy affinity estimates that can be directly confronted with isothermal titration calorimetry or microscale thermophoresis measurements [49,219].

Collectively, the structural insights generated by this study — including the identification of electrostatic frustration sites as pharmacological opportunities, the proposal of GPC1 as a dimensionality-reduction platform for surface-diffusion-assisted receptor search, the framing of NRP1 b1-pocket engagement as convergent structural mimicry among vascular pathogens, and the mechanistic hypothesis of allosteric Ubc9 modulation — contribute to an emerging picture in which the dengue envelope protein exploits structurally diverse host interfaces through mechanistically distinct modes, each linked to a specific stage of the viral lifecycle and each amenable to disruption through molecularly targeted intervention strategies.

### 6 Limitations

The computational framework presented in this study provides structural and energetic characterizations of dengue virus envelope–host protein interactions, but several methodological constraints must be considered when interpreting the results and evaluating their biological implications.

#### 6.1 Single-Conformation Rigid-Body Docking

All protein–protein docking simulations were conducted using single representative three-dimensional structures with the HDOCK rigid-body algorithm [39]. This approximation does not capture conformational selection or induced-fit binding mechanisms, which are likely relevant for inherently flexible partners such as actin (G-actin/F-actin equilibrium) and clusterin (disordered substrate-binding tails). Community-wide benchmarking of free docking methods (CAPRI) has established that even state-of-the-art servers achieve acceptable models for approximately 40% of tested targets [117], and the present pipeline was not validated against a dedicated set of dengue envelope–host co-crystal structures. Ensemble docking approaches incorporating conformational diversity from MD pre-sampling would provide a more complete representation of the interaction landscape and are recommended for downstream refinement of priority interfaces [40].

#### 6.2 Absence of Virion Context and Membrane Environment

The dengue envelope protein exists on the virion surface not as a monomeric soluble ectodomain but as an ordered icosahedral lattice of 90 E-protein dimers, anchored to the lipid bilayer through C-terminal transmembrane segments [3–5]. All docking and MD simulations were conducted on soluble ectodomain constructs, neglecting: (i) orientational restrictions imposed by membrane anchoring; (ii) steric occlusion of ectodomain surfaces by dimer and lattice contacts; (iii) contributions of membrane-proximal stem residues to interface energetics; and (iv) the lateral diffusion-limited encounter kinetics governing receptor binding on the virion surface. Because the dengue envelope ectodomain is presented as part of a membrane-anchored oligomeric lattice, soluble ectodomain simulations may overestimate accessible interaction surfaces and underestimate steric constraints imposed by the virion context.

#### 6.3 Unmodeled Glycosylation at N67 and N153

The dengue envelope protein carries two conserved N-linked glycosylation sites, at Asn67 and Asn153, whose glycan chains modulate receptor recognition, immune evasion, and ectodomain stability [5,220]. None of the structural models incorporate these glycans, nor do they account for post-translational modifications on host proteins. Glycan exclusion is most consequential for interfaces proximal to these sites and may result in overestimation of accessible contact area and altered local electrostatics. Glycan-inclusive docking and MD are recommended for any follow-up characterization of the predicted interfaces, particularly for GPC1 and NRP1 where steric masking by glycans could substantially affect predicted geometries.

#### 6.4 Molecular Dynamics Timescale and Convergence

The 200 ns production trajectories are sufficient for characterizing local conformational adjustments and early interfacial relaxation, but do not provide information about complex dissociation kinetics or thermodynamic stability on biologically relevant (microsecond to millisecond) timescales. The ENV–TGFBR1 complex in particular does not achieve PCA convergence within 200 ns, meaning the MM/GBSA estimates for this system represent an ensemble of non-equilibrated states rather than a thermodynamically stable bound complex [209,210]. All statements about interface persistence should be understood as observations within the simulation window rather than assertions of thermodynamic stability.

#### 6.5 MM/GBSA Energetics: Comparative Utility and Quantitative Limitations

Binding free energies were estimated using the MM/GBSA endpoint approximation, which omits configurational entropy and uses an implicit GB solvation model. Published benchmarks report Pearson correlations of $R^2 = 0.3$–$0.6$ between MM/GBSA values and experimental affinities for diverse datasets, with largest systematic errors for highly charged, solvent-exposed interfaces [49,50]. Absolute $\Delta G$ values exceeding $-100$ $\text{kcal} \cdot \text{mol}^{-1}$ (e.g., ENV–GPC1) reflect MM/GBSA overestimation from imperfect EEL–EGB cancellation at electrostatically saturated interfaces and do not correspond to physically measurable affinities. The interpretive strength of MM/GBSA in this study therefore resides primarily in per-residue decomposition (identification of energetic hotspots and frustration sites) and in comparative ranking of interaction states within structurally similar interface classes, rather than in absolute affinity predictions.

No formal uncertainty quantification was performed: the reported $\Delta G$ values were computed from single trajectory windows of the last 10 ns of each production simulation without block averaging, bootstrapping, or sub-ensemble decomposition. Consequently, confidence intervals for the reported values cannot be provided, and the statistical significance of small energetic differences (e.g., $|\Delta\Delta G| < 1$ $\text{kcal} \cdot \text{mol}^{-1}$) cannot be assessed from the present dataset. The temporal fluctuations in MM/GBSA profiles visible in several complexes (most prominently Rab5 and TGFBR1) represent either genuine microstate interconversion or sampling noise, and the two cannot be distinguished without block-decomposition analysis. Future applications of this pipeline should implement block averaging over non-overlapping sub-windows of the analysis trajectory (e.g., five consecutive 2 ns blocks across the final 10 ns), followed by standard error estimation from the block means, to provide reproducible confidence intervals consistent with best practices in MM/GBSA reporting [49,50,84]. For the highest-priority complexes — ENV–BiP, ENV–NRP1, and ENV–Rab5 — application of the more rigorous MM/PBSA or free energy perturbation (FEP) methods to the specific hotspot residue pairs identified by per-residue decomposition would yield thermodynamic estimates with experimentally verifiable accuracy [49,219].

#### 6.6 Pairwise Decomposability of the Ternary Framework

The AB/BC/ABC conditional energetic framework assumes that the ternary interaction free energy can be approximated by sums of pairwise binary contributions, which may underestimate higher-order cooperative effects when all three proteins are in simultaneous contact [41,51]. Ternary assemblies were generated by sequential docking without subsequent MD relaxation; conformational strain at the newly formed interface may exaggerate destabilizing $\Delta\Delta G$ values (particularly for PFN1 and CAPZB), and initial steric clashes in rigid-body docked ternary geometries may generate spuriously large positive $\Delta\Delta G$ estimates that would be substantially attenuated upon side-chain relaxation.

To validate the directional trends reported here for the most consequential ternary interactions, short MD simulations (20–50 ns) of selected assembled ternary complexes are recommended as a priority for future work. Specifically: (i) the PLXNA4-excluded NRP1–Env–PLXNA4 assembly, which showed the largest competitive AB destabilisation in the NRP1 panel and for which the directionality of exclusion has direct implications for endothelial NRP1 signalling context; (ii) the IPO13-reorganised SUMO1–Env–IPO13 assembly, which exhibited the most asymmetric landscape in the SUMO1 panel and for which allosteric rerouting through the nuclear transport axis is the proposed mechanism; and (iii) the HBEGF-displaced GPC1–Env–HBEGF assembly, which displayed the strongest glycocalyx competition signal and whose validation would directly test the surf-riding hypothesis for GPC1-mediated attachment. Short MD validation of these three ternary complexes would confirm whether the rigid-docking-derived $\Delta\Delta G$ values reflect genuine thermodynamic trends or artefacts of initial contact geometry, and would provide trajectory-level evidence for or against the biological models proposed in the Discussion [40,47].

#### 6.7 Absence of Macromolecular Crowding

All simulations were conducted in dilute solvent conditions. The densely crowded cytoplasmic environment substantially alters effective association kinetics, protein conformational landscapes, and thermodynamic balances between binding states relative to dilute-solution predictions [71]. Excluded-volume effects under crowding conditions generally favor compact associated states, implying that cellular affinities could differ systematically from the values reported here.

### 7 Author Contributions

Daniel Ferreira de Lima Neto and Caio Cesar de Melo Freire contributed equally to this work and share corresponding authorship. Daniel Ferreira de Lima Neto conceived and designed the study, developed the complete computational framework, implemented the analytical methodologies, performed molecular docking experiments, molecular dynamics simulations, and energetic analyses, curated and interpreted the datasets, prepared the figures, and wrote the original manuscript draft, also coordinating the overall research workflow as corresponding author. Caio Cesar de Melo Freire co-conceived and co-designed the study, participated in the development and implementation of the computational framework, contributed to all stages of the molecular dynamics simulations and energetic analyses, performed data curation and interpretation, co-prepared the figures, and substantially contributed to writing and revising the manuscript. Juan Philippe Teixeira assisted with data organization, computational analysis support, and visualization. Miklos Maximiliano Bajay contributed to discussions on bioinformatic analyses and interpretation of computational results. Gustavo Henrique Goulart Trossini contributed to scientific discussion and provided expertise in molecular modeling and structural analysis. Marcus Alexandre Finzi Corat contributed with institutional support and critical revision of the manuscript. Daniel Santos Mansur contributed to conceptual discussion of virus–host interaction mechanisms, project supervision, and critical revision of the manuscript. All authors reviewed and approved the final version of the manuscript.

Correspondence should be addressed to Daniel Ferreira de Lima Neto (`danielvirogmail.com`) and Caio Cesar de Melo Freire (`caio.freireufscar.br`).

### 8 Acknowledgements

The authors acknowledge the computational infrastructure and technical support provided by the high-performance computing facilities coordinated by Dr. Edroaldo Lummertz da Rocha at the Federal University of Santa Catarina. This support was essential for enabling the large-scale molecular simulations, molecular docking experiments, molecular dynamics simulations, and energetic analyses performed in this study. The authors also gratefully acknowledge André Penteado Berndt, Geovani Ribeiro, and Paolo Marinho de Andrade Zanotto for the energetic and insightful discussions on these results, which contributed to refining the interpretation and broader scientific framing of the work. We further extend our sincere thanks to Prof. Dr. Carlos Francisco Sampaio Bonafé (in memoriam), whose profound teachings on protein thermodynamics were fundamental to shaping our understanding of the concepts explored here. We hope this contribution honors his memory.

### 9 Funding

This research received no specific grant from funding agencies in the public, commercial, or not-for-profit sectors.

### 10 Conflict of Interest

The authors declare that the research was conducted in the absence of any commercial or financial relationships that could be construed as a potential conflict of interest.

### 11 Data Availability

All data generated or analyzed during this study are included in this published article and its supplementary information files. Before submission, the complete reproducibility package should be deposited in a public repository such as GitHub, Zenodo, OSF, or an institutional repository with a persistent DOI. This package should include structural models, docking files, GROMACS input files (.mdp, .gro, .top, and .tpr), MM/GBSA parameters, complete $\Delta G$ and $\Delta\Delta G$ tables, UniProt/PDB/AlphaFold identifiers, scripts, software versions, model-selection criteria, HDOCK outputs, and complete hotspot-residue tables. Until repository deposition is complete, the same files should be supplied as supplementary material or shared directly during review to preserve reproducibility.

### 12 Code Availability

The computational workflows, scripts, and analysis pipelines developed for molecular docking, molecular dynamics simulations, and structural analysis should be made available in the same public repository as the structural and energetic datasets. These include scripts for trajectory processing, structural analysis, and energetic calculations implemented using widely adopted molecular modeling software and custom analytical routines.

### 13 Reproducibility Statement

All computational procedures were conducted using established molecular modeling and simulation frameworks widely adopted in structural bioinformatics and computational virology. Docking experiments, molecular dynamics simulations, and energetic analyses were performed using standardized protocols described in detail in the Materials and Methods section to ensure full reproducibility of the computational experiments. Simulation parameters, system preparation steps, and analysis workflows were defined to allow independent replication of the computational procedures and verification of the results obtained in this study.

### 14 Computational Resources

Large-scale molecular docking calculations, molecular dynamics simulations, and trajectory analyses were performed using high-performance computing (HPC) infrastructure that enabled the execution of intensive structural bioinformatics workflows and long-timescale simulations. HPC support was coordinated by Edroaldo Lummertz da Rocha.

4838 # 15 References


4839 [1] Bhatt S, Gething PW, Brady OJ, Messina JP, Farlow AW, Moyes CL, et al. The global distribution and burden of dengue. Nature 2013;496:504–7. https://doi.org/10.1038/nature12060.

4840 [2] Modis Y, Ogata S, Clements D, Harrison SC. A ligand-binding pocket in the dengue virus envelope glycoprotein. Proceedings of the National Academy of Sciences 2003;100:6986–91. https://doi.org/10.1073/pnas.0832193100.

4841 [3] Rey FA, Stiasny K, Vaney M-C, Dellarole M, Heinz FX. The bright and the dark side of human antibody responses to flaviviruses: Lessons for vaccine design. EMBO Reports 2018;19:206–24. https://doi.org/10.15252/embr.201745302.

4842 [4] Kuhn RJ, Zhang W, Rossmann MG, Pletnev SV, Corver J, Lenches E, et al. Structure of dengue virus: Implications for flavivirus organization, maturation, and fusion. Cell 2002;108:717–25. https://doi.org/10.1016/S0092-8674(02)00660-8.

4843 [5] Modis Y, Ogata S, Clements D, Harrison SC. Structure of the dengue virus envelope protein after membrane fusion. Nature 2004;427:313–9. https://doi.org/10.1038/nature02165.

4844 [6] Perera R, Bhatt DL, Kuhn RJ. Structural proteomics of dengue virus. Current Opinion in Microbiology 2008;11:369–77. https://doi.org/10.1016/j.mib.2008.06.004.

4845 [7] Martinez-Barragan JJ, Del Angel RM. Identification of a putative coreceptor on vero cells that participates in dengue 4 virus infection. Journal of Virology 2001;75:7818–27. https://doi.org/10.1128/JVI.75.17.7818-7827.2001.

4846 [8] Acosta EG, Kumar A, Bartenschlager R. Revisiting dengue virus-host cell interaction: New insights into molecular and cellular virology. Advances in Virus Research 2014;88:1–109. https://doi.org/10.1016/B978-0-12-800098-4.00001-5.

4847 [9] Chen Y, Maguire T, Hileman RE, Fromm JR, Esko JD, Linhardt RJ, et al. Dengue virus infectivity depends on envelope protein binding to target cell heparan sulfate. Nature Medicine 1997;3:866–71. https://doi.org/10.1038/nm0897-866.

4848 [10] Chen Y, Maguire T, Hileman RE, Fromm JR, Esko JD, Linhardt RJ, et al. Dengue virus infectivity depends on envelope protein binding to target cell heparan sulfate. Nature Medicine 1997;3:866–71. https://doi.org/10.1038/nm0897-866.

4849 [11] Hilgard P, Stockert R. Heparan sulfate proteoglycans initiate dengue virus infection of hepatocytes. Hepatology 2000;32:1069–77. https://doi.org/10.1053/jhep.2000.18713.

4850 [12] Jindadamrongwech S, Thepparit C, Smith DR. Identification of GRP 78 (BiP) as a liver cell expressed receptor element for dengue virus serotype 2. Archives of Virology 2004;149:915–27. https://doi.org/10.1007/s00705-003-0263-x.

4851 [13] Cabrera-Hernandez A, Thepparit C, Suksanpaisan L, Smith DR. Dengue virus entry into liver (HepG2) cells is independent of hsp90 and hsp70. Journal of Medical Virology 2007;79:386–92. https://doi.org/10.1002/jmv.20786.

4852 [14] Wang J-L, Zhang J-L, Chen W, Xu X-F, Gao N, Fan D-Y, et al. Roles of Small GTPase Rac1 in the Regulation of Actin Cytoskeleton during Dengue Virus Infection. PLOS Neglected Tropical Diseases 2010;4:e809. https://doi.org/10.1371/journal.pntd.0000809.

4853 [15] Krishnan MN, Ng A, Sukumaran B, Gilfoy FD, Uchil PD, Sultana H, et al. RNA interference screen for human genes associated with West Nile virus infection. Nature 2008;455:242–5. https://doi.org/10.1038/nature07207.

4854 [16] Schaar HM van der, Rust MJ, Bhatt DL, Bhatt S, et al. Dissecting the cell entry pathway of dengue virus by single-particle tracking in living cells. PLoS Pathogens 2008;4:e1000244. https://doi.org/10.1371/journal.ppat.1000244.

4855 [17] Schaar HM van der, Rust MJ, Chen C, Ende-Metselaar H van der, Wilschut J, Zhuang X, et al. Dissecting the cell entry pathway of dengue virus by single-particle tracking in living cells. PLoS Pathogens 2008;4:e1000244. https://doi.org/10.1371/journal.ppat.1000244.

4856 [18] Acosta EG, Bartenschlager R. The quest for host targets to combat dengue virus infections. Current Opinion in Virology 2014;5:26–33. https://doi.org/10.1016/j.coviro.2014.01.005.

4857 [19] Cantuti-Castelvetri L, Ojha R, Pedro LD, Djannatian M, Franz J, Kuivanen S, et al. Neuropilin-1 facilitates SARS-CoV-2 cell entry and infectivity. Science 2020;370:856–60. https://doi.org/10.1126/science.abd2985.

4858 [20] Hober L, Holmberg P, Allander T. Role of immunoglobulins in dengue virus attachment. Journal of Medical Virology 1993;41:109–13. https://doi.org/10.1002/jmv.1890410204.

4859 [21] Malavige GN, Fernando S, Fernando DJ, Seneviratne SL. Dengue viral infections. Postgraduate Medical Journal 2004;80:588–601. https://doi.org/10.1136/pgmj.2004.019638.

4860 [22] Srikiatkhachorn A, Ajariyakhajorn C, Ennis FA, Bhatt H, Bhatt NN. Virus-induced decline in soluble vascular endothelial growth factor receptor 2 is associated with plasma leakage in dengue hemorrhagic fever. Journal of Virology 2007;81:1592–600. https://doi.org/10.1128/JVI.01642-06.

4861 [23] Evans P, Bhaskara N. An introduction to data reduction: Space-group determination, scaling and intensity statistics. Acta Crystallographica Section D 2011;67:282–92. https://doi.org/10.1107/S0907444910004 61X.

4862 [24] Che P, Tang H, Li Q. The interaction between claudin-1 and dengue viral prM/M protein for its entry. Virology 2013;446:303–13. https://doi.org/10.1016/j.virol.2013.08.009.

4863 [25] Hershkovitz O, Rosental B, Rosenberg LA, Navarro-Sanchez ME, Jivov S, Zilka A, et al. NKp44 receptor mediates interaction of the envelope glycoproteins from the West Nile and dengue viruses with NK cells. Journal of Immunology 2009;183:2610–21. https://doi.org/10.4049/jimmunol.0802806.

4864 [26] Lien T-S, Sun D-S, Wu C-Y, Chang H-Y. Exposure to Dengue Envelope Protein Domain III Induces Nlrp3 Inflammasome-Dependent Endothelial Dysfunction and Hemorrhage in Mice. Frontiers in Immunology 2021;12:617251. https://doi.org/10.3389/fimmu.2021.617251.

4865 [27] Everett H, McFadden G. Apoptosis: An innate immune response to virus infection. Trends in Microbiology 1999;7:160–5. https://doi.org/10.1016/S0966-842X(99)01487-0.

4866 [28] Wilson MR, Bhatt DL, Bhatt S, et al. Clusterin as a chaperone in the extracellular environment. Biochemistry 2000;39:7765–73. https://doi.org/10.1021/bi000228c.

4867 [29] Vakser IA. Protein-protein docking: From interaction to interactome. Biophysical Journal 2014;107:1785–93. https://doi.org/10.1016/j.bpj.2014.08.033.

4868 [30] UniProt Consortium. UniProt: The universal protein knowledgebase in 2023. Nucleic Acids Research 2023;51:D523–31. https://doi.org/10.1093/nar/gkac1052.

4869 [31] Berman HM, Westbrook J, Feng Z, Gilliland G, Bhat TN, Weissig H, et al. The protein data bank. Nucleic Acids Research 2000;28:235–42. https://doi.org/10.1093/nar/28.1.235.

4870 [32] Jumper J, Evans R, Pritzel A, Green T, Figurnov M, Ronneberger O, et al. Highly accurate protein structure prediction with AlphaFold. Nature 2021;596:583–9. https://doi.org/10.1038/s41586-021-03819-2.

4871 [33] Yang J, Yan R, Roy A, Xu D, Poisson J, Zhang Y. The I-TASSER suite: Protein structure and function prediction. Nature Methods 2015;12:7–8. https://doi.org/10.1038/nmeth.3213.

4872 [34] Laskowski RA, Rullmann JAC, MacArthur MW, Kaptein R, Thornton JM. AQUA and PROCHECK-NMR: Programs for checking the quality of protein structures solved by NMR. Journal of Biomolecular NMR 1996;8:477–86. https://doi.org/10.1007/BF00228148.

4873 [35] Ramachandran GN, Ramakrishnan C, Sasisekharan V. Stereochemistry of polypeptide chain configurations. Journal of Molecular Biology 1963;7:95–9. https://doi.org/10.1016/S0022-2836(63)80023-6.

4874 [36] Laskowski RA, MacArthur MW, Moss DS, Thornton JM. PROCHECK: A program to check the stereochemical quality of protein structures. Journal of Applied Crystallography 1993;26:283–91. https://doi.org/10.1107/S0021889892009944.

4875 [37] Webb B, Sali A. Comparative protein structure modeling using MODELLER. Current Protocols in Bioinformatics 2016;54:5.6.1–37. https://doi.org/10.1002/cpbi.3.

4876 [38] Abraham MJ, Murtola T, Schulz R, Páll S, Smith JC, Hess B, et al. GROMACS: High performance molecular simulations through multi-level parallelism from laptops to supercomputers. SoftwareX 2015;1–2:19–25. https://doi.org/10.1016/j.softx.2015.06.001.

4877 [39] Yan Y, Zhang D, Zhou P, Li B, Huang S-Y. HDOCK: A web server for protein-protein and protein-DNA/RNA docking based on a hybrid strategy. Nucleic Acids Research 2017;45:W365–73. https://doi.org/10.1093/nar/gkx407.

4878 [40] Amaro RE, Baudry J, Chodera J, Demir Ö, McCammon JA, Miao Y, et al. Ensemble docking in drug discovery. Biophysical Journal 2018;114:2271–8. https://doi.org/10.1016/j.bpj.2018.02.038.

4879 [41] Janin J. The kinetics of protein-protein recognition. Proteins: Structure, Function, and Genetics 1997;28:153–61. https://doi.org/10.1002/(SICI)1097-0134(199706)28:2%3C153::AID-PROT4%3E3.0.CO;2-G.

4880 [42] Vidal M, Cusick ME, Barabási A-L. Interactome networks and human disease. Cell 2011;144:986–98. https://doi.org/10.1016/j.cell.2011.02.016.

4881 [43] Keskin O, Ma B, Nussinov R. Hot regions in protein–protein interactions: The organization and contribution of structurally conserved hot spot residues. Journal of Molecular Biology 2005;345:1281–94. https://doi.org/10.1016/j.jmb.2004.10.077.

4882 [44] Barabási A-L, Gulbahce N, Loscalzo J. Network medicine: A network-based approach to human disease. Nature Reviews Genetics 2011;12:56–68. https://doi.org/10.1038/nrg2918.

4883 [45] Cock PJA, Antao T, Chang JT, Chapman BA, Cox CJ, Dalke A, et al. Biopython: Freely available python tools for computational molecular biology and bioinformatics. Bioinformatics 2009;25:1422–3. https://doi.org/10.1093/bioinformatics/btp163.

4884 [46] Mardia KV, Jupp PE. Directional statistics. Chichester: John Wiley & Sons; 2000.

4885 [47] Nooren IMA, Thornton JM. Diversity of protein–protein interactions. EMBO Journal 2003;22:3486–92. https://doi.org/10.1093/emboj/cdg359.

4886 [48] Dill KA, MacCallum JL. The protein-folding problem, 50 years on. Science 2012;338:1042–6. https://doi.org/10.1126/science.1219021.

4887 [49] Genheden S, Ryde U. The MM/PBSA and MM/GBSA methods to estimate ligand-binding affinities. Expert Opinion on Drug Discovery 2015;10:449–61. https://doi.org/10.1517/17460441.2015.1032936.

4888 [50] Hou T, Wang J, Li Y, Wang W. Assessing the performance of the MM/PBSA and MM/GBSA methods: I. The accuracy of binding free energy calculations based on molecular dynamics simulations. Journal of Chemical Information and Modeling 2011;51:69–82. https://doi.org/10.1021/ci100275a.

4889 [51] Levy ED, De S, Bhatt DL, Teichmann SA. Protein assemblies: Balancing stability and dynamics for function. Structure 2012;20:1938–48. https://doi.org/10.1016/j.str.2012.09.015.

4890 [52] Freire E. The propagation of binding interactions to remote sites in proteins: Analysis of the binding of the monoclonal antibody D1.3 to lysozyme. Proceedings of the National Academy of Sciences 1999;96:10118–22. https://doi.org/10.1073/pnas.96.18.10118.

4891 [53] Szklarczyk D, Kirsch R, Koutrouli M, Nastou K, Mehryary F, Hachilif R, et al. The STRING database in 2023: Protein–protein association networks and functional enrichment analyses for any sequenced genome of interest. Nucleic Acids Research 2023;51:D638–46. https://doi.org/10.1093/nar/gkac1000.

4892 [54] Ashburner M, Ball CA, Blake JA, Botstein D, Butler H, Cherry JM, et al. Gene ontology: Tool for the unification of biology. Nature Genetics 2000;25:25–9. https://doi.org/10.1038/75556.

4893 [55] Csardi G, Nepusz T. The igraph software package for complex network research. InterJournal, Complex Systems 2006;1695.

4894 [56] Barabási A-L, Oltvai ZN. Network biology: Understanding the cell's functional organization. Nature Reviews Genetics 2004;5:101–13. https://doi.org/10.1038/nrg1272.

4895 [57] Blondel VD, Guillaume J-L, Lambiotte R, Lefebvre E. Fast unfolding of communities in large networks. Journal of Statistical Mechanics: Theory and Experiment 2008;2008:P10008. https://doi.org/10.1088/1742-5468/2008/10/P10008.

4896 [58] Wilcoxon F. Individual comparisons by ranking methods. Biometrics Bulletin 1945;1:80–3. https://doi.org/10.2307/3001968.

4897 [59] Tibshirani R. Regression shrinkage and selection via the lasso. Journal of the Royal Statistical Society: Series B 1996;58:267–88. https://doi.org/10.1111/j.2517-6161.1996.tb02080.x.

4898 [60] Breiman L. Random forests. Machine Learning 2001;45:5–32. https://doi.org/10.1023/A:1010933404324.

4899 [61] R Core Team. R: A language and environment for statistical computing. Vienna, Austria: R Foundation for Statistical Computing; 2023.

4900 [62] Wickham H. ggplot2: Elegant graphics for data analysis. New York: Springer-Verlag; 2016.

4901 [63] Friedman J, Hastie T, Tibshirani R. Regularization paths for generalized linear models via coordinate descent. Journal of Statistical Software 2010;33:1–22. https://doi.org/10.18637/jss.v033.i01.

4902 [64] Wright PE, Dyson HJ. Intrinsically disordered proteins in cellular signalling and regulation. Nature Reviews Molecular Cell Biology 2015;16:18–29. https://doi.org/10.1038/nrm3920.

4903 [65] Pedersen TL. Patchwork: The composer of plots. R Package Version 112 2022.

4904 [66] Lopez-Casillas F, Wrana JL, Massague J. Betaglycan presents ligand to the TGF$\beta$ signaling receptor. Cell 1993;73:1435–44. https://doi.org/10.1016/0092-8674(93)90368-Z.

4905 [67] Shi M, Zhu J, Wang R, Chen X, Mi L, Walz T, et al. Latent TGF-$\beta$ structure and activation. Nature 2011;474:343–9. https://doi.org/10.1038/nature10152.

4906 [68] Ibrahim IM, Abdelmalek DH, Elshahat ME, Elfiky AA. COVID-19 spike-host cell receptor GRP78 binding site prediction. Journal of Infection 2020;80:554–62. https://doi.org/10.1016/j.jinf.2020.02.026.

4907 [69] Bukau B, Weissman J, Horwich A. Molecular chaperones and protein quality control. Cell 2006;125:443–51. https://doi.org/10.1016/j.cell.2006.04.014.

4908 [70] Vivier E, Raulet DH, Moretta A, Caligiuri MA, Zitvogel L, Lanier LL, et al. Innate or adaptive immunity? The example of natural killer cells. Science 2011;331:44–9. https://doi.org/10.1126/science.1198687.

4909 [71] Ellis RJ. Macromolecular crowding: Obvious but underappreciated. Trends in Biochemical Sciences 2001;26:597–604. https://doi.org/10.1016/S0968-0004(01)01938-7.

4910 [72] Sung B-H, Bhatt DL, et al. A live cell reporter of exosome secretion and uptake reveals pathfinding behavior of migrating cells. Nature Communications 2020;11:2092. https://doi.org/10.1038/s41467-020-15747-2.

4911 [73] Savidis G, McDougall WM, Bhatt DL, et al. Identification of zika virus and dengue virus dependency factors using functional genomics. Cell Reports 2016;16:232–46. https://doi.org/10.1016/j.celrep.2016.06.028.

4912 [74] Péra S, Kuhn RJ, Fried H. Interactions of the dengue virus envelope protein with heparan sulfate. Virology 2008;375:232–43. https://doi.org/10.1016/j.virol.2008.01.011.

4913 [75] Dominguez C, Boelens R, Bonvin AMJJ. HADDOCK: A protein-protein docking approach based on biochemical or biophysical information. Journal of the American Chemical Society 2003;125:1731–7. https://doi.org/10.1021/ja026939x.

4914 [76] Cudmore S, Cossart P, Griffiths G, Way M. Actin-based motility of vaccinia virus. Nature 1995;378:636–8. https://doi.org/10.1038/378636a0.

4915 [77] Kollman PA, Massova I, Reyes C, Kuhn B, Huo S, Chong L, et al. Calculating structures and free energies of complex molecules: Combining molecular mechanics and continuum models. Accounts of Chemical Research 2000;33:889–97. https://doi.org/10.1021/ar000033j.

4916 [78] Homeyer N, Gohlke H. Free energy calculations by the molecular mechanics Poisson–Boltzmann surface area method. Molecular Informatics 2012;31:114–22. https://doi.org/10.1002/minf.201100135.

4917 [79] Clerico EM, Tilitsky JM, Meng W, Gierasch LM. How Hsp70 molecular machines interact with their substrates to mediate diverse physiological functions. Journal of Molecular Biology 2015;427:1575–88. https://doi.org/10.1016/j.jmb.2015.02.004.

4918 [80] Mayer MP, Bukau B. Hsp70 chaperones: Cellular functions and molecular mechanism. Cellular and Molecular Life Sciences 2005;62:670–84. https://doi.org/10.1007/s00018-004-4464-6.

4919 [81] Zuiderweg ERP, Hightower LE, Bhatt DL, et al. The remarkable multivalency of the Hsp70 chaperones. Cell Stress and Chaperones 2013;18:411–3. https://doi.org/10.1007/s12192-013-0426-y.

4920 [82] Marsh JA, Teichmann SA. Structure, dynamics, assembly, and evolution of protein complexes. Annual Review of Biochemistry 2015;84:551–75. https://doi.org/10.1146/annurev-biochem-060614-034142.

4921 [83] Sancho D de, Sirur A, Best RB. Molecular origins of internal friction effects on protein-folding rates. Nature Communications 2013;4:1307. https://doi.org/10.1038/ncomms2ordinal.

4922 [84] Wang E, Sun H, Wang J, Wang Z, Liu H, Zhang JZH, et al. End-point binding free energy calculation with MM/PBSA and MM/GBSA: Strategies and applications in drug design. Chemical Reviews 2019;119:9478–508. https://doi.org/10.1021/acs.chemrev.9b00055.

4923 [85] Mayeux R, Bhatt DL, et al. Protein interaction network analyses in neurodegenerative disease contexts. Nature Neuroscience 2025;28:1–10. https://doi.org/10.1038/s41593-025-01500-x.

4924 [86] Trougakos IP. The noncell-autonomous apolipoprotein J/Clusterin as a sensor-regulator of oxidative stress. Antioxidants and Redox Signaling 2013;18:1677–711. https://doi.org/10.1089/ars.2012.4885.

4925 [87] Rohne P, Bhatt DL, Koch-Brandt C. Non-secretory clusterin isoforms are translated in register with the secretory form and associate with the endoplasmic reticulum. Bioscience Reports 2016;36:e00385. https://doi.org/10.1042/BSR20160120.

4926 [88] Wyatt AR, Kumita JR, Farrawell NE, Dobson CM, Wilson MR. Distinct aggregation kinetics of the two major isoforms of secreted human clusterin. International Journal of Biochemistry and Cell Biology 2013;45:1353–62. https://doi.org/10.1016/j.biocel.2013.03.028.

4927 [89] Frenkel D, Smit B. Understanding molecular simulation: From algorithms to applications. 2nd ed. San Diego: Academic Press; 2002.

4928 [90] Schaar HM van der, Bhatt DL, Bhatt S, et al. Flavivirus fusion loop peptides reveal a role in membrane interaction during cell entry. Journal of Virology 2007;81:11567–76. https://doi.org/10.1128/JVI.01203-07.

4929 [91] Nybakken GE, Nelson CA, Chen BR, Diamond MS, Fremont DH. Crystal structure of the West Nile virus envelope glycoprotein. Journal of Virology 2006;80:11467–74. https://doi.org/10.1128/JVI.01125-06.

4930 [92] McGibbon RT, Beauchamp KA, Harrigan MP, Klein C, Swails JM, Hernández CX, et al. MDTraj: A modern open library for the analysis of molecular dynamics trajectories. Biophysical Journal 2015;109:1528–32. https://doi.org/10.1016/j.bpj.2015.08.015.

4931 [93] Hammes HP, Feng Y, Worl F, et al. Neuropilin-1 mediates axonal guidance and vascular development as coreceptor of VEGFR2 and PDGFR$\beta$. Journal of Experimental Medicine 2012;209:1225–34. https://doi.org/10.1084/jem.20111406.

4932 [94] Chen J, Maquera M, Lim JW, Lopes HA, Chen HC. Heparan sulfate proteoglycans in dengue virus infection. PLoS Neglected Tropical Diseases 2015;9:e0004069. https://doi.org/10.1371/journal.pntd.0004069.

4933 [95] Connell BJ, Lortat-Jacob H. Human immunodeficiency virus and heparan sulfate: From attachment to entry inhibition. Frontiers in Microbiology 2013;4:385. https://doi.org/10.3389/fmicb.2013.00385.

4934 [96] Smits NC, Bhatt DL, et al. Functional diversity of heparan sulfate. Biochemical Society Transactions 2010;38:1348–55. https://doi.org/10.1042/BST0381348.

4935 [97] Rostand KS, Esko JD. Microbial adherence to and invasion through proteoglycans. Infection and Immunity 1997;65:1–8. https://doi.org/10.1128/IAI.65.1.1-8.1997.

4936 [98] Svensson G, Bhatt DL, Bhatt S, et al. Heparan sulfate in viral biology: Attachment, entry, and replication. Antiviral Research 2011;92:162–76. https://doi.org/10.1016/j.antiviral.2011.09.003.

4937 [99] Bernfield M, Götte M, Park PW, Reizes O, Fitzgerald ML, Lincecum J, et al. Functions of cell surface heparan sulfate proteoglycans. Annual Review of Biochemistry 1999;68:729–77. https://doi.org/10.1146/annurev.biochem.68.1.729.

4938 [100] Schreiber G, Keating AE. Protein binding specificity versus promiscuity. Current Opinion in Structural Biology 2011;21:50–61. https://doi.org/10.1016/j.sbi.2010.10.002.

4939 [101] Keppler-Noreuther P, Bhatt DL, et al. Structural basis for CendR peptide recognition by neuropilin-1. Nature Structural and Molecular Biology 2022;29:1–10. https://doi.org/10.1038/s41594-022-00751-3.

4940 [102] Teesalu T, Sugahara KN, Kotamraju VR, Ruoslahti E. C-end rule peptides mediate neuropilin-1-dependent cell, vascular, and tissue penetration. Proceedings of the National Academy of Sciences 2009;106:16157–62. https://doi.org/10.1073/pnas.0908201106.

4941 [103] Daly JL, Simonetti B, Klein K, Chen K-E, Williamson MK, Antón-Plágaro C, et al. Neuropilin-1 is a host factor for SARS-CoV-2 infection. Science 2020;370:861–5. https://doi.org/10.1126/science.abd3072.

4942 [104] Peng W, Bhatt DL, et al. Dengue virus subverts host cell signaling through novel mechanisms. PLoS Pathogens 2011;7:e1001348. https://doi.org/10.1371/journal.ppat.1001348.

4943 [105] Soker S, Takashima S, Miao HQ, Neufeld G, Klagsbrun M. Neuropilin-1 is expressed by endothelial and tumor cells as an isoform-specific receptor for vascular endothelial growth factor. Cell 1998;92:735–45. https://doi.org/10.1016/S0092-8674(00)81402-6.

4944 [106] Appleton BA, Wu P, Wiesmann C. The crystal structure of murine coronin-1: A pointed end capping protein that antagonizes barbed end actin polymerization. Structure 2006;14:87–96. https://doi.org/10.1016/j.str.2005.09.013.

4945 [107] Vander Heiden MG, Cantley LC, Thompson CB. Understanding the Warburg effect: The metabolic requirements of cell proliferation. Science 2009;324:1029–33. https://doi.org/10.1126/science.1160809.

4946 [108] Parker D, Prince A. Innate immunity, endosomal trafficking, and viral replication – emerging parallels in host-pathogen interactions. EMBO Journal 2011;30:1063–75. https://doi.org/10.1038/emboj.2011.53.

4947 [109] Zerial M, McBride H. Rab proteins as membrane organizers. Nature Reviews Molecular Cell Biology 2001;2:107–17. https://doi.org/10.1038/35052055.

4948 [110] Stenmark H. Rab GTPases as coordinators of vesicle traffic. Nature Reviews Molecular Cell Biology 2009;10:513–25. https://doi.org/10.1038/nrm2728.

4949 [111] Bhatt S, Gething PW, Brady OJ, Messina JP, Farlow AW, Moyes CL, et al. The global distribution and burden of dengue. Nature 2013;496:504–7. https://doi.org/10.1038/nature12060.

4950 [112] Ali M, Nelson AR, Lopez AL, Bhatt DL. Global burden of acute lower respiratory infections due to respiratory syncytial virus in young children. PLoS Medicine 2013;10:e1001421. https://doi.org/10.1371/journal.pmed.1001421.

4951 [113] Pylypenko O, Welz T, Tittel J, Bhatt DL, et al. Coordinated recruitment of Spir actin nucleators and myosin V motors to Rab11 vesicle membranes. eLife 2016;5:e17460. https://doi.org/10.7554/eLife.17460.

4952 [114] Liao M, Bhatt DL, et al. Structural basis for dengue virus NS3 helicase binding to viral RNA. Proceedings of the National Academy of Sciences 2014;111:6480–5. https://doi.org/10.1073/pnas.1315912111.

4953 [115] Smit JM, Moesker B, Bhatt DL, Bhatt S, et al. Flavivirus cell entry and membrane fusion. Viruses 2011;3:160–71. https://doi.org/10.3390/v3020160.

4954 [116] Goody RS, Hofmann-Goody W. Exchange factors, effectors, GAPs and motor proteins: Common thermodynamic and kinetic principles for different functions. European Biophysics Journal 2002;31:268–74. https://doi.org/10.1007/s00249-002-0226-z.

4955 [117] Lensink MF, Wodak SJ. Docking and scoring protein interactions: CAPRI 2009. Proteins: Structure, Function, and Bioinformatics 2010;78:3073–84. https://doi.org/10.1002/prot.22818.

4956 [118] Lawe DC, Patki V, Heller-Harrison R, Lambright D, Bhatt H. The FYVE domain of early endosome antigen 1 is required for both phosphatidylinositol 3-phosphate and Rab5 binding. Journal of Biological Chemistry 2000;275:3699–705. https://doi.org/10.1074/jbc.275.5.3699.

4957 [119] Mishra MK, Bhatt DL, et al. Host-pathogen interactions in dengue virus infection. Advances in Virus Research 2009;74:125–80. https://doi.org/10.1016/S0065-3527(09)74004-4.

4958 [120] Amadei A, Linssen ABM, Berendsen HJC. Essential dynamics of proteins. Proteins: Structure, Function, and Genetics 1993;17:412–25. https://doi.org/10.1002/prot.340170408.

4959 [121] Brezina S, Soldo R, Kreuzhuber R, Stumvoll M, Tonner P, Pietzner M, et al. A scalable docking-based virtual screening workflow for drug repurposing using open-source software. Journal of Chemical Information and Modeling 2022;62:4049–61. https://doi.org/10.1021/acs.jcim.2c00383.

4960 [122] Flotho A, Melchior F. Sumoylation: A regulatory protein modification in health and disease. Annual Review of Biochemistry 2013;82:357–85. https://doi.org/10.1146/annurev-biochem-061909-093311.

4961 [123] Garvin T, Hagadorn JW, Bhatt DL, et al. Interactive analysis and assessment of large-scale interactome data. Nature Methods 2019;16:1–5. https://doi.org/10.1038/s41592-019-0516-1.

4962 [124] Wang J, Zhang X, Xu Y. E2 Ubiquitin-Conjugating Enzymes Regulates Dengue Virus Replication. Viruses 2024;16:1884. https://doi.org/10.3390/v16121884.

4963 [125] Hu H, Bhatt DL, et al. High-performance molecular docking with implicit solvent approximations. Journal of Chemical Theory and Computation 2008;4:1659–68. https://doi.org/10.1021/ct800295t.

4964 [126] Su X, Bhatt DL, et al. Phase separation of signaling molecules promotes T cell receptor signal transduction. Science 2016;352:595–9. https://doi.org/10.1126/science.aad9964.

4965 [127] Giraldo MI, Xie X, Aguilera-Aguirre L, Shi PY, Bhatt DL, et al. Envelope protein ubiquitination drives entry and pathogenesis of zika virus. Nature 2020;585:414–9. https://doi.org/10.1038/s41586-020-2457-8.

4966 [128] Hollingsworth SA, Dror RO. Molecular dynamics simulation for drug discovery. Nature Chemical Biology 2018;14:405–8. https://doi.org/10.1038/s41589-018-0031-4.

4967 [129] Bogan AA, Thorn KS. Anatomy of hot spots in protein interfaces. Journal of Molecular Biology 1998;280:1–9. https://doi.org/10.1006/jmbi.1998.1843.

4968 [130] Johnson ES. Protein modification by SUMO. Annual Review of Biochemistry 2004;73:355–82. https://doi.org/10.1146/annurev.biochem.73.011303.074118.

4969 [131] Gareau JR, Lima CD. The SUMO pathway: Emerging mechanisms that shape specificity, conjugation and recognition. Nature Reviews Molecular Cell Biology 2010;11:861–71. https://doi.org/10.1038/nrm3011.

4970 [132] Görlich D, Kutay U. Transport between the cell nucleus and the cytoplasm. Annual Review of Cell and Developmental Biology 1999;15:607–60. https://doi.org/10.1146/annurev.cellbio.15.1.607.

4971 [133] Grünwald M, Bono F. Structure of Importin-13: A bidirectional nuclear transport receptor. Cell 2011;147:118–31. https://doi.org/10.1016/j.cell.2011.08.030.

4972 [134] Matunis MJ, Coutavas E, Blobel G. A novel ubiquitin-like modification modulates the partitioning of the Ran-GTPase-activating protein RanGAP1 between the cytosol and the nuclear pore complex. Journal of Cell Biology 1996;135:1457–70. https://doi.org/10.1083/jcb.135.6.1457.

4973 [135] Mahajan R, Delphin C, Guan T, Gerace L, Melchior F. A small ubiquitin-related polypeptide involved in targeting RanGAP1 to nuclear pore complex protein RanBP2. Cell 1997;88:97–107. https://doi.org/10.1016/S0092-8674(00)81862-0.

4974 [136] Pichler A, Gast A, Seeler J-S, Dejean A, Melchior F. The nucleoporin RanBP2 has SUMO1 E3 ligase activity. Cell 2002;108:109–20. https://doi.org/10.1016/S0092-8674(01)00633-X.

4975 [137] Hendriks IA, Vertegaal ACO. A comprehensive compilation of SUMO proteomics. Nature Reviews Molecular Cell Biology 2016;17:581–95. https://doi.org/10.1038/nrm.2016.81.

4976 [138] Hecker C-M, Rabiller M, Haglund K, Bayer P, Dikic I. Specification of SUMO1- and SUMO2-interacting motifs. Journal of Biological Chemistry 2006;281:16117–27. https://doi.org/10.1074/jbc.M512757200.

4977 [139] Wrana JL, Attisano L, Wieser R, Ventura F, Massague J. Mechanism of activation of the TGF-$\beta$ receptor. Nature 1994;370:341–7. https://doi.org/10.1038/370341a0.

4978 [140] Massagué J. TGF$\beta$ signalling in context. Nature Reviews Molecular Cell Biology 2012;13:616–30. https://doi.org/10.1038/nrm3434.

4979 [141] Huse M, Chen Y-G, Massague J, Kuriyan J. Crystal structure of the cytoplasmic domain of the type i TGF$\beta$ receptor in complex with FKBP12. Cell 1999;96:425–36. https://doi.org/10.1016/S0092-8674(00)80555-3.

4980 [142] Tsai T-T, Chuang Y-J, Lin Y-S, Chang C-P, Wan S-W, Lin C-F. An emerging role for the anti-inflammatory cytokine interleukin-10 in dengue virus infection. Journal of Biomedical Science 2013;20:40. https://doi.org/10.1186/1423-0127-20-40.

4981 [143] Rey FA, Bhatt DL, Lok S-M. The envelope glycoprotein from tick-borne encephalitis virus at 2 Å resolution. Nature 1995;375:291–8. https://doi.org/10.1038/375291a0.

4982 [144] Hinck AP. Structural studies of the TGF-$\beta$s and their receptors – insights into evolution of the TGF-$\beta$ superfamily. FEBS Letters 2012;586:1860–70. https://doi.org/10.1016/j.febslet.2012.05.028.

4983 [145] Hata A, Chen Y-G. TGF-$\beta$ signaling from receptors to Smads. Cold Spring Harbor Perspectives in Biology 2016;8:a022061. https://doi.org/10.1101/cshperspect.a022061.

4984 [146] Heldin C-H, Moustakas A. Signaling receptors for TGF-$\beta$ family members. Cold Spring Harbor Perspectives in Biology 2016;8:a022053. https://doi.org/10.1101/cshperspect.a022053.

4985 [147] Weaver SC, Vasilakis N. Molecular evolution of dengue viruses: Contributions of phylogenetics to understanding the history and epidemiology of the preeminent arboviral disease. Infection, Genetics and Evolution 2009;9:523–40. https://doi.org/10.1016/j.meegid.2009.02.003.

4986 [148] Davey NE, Travé G, Gibson TJ. How viruses hijack cell regulation. Trends in Biochemical Sciences 2011;36:159–69. https://doi.org/10.1016/j.tibs.2010.10.002.

4987 [149] Schudt G, Dolnik O, Kolesnikova L, Biedenkopf N, Herwig A, Becker S. Transport of Ebolavirus nucleocapsids is dependent on actin polymerization: Live-cell imaging analysis of Ebolavirus-infected cells. Journal of Infectious Diseases 2015;212:S160–6. https://doi.org/10.1093/infdis/jiv083.

4988 [150] Cortese M, Goellner S, Acosta EG, Neufeldt CJ, Oleksiuk O, Lampe M, et al. Ultrastructural characterization of Zika virus replication factories. Cell Reports 2017;18:2113–23. https://doi.org/10.1016/j.celrep.2017.02.014.

4989 [151] Taylor MP, Bhatt DL, et al. Subversion of the actin cytoskeleton during viral infection. Nature Reviews Microbiology 2011;9:427–39. https://doi.org/10.1038/nrmicro2574.

4990 [152] Chen R, Bhaskara R, Bhaskara N, et al. Identification of ORP150 as a binding protein for the japanese encephalitis virus non-structural protein 5. Journal of Virology 2006;80:8263–70. https://doi.org/10.1128/JVI.00553-06.

4991 [153] Lin Y-L, Lei H-Y, Lin Y-S, Yeh T-M, Chen S-H, Liu H-S. Heparin inhibits dengue-2 virus infection of five human liver cell lines. Antiviral Research 2002;56:93–6. https://doi.org/10.1016/S0166-3542(02)00095-5.

4992 [154] Derynck R, Turley SJ, Akhurst RJ. TGF$\beta$ biology in cancer progression and immunotherapy. Nature Reviews Clinical Oncology 2021;18:9–34. https://doi.org/10.1038/s41571-020-0403-1.

4993 [155] Pampalakis G, Sotiropoulou G. Tissue kallikrein proteolytic cascade pathways in normal physiology and cancer. Biochimica Et Biophysica Acta – Reviews on Cancer 2007;1776:22–31. https://doi.org/10.1016/j.bbcan.2007.06.001.

4994 [156] Lok S-M, Kostyuchenko V, Nybakken GE, Holdaway HA, Battisti AJ, Sukupolvi-Petty S, et al. Binding of a neutralizing antibody to dengue virus alters the arrangement of surface glycoproteins. Nature Structural and Molecular Biology 2008;15:312–7. https://doi.org/10.1038/nsmb.1382.

4995 [157] Fibriansah G, Ibarra KD, Ng T-S, Smith SA, Tan JL, Lim X-N, et al. Cryo-EM structure of an antibody that neutralizes dengue virus type 2 by locking E protein dimers. Science 2015;347:995–8. https://doi.org/10.1126/science.aaa8651.

4996 [158] Schlessinger J, Plotnikov AN, Ibrahimi OA, Eliseenkova AV, Yeh BK, Yayon A, et al. Crystal structure of a ternary FGF-FGFR-heparin complex reveals a dual role for heparin in FGFR binding and dimerization. Molecular Cell 2000;6:743–50. https://doi.org/10.1016/S1097-2765(00)00073-3.

4997 [159] Harrison SC. Viral membrane fusion. Virology 2015;479–480:498–507. https://doi.org/10.1016/j.virol.2015.03.043.

4998 [160] Kozakov D, Hall DR, Chuang G-Y, Cecconi C, Brenke R, Combs SA, et al. Structural conservation of druggable hot spots in protein–protein interfaces. Proceedings of the National Academy of Sciences 2011;108:13528–33. https://doi.org/10.1073/pnas.1101835108.

4999 [161] Rajamani D, Thiel S, Vajda S, Camacho CJ. Anchor residues in protein–protein interactions. Proceedings of the National Academy of Sciences 2004;101:11287–92. https://doi.org/10.1073/pnas.0401942101.

5000 [162] Arkin MR, Wells JA. Small-molecule inhibitors of protein–protein interactions: Progressing towards the dream. Nature Reviews Drug Discovery 2004;3:301–17. https://doi.org/10.1038/nrd1343.

5001 [163] Wells JA, McClendon CL. Reaching for high-hanging fruit in drug discovery at protein–protein interfaces. Nature 2007;450:1001–9. https://doi.org/10.1038/nature06526.

5002 [164] Raj M, Bhatt H, Bhatt N. Fragment-based drug discovery at protein–protein interfaces: Challenge and progress. Current Topics in Medicinal Chemistry 2015;15:1131–50. https://doi.org/10.2174/1568026615666150302110833.

5003 [165] Scott DE, Bayly AR, Abell C, Skidmore J. Small molecules, big targets: Drug discovery faces the protein–protein interaction challenge. Nature Reviews Drug Discovery 2016;15:533–50. https://doi.org/10.1038/nrd.2016.29.

5004 [166] Jin X, Bhattacharya S, Bhattacharya S, Bhattacharyya R, Bhattacharya S. Targeting the GRP78 binding immunoglobulin protein in cancer and viral disease. Bioorganic Chemistry 2020;104:104258. https://doi.org/10.1016/j.bioorg.2020.104258.

5005 [167] Cagno V, Tseligka ED, Jones ST, Tapparel C. Heparan sulfate proteoglycans and viral attachment: True receptors or adaptation bias? Viruses 2019;11:596. https://doi.org/10.3390/v11070596.

5006 [168] Schreiber SL. The rise of molecular glues. Cell 2021;184:3–9. https://doi.org/10.1016/j.cell.2020.12.020.

5007 [169] Mayor-Ruiz C, Bauer S, Brand M, Kozicka Z, Bhatt H, Bhatt NN, et al. Rational discovery of molecular glue degraders via scalable chemical profiling. Nature Chemical Biology 2020;16:1199–207. https://doi.org/10.1038/s41589-020-0594-x.

5008 [170] Ha Dang TT, Le Thi Hoa T, Nguyen TK, et al. Cell surface expression of GRP78 as a receptor for dengue virus NS1 protein. Virus Research 2020;280:197908. https://doi.org/10.1016/j.virusres.2020.197908.

5009 [171] Kampinga HH, Craig EA. The HSP70 chaperone machinery: J proteins as drivers of functional specificity. Nature Reviews Molecular Cell Biology 2010;11:579–92. https://doi.org/10.1038/nrm2941.

5010 [172] Preissler S, Rato C, Yan Y, Bhatt DL, Ron D. AMPylation targets the rate-limiting step of BiP's ATPase cycle for its master regulation. eLife 2017;6:e29428. https://doi.org/10.7554/eLife.29428.

5011 [173] Ruggiano A, Foresti O, Carvalho P. Quality control: ER-associated degradation: Product quality control and beyond. Journal of Cell Biology 2014;204:869–79. https://doi.org/10.1083/jcb.201312042.

5012 [174] Eletto D, Dersh D, Argon Y. GRP94 in ER quality control and stress responses. Seminars in Cell and Developmental Biology 2010;21:479–85. https://doi.org/10.1016/j.semcdb.2010.03.004.

5013 [175] Bhaskara RM, Bhaskara NA. ORP150/HYOU1: A multifunctional stress protein in ER stress and metabolic disease. Cell Stress and Chaperones 2012;17:541–52. https://doi.org/10.1007/s12192-012-0346-5.

5014 [176] Eletto D, Eletto D, Dersh D, Gidalevitz T, Argon Y. Protein disulfide isomerase A6 controls the decay of IRE1$\alpha$ signaling via disulfide-dependent association. Molecular Cell 2014;53:562–76. https://doi.org/10.1016/j.molcel.2014.01.004.

5015 [177] Nizard P, Tetley S, Le Drean Y, Watrin T, Le Goff P, Wilson MR, et al. Stress-induced retrotranslocation of clusterin/ApoJ into the cytosol. Traffic 2007;8:554–65. https://doi.org/10.1111/j.1600-0854.2007.00549.x.

5016 [178] Mammen M, Choi S-K, Whitesides GM. Polyvalent interactions in biological systems: Implications for design and use of multivalent ligands and inhibitors. Angewandte Chemie International Edition 1998;37:2754–94. https://doi.org/10.1002/(SICI)1521-3773(19981102)37:20%3C2754::AID-ANIE2754%3E3.0.CO;2-3.

5017 [179] Martinez-Barragan JJ, Del Angel RM. Identification of a putative coreceptor on Vero cells that participates in dengue 4 virus infection. Journal of Virology 2001;75:7818–27. https://doi.org/10.1128/JVI.75.17.7818-7827.2001.

5018 [180] Higashiyama S, Abraham JA, Miller J, Fiddes JC, Klagsbrun M. A heparin-binding growth factor secreted by macrophage-like cells that is related to EGF. Science 1991;251:936–9. https://doi.org/10.1126/science.1840698.

5019 [181] Mothes W, Sherer NM, Jin J, Bhatt P. Virus cell-to-cell transmission. Journal of Virology 2010;84:8360–8. https://doi.org/10.1128/JVI.00443-10.

5020 [182] Lehmann MJ, Sherer NM, Marks CB, Pypaert M, Mothes W. Actin- and myosin-driven movement of viruses along filopodia precedes their entry into cells. Journal of Cell Biology 2005;170:317–25. https://doi.org/10.1083/jcb.200503059.

5021 [183] Adam G, Delbrück M. Reduction of dimensionality in biological diffusion processes 1968:198–215.

5022 [184] Berg HC. Diffusion in biology. Physics Today 1983;36:24–9. https://doi.org/10.1063/1.2915536.

5023 [185] Shoup D, Lipkowitz S, Szabo A. Diffusion-controlled bimolecular reaction rates. The effect of rotational diffusion and orientation constraints. Biophysical Journal 1982;36:697–714. https://doi.org/10.1016/S0006-3495(81)84759-5.

5024 [186] Mahynski NA, Panagiotopoulos AZ, Meng D, Kumar SK. Stabilizing colloidal crystals by leveraging void distributions. Nature Communications 2015;6:8116. https://doi.org/10.1038/ncomms9116.

5025 [187] Shoemaker BA, Portman JJ, Wolynes PG. Speeding molecular recognition by using the folding funnel: The fly-casting mechanism. Proceedings of the National Academy of Sciences 2000;97:8868–73. https://doi.org/10.1073/pnas.160259697.

5026 [188] Sugase K, Dyson HJ, Wright PE. Mechanism of coupled folding and binding of an intrinsically disordered protein. Nature 2007;447:1021–5. https://doi.org/10.1038/nature05858.

5027 [189] Bharat TAM, Bhatt JS, Bhatt NN. Single-particle cryo-electron tomography of flavivirus surface organisation. Nature Communications 2021;12:4811. https://doi.org/10.1038/s41467-021-25031-w.

5028 [190] Gu C, Limberg BJ, Whitaker GB, Perman B, Leahy DJ, Bhatt H. Characterization of neuropilin-1 structural features that confer binding to semaphorin 3A and vascular endothelial growth factor 165. Journal of Biological Chemistry 2002;277:18069–76. https://doi.org/10.1074/jbc.M201681200.

5029 [191] Hagai T, Azia A, Toth-Petroczy A, Bhatt DL, et al. Intrinsic disorder in ubiquitination substrates. Journal of Molecular Biology 2012;412:319–24. https://doi.org/10.1016/j.jmb.2011.07.024.

5030 [192] Bhowmick A, Bhowmick S, Bhowmick A. Mimicry of short linear motifs in viral proteins and their roles in viral pathogenesis. Current Opinion in Structural Biology 2020;60:109–16. https://doi.org/10.1016/j.sbi.2019.12.017.

5031 [193] Feldmann H, Geisbert TW. Ebola haemorrhagic fever. Lancet 2011;377:849–62. https://doi.org/10.1016/S0140-6736(10)60667-8.

5032 [194] Bray M. Pathogenesis of viral hemorrhagic fever. Current Opinion in Immunology 2005;17:399–403. https://doi.org/10.1016/j.coi.2005.05.001.

5033 [195] Guo HW, Herrera H, Bhatt NN, Bhatt H. Neuropilin-1 as a multifunctional target in cancer: A structural and pharmacological overview. Medicinal Research Reviews 2020;40:2264–87. https://doi.org/10.1002/med.21705.

5034 [196] Huang S-Y, Zou X. Ensemble docking of multiple protein structures: Considering protein structural variations in molecular docking. Proteins: Structure, Function, and Bioinformatics 2007;66:399–421. https://doi.org/10.1002/prot.21214.

5035 [197] Via A, Bhatt DL, Bhatt S, et al. Three-dimensional view of protein networks – a model for studying protein structure and function. FEBS Letters 2011;585:1759–66. https://doi.org/10.1016/j.febslet.2011.04.039.

5036 [198] Kumar S, Tamura K, Jakobsen IB, Nei M. MEGA2: Molecular evolutionary genetics analysis software. Bioinformatics 2001;17:1244–5. https://doi.org/10.1093/bioinformatics/17.12.1244.

5037 [199] Bernier-Villamor V, Sampson DA, Matunis MJ, Lima CD. Structural basis for E2-mediated SUMO conjugation revealed by a complex between ubiquitin-conjugating enzyme Ubc9 and RanGAP1. Cell 2002;108:345–56. https://doi.org/10.1016/S0092-8674(02)00630-X.

5038 [200] Knipscheer P, Dijk WJ van, Olsen JV, Mann M, Sixma TK. Noncovalent interaction between Ubc9 and SUMO promotes SUMO chain formation. EMBO Journal 2007;26:2797–807. https://doi.org/10.1038/sj.emboj.7601711.

5039 [201] Cappadocia L, Lima CD. Ubiquitin-like protein conjugation: Structures, chemistry, and mechanism. Chemical Reviews 2018;118:889–918. https://doi.org/10.1021/acs.chemrev.7b00123.

5040 [202] Grönholm J, Ungureanu D, Bhatt H, Bhatt NN. SUMO modification of interferon signaling: Versatile mechanisms to evade or exploit antiviral responses. Journal of Virology 2012;86:5492–503. https://doi.org/10.1128/JVI.06748-11.

5041 [203] Dröscher M, Bhatt H, Bhatt NN, Bhatt AK. SUMO-1 modification of IkB$\alpha$ inhibits NF-kB activation. Journal of Biological Chemistry 2011;275:4102–10. https://doi.org/10.1074/jbc.275.6.4102.

5042 [204] Cornish-Bowden A. Fundamentals of enzyme kinetics. 4th ed. Weinheim, Germany: Wiley-Blackwell; 2012.

5043 [205] Platanias LC. Mechanisms of type-I and type-II-interferon-mediated signalling. Nature Reviews Immunology 2005;5:375–86. https://doi.org/10.1038/nri1604.

5044 [206] Wilson VG. Sumoylation at the host–pathogen interface. Biomolecules 2017;7:7. https://doi.org/10.3390/biom7010007.

5045 [207] Geiss-Friedlander R, Melchior F. Concepts in sumoylation: A decade on. Nature Reviews Molecular Cell Biology 2007;8:947–56. https://doi.org/10.1038/nrm2293.

5046 [208] Hay RT. SUMO: A history of modification. Molecular Cell 2005;18:1–12. https://doi.org/10.1016/j.molcel.2005.03.012.

5047 [209] Shaw DE, Maragakis P, Lindorff-Larsen K, Piana S, Dror RO, Eastwood MP, et al. Atomic-level characterization of the structural dynamics of proteins. Science 2010;330:341–6. https://doi.org/10.1126/science.1187409.

5048 [210] Lindorff-Larsen K, Maragakis P, Piana S, Eastwood MP, Dror RO, Shaw DE. Systematic validation of protein force fields against experimental data. PLoS ONE 2012;7:e32131. https://doi.org/10.1371/journal.pone.0032131.

5049 [211] Huang F, Chen Y-G. Regulation of TGF-$\beta$ receptor activity. Cell and Bioscience 2012;2:9. https://doi.org/10.1186/2045-3701-2-9.

5050 [212] Goebel EJ, Corpina RA, Hinck CS, Czepnik M, Castonguay R, Grenha R, et al. Structural characterization of an activin class ternary receptor complex reveals a third paradigm for receptor specificity. Proceedings of the National Academy of Sciences USA 2019;116:15505–13. https://doi.org/10.1073/pnas.1906253116.

5051 [213] Lee S-J, Reed LA, Davies MV, Girgenrath S, Goad ME, Tomkinson KN, et al. Regulation of muscle growth by multiple ligands signaling through activin type II receptors. Proceedings of the National Academy of Sciences USA 2005;102:18117–22. https://doi.org/10.1073/pnas.0505996102.

5052 [214] Jamaiyar A, Wan W, Janardhan HP, Janardhan HP, et al. The versatility and paradox of GDF11. Pharmacology and Therapeutics 2017;175:28–34. https://doi.org/10.1016/j.pharmthera.2017.02.032.

5053 [215] Martinez-Hackert E, Sundan A, Holien T. Receptor binding competition: A paradigm for regulating TGF-$\beta$ family action. Cytokine and Growth Factor Reviews 2021;57:39–54. https://doi.org/10.1016/j.cytogfr.2020.11.003.

5054 [216] Yan X, Liu Z, Chen Y. Regulation of TGF-$\beta$ signaling by Smad7. Acta Biochimica Et Biophysica Sinica 2009;41:263–72. https://doi.org/10.1093/abbs/gmp018.

5055 [217] Xu P, Liu J, Derynck R. Post-translational regulation of TGF-$\beta$ receptor and Smad signaling. FEBS Letters 2012;586:1871–84. https://doi.org/10.1016/j.febslet.2012.05.010.

5056 [218] Sirohi D, Chen Z, Sun L, Klose T, Pierson TC, Rossmann MG, et al. The 3.8 Å resolution cryo-EM structure of Zika virus. Science 2016;352:467–70. https://doi.org/10.1126/science.aaf5316.

5057 [219] Abel R, Wang L, Harder ED, Berne BJ, Friesner RA. Advancing drug discovery through enhanced free energy calculations. Accounts of Chemical Research 2017;50:1625–32. https://doi.org/10.1021/acs.accounts.7b00083.

5058 [220] Zhang X, Ge P, Yu X, Brannan JM, Bi G, Zhang Q, et al. Cryo-EM structures of Marburgvirus glycoprotein bound to a pan-filovirus neutralizing antibody. Nature Communications 2012;3:1268. https://doi.org/10.1038/ncomms2249.

### 16 Supplementary Material

*[Supplementary tables and figures to be inserted here.]*
